# Multi-omic data helps improve prediction of personalised tumor suppressors and oncogenes

**DOI:** 10.1101/2022.01.13.476163

**Authors:** Malvika Sudhakar, Raghunathan Rengaswamy, Karthik Raman

## Abstract

The progression of tumorigenesis starts with a few mutational and structural driver events in the cell. Various cohort-based computational tools exist to identify driver genes but require a large number of samples to produce reliable results. Many studies use different methods to identify driver mutations/genes from mutations that have no impact on tumour progression; however, a small fraction of patients show no mutational events in any known driver genes. Current unsupervised methods map somatic and expression data onto a network to identify the perturbation in the network. Our method is the first machine learning model to classify genes as tumour suppressor gene (TSG), oncogene (OG) or neutral, thus assigning the functional impact of the gene in the patient. In this study, we develop a multi-omic approach, PIVOT (Personalised Identification of driVer OGs and TSGs), to train on experimentally or computationally validated mutational and structural driver events. Given the lack of any gold standards for the identification of personalised driver genes, we label the data using four strategies and, based on classification metrics, show gene-based labelling strategies perform best. We build different models using SNV, RNA, and multi-omic features to be used based on the data available. Our models trained on multi-omic data improved predictions compared to mutation and expression data, achieving an accuracy ≥ 0.99 for BRCA, LUAD and COAD datasets. We show network and expression-based features contribute the most to PIVOT. Our predictions on BRCA, COAD and LUAD cancer types reveal commonly altered genes such as TP53, and PIK3CA, which are predicted drivers for multiple cancer types. Along with known driver genes, our models also identify new driver genes such as PRKCA, SOX9 and PSMD4. Our multi-omic model labels both CNV and mutations with a more considerable contribution by CNV alterations. While predicting labels for genes mutated in multiple samples, we also label rare driver events occurring in as few as one sample. We also identify genes with dual roles within the same cancer type. Overall, PIVOT labels personalised driver genes as TSGs and OGs and also identifies rare driver genes.

PIVOT is available at https://github.com/RamanLab/PIVOT.

## 1 INTRODUCTION

Alterations in the genome drive the progression of cancer (Stratton et al., 2009). Mutations in certain genes, called driver genes, give cancer cells an added growth advantage (Vogelstein et al., 2013). These mutations, as well as other genomic changes, such as copy number variations (CNVs), accumulate as the tumour progresses. The genomic landscape of cancer is complex (Stratton et al., 2009; Vogelstein et al., 2013), with differences between cancer types in the number of mutations observed (Kandoth et al., 2013) or the mutation signatures (Alexandrov et al., 2013b,a). The genes mutated vary between cancer types and within subtypes of cancer. We now know that cells are heterogeneous within the same tumour, and heterogeneity confounds our understanding of the evolution of tumours (Greaves and Maley, 2012; Burrell et al., 2013). Mutational signatures vary in different cancer types (Alexandrov et al., 2013b). These specific patterns of mutations imply the need for cancer-specific driver mutation prediction tools. Both pan-cancer and tissue-specific identification of driver genes are essential for understanding cancer.

Various computational methods exist for identifying driver genes. Tools either classify mutations as driver events (Banerjee et al., 2021; Mao et al., 2013; Tokheim and Karchin, 2019), or the genes mutated as driver genes (Tokheim et al., 2016). Driver mutation prediction relies on the functional impact or neighbourhood sequence. While some tools are specific to cancer (Tokheim and Karchin, 2019), other functional impact-based tools like SIFT (Ng and Henikoff, 2003) or PolyPhen2 (Adzhubei et al., 2010) are not. Some tools are limited in their ability to predict only single nucleotide variations, more specifically missense mutations (Tokheim and Karchin, 2019). Hence, using tools that predict driver mutations to predict personalised driver genes are limited in their scope.

Other tools exist that predict driver genes instead of mutations and use background mutation rate (BMR) or ratio-metric features for prediction. Computational methods using BMR such as MutSigCV (Lawrence et al., 2013) assume higher mutation rates in driver genes when compared to the background mutation rate. BMR-based methods are biased towards driver genes with high mutation frequency (Sudhakar et al., 2022). This shortcoming is overcome by ratio-metric features, which give importance to the functional impact of the mutation on the gene rather than the frequency of mutations (Davoli et al., 2013; Tokheim et al., 2016; Sudhakar et al., 2022). These methods are essential to identify most of the driver genes observed in a cohort but are elusive to rare driver genes.

Methods using the concept of mutual exclusivity of genes overcome the challenge by identifying a set of mutually exclusive genes in samples (Bokhari and Arodz, 2017; Leiserson et al., 2013). Somatic mutation data is used to identify sets of genes that improve coverage for the entire cohort. The mutual exclusivity approach is further improved by including biological knowledge as network information. QuaDMutNetEx (Bokhari and Arodz, 2017) uses biological interactions between proteins to find a set of mutually exclusive genes that perturb a pathway. Along with the mutual exclusivity of genes, the algorithm identifies a set of genes, which the algorithm can map onto the network to form a connected component. While this method may miss out on genes on different complementary pathways, the authors suggest iterating QuaDMutNetEx after excluding previously identified genes to help identify other essential driver genes. Network-based approaches may help identify low mutation frequency genes in a cohort by including biological interactions between proteins. While cohort-based methods help in understanding the biological mechanism of the disease, they are not very useful in a clinical setup. Additionally, a large number of samples are required for cohort-based methods to produce reliable results. They also cannot be used to find very rare driver genes.

While a large number of genomic events in the cancer genome are single nucleotide variations (SNVs), other genomic rearrangements such as CNVs, gene fusions and epigenomic changes are also known to occur. While the above-mentioned methods help identify many genes, many samples remain with no mutations in known driver genes (Campbell and et al., 2020). This implies that rare driver genes are missed out by cohort-based methods. BMR or ratio-metric methods also do not capture the effects of these methods. Another approach to identifying driver genes mutated at very low frequency in samples is to identify *personalised* driver genes, i.e. driver genes for individual samples rather than a cohort. Cohort-based studies rely on a large sample size to identify patterns consistent across samples, while identification of personalised driver genes is especially relevant for sub-types of cancers where large cohort studies are not possible or show very few mutations.

Identification of personalised driver genes helps identify actionable targets in patients without known driver mutations. The methods for the identification of personalised driver genes are based on unsupervised algorithms because we lack the ground truth. Instead, the methods use a network-based algorithm to identify perturbed pathways. The graph, along with somatic alterations and differential gene expression profile of the patient, is used to predict driver genes. DawnRank (Hou and Ma, 2014) and SCS (Guo et al., 2018) use a directed graph with loops for auto-regulation. The directed graph is a collapsed network built using multiple protein-protein interaction (PPI) networks. DawnRank uses a modification of the Page-Rank algorithm to rank genes with downstream perturbed genes, while SCS uses Random Walker with Restart algorithm (RWR). Prodigy (Dinstag and Shamir, 2019) uses network as well as pathway data to identify genes, which deregulate a large number of pathways. The method uses a prize-collecting Steiner tree algorithm to find genes with SNV mutations. All methods identify rare driver genes compared with existing network-based methods for driver gene identification.

Network-based personalised driver gene tools integrate somatic mutation and gene expression data and identify genes using an unsupervised method. A subset of mutations, SNV, are used in PRODIGY though the method can be extended to other mutation types. Further, the functional impact of mutations is ignored when mutation data is converted into presence/absence calls. Data ingested is limited to network, mutation and expression data, though methods like DawnRank also analyse CNV data. With many high throughput multi-omic data available, the prediction of driver genes can be improved by including multi-omic data. Moreover, the expression data is included as differentially expressed genes (DEGs), calculated based on the cohort and not an individual sample. These methods rank the driver genes but do not classify genes as TSG or OG.

In this study, we define a machine learning (ML) classification problem to identify personalised driver genes and address the challenges. We define strategies for labelling genes and identify the best method. We employ features based on mutation, expression, CNV and miRNA expression data and understand their contribution to classification. We finally build mutation, RNA and multi-omic models to identify personalised diver genes and assign functional classes. We classify genes of three TCGA cancer cohorts as TSG or OG for individual samples and identify new driver genes.

## 2 RESULTS

Our method, PIVOT (Personalised Identification of driVer OGs and TSGs), is the first ML-based supervised approach for identifying personalised driver genes to the best of our knowledge. Unlike previous methods that distinguish between driver and non-driver genes, PIVOT can also identify whether it functions as a TSG or OG in the specific cancer type. We build separate models using features extracted from different modalities of -omics data: SNV, gene expression, and multi-omics. The SNV model is trained on mutation data not limited to single nucleotide variation but all mutation types. The multi-omic data integrate SNV, gene expression, CNV, miRNA data, and also network information. Figure 1 shows the different combinations of labelling strategies, data, feature sets and models used. We further identify common cancer domains mutated frequently and find that only a subset of domains contribute to the models. We show that integrating network information with gene expression data improves the overall predictive power of the classification models. Lastly, we observe that our multi-omic model generates better predictions than the models based on SNV and gene expression data alone and can be used to predict novel TSGs and OGs in individual samples.

**Figure 1.**
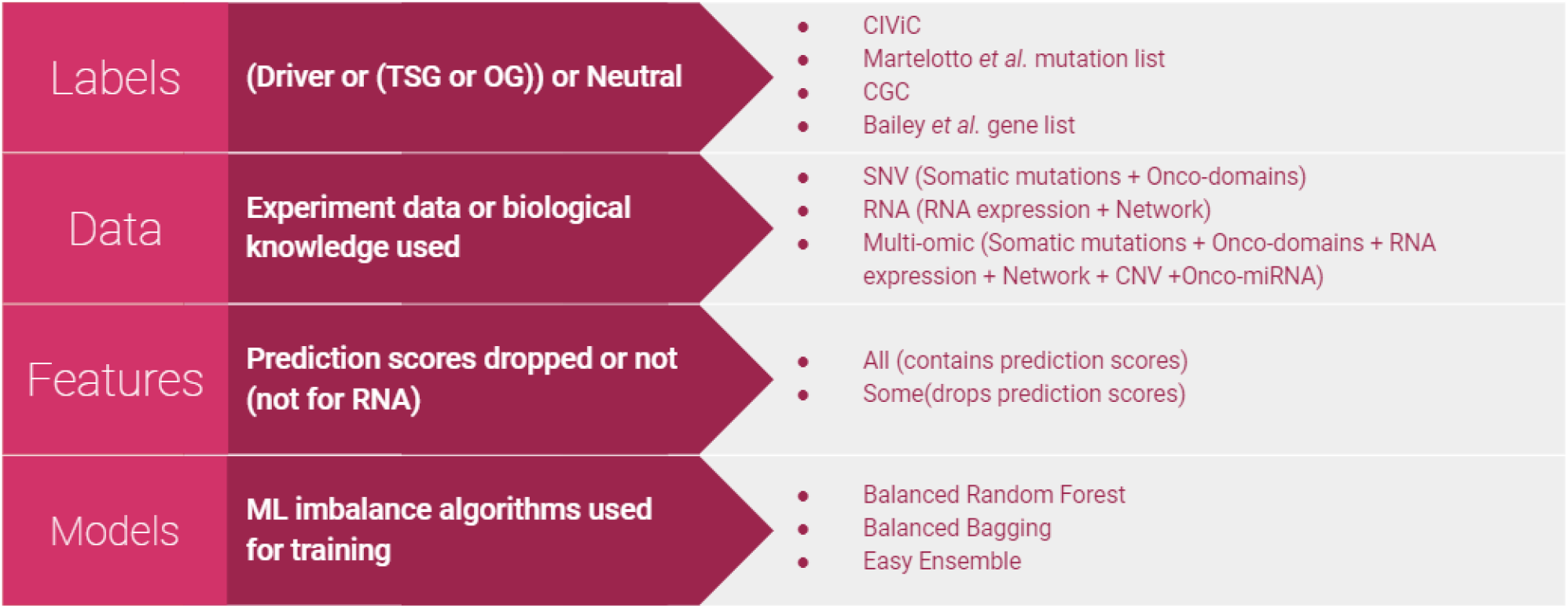
Different types of classifiers. Four different labelling strategies were used to label the altered genes. Models were built using SNV, RNA or mulit-omic data. The number of features used was varied for SNV and multi-omic data to either include or drop features based on prediction scores. For building classifiers, we used algorithms specific for imbalanced data as the number of neutral genes are far higher than than TSGs or OGs. Classifiers are built using all combinations of labels, data, features and models.

### 2.1 Gene-based labelling helps learn better models

One of the significant challenges in formulating a supervised classification problem for driver gene identification is the lack of a labelled gold standard dataset. We define four labeling strategies to assign labels and build models using the SNV data, two at mutation level and two at the gene level. We used the CIViC (Griffith et al., 2017) database and a list of mutations published by Martelotto et al. (2014), to label genes containing known driver mutations as driver or TSG and OG. The models developed show high accuracy for BRCA. The precision and recall of driver classes are ≥ 0.95 (Supplementary Table 1). Although the models perform well, we use a highly curated list of less than 10 genes to train the model. Although we could successfully label all the mutations from the BRCA dataset, both methods fail to label the COAD, LGG and LUAD datasets. Notably, the number of samples in BRCA is twice the number of samples compared to other datasets. We conclude that although mutation labeling-based approaches successfully predict personalised driver genes, they are often limited in their ability to label mutations from all cancer types.

Cancer Gene Census (CGC) (Sondka et al., 2018) is the gold standard database for known driver genes labelled as either TSG and OG. Similarly, the list of cancer-type specific genes published by Bailey et al. (2018) consists of predicted TSG and OG list, which has been manually curated. Datasets labelled using CGC genes show accuracy ≤ 0.73, where the neutral class mainly contributes to the score (Supplementary Table 1). Given the heavy class imbalance, accuracy is not the best metric to judge a model. While the accuracy of the models is low compared to other labelling strategies, the F1-score (harmonic mean of precision and recall) is found to be ≤ 0.52 for TSGs. In some models, the F1 score of the TSG and OG is ≤ 0.20. The list of genes in CGC is not cancer-type specific. Using CGC as the source to label cancer-type specific personalised driver genes results in models that perform well on the training set but not on the test dataset.

The best classification performances for identifying personalized driver genes were obtained using the labels derived from the list of TSGs and OGs published by Bailey et al. The cancer-specific genes obtained using this strategy resulted in a larger training dataset consisting of TSGs and OGs, compared to the mutation-based labelling approach. The method consistently performs well across all four datasets with the best model accuracy of 0.89 (Supplementary Table 1). Across all feature sets used to build the best model, the accuracy is ≥ 0.80 except for the best model for BRCA data using a subset of the SNV features. We conclude that the increase in the size of the training dataset using TSGs and OGs from Bailey et al., and the specificity of the genes used for training contribute to building better models for identifying personalised driver genes.

### 2.2 Mutation data is helpful for identifying personalised genes

We used PIVOT on four TCGA datasets (Breast Cancer: BRCA, Colorectal Adenocarcinoma: COAD, Lower Grade Glioma: LGG and Lung Adenocarcinoma: LUAD) to predict genes as neutral or drivers. Depending on the labelling strategy, driver genes were further classified as TSG or OG. For BRCA, we observed the best accuracy for the model trained on CIViC labels using all SNV features (Supplementary Table 1). The F1 score for the best model was ≥ 0.98 for driver and neutral class. All ML algorithms, balanced bagging, balanced random forest and Easy ensemble gave comparable results. Based on the F1 scores of TSG and OG for the best model in each labelling strategy, we built better models labelled using genes form Martelotto et al., followed by those from Bailey et al. and CGC. Different ML algorithms perform better based on the labelling strategy or the number of features used for prediction, though balanced bagging consistently performed best or close to best.

No data was labelled using CIViC or Martelotto et al. driver genes for the other three datasets (Supplementary Table 1). Models built on data labelled using TSGs and OGs published by Bailey et al consistently performed better than models built on data labelled using CGC (Supplementary Table 1). Further, irrespective of the ML algorithm used, the recall on TSG is always higher than the precision. Predicting OGs using SNV is more straightforward than predicting TSG, as evident from the higher F1 score for OGs compared to the TSGs. The SNV features are primarily based on the scores given by mutation prediction tools that predict the damaging nature of the missense mutation. Since the training dataset mainly consists of missense mutations, it is intuitive that predicting OGs are easier. In general, models learnt on data labelled using Bailey et al. perform well using all the SNV features.

### 2.3 Mutation-based categorical features are not sufficient to predict driver genes

SNV features are based on the functional impact of the mutation, domains mutated, and the prediction scores by various driver or mutation impact predicting tools. These tools cannot score all mutations and are hence dropped while training. We train our models using two feature sets, one that uses all features and the second that uses a subset of these features. The advantage of a smaller feature set is an increased number of training data. Since features with a large number of missing data are dropped, the number of data points dropped because of missing data reduces. The statistics for labelled data used for training and testing is available in Supplementary Table 1. Most SNV features consist of prediction scores and a corresponding categorical feature defining the impact of the mutation. For example, feature *SIFT*_*score*_ consists of a prediction score between 0 and 1, while the corresponding feature *SIFT*_*pred*_ is a categorical feature with “D” defining damaging and “N” for neutral mutations. We cannot impute missing values for *SIFT*_*score*_ and *SIFT*_*pred*_, as the given mutation type might be out of the scope of the tool. Hence in the smaller feature subset, we drop *SIFT*_*pred*_ feature and encode *SIFT*_*pred*_ as an ordinal feature with the lowest value for missing data (explained in Methods).

We ran the SNV models with two feature sets, “all” and “small”, where “small” consists of categorical features. It is to be noted that the number of features in the two feature sets vary between the datasets because of the difference in domains associated with a cancer type. In BRCA, irrespective of the labelling strategy, the models consistently perform better on “all” features (Supplementary Table 1). Since only a subset of features is used, lower scores show that non-categorical features are essential for classification (Supplementary Table 1). For other datasets labelled using Bailey et al. genes, we find that the “small” feature set performs equally well as compared to the “all” feature set (Supplementary Table 1). Feature importance ranking shows that, unlike BRCA, for other datasets, categorical features rank high for both “all” and “small” feature sets, explaining the slight difference in F1 scores (Fig. 2). While score-based features rank in the top 20 of all datasets, these features contribute more to BRCA (Supplementary Table 2).

**Figure 2.**
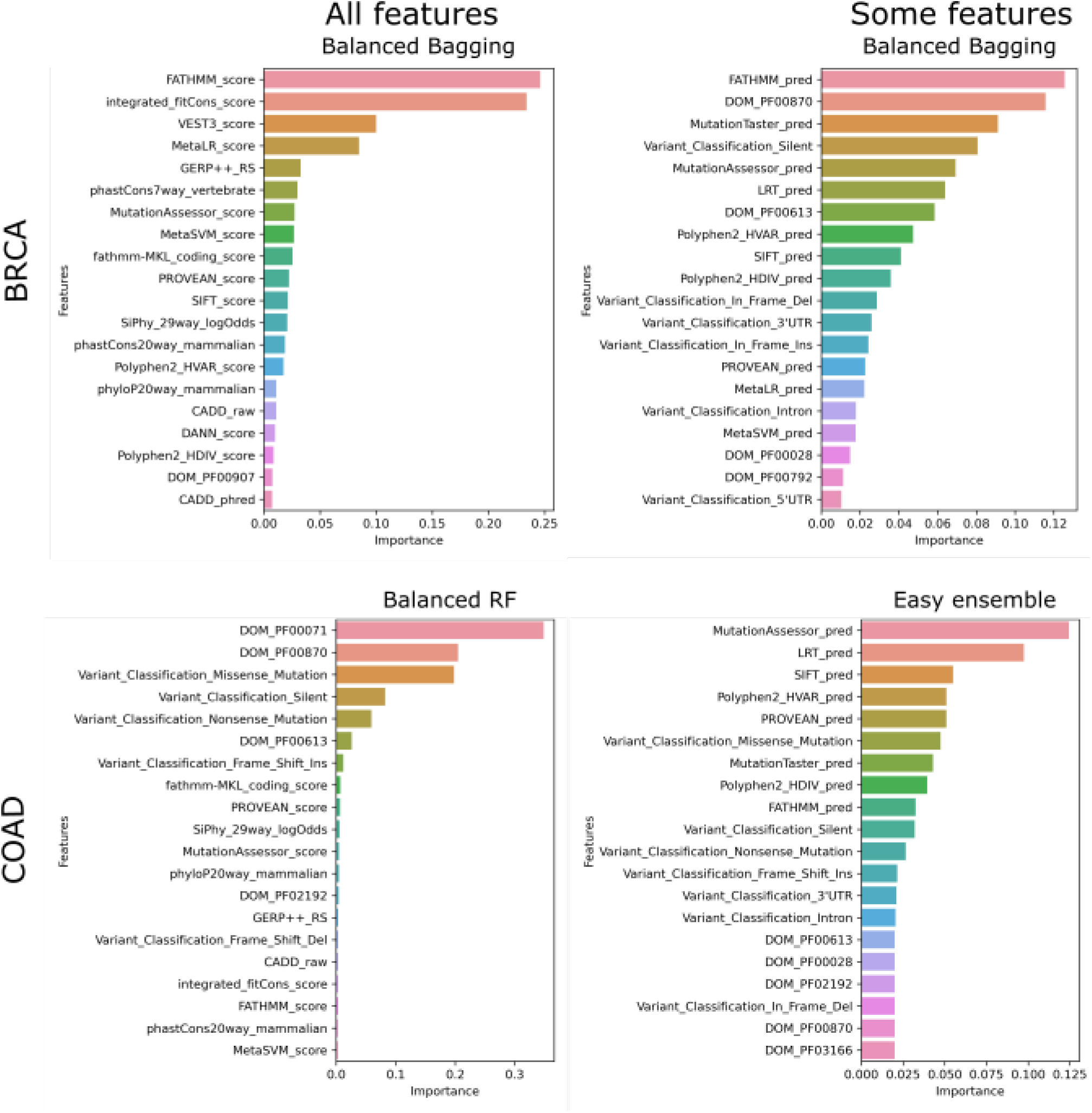
Top features contributing to models. The top20 features and their contribution to the best model is plotted for “all” and “some” feature sets.

### 2.4 Cancer domains used for predicting driver genes

We include onco-domain information into the SNV features. For each cancer type, we define features to capture the presence or absence of mutations in cancer-type specific known onco-domains. Not all domains contribute equally to the classifier’s performance, with a majority having no contribution. We identify the top contributing domain in each cancer type. The subset of domains identified during SNV classification intersects with domain features contributing to multi-omic models (Table 1). BRCA identifies three domains: p53 DNA-binding domain, T-box and cadherin binding domain. T-box is a DNA binding domain used in transcriptional activation/repression roles. TP53 is a known TSG pan-cancer, while cadherins are trans-membrane proteins used for adhesion. Similarly, we identify the top domains for all four cancer types. We find that the p53 DNA-binding domain is the only domain feature identified for all cancer-types by SNV and multi-omic models. The individual contribution of domains to SNV and multi-omic models is listed in Supplementary Table 3.

**Table 1.**
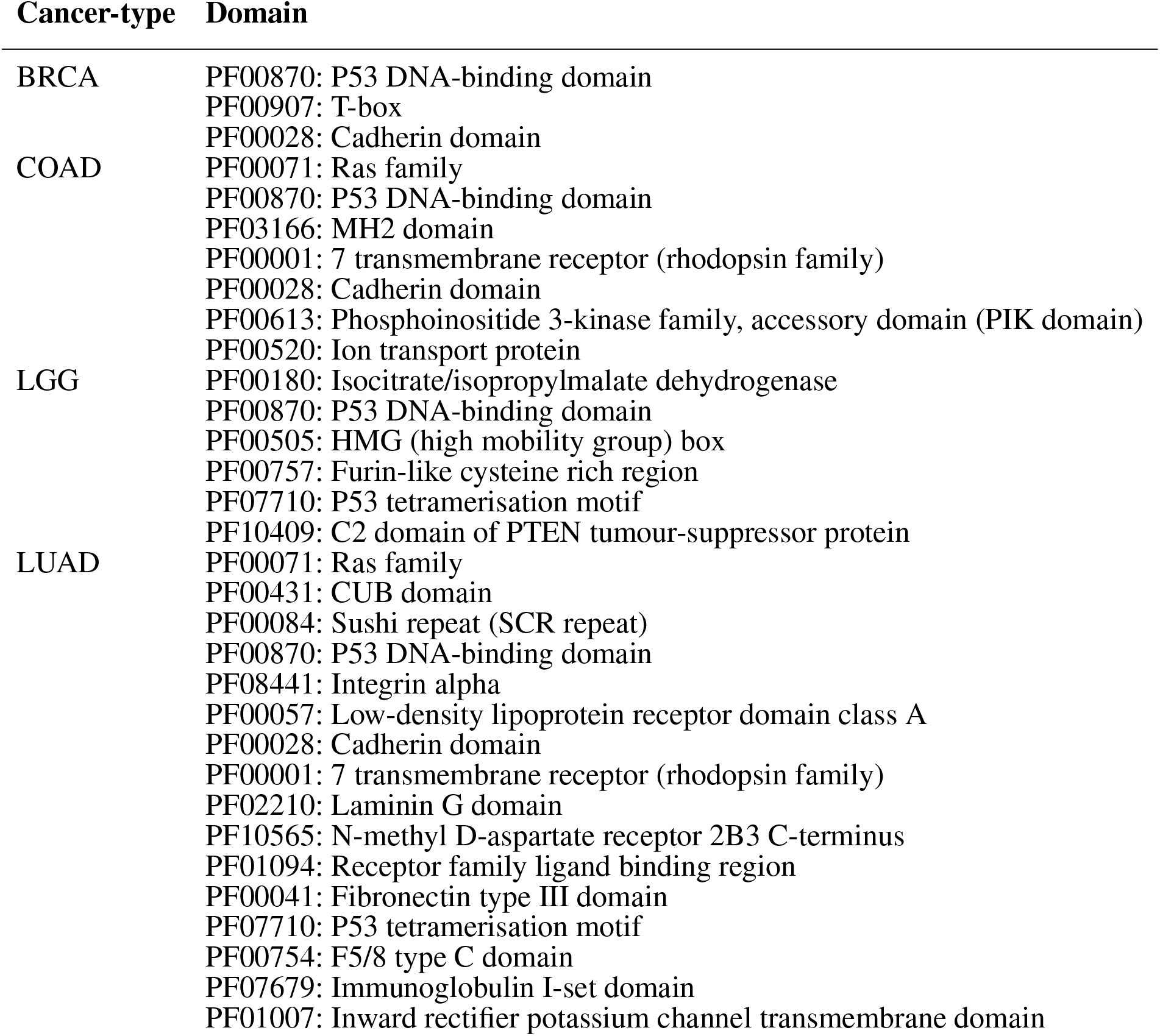
Domains contributing to models for the cancer-type

### 2.5 Expression and network-based features improve prediction accuracy

While mutation data helps classify genes into TSGs and OGs, there is scope for improvement. We used expression data and PPI networks for generating features and predicting personalised driver genes. We built models for BRCA, COAD and LUAD cancer-types using all feasible labelling strategies. In contrast to SNV models, we find ≥ 96% accuracy across the best models for all three cancer types (Supplementary Table 4). The F1 score of OG and TSG is ≥ 94% across all models, irrespective of the labels. An analysis of the features revealed network properties of genes such as closeness centrality and degree are the significant contributors to the identification of TSGs and OGs (Fig. 3). While RNA expression-based features logFC and logCPM contribute, they are not sufficient for classification. Network and RNA expression features improve the classification accuracy above SNV features.

**Figure 3.**
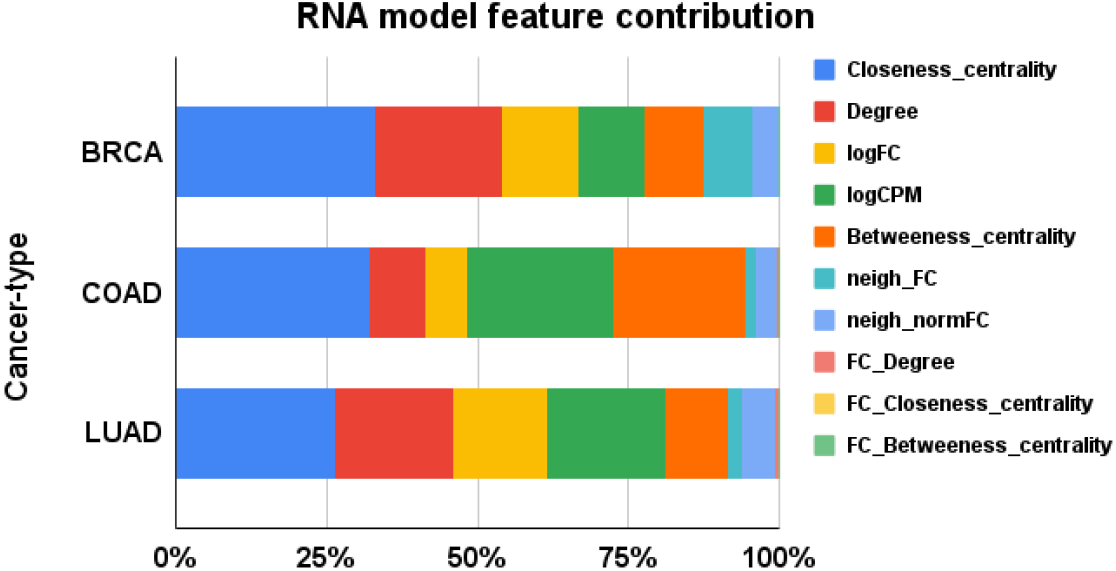
Expression and network-based features contributing to models. For each cancer type, the average contribution of a feature across all models is plotted.

### 2.6 Multi-omic data generate the best classification performances

SNV and RNA models are sufficient to predict the driver genes, but including both improves the prediction. We used features extracted from SNV, RNA features, CNV and miRNA data to build the multi-omic models. The best prediction model uses Balanced Bagging across all cancer types using Bailey et al. genes for labelling (Supplementary Table 5). Unlike the SNV models, the smaller feature subset performs better across all cancer types. The model accuracy is 1.0 for BRCA, with an F1 score of 0.99 for OG and 0.97 for TSG. For COAD, we observe an accuracy of 0.99 with an F1 score of 0.97 and 0.94 for OG and TSG, respectively. Similarly, for the LUAD cancer type, we achieve an accuracy of 0.99 and an F1 score of 0.97 and 0.94 for OG and TSG respectively. Detailed classification metrics for all multi-omic models are available as Supplementary Table 5. Feature importance shows network and RNA expression are top-ranking features (Fig. 4) among other multi-omic features (Supplementary Table 2). Along with CNV features, we also find some miRNA features contributing to the overall classification performance (Supplementary Table 9).

**Figure 4.**
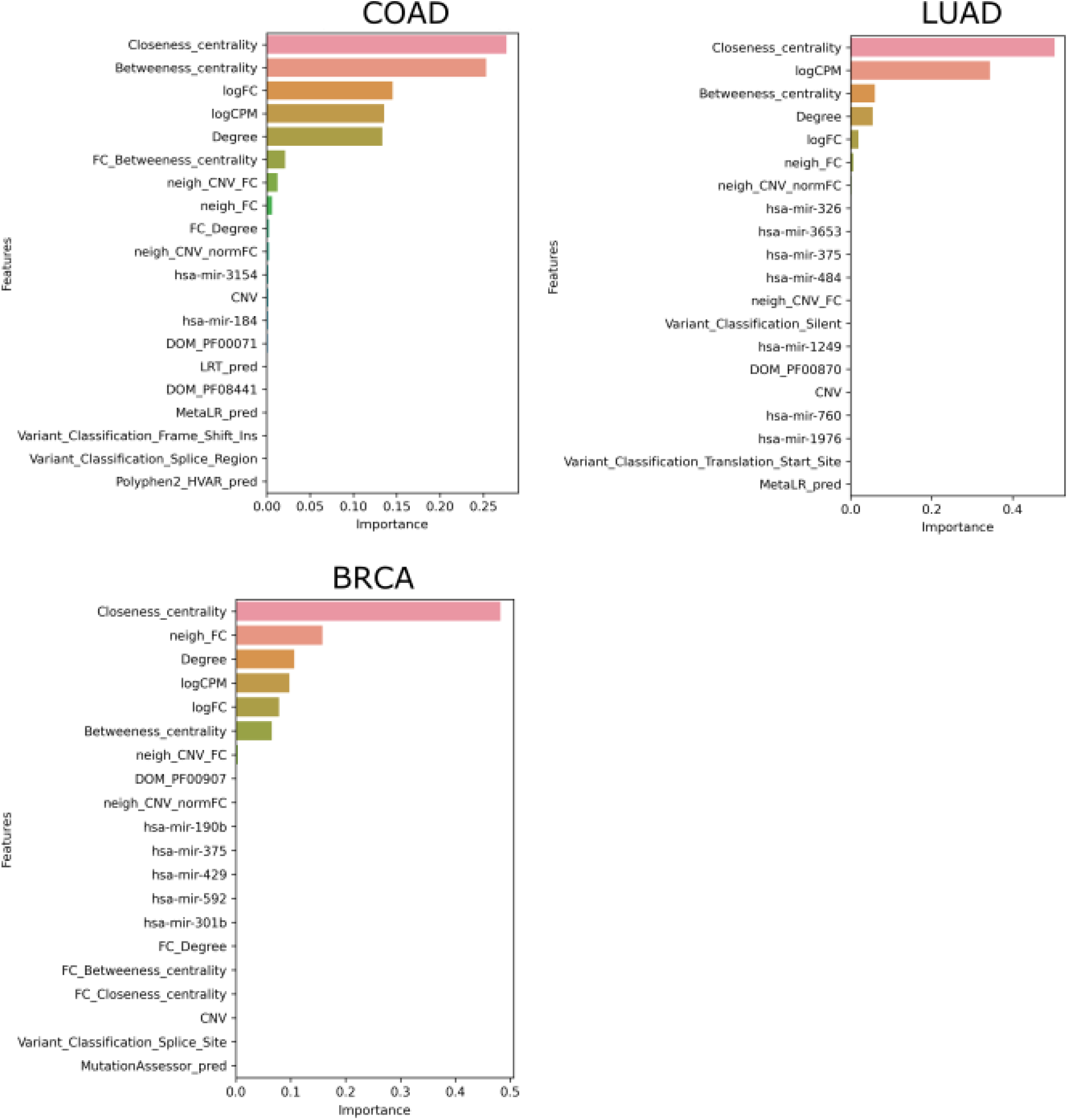
Multi-omic features contributing to the best model for cancer-type. For each cancer type, the contribution of the top 20 features is plotted.

### 2.7 Rare driver genes predicted across cancer-types

We used the multi-omic model to predict TSGs and OGs for BRCA, COAD and LUAD cancer types. Most samples predicted at least one TSG or OG. Out of 984 samples, we generated features for 972 samples and identified driver genes in 963. Similarly, only 3 samples in COAD predicted no driver genes and the number was larger at 62 for LUAD. Surprisingly, the number of unique genes predicted in each cancer type was large. In BRCA a total of 1342 unique genes were identified, followed by 1155 and 1152 in LUAD and COAD respectively. The distribution of genes identified across samples was similar in all cancer-types with most samples with < 100 genes identified as drivers (Supplementary Figures S43, S49 and S56). A large number of genes identified, consist of mutation as well as CNV alterations with CNVs contributing with as high as 100 genes altered in one sample (Supplementary Figures S44, S50 and S57). In contrast, most samples had an average of 10 mutations identified as driver genes (Supplementary Figures S45, S51 and S58) as previously reported in the literature (Vogelstein et al., 2000).

Genes identified across a large number of samples are well-known driver genes. The top 10 genes in each cancer type are listed in Table 2. Comparison with the list of genes in CGC showed an overlap of 228, 169, and 188 genes for BRCA, COAD and LUAD respectively. The list of top genes also consists of genes, such as *PRKCA* (Lin et al., 2017; Pham et al., 2017; Beetch et al., 2021; Kelemen et al., 2009), *SOX9* (Lizarraga et al., 2019; Carrasco-Garcia et al., 2016; Lü et al., 2008) and *NFKBIA* (Furukawa et al., 2013), that are not present in CGC but found in the literature for their role in respective cancer types. We also identify a large number of rare driver genes identified in as few as one sample in BRCA (Supplementary Table 6), COAD (Supplementary Table 7) and LUAD (Supplementary Table 8). In BRCA (Figure 5) and LUAD (Supplementary Figure S62), we observe a bimodal distribution with a large number of genes identified in a large number of samples along with genes that are mutated in less than 10 samples. Genes predicted in COAD cancer type on the other hand show consensus in a few samples (Supplementary Figure S55). The distribution is similar for the known driver gene listed in CGC and predicted by the model, indicating genes identified in a few samples are not false positives. We find genes, such as *TK1* in LUAD (Malvi et al., 2019; Xu et al., 2012) and *ELAVL1* in BRCA (Liu et al., 2019; Chou et al., 2015), predicted in only one sample but known to have a role in respective cancer types. The list of rare predicted genes provides potential genes to target and study to understand their role in the progression of tumours.

**Table 2.** Top genes identified for all cancer-types. The genes are listed along with the number of samples they were labelled TSGs or OGs. Their presence or absence in CGC is also given.

| BRCA |  |  | COAD |  |  | LUAD |  |  |
| --- | --- | --- | --- | --- | --- | --- | --- | --- |
| Gene | Samples | CGC | Gene | Samples | CGC | Gene | Samples | CGC |
| PIK3CA | 453 | Yes | APC | 408 | Yes | TP53 | 268 | Yes |
| TP53 | 383 | Yes | TP53 | 212 | Yes | KRAS | 186 | Yes |
| PRKCA | 239 | No | KRAS | 153 | Yes | STK11 | 135 | Yes |
| MYC | 235 | Yes | PIK3CA | 122 | Yes | CDKN2A | 133 | Yes |
| GATA3 | 230 | Yes | ARID1A | 87 | Yes | EGFR | 112 | Yes |
| PSMC5 | 229 | No | FBXW7 | 86 | Yes | KEAP1 | 111 | Yes |
| IKBKB | 228 | Yes | SOX9 | 81 | No | NF1 | 105 | Yes |
| PSMD12 | 228 | No | PRKDC | 73 | No | PSMD4 | 105 | No |
| MAP2K6 | 226 | No | SMAD4 | 72 | Yes | NFKBIA | 103 | No |
| RPS6KB1 | 225 | No | MTOR | 60 | Yes | MYC | 99 | Yes |

**Figure 5.**
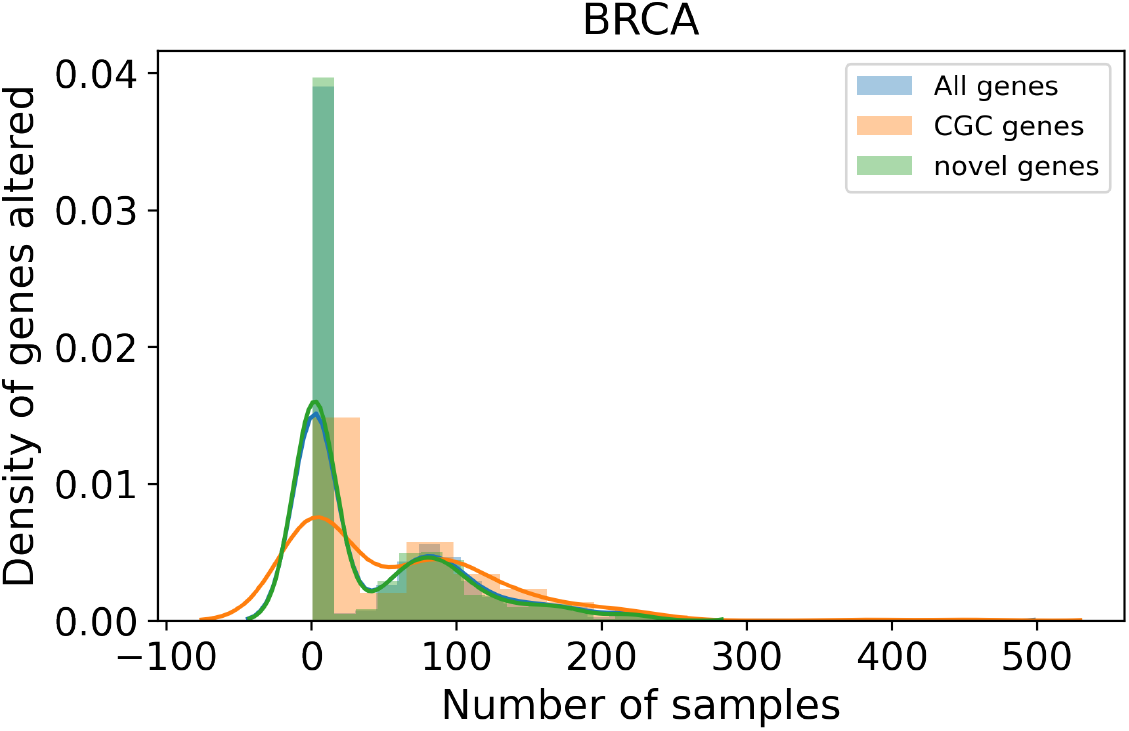
Distribution of samples for genes predicted as a driver. For BRCA, the distribution of samples a gene is identified as a driver.

### 2.8 Gene with dual roles in different samples identified

We observed that the number of TSGs in a given cancer type is always greater than OGs. In LUAD, 1030 TSGs and 195 OGs were predicted, while the number was 991 and 798 in BRCA and 938 and 266 in COAD. It is interesting to note that some genes are labelled differently in different samples, with as many as 447 genes in BRCA, 70 in LUAD and 52 in COAD. Genes labelled as both TSG and OG suggest genes behaving differently in not only cancer types but within sub-types of cancer. *JAK1* was one such gene identified as both TSG and OG in BRCA and COAD cancer types. *JAK1* is a signal transducer and activates the JAK/STAT pathway. It has been shown to be consistently active leading to cell survival in colon cancer (An et al., 2014) and lower survival rates (Tang et al., 2018). Similarly, *JAK1* is activated by PRLR signalling in a subset of breast cancers (Neilson et al., 2007), while underexpression of *JAK1* is needed for the invasion of immune response (Albacker et al., 2017; Chen et al., 2019). The role of *JAK1* is highly dependent on the cell conditions and can vary across different cancer sub-types (Yeh et al., 2007). *GNA11* is another gene that was classified as both a TSG and an OG in all three cancer types. The role of *GNA11* as an oncogene and the occurrence of mutations in tumour samples is well studied. But, unlike an oncogene downregulation of *GNA1* was observed in human breast cancers. Genes identified with multiple labels might be used to understand diverging roles of a gene in cancer.

## 3 DISCUSSION

Identifying driver genes is still a challenging problem with new tools being developed to identify driver genes. Tumours are highly heterogeneous, even within the same cancer type. While some genes such as *TP53* (Vogelstein et al., 2000; Petitjean et al., 2007) are mutated in a significant fraction of tumours, a small fraction of patients with no mutations in known cancer driver genes exist Campbell and et al. (2020). The treatment strategy is geared towards personalised medicine based on the presence of prognostic markers to optimise for patient’s recovery (Verma, 2012). Targeted therapy is given based on the genomic alterations in known driver genes (Cho et al., 2012; Ross et al., 2009; Amado et al., 2008). Identification of personalised driver genes is the first step to push the field of personalised medicine further. Personalised genes will not only help identify potential targets but identify genes mutated in only a small subset of samples.

We developed a machine learning model that employs multi-omic data to identify personalised driver genes. To the best of our knowledge, this study is the first supervised ML approach for identifying personalised driver genes. Our models label the genes based on their functionality as TSGs and OGs, another first in the field of personalised driver genes. We employ SNV, RNA, CNV, miRNA, network, and Pfam (Mistry et al., 2021) domains to build the models. Not all data from multiple omics will be available at all times. Hence, we build SNV and RNA models when only SNV or RNA data is available. Many driver genes are also hub genes, and integrating STRING network (Szklarczyk et al., 2019) information greatly improves the prediction as observed by our RNA and multi-omic models. All three node-based network features rank consistently high. While driver genes are known to have a high degree, the distribution of the degree of predicted genes shows genes with a low degree are also predicted and the model is not biased towards genes with a high degree.

Supervised classification models require labels to train the models. The field of personalised driver gene prediction is largely unexplored, and no gold standards are available to validate the results. Formulation of the supervised problem requires a label. The mutation-based labelling methods consist of highly curated mutations and genes, which build models with high accuracy for only one cancer type (BRCA). While models built on labelled mutations score high on classification metrics, their ability to predict genes not observed may vary. Further, ML algorithms predict consistently only when trained on large datasets that mimic all driver genes. We increased the training data by employing gene-based labelling methods and dropping SNV features with large missing data. Our final multi-omic models built on fewer features and many cancer-type specific driver genes perform the best.

The lack of gold standard further makes comparison difficult among different methods. Most driver gene lists are either pan-cancer or cancer-type specific. Comparison for personalised driver genes requires a list of diver genes specific for a patient. In the absence of any ground truth most methods use CGC gene list to compare predictions. While building models we show the performance is poorest on CGC genes as the list of gene is not specific to cancer-type. There is a need for establishing gold-standard for personalised driver genes to further the field.

Compared to previous methods that employ only mutation and expression data, we include known biological knowledge regarding domains and onco-miRNA expression. Further, the previous network-based unsupervised methods consider the presence or absence of mutations in the gene. The functional impact of the mutation is lost in the compression. We use mutation type and prediction scores by multiple tools, and we observe functional impact based features ranking high in the absence of network or RNA data. Further, the methods use expression data to identify DEGs to map onto the network. The method to identify DEGs is based on cohort, making it difficult to use in a clinical setting. We identify DEGs for an individual sample based on pre-computed values of biological variation for the cancer type. Any new sample can be processed to features and predict TSG and OG.

Our method has its limitation commonly observed with ML-based models. ML algorithms assume the training data covers all possible driver genes and the features capture all the required information to predict the genes. Given the lack of experimental data on personalised driver genes, we assume known driver mutations and all mutations in the curated driver gene list are drivers in the observed tumour. We use network data from STRING (Szklarczyk et al., 2019), and other networks such as Reactome can be used instead of or in combination. Currently, we use miRNA expression of known cancer miRNAs. We can generate gene-specific features to capture miRNA and mRNA interaction. Further, we can include scores from other personalised driver prediction tools as features to develop an ensemble model to improve predictions. The research space for the identification of personalised driver genes is mostly under-explored.

## 4 CONCLUSION

In this study, we make three significant contributions. First, we define the identification of personalised drivers as a supervised problem by defining and studying various labelling strategies. We conclude the best labelling strategy is using gene-based labelling that is cancer-specific. Second, we build multiple models on mutation, RNA and multi-omic features to identify the contribution of individual omic based features. We show network-based and expression features contribute the most to models, with the multi-omic models performing the best. In case of missing omic data, mutation or RNA models can be used for predicting driver genes. Lastly, our models can label genes as TSG and OG in a tumour. The functional labelling of genes is helpful for the identification of potential treatment strategies. Our method, PIVOT, is capable of predicting TSG and OG for individual samples, with multiple models available for predicting based on the availability of data.

## 5 METHODS

The models for identification of personalised TSGs and OGs are built using different modalities of data, feature sets. The data is labelled using four different strategies. The data is split into training and test and models are tuned using cross-validation on the training data. The best models are selected using metrics on the test data. The overview of model building is shown in Figure 6.

**Figure 6.**
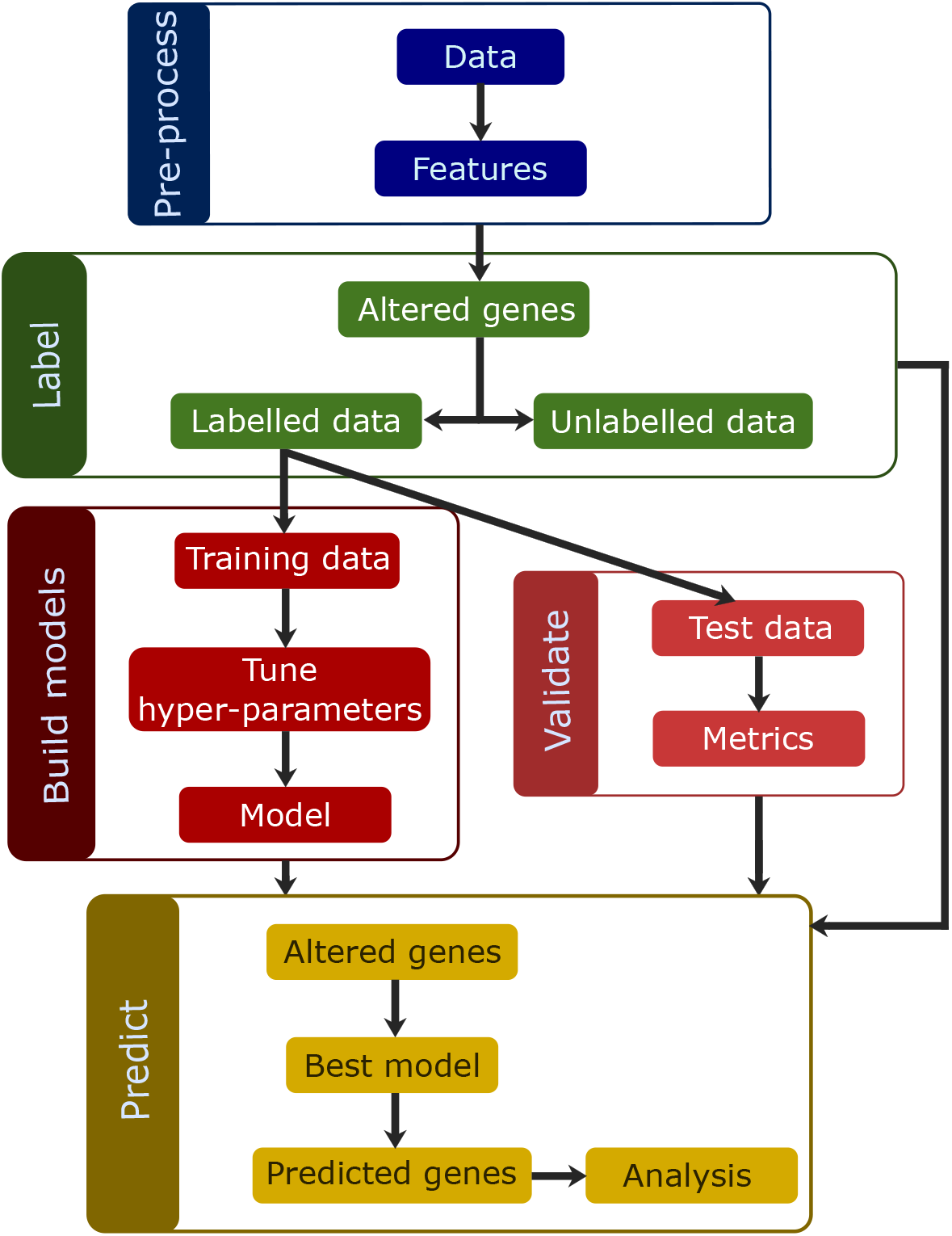
Overview of the methods. The data is pre-processed to generate features to be used for classification. The data is then labelled using one of the four labelling strategies, and samples are split into train and test. The training data is used to tune and build models. The metrics are calculated for the predictions made on test data and used to select the best model. The best model is then used to predict TSGs and OGs on the altered genes for all samples.

### 5.1 Data

The TCGA data was downloaded from GDC for the four cancer types, BRCA, COAD, LGG and LUAD. For each cancer type the mutation, expression, CNV and miRNA data was downloaded. The mutation file was downloaded as a maf file generated using Mutect2. The data was annotated using ANNOVAR and processed to include domain-based features. The expression data from RNA-sequencing experiments alone were downloaded for BRCA, COAD and LUAD as raw HT-seq counts. The data was processed using edgeR to obtain deferentially expressed genes (DEGs) for each patient. For BRCA, COAD and LUAD we downloaded CNV data as GISTIC gene-level copy number score. For labelling genes, driver genes lists are downloaded from the CIViC database, Cancer Gene Census (CGC), Martelotto et al. and Bailey et al. The list of neutral genes are also published by Bailey et al. included in GitHub PIVOT folder under folder data, subfolder driver. A list of onco-domains, miRNA associated with cancer-type is also included in GitHub PIVOT folder data. STRING database was downloaded for the protein-protein interaction (PPI) network. Only edges with experimental or database scores above 700 were retained. The graph was processed to generate degree, closeness centrality, betweenness centrality and all neighbours for a gene. The data is stored as pickle files and accessible via the GitHub PIVOT repository.

### 5.2 Feature generation

#### 5.2.1 Mutational features

For SNV features, the mutation data were annotated using ANNOVAR (Wang et al., 2010) to include prediction scores for various tools. The mutation type was one-hot encoded. All other categorical predicted features were converted into ordinal categories, where missing data was given value 0. The list of domains (Hashemi et al., 2017) was processed to retrieve Pfam ids. All onco-domains for the given cancer type were used as features, and the domain feature was assigned value one if the mutation is associated with the domain. The mutation and list of domains associated are given in the mutation maf file. The number of features vary in each cancer type and are dependent on the number of cancer domain identified.

#### 5.2.2 Expression features

For RNA features, the data was processed for each sample to generate a list of DEGs using edgeR. Differential genes are usually reported for a cohort. It is advisable to have more than one sample in each condition. In a clinical setting, multiple samples may not be available. We generate DEGs for each sample. Not all samples contain adjacent normal for a cancer type. We group all normal samples and run DEG against all tumour samples individually. The biological variance (bcv) is specified when only one sample is present. We calculate the common bcv for all tumour samples against normal. The common bcv is further used for the identification of DEGs for individual samples and saved as individual files. The logFC and logCPM values are used as features. Other features include node properties degree, closeness centrality, and betweenness centrality. Another three features are produced, multiplying the logFC to the node properties. The differential expression of neighbours is captured by features *neigh_FC* and *neigh_normFC*. For a given gene, *neigh_FC* is calculated as the sum of logFC of all neighbours of the gene, with fold change > 2 or < −2. *Neigh_normFC* is normalised for the number of genes differentially expressed in the neighbourhood.

#### 5.2.3 Multi-omic features

Multi-omic features concatenate the SNV, RNA features, CNV and miRNA data. The effect of CNV on neighbourhood is calculated by *neigh_CNV_FC* and *neigh_CNV_normFC* similar to RNA features. All genes with copy number variation ≠ 0, the sum of logFC values > 2 or < − 2 of all neighbouring genes defines *neigh_CNV_FC*. The neighbourhood is defined by all genes that can be accessed with ≤ *n* hops. We consider *n* = 1 for this analysis. The list of miRNA associated with cancer type was downloaded from OncomiR Cancer Database (OCMD; Sarver et al. (2018)) and OncomiR (Wong et al., 2018) databases and the intersection of both databases was used for the analysis. Expression of the known oncogenic miRNAs was used as features for predicting.

### 5.3 Classification

The data is labelled using four different gene lists. CIViC and Martelotto et al. list the gene as well as the mutation location. Only mutations with the exact location and base change are used labelled. CIViC labels genes as *driver*, while all other methods label as genes as TSG or OG. CGC and Bailey et al. label all mutations in a gene with the same label. Further, we labelled genes with copy number variations in driver genes based on labels assigned by gene-based labelling strategies for multi-omic analysis. Classification of CIViC labels was a binary classification problem, while the rest were multi-class classification problems. We used 70:30 split the samples into train and test data. All the mutations and/or CNV alterations were labelled using one of the four lists of driver genes described earlier. It is to be noted the number of data points in train and test split vary for the different labelling strategies (Supplementary Tables 1,4 and 5). The data is highly imbalanced, with a large number of neutral genes and very few TSG and OG. We used sampling algorithms for imbalanced data from *imblearn* package. Models were built using balanced random forest, Balanced bagging and easy ensemble. Five-fold cross-validation was conducted to tune hyperparameters. The precision-recall (PR) curve and the receiver operator curve (ROC) was plotted for all classes and all models. Further, the top 20 features contributing to the model are plotted. The accuracy, F1 score, precision and recall for the training and test set was calculated. All the models, plots and output metrics are available in the GitHub folder and as supplementary data.

### 5.4 Feature, domain and miRNA analysis

For each cancer type, the consensus feature contribution was calculated as the average feature contribution for all models built on the dataset. The top domains for SNV and multi-omic datasets were identified based on all domains with feature contribution > 0. Similarly, the top miRNA features with consensus feature contribution > 0 for the cancer type were listed.

## Supporting information

Summary statistics for SNV models.

Feature contribution for all models and their consensus.

List of domains and their contributions.

Summary statistics for RNA models.

Summary statistics for multi-omic models.

List of predictions for BRCA cancer type.

List of predictions for COAD cancer type.

List of predictions for LUAD cancer type.

List of miRNA features and their contributions.

List of all supplementary figures.

## AUTHOR CONTRIBUTIONS

MS, RR, and KR conceived and designed the study. MS built the ML models. MS, RR, and KR were involved in analysing and interpreting data. MS, RR and KR drafted the manuscript. RR and KR supervised the study. All authors read and approved the final manuscript.

## ACKNOWLEDGMENTS

The results published here are in whole or part based upon data generated by the TCGA Research Network: https://www.cancer.gov/tcga.

## SUPPLEMENTAL DATA

**Supplementary Table 1. Summary statistics for SNV models**. Details on the number of samples and data points trained and tested on for each model. The accuracy, F1-score, precision and recall for training and test set for all labelling strategies and cancer types are calculated.

**Supplementary Table 2: Feature contribution for all models and their consensus**. For each model built on a cancer type, the feature contribution was calculated. The average values across the models is listed under consensus.

**Supplementary Table 3: List of domains and their contributions**. The Pfam domains, domain features and their average contribution to SNV and multi-omic models are listed.

**Supplementary Table 4: Summary statistics for RNA models**. Lists the number of samples and data points trained and tested on for each model. The accuracy, F1-score, precision and recall for training and test set for all labelling strategies and cancer types are calculated.

**Supplementary Table 5: Summary statistics for multi-omic models**. Lists the number of samples and data points trained and tested on for each model. The accuracy, F1-score, precision and recall for training and test set for all labelling strategies and cancer types are calculated.

**Supplementary Table 6. List of predictions for BRCA cancer type**.

**Supplementary Table 7. List of predictions for COAD cancer type**.

**Supplementary Table 8. List of predictions for LUAD cancer type**.

**Supplementary Table 9: List of miRNA features and their contributions**. The miRNA and their average contribution to multi-omic models are listed for individual cancer types.

**Supplementary Figures: List of all supplementary figures**.

## DATA AVAILABILITY STATEMENT

The TCGA datasets were downloaded using Genomic Data Commons (GDC). The processed data analysed for this study can be found in the GitHub repository PIVOT under folder data (https://github.com/RamanLab/PIVOT).

