## Supplementary material for "Multi-omic data helps improve prediction of personalised tumor suppressors and oncogenes": List of all supplementary figures.

### List Of Supplementary Figures


- Figure S1A. Top 20 features for BRCA using SNV data with Bailey et al. labels built on “all” features using Balanced bagging.
- Figure S1B. Precision-recall curve for BRCA using SNV data with Bailey et al. labels built on “all” features using Balanced bagging.
- Figure S1C Receiver operating characteristic for BRCA using SNV data with Bailey et al. labels built on “all” features using Balanced bagging.
- Figure S2A. Top 20 features for BRCA using SNV data with Bailey et al. labels built on “some” features using Balanced bagging.
- Figure S2B. Precision-recall curve for BRCA using SNV data with Bailey et al. labels built on “some” features using Balanced bagging.
- Figure S2C Receiver operating characteristic for BRCA using SNV data with Bailey et al. labels built on “some” features using Balanced bagging.
- Figure S3A. Top 20 features for BRCA using SNV data with CGC labels built on “all” features using Balanced bagging.
- Figure S3B. Precision-recall curve for BRCA using SNV data with CGC labels built on “all” features using Balanced bagging.
- Figure S3C.  Receiver operating characteristic for BRCA using SNV data with CGC labels built on “all” features using Balanced bagging.
- Figure S4A. Top 20 features for BRCA using SNV data with CGC labels built on “some” features using Balanced Random Forest.
- Figure S4B. Precision-recall curve for BRCA using SNV data with CGC labels built on “some” features using Balanced Random Forest.
- Figure S4C.  Receiver operating characteristic for BRCA using SNV data with CGC labels built on “some” features using Balanced Random Forest.
- Figure S5A. Top 20 features for BRCA using SNV data with CIViC labels built on “all” features using Easy Ensemble.
- Figure S5B. Precision-recall curve for BRCA using SNV data with CIViC labels built on “all” features using Easy Ensemble.
- Figure S5C.  Receiver operating characteristic for BRCA using SNV data with CIViC labels built on “all” features using Easy Ensemble.
- Figure S6A. Top 20 features for BRCA using SNV data with CIViC labels built on “some” features using Balanced Bagging.
- Figure S6B. Precision-recall curve for BRCA using SNV data with CIViC labels built on “some” features using Balanced Bagging.
- Figure S6C.  Receiver operating characteristic for BRCA using SNV data with CIViC labels built on “some” features using Balanced Bagging.
- Figure S7A. Top 20 features for BRCA using SNV data with Marellotto et al. labels built on “all” features using Easy Ensemble.

- Figure S7B. Precision-recall curve for BRCA using SNV data with Marellotto et al. labels built on “all” features using Easy Ensemble.
- Figure S7C.  Receiver operating characteristic for BRCA using SNV data with Marellotto et al. labels built on “all” features using Easy Ensemble.
- Figure S8A. Top 20 features for BRCA using SNV data with Marellotto et al. labels built on “some” features using Balanced Bagging.
- Figure S8B. Precision-recall curve for BRCA using SNV data with Marellotto et al. labels built on “some” features using Balanced Bagging.
- Figure S8C.  Receiver operating characteristic for BRCA using SNV data with Marellotto et al. labels built on “some” features using Balanced Bagging.
- Figure S9A. Top 20 features for COAD using SNV data with Bailey et al. labels built on “all” features using Balanced Random Forest.
- Figure S9B. Precision-recall curve for COAD using SNV data with  Bailey et al. labels built on “all” features using Balanced Random Forest.
- Figure S9C Receiver operating characteristic for COAD using SNV data with Bailey et al. labels built on “all” features using Balanced Random Forest.
- Figure S10A. Top 20 features for COAD using SNV data with Bailey et al. labels built on “some” features using Easy Ensemble.
- Figure S10B. Precision-recall curve for COAD using SNV data with  Bailey et al. labels built on “some” features using Easy Ensemble.
- Figure S10C Receiver operating characteristic for COAD using SNV data with Bailey et al. labels built on “some” features using Easy Ensemble.
- Figure S11A. Top 20 features for COAD using SNV data with CGC labels built on “all” features using Balanced Bagging.
- Figure S11B. Precision-recall curve for COAD using SNV data with CGC labels built on “all” features using Balanced Bagging.
- Figure S11C Receiver operating characteristic for COAD using SNV data with CGC labels built on “all” features using Balanced Bagging.
- Figure S12A. Top 20 features for COAD using SNV data with CGC labels built on “some” features using Balanced Random Forest.
- Figure S12B. Precision-recall curve for COAD using SNV data with CGC labels built on “some” features using Balanced Random Forest.
- Figure S12C Receiver operating characteristic for COAD using SNV data with CGC labels built on “some” features using Balanced Random Forest.
- Figure S13A. Top 20 features for LGG using SNV data with Bailey et al. labels built on “all” features using Balanced Random Forest.
- Figure S13B. Precision-recall curve for LGG using SNV data with Bailey et al. labels built on “all” features using Balanced Random Forest.
- Figure S13C Receiver operating characteristic for LGG using SNV data with Bailey et al. labels built on “all” features using Balanced Random Forest.
- Figure S14A. Top 20 features for LGG using SNV data with Bailey et al. labels built on “some” features using Easy Ensemble.
- Figure S14B. Precision-recall curve for LGG using SNV data with  Bailey et al. labels

built on “some” features using Easy Ensemble.
- Figure S14C Receiver operating characteristic for LGG using SNV data with Bailey et al. labels built on “some” features using Easy Ensemble.
- Figure S15A. Top 20 features for LGG using SNV data with CGC labels built on “all” features using Balanced Bagging.
- Figure S15B. Precision-recall curve for LGG using SNV data with CGC labels built on “all” features using Balanced Bagging.
- Figure S15C Receiver operating characteristic for LGG using SNV data with CGC labels built on “all” features using Balanced Bagging.
- Figure S16A. Top 20 features for LGG using SNV data with CGC labels built on “some” features using Balanced Random Forest.
- Figure S16B. Precision-recall curve for LGG using SNV data with CGC labels built on “some” features using Balanced Random Forest.
- Figure S16C Receiver operating characteristic for LGG using SNV data with CGC labels built on “some” features using Balanced Random Forest.
- Figure S17A. Top 20 features for LUAD using SNV data with Bailey et al. labels built on “all” features using Balanced Random Forest.
- Figure S17B. Precision-recall curve for LUAD using SNV data with Bailey et al. labels built on “all” features using Balanced Random Forest.
- Figure S17C Receiver operating characteristic for LUAD using SNV data with Bailey et al. labels built on “all” features using Balanced Random Forest.
- Figure S18A. Top 20 features for LUAD using SNV data with Bailey et al. labels built on “some” features using Balanced Random Forest.
- Figure S18B. Precision-recall curve for LUAD using SNV data with Bailey et al. labels built on “some” features using Balanced Random Forest.
- Figure S18C Receiver operating characteristic for LUAD using SNV data with Bailey et al. labels built on “some” features using Balanced Random Forest.
- Figure S19A. Top 20 features for BRCA using RNA data with Bailey et al. labels built using Balanced bagging.
- Figure S19B. Precision-recall curve for BRCA using RNA data with Bailey et al. labels built using Balanced bagging.
- Figure S19C Receiver operating characteristic for BRCA using RNA data with Bailey et al. labels built using Balanced bagging.
- Figure S20A. Top 20 features for BRCA using RNA data with CGC labels built using Balanced bagging.
- Figure S20B. Precision-recall curve for BRCA using RNA data with CGC labels built using Balanced bagging.
- Figure S20C Receiver operating characteristic for BRCA using RNA data with CGC labels built using Balanced bagging.
- Figure S21A. Top 20 features for BRCA using RNA data with CIViC labels built using Balanced bagging.
- Figure S21B. Precision-recall curve for BRCA using RNA data with CIViC labels built using Balanced bagging.

---

- Figure S21C Receiver operating characteristic for BRCA using RNA data with CIViC labels built using Balanced bagging.
- Figure S22A. Top 20 features for BRCA using RNA data with Marellotto et al. labels built using Easy Ensemble.
- Figure S22B. Precision-recall curve for BRCA using RNA data with Marellotto et al. labels built using Easy Ensemble.
- Figure S22C Receiver operating characteristic for BRCA using RNA data with Marellotto et al. labels built using Easy Ensemble.
- Figure S23A. Top 20 features for COAD using RNA data with Bailey et al. labels built using Balanced bagging.
- Figure S23B. Precision-recall curve for COAD using RNA data with Bailey et al. labels built using Balanced bagging.
- Figure S23C Receiver operating characteristic for COAD using RNA data with Bailey et al. labels built using Balanced bagging.
- Figure S24A. Top 20 features for COAD using RNA data with CGC labels built using Balanced bagging.
- Figure S24B. Precision-recall curve for COAD using RNA data with CGC labels built using Balanced bagging.
- Figure S24C Receiver operating characteristic for COAD using RNA data with CGC labels built using Balanced bagging.
- Figure S25A. Top 20 features for LUAD using RNA data with Bailey et al. labels built using Balanced bagging.
- Figure S25B. Precision-recall curve for LUAD using RNA data with Bailey et al. labels built using Balanced bagging.
- Figure S25C Receiver operating characteristic for LUAD using RNA data with Bailey et al. labels built using Balanced bagging.
- Figure S26A. Top 20 features for LUAD using RNA data with CGC labels built using Balanced bagging.
- Figure S26B. Precision-recall curve for LUAD using RNA data with CGC labels built using Balanced bagging.
- Figure S26C Receiver operating characteristic for LUAD using RNA data with CGC labels built using Balanced bagging.
- Figure S27A. Top 20 features for BRCA using multi-omic data with Bailey et al. labels built on “all” features using Balanced bagging.
- Figure S27B. Precision-recall curve for BRCA using multi-omic data with Bailey et al. labels built on “all” features using Balanced bagging.
- Figure S27C Receiver operating characteristic for BRCA using multi-omic data with Bailey et al. labels built on “all” features using Balanced bagging.
- Figure S28A. Top 20 features for BRCA using multi-omic data with Bailey et al. labels built on “some” features using Balanced bagging.
- Figure S28B. Precision-recall curve for BRCA using multi-omic data with Bailey et al. labels built on “some” features using Balanced bagging.
- Figure S28C Receiver operating characteristic for BRCA using multi-omic data with

Bailey et al. labels built on “some” features using Balanced bagging.

- Figure S29A. Top 20 features for BRCA using multi-omic data with CGC labels built on “all” features using Balanced bagging.
- Figure S29B. Precision-recall curve for BRCA using multi-omic data with CGC labels built on “all” features using Balanced bagging.
- Figure S29C.  Receiver operating characteristic for BRCA using multi-omic data with CGC labels built on “all” features using Balanced Bagging.
- Figure S30A. Top 20 features for BRCA using multi-omic data with CGC labels built on “some” features using Balanced Bagging.
- Figure S30B. Precision-recall curve for BRCA using multi-omic data with CGC labels built on “some” features using Balanced Bagging.
- Figure S30C.  Receiver operating characteristic for BRCA using multi-omic data with CGC labels built on “some” features using Balanced Bagging.
- Figure S31A. Top 20 features for BRCA using multi-omic data with CIViC labels built on “all” features using Balanced Bagging.
- Figure S31B. Precision-recall curve for BRCA using multi-omic data with CIViC labels built on “all” features using Balanced Bagging.
- Figure S31C.  Receiver operating characteristic for BRCA using multi-omic data with CIViC labels built on “all” features using Balanced Bagging.
- Figure S32A. Top 20 features for BRCA using multi-omic data with CIViC labels built on “some” features using Balanced Bagging.
- Figure S32B. Precision-recall curve for BRCA using multi-omic data with CIViC labels built on “some” features using Balanced Bagging.
- Figure S32C.  Receiver operating characteristic for BRCA using multi-omic data with CIViC labels built on “some” features using Balanced Bagging.
- Figure S33A. Top 20 features for BRCA using multi-omic data with Marellotto et al. labels built on “all” features using Balanced Bagging.
- Figure S33B. Precision-recall curve for BRCA using multi-omic data with Marellotto et al. labels built on “all” features using Balanced Bagging.
- Figure S33C.  Receiver operating characteristic for BRCA using multi-omic data with Marellotto et al. labels built on “all” features using Balanced Bagging.
- Figure S34A. Top 20 features for BRCA using multi-omic data with Marellotto et al. labels built on “some” features using Balanced Bagging.
- Figure S34B. Precision-recall curve for BRCA using multi-omic data with Marellotto et al. labels built on “some” features using Balanced Bagging.
- Figure S34C.  Receiver operating characteristic for BRCA using multi-omic data with Marellotto et al. labels built on “some” features using Balanced Bagging.
- Figure S35A. Top 20 features for COAD using multi-omic data with Bailey et al. labels built on “all” features using Balanced bagging.
- Figure S35B. Precision-recall curve for COAD using multi-omic data with Bailey et al. labels built on “all” features using Balanced bagging.
- Figure S35C Receiver operating characteristic for COAD using multi-omic data with Bailey et al. labels built on “all” features using Balanced bagging.

- Figure S36A. Top 20 features for COAD using multi-omic data with Bailey et al. labels built on “some” features using Balanced bagging.
- Figure S36B. Precision-recall curve for COAD using multi-omic data with Bailey et al. labels built on “some” features using Balanced bagging.
- Figure S36C Receiver operating characteristic for COAD using multi-omic data with Bailey et al. labels built on “some” features using Balanced bagging.
- Figure S37A. Top 20 features for COAD using multi-omic data with CGC labels built on “all” features using Balanced bagging.
- Figure S37B. Precision-recall curve for COAD using multi-omic data with CGC labels built on “all” features using Balanced bagging.
- Figure S37C.  Receiver operating characteristic for COAD using multi-omic data with CGC labels built on “all” features using Balanced Bagging.
- Figure S38A. Top 20 features for COAD using multi-omic data with CGC labels built on “some” features using Balanced Bagging.
- Figure S38B. Precision-recall curve for COAD using multi-omic data with CGC labels built on “some” features using Balanced Bagging.
- Figure S38C.  Receiver operating characteristic for COAD using multi-omic data with CGC labels built on “some” features using Balanced Bagging.
- Figure S39A. Top 20 features for LUAD using multi-omic data with Bailey et al. labels built on “all” features using Balanced bagging.
- Figure S39B. Precision-recall curve for LUAD using multi-omic data with Bailey et al. labels built on “all” features using Balanced bagging.
- Figure S39C Receiver operating characteristic for LUAD using multi-omic data with Bailey et al. labels built on “all” features using Balanced bagging.
- Figure S40A. Top 20 features for LUAD using multi-omic data with Bailey et al. labels built on “some” features using Balanced bagging.
- Figure S40B. Precision-recall curve for LUAD using multi-omic data with Bailey et al. labels built on “some” features using Balanced bagging.
- Figure S40C Receiver operating characteristic for LUAD using multi-omic data with Bailey et al. labels built on “some” features using Balanced bagging.
- Figure S41A. Top 20 features for LUAD using multi-omic data with CGC labels built on “all” features using Balanced bagging.
- Figure S41B. Precision-recall curve for LUAD using multi-omic data with CGC labels built on “all” features using Balanced bagging.
- Figure S41C.  Receiver operating characteristic for LUAD using multi-omic data with CGC labels built on “all” features using Balanced Bagging.
- Figure S42A. Top 20 features for LUAD using multi-omic data with CGC labels built on “some” features using Balanced Bagging.
- Figure S42B. Precision-recall curve for LUAD using multi-omic data with CGC labels built on “some” features using Balanced Bagging.
- Figure S42C.  Receiver operating characteristic for LUAD using multi-omic data with CGC labels built on “some” features using Balanced Bagging.
- Figure S43.  Distribution of genes predicted for samples in BRCA.

- Figure S44. Distribution of mutated genes predicted for samples in BRCA.
- Figure S45. Distribution of CNV altered genes predicted for samples in BRCA.
- Figure S46. Distribution of degree of genes predicted for samples in BRCA.
- Figure S47. Distribution of predicted oncogenes for samples in BRCA.
- Figure S48. Distribution of predicted tumour suppressor genes in BRCA.
- Figure S49. Distribution of genes predicted for samples in COAD.
- Figure S50. Distribution of mutated genes predicted for samples in COAD.
- Figure S51. Distribution of CNV altered genes predicted for samples in COAD.
- Figure S52. Distribution of degree of genes predicted for samples in COAD.
- Figure S53. Distribution of predicted oncogenes for samples in COAD.
- Figure S54. Distribution of predicted tumour suppressor genes in COAD.
- Figure S55. Distribution of samples for genes predicted as driver in COAD.
- Figure S56. Distribution of genes predicted for samples in LUAD.
- Figure S57. Distribution of mutated genes predicted for samples in LUAD.
- Figure S58. Distribution of CNV altered genes predicted for samples in LUAD.
- Figure S59. Distribution of degree of genes predicted for samples in LUAD.
- Figure S60. Distribution of predicted oncogenes for samples in LUAD.
- Figure S61. Distribution of predicted tumour suppressor genes in LUAD.
- Figure S62. Distribution of samples for genes predicted as driver in LUAD.


---

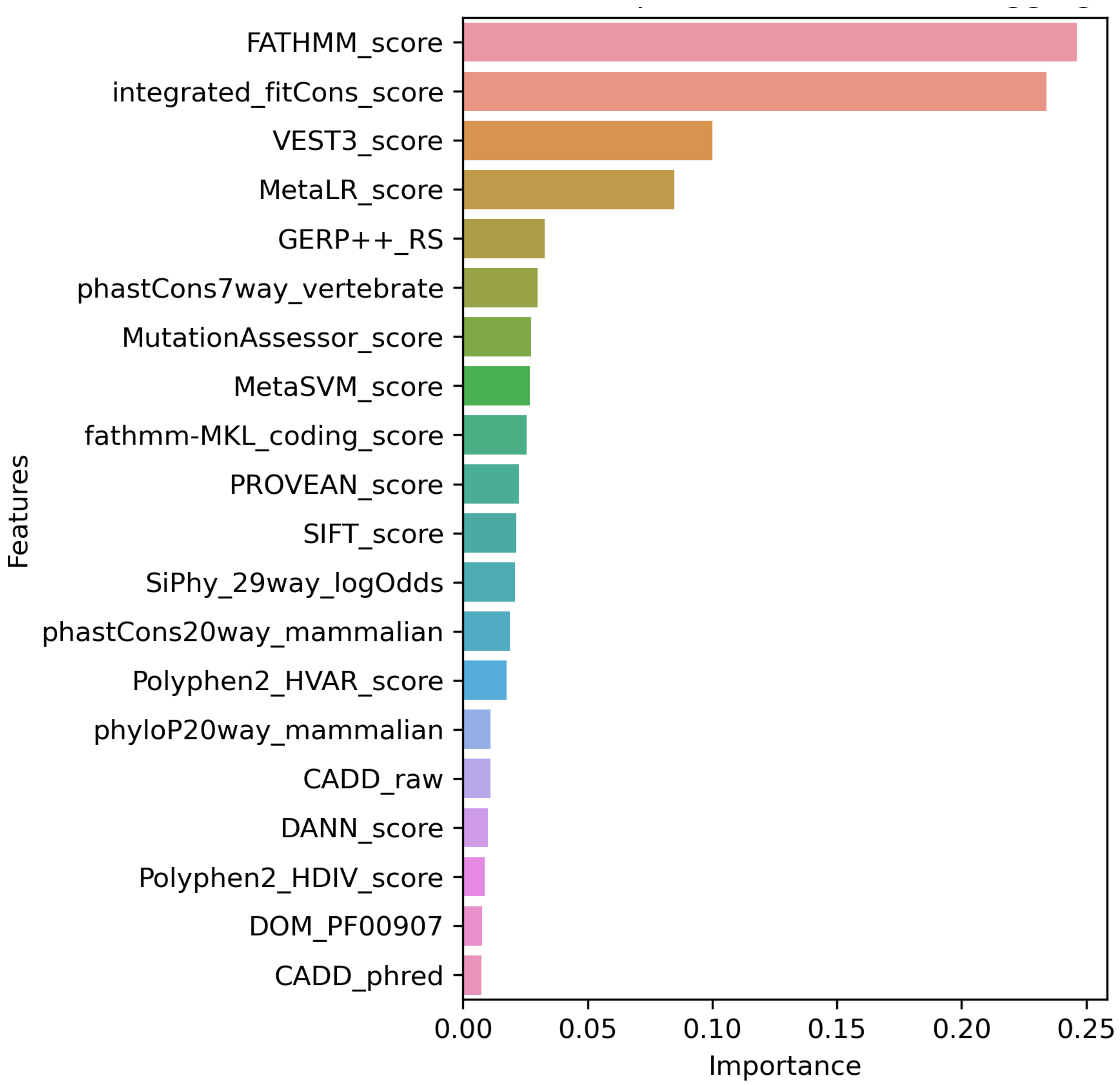


**Figure S1A. Top 20 features for BRCA using SNV data with Bailey *et al.* labels built on "all" features using Balanced bagging.**

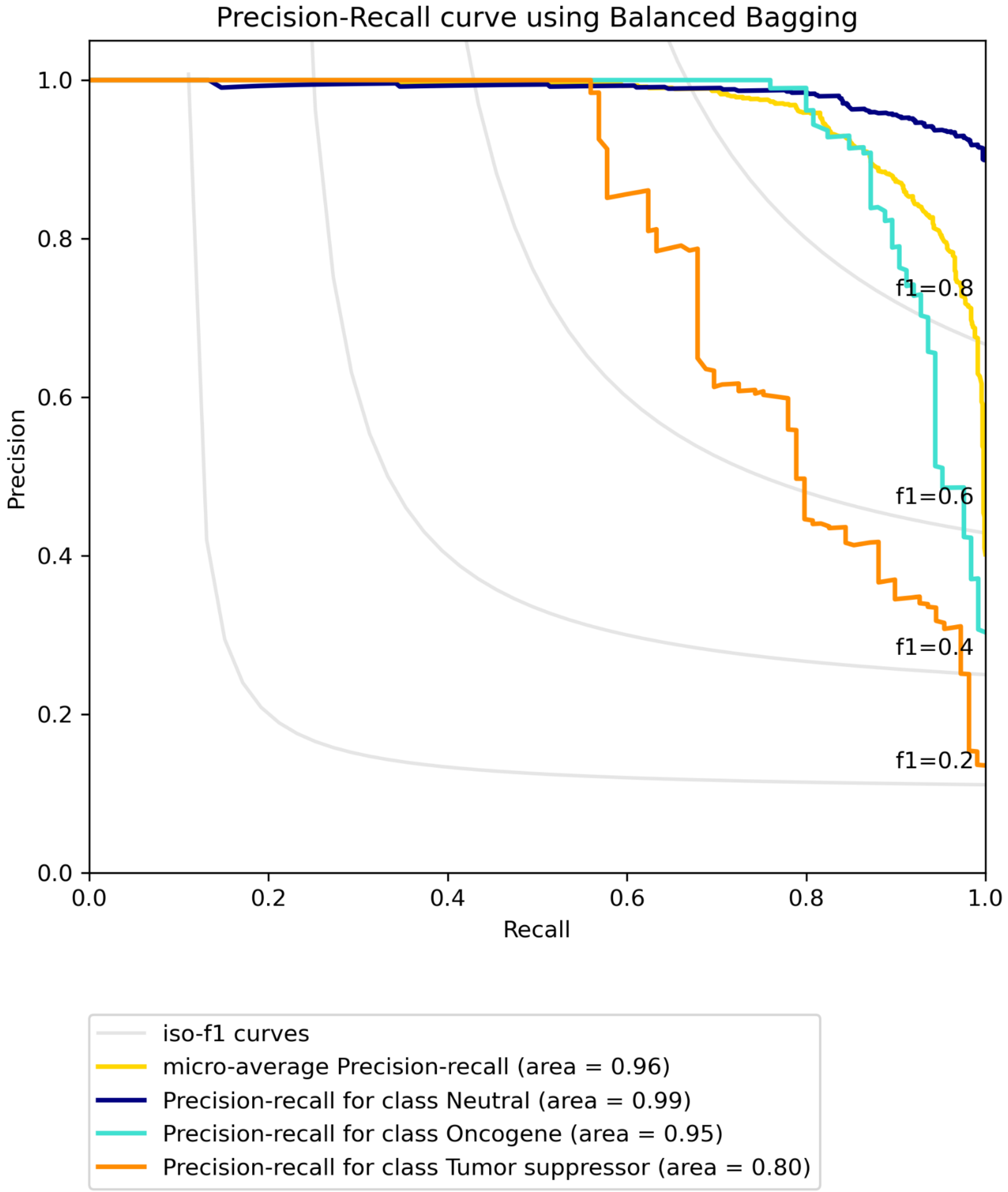


**Figure S1B. Precision-recall curve for BRCA using SNV data with Bailey *et al.* labels built on "all" features using Balanced bagging.**

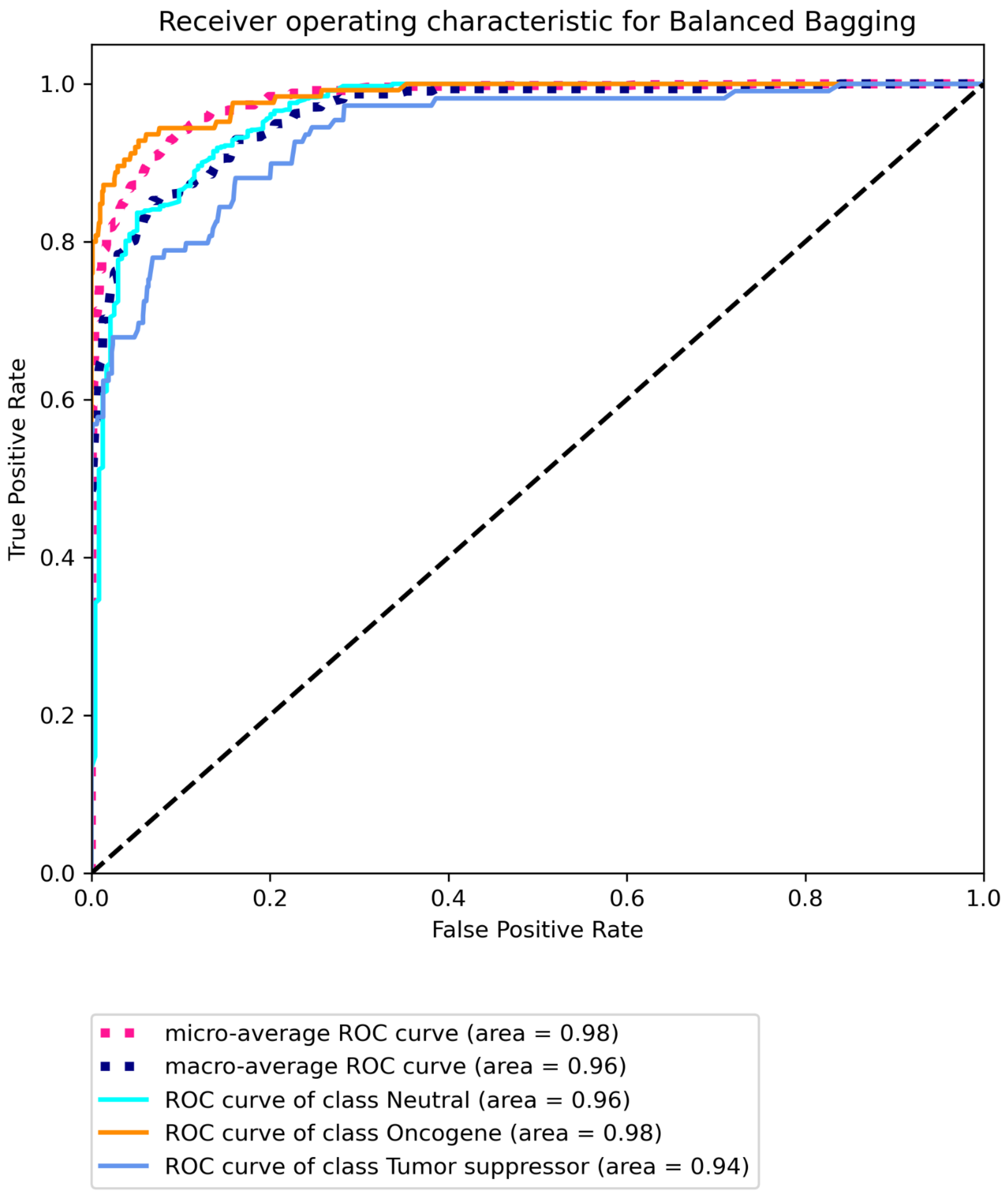


**Figure S1C Receiver operating characteristic for BRCA using SNV data with Bailey *et al.* labels built on "all" features using Balanced bagging.**

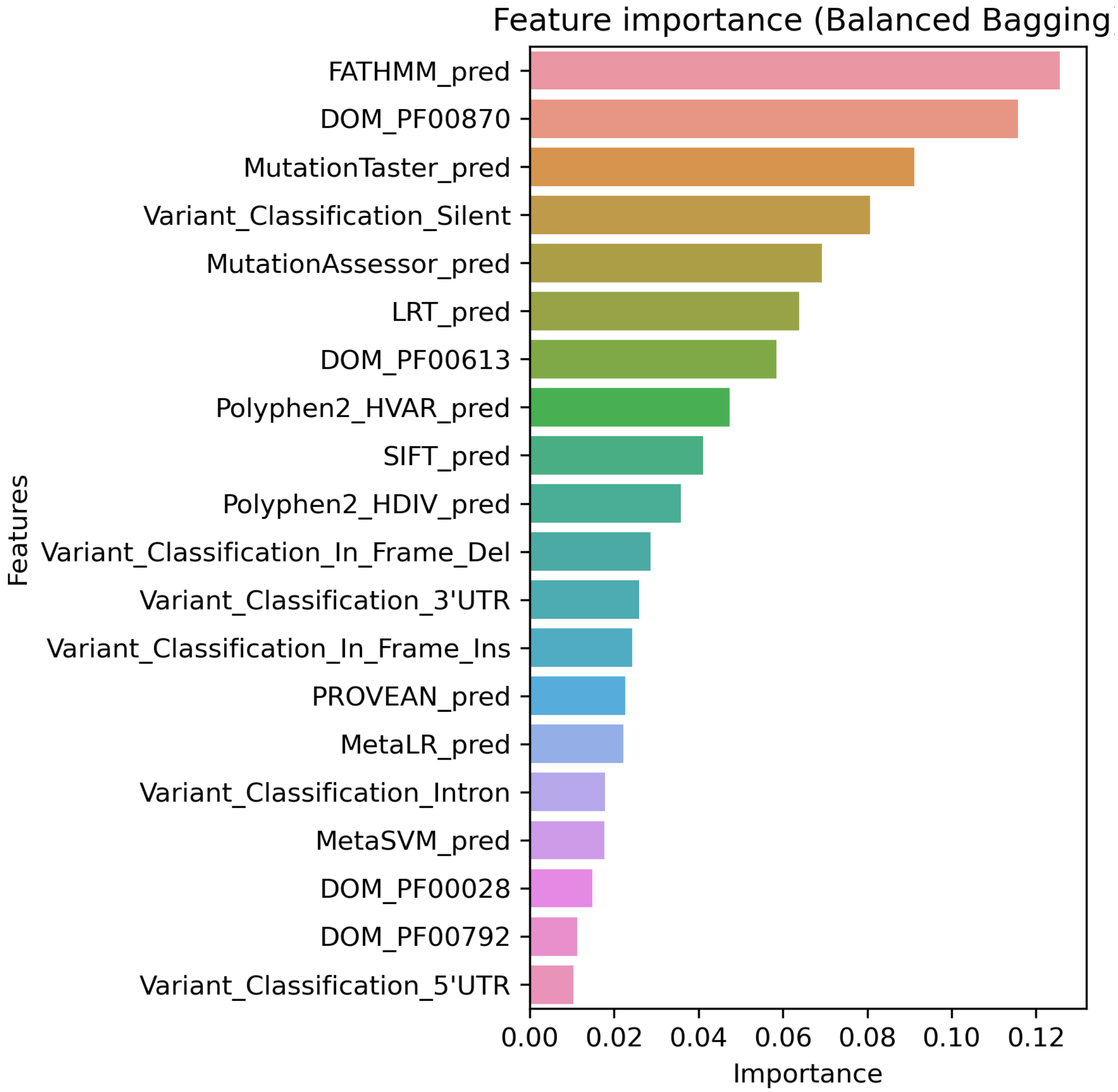


**Figure S2A. Top 20 features for BRCA using SNV data with Bailey *et al.* labels built on "some" features using Balanced bagging.**

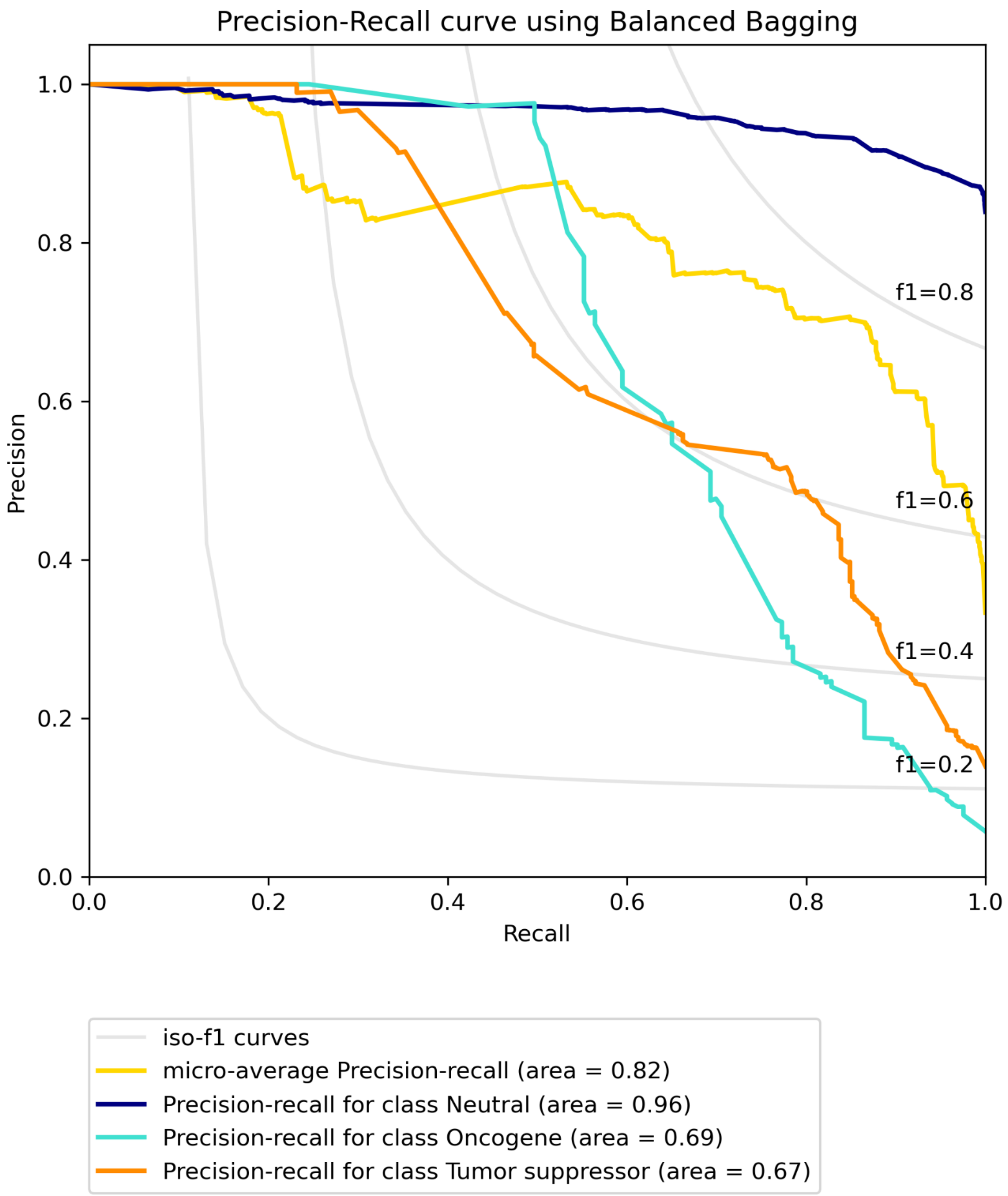


**Figure S2B. Precision-recall curve for BRCA using SNV data with Bailey *et al.* labels built on "some" features using Balanced bagging.**

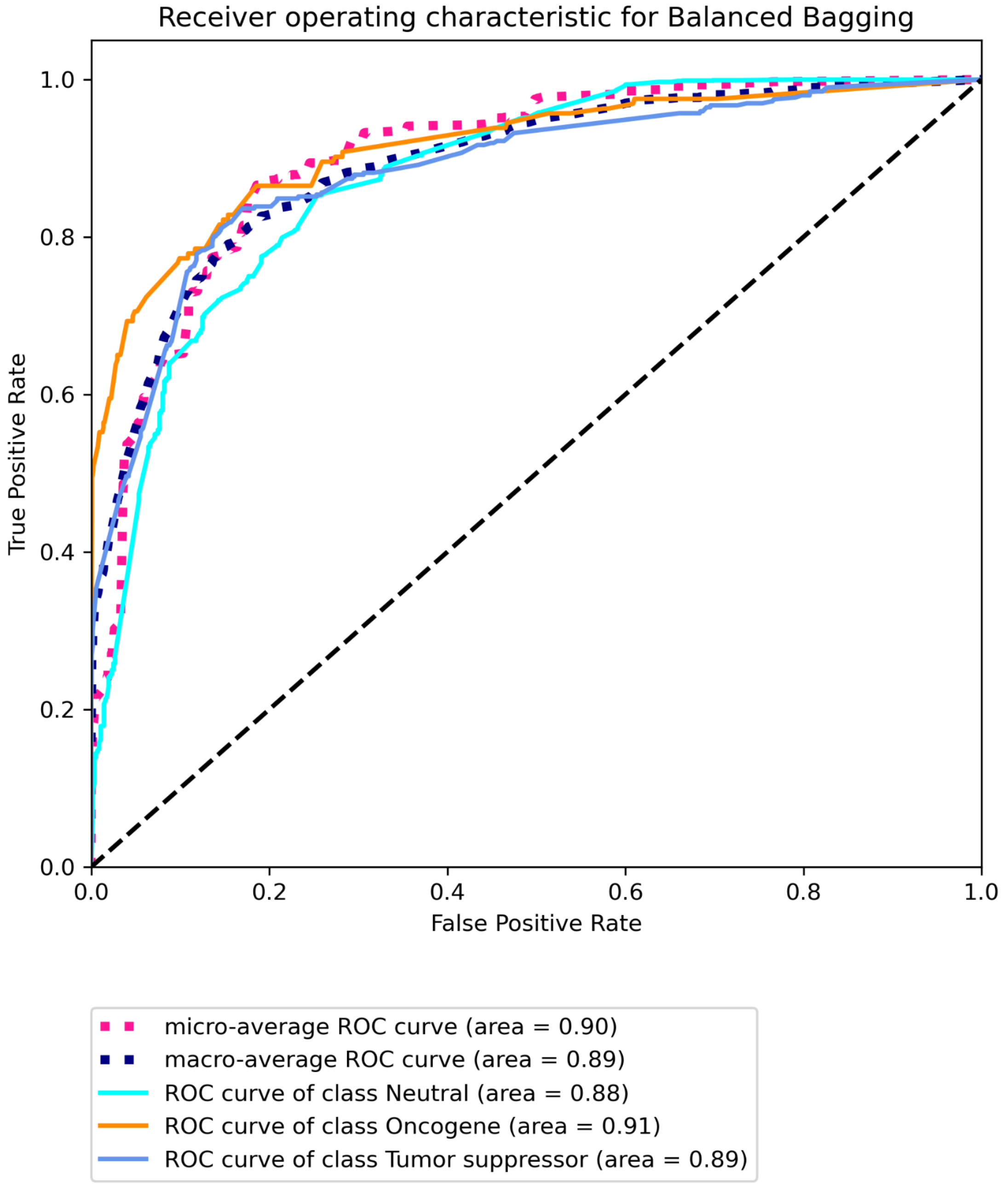


**Figure S2C Receiver operating characteristic for BRCA using SNV data with Bailey *et al.* labels built on "some" features using Balanced bagging.**

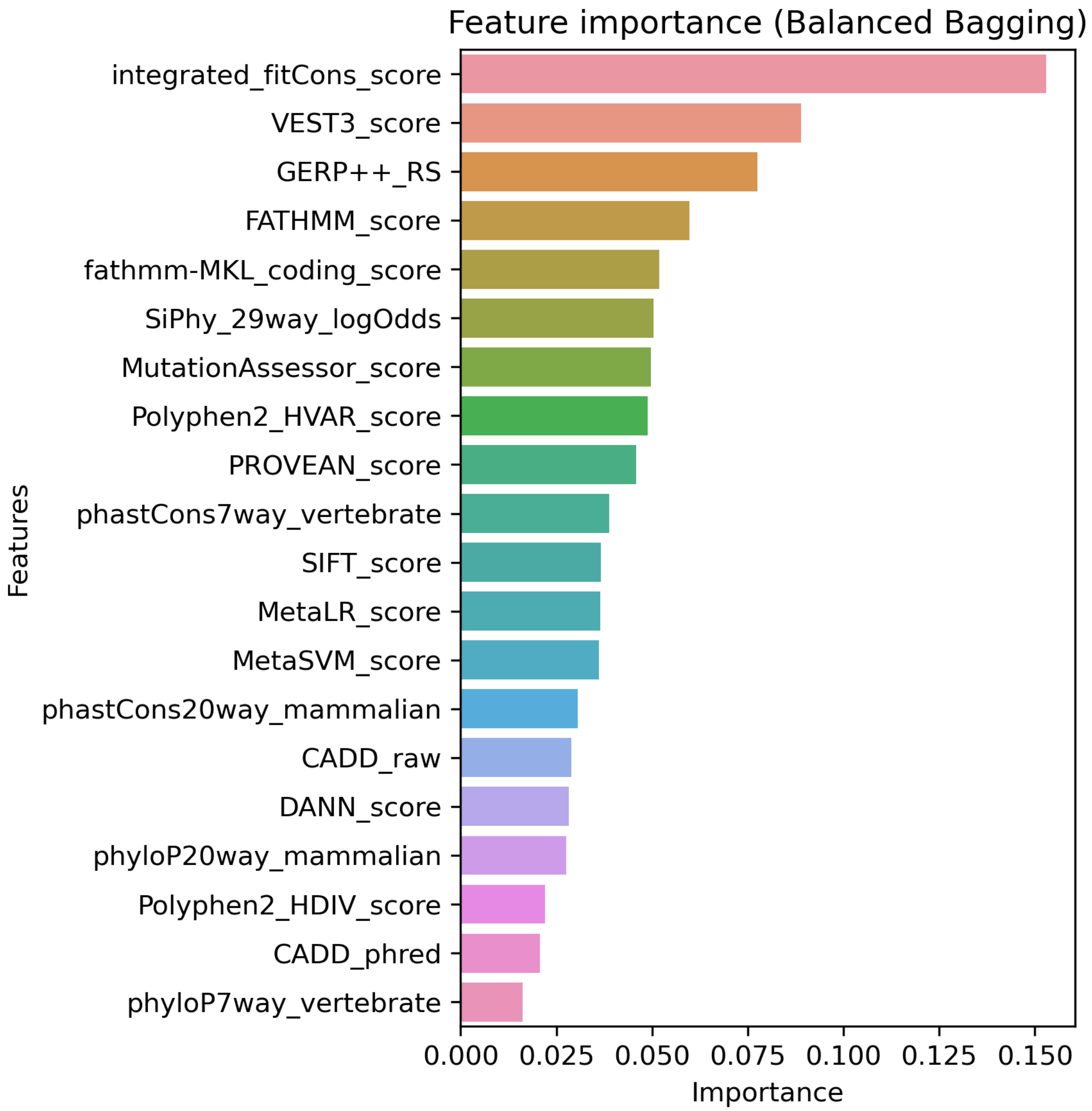


**Figure S3A. Top 20 features for BRCA using SNV data with CGC labels built on "all" features using Balanced bagging.**

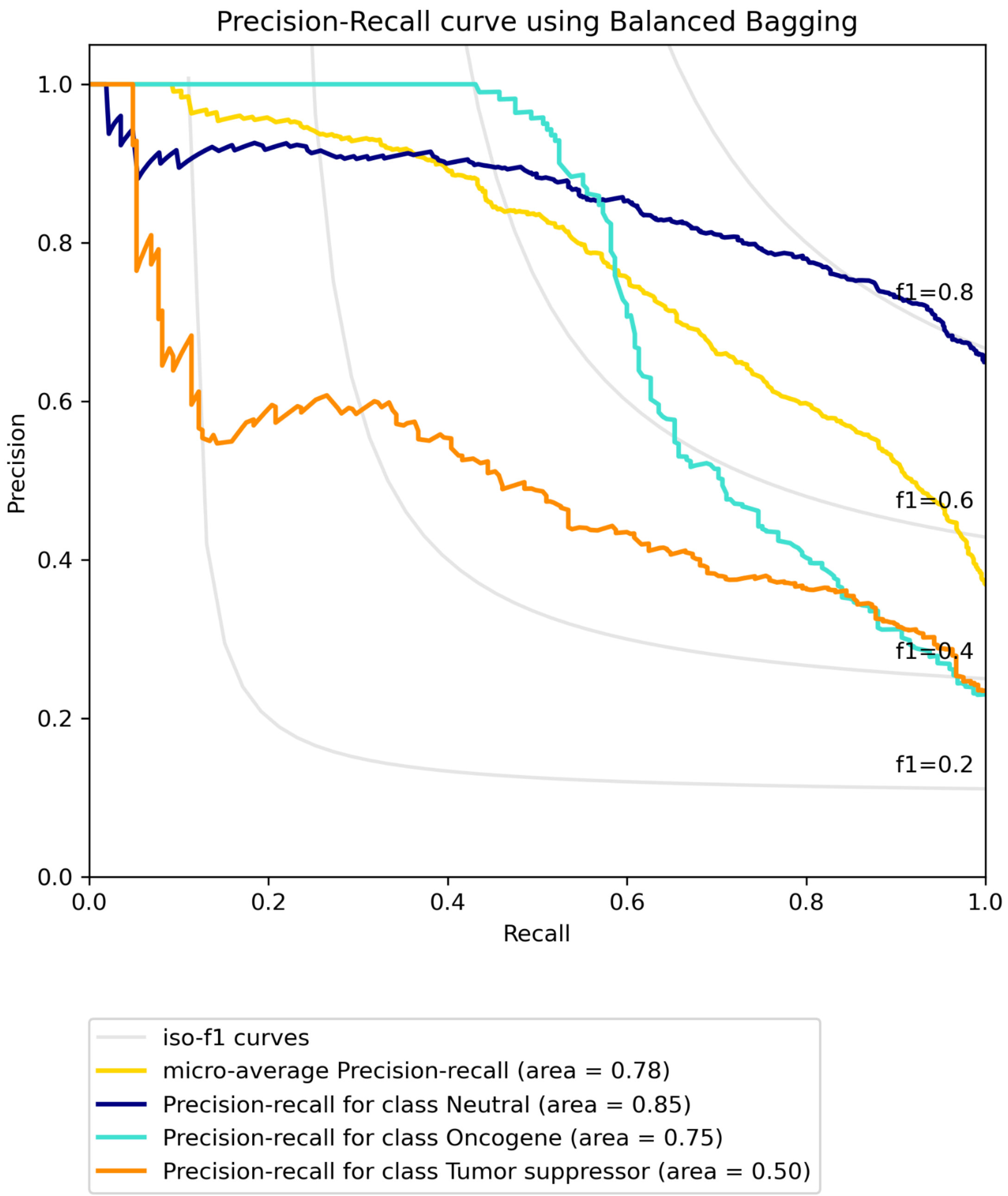


**Figure S3B. Precision-recall curve for BRCA using SNV data with CGC labels built on "all" features using Balanced bagging.**

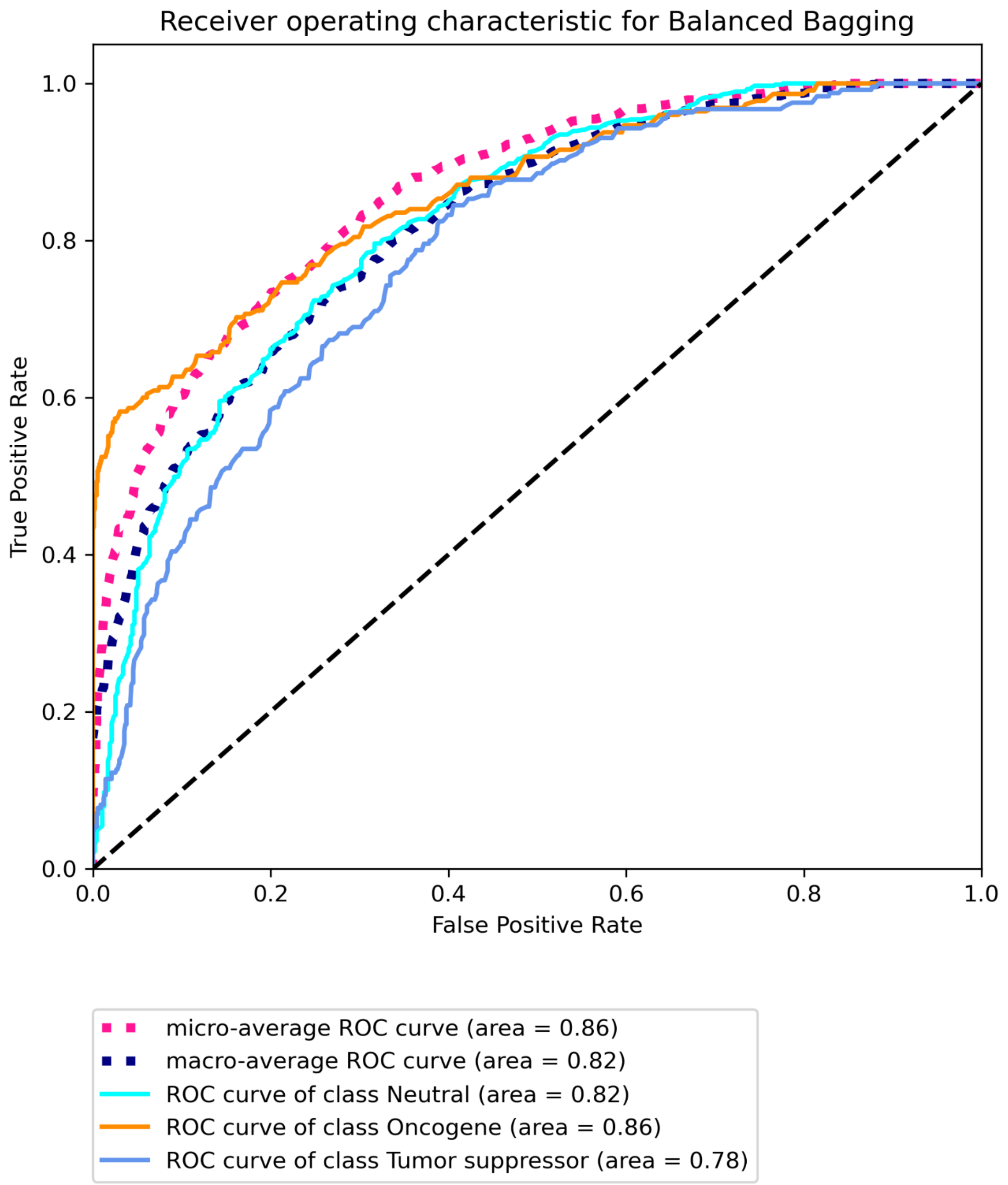


**Figure S3C. Receiver operating characteristic for BRCA using SNV data with CGC labels built on "all" features using Balanced bagging.**

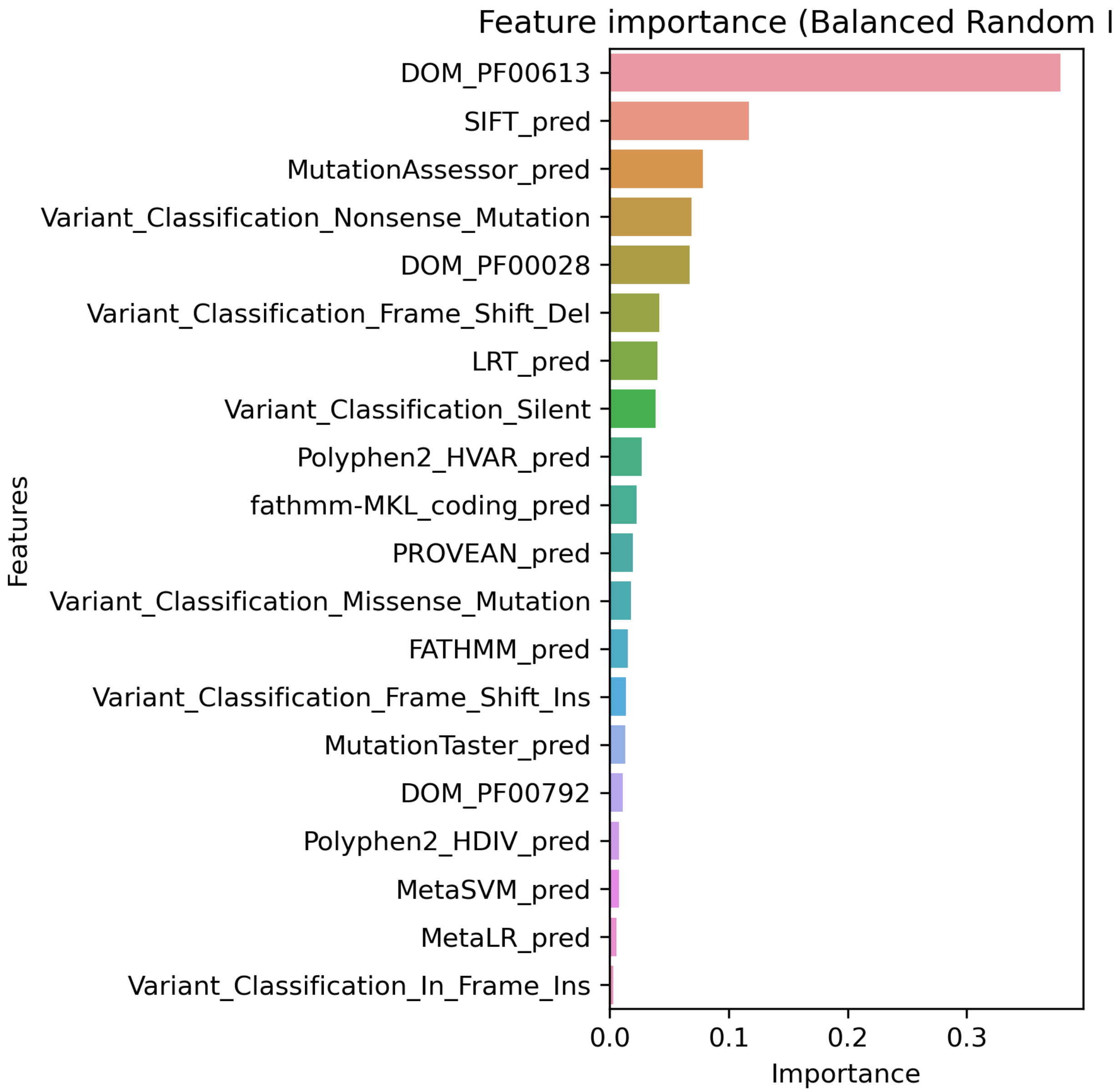


**Figure S4A. Top 20 features for BRCA using SNV data with CGC labels built on "some" features using Balanced Random Forest.**

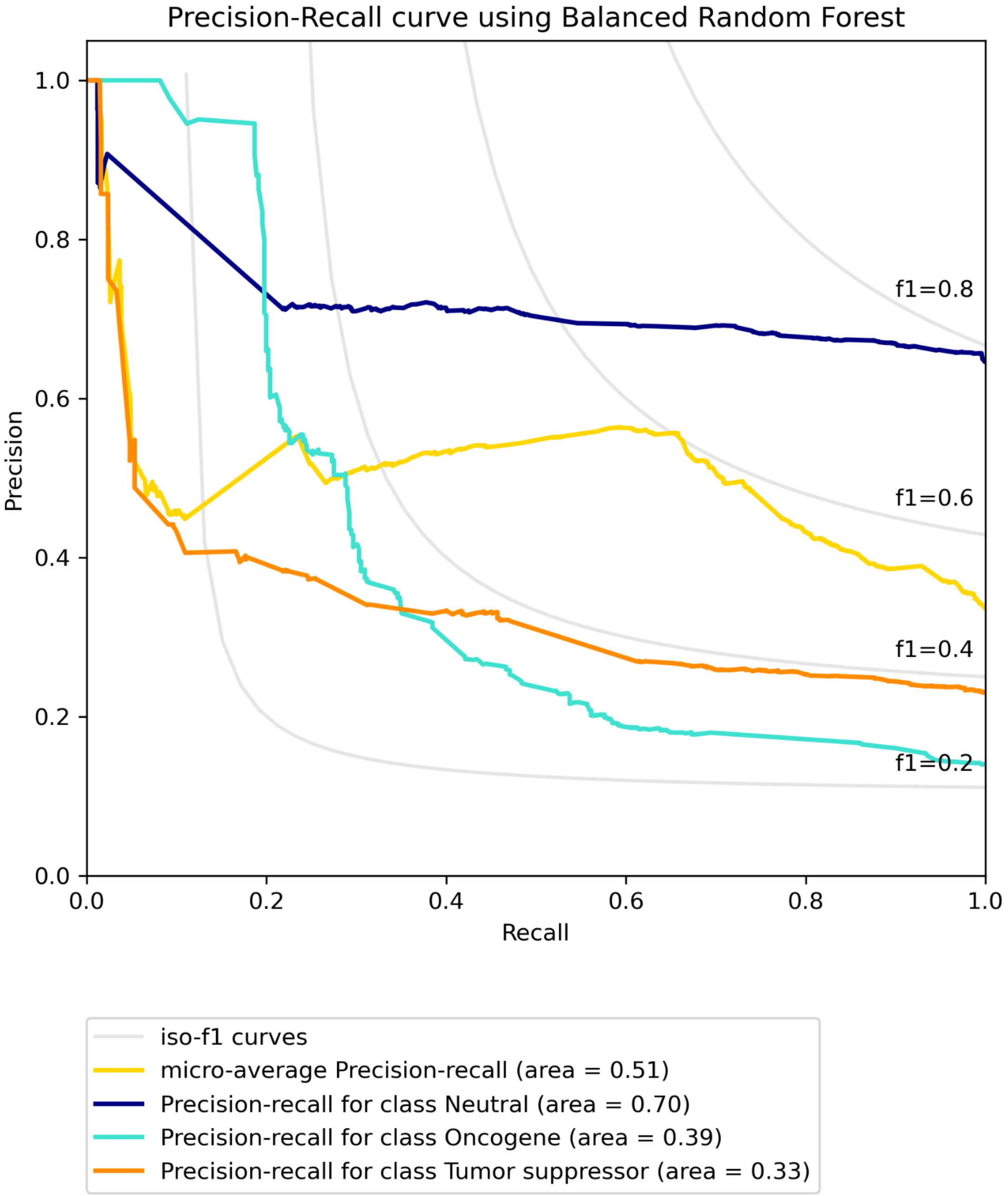


**Figure S4B. Precision-recall curve for BRCA using SNV data with CGC labels built on "some" features using Balanced Random Forest.**

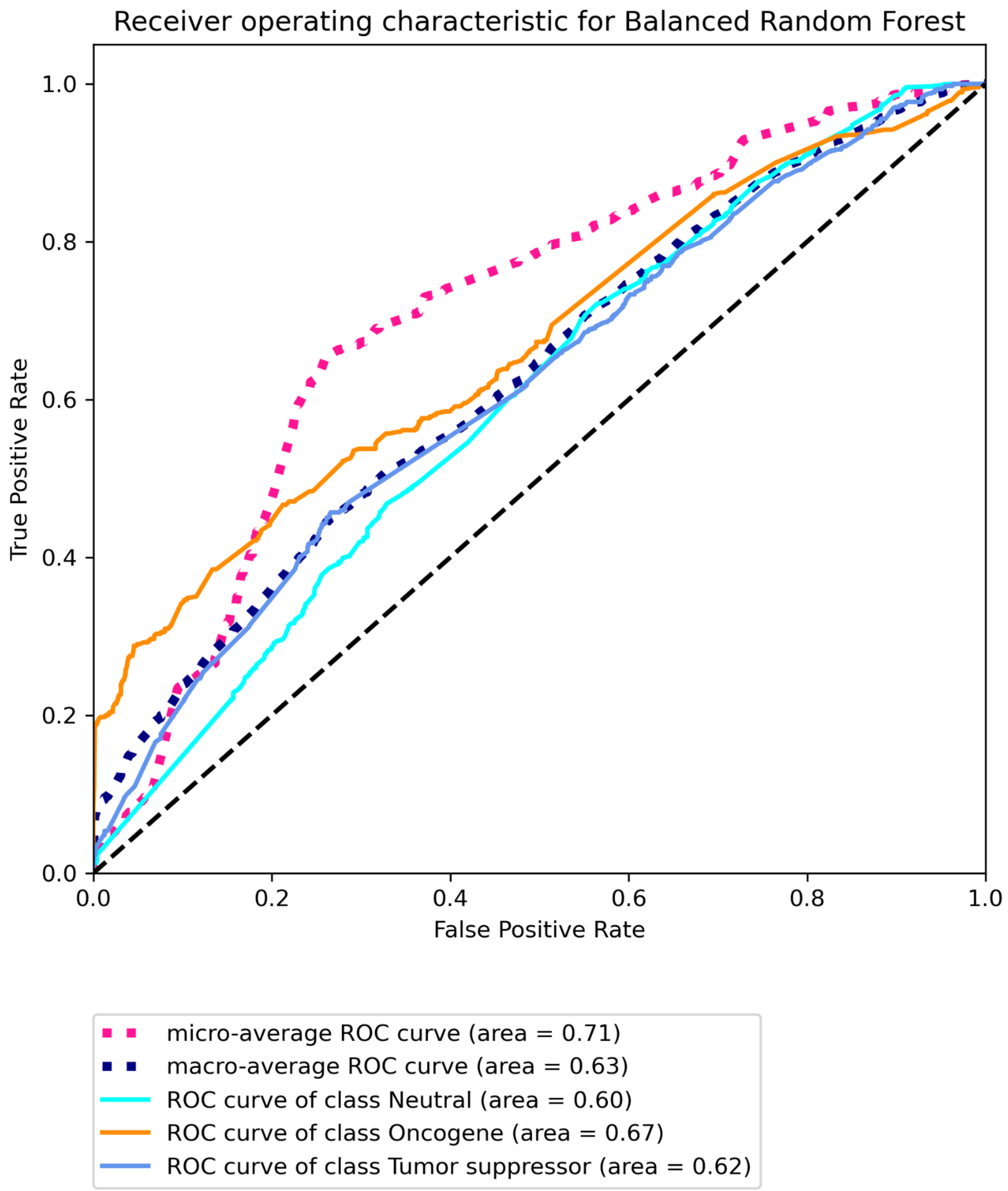


**Figure S4C. Receiver operating characteristic for BRCA using SNV data with CGC labels built on “some” features using Balanced Random Forest.**

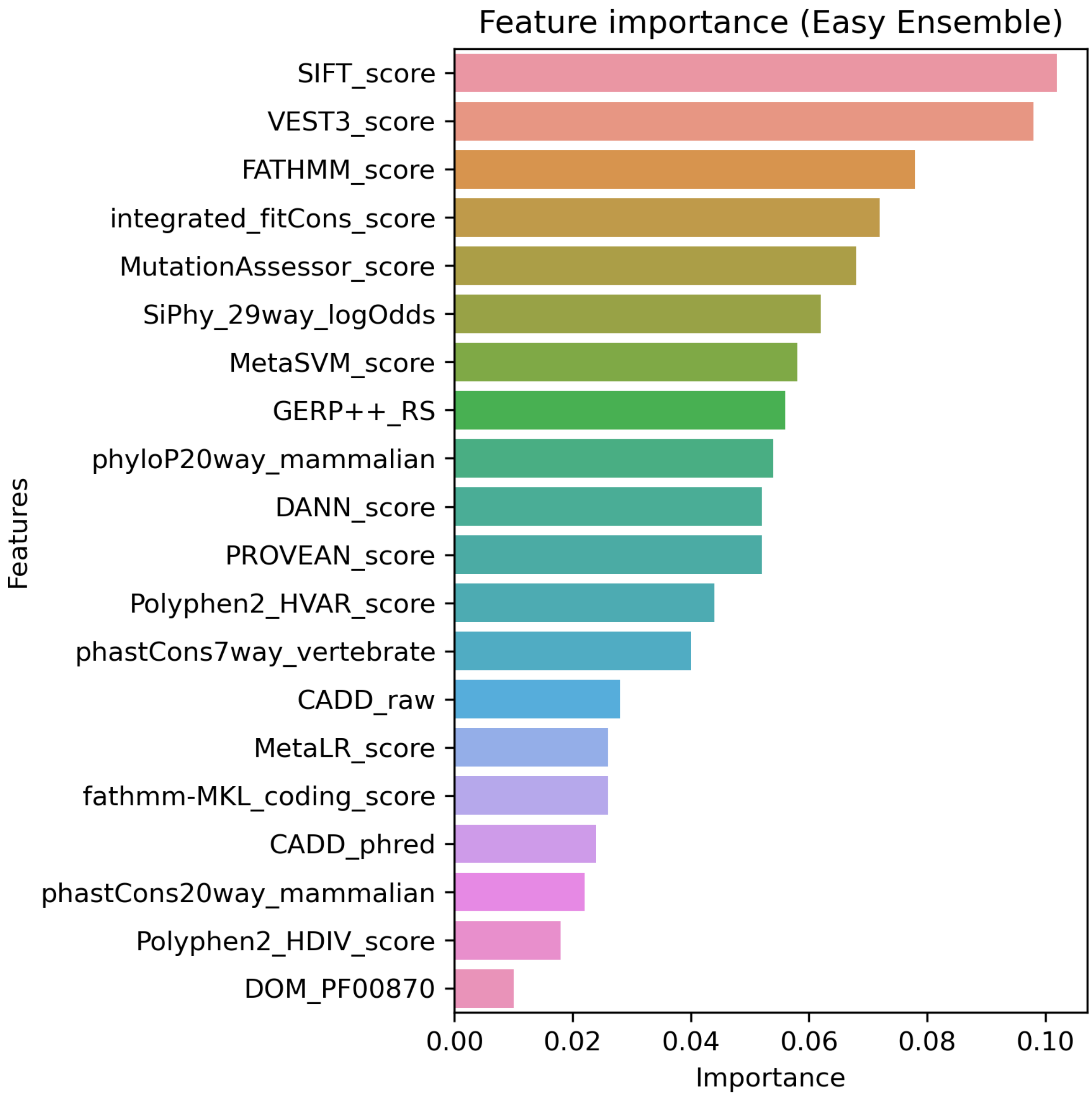


**Figure S5A. Top 20 features for BRCA using SNV data with CIViC labels built on "all" features using Easy Ensemble.**

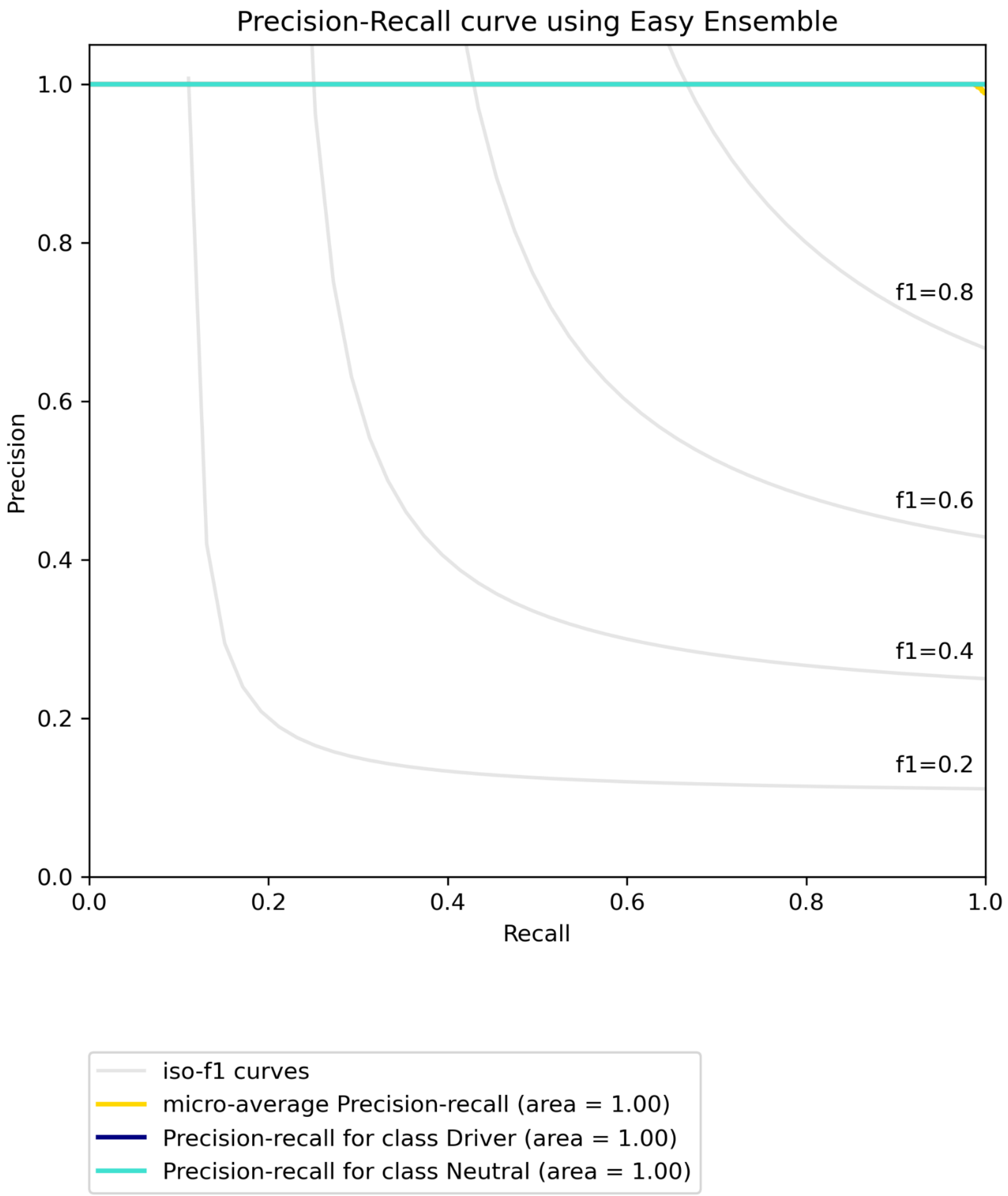


**Figure S5B. Precision-recall curve for BRCA using SNV data with CIViC labels built on "all" features using Easy Ensemble.**

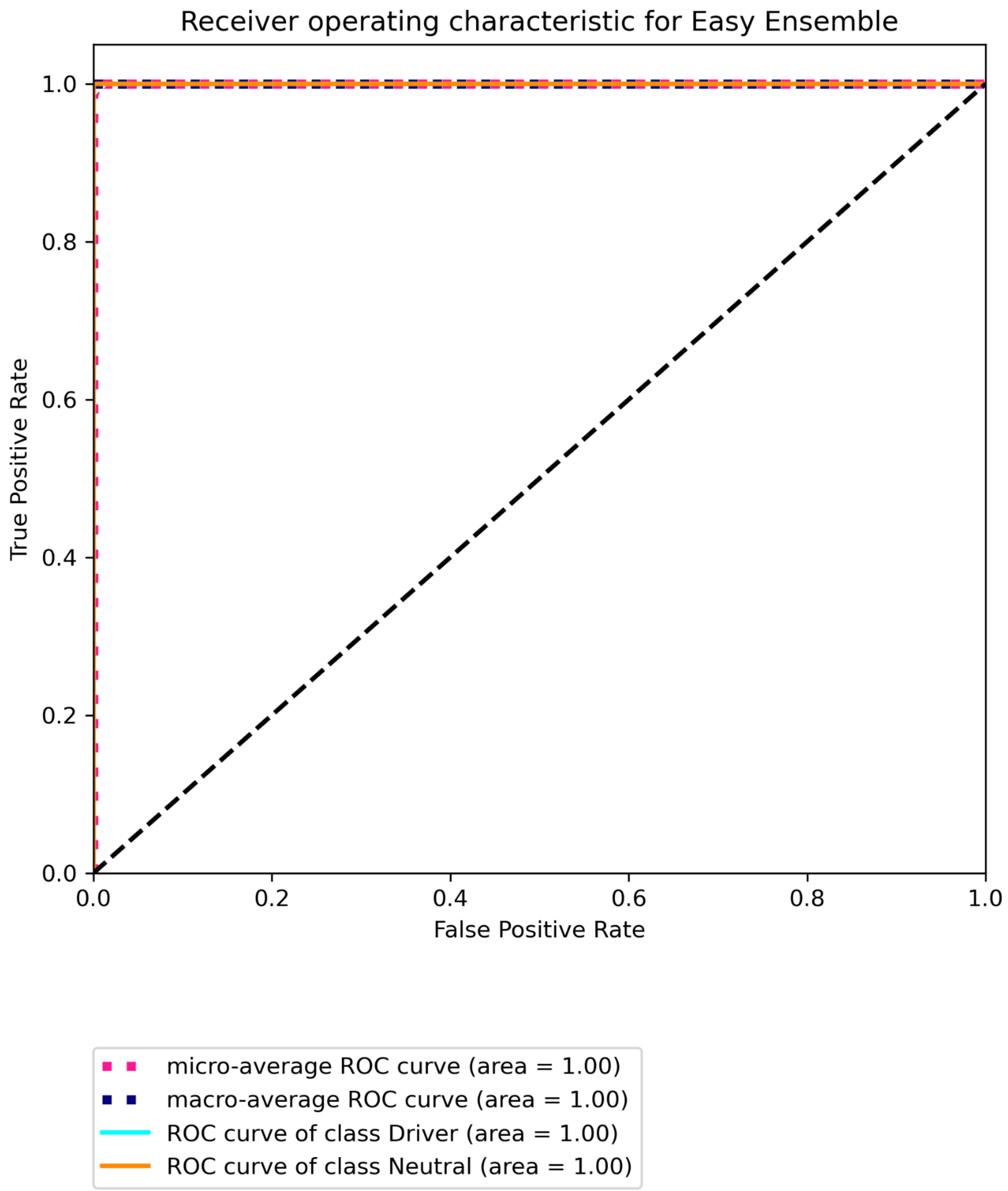


**Figure S5C. Receiver operating characteristic for BRCA using SNV data with CIViC labels built on "all" features using Easy Ensemble.**

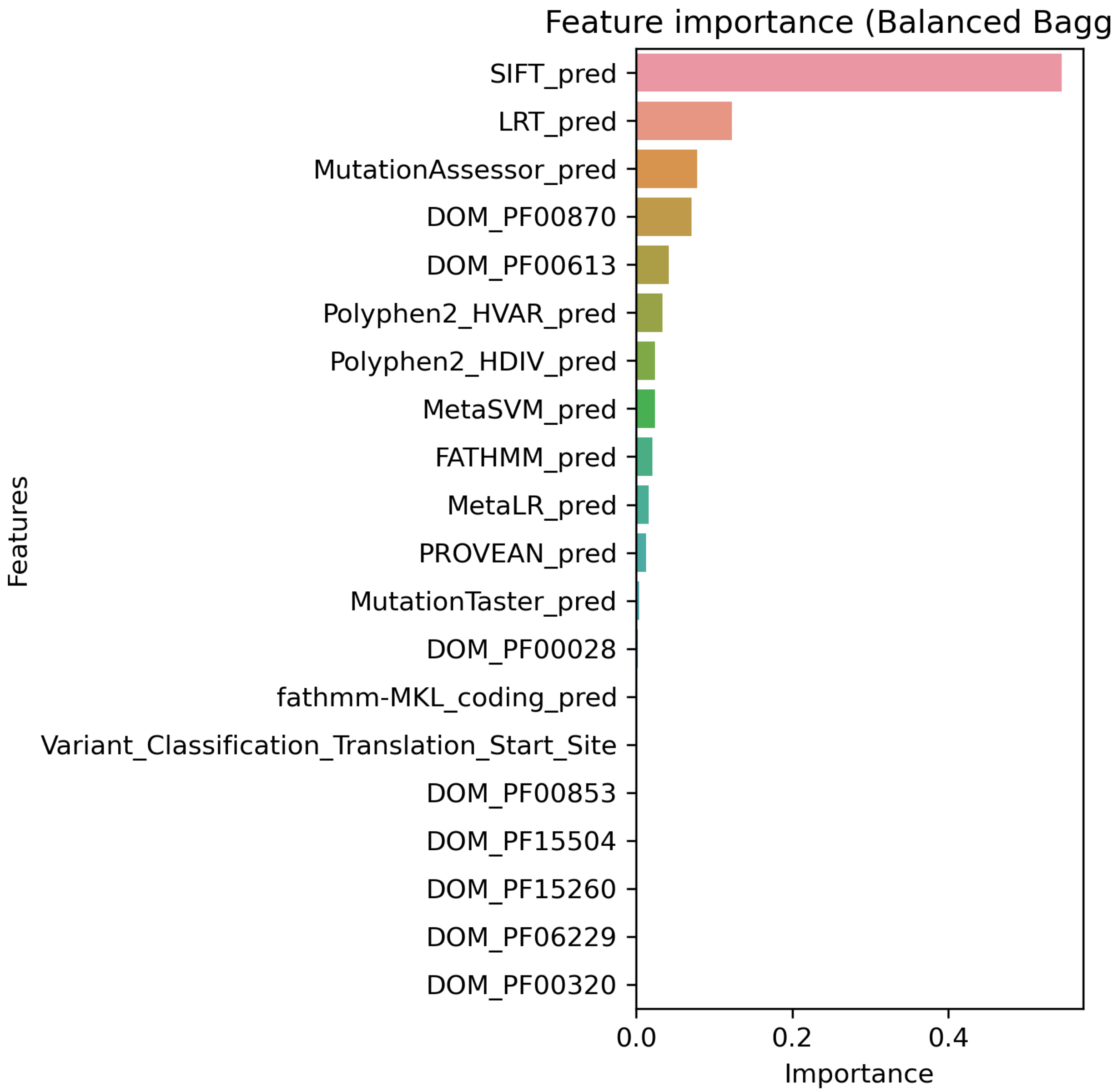


**Figure S6A. Top 20 features for BRCA using SNV data with CIViC labels built on "some" features using Balanced Bagging.**

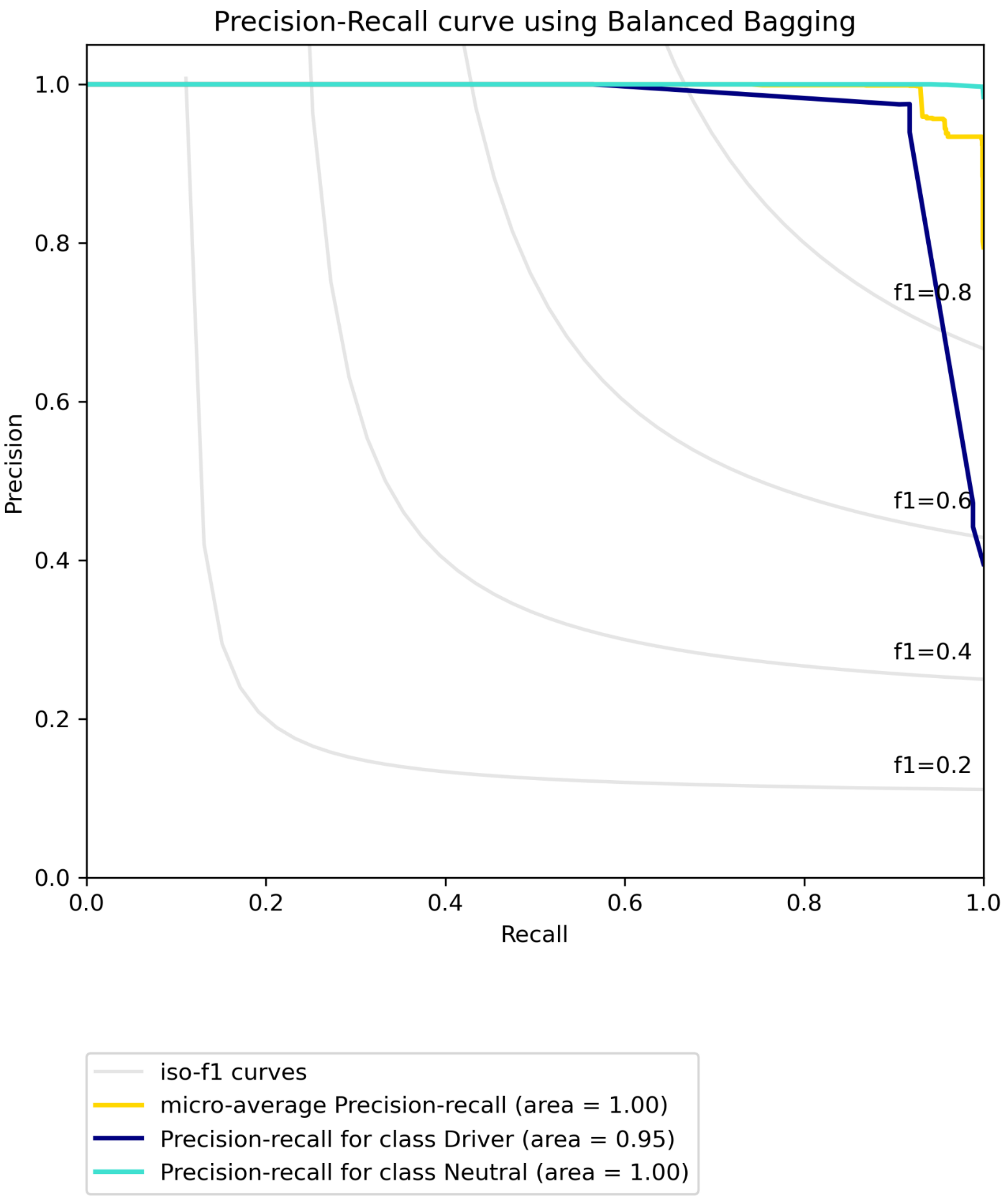


**Figure S6B. Precision-recall curve for BRCA using SNV data with CIViC labels built on "some" features using Balanced Bagging.**

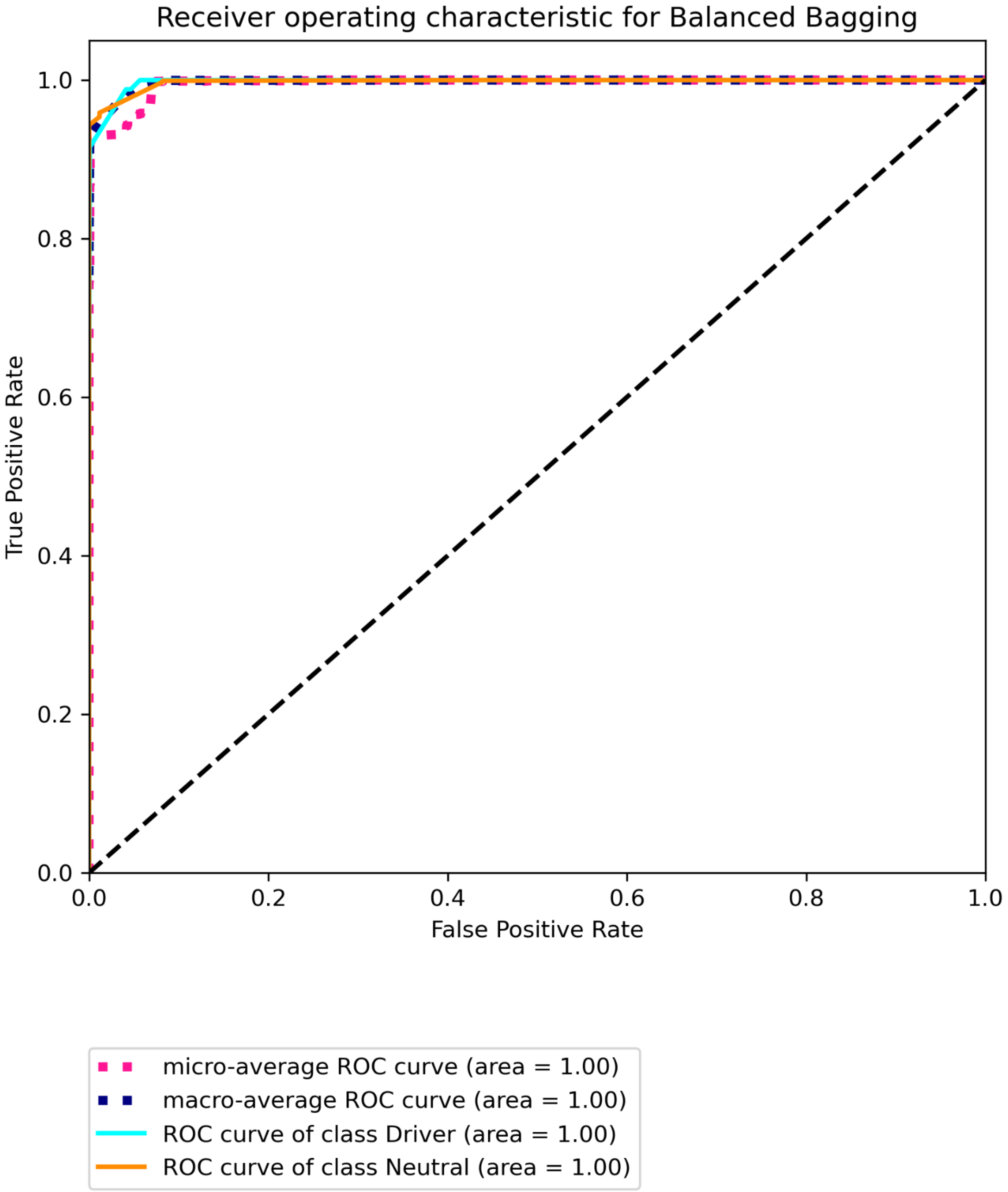


**Figure S6C. Receiver operating characteristic for BRCA using SNV data with CIViC labels built on "some" features using Balanced Bagging.**

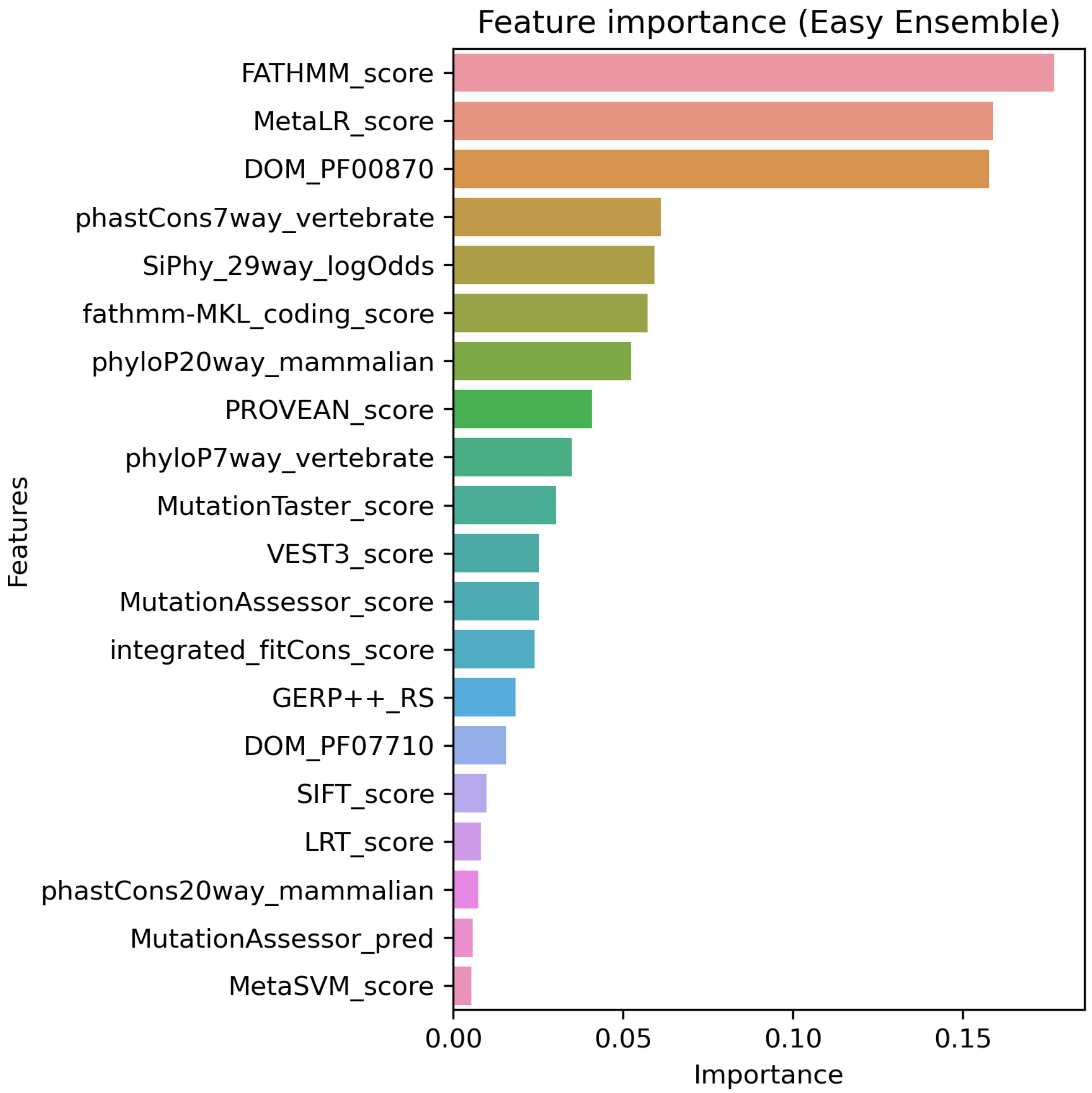


**Figure S7A. Top 20 features for BRCA using SNV data with Marellotto *et al.* labels built on "all" features using Easy Ensemble.**

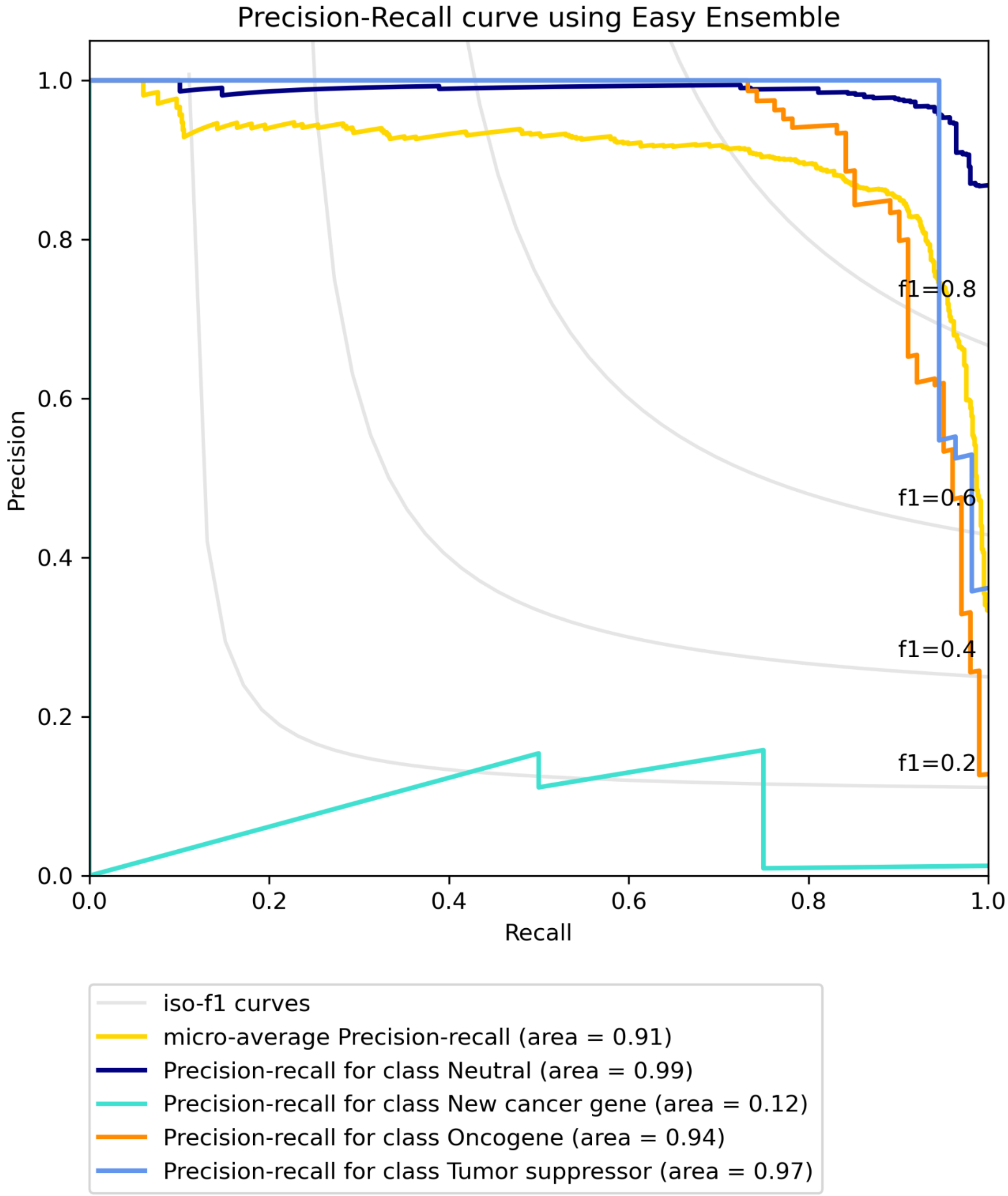


**Figure S7B. Precision-recall curve for BRCA using SNV data with Marellotto *et al.* labels built on "all" features using Easy Ensemble.**

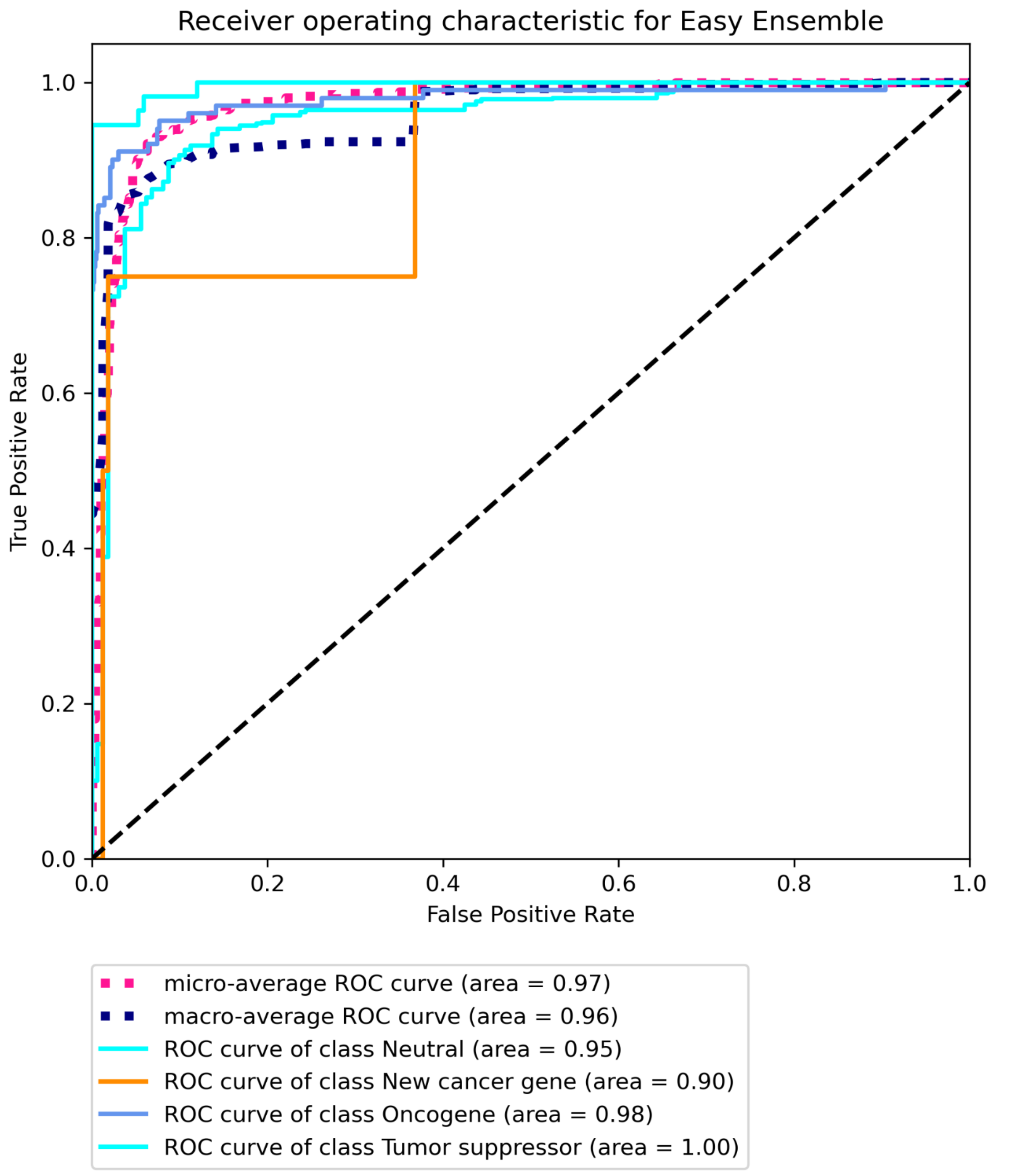


**Figure S7C. Receiver operating characteristic for BRCA using SNV data with Marellotto *et al.* labels built on "all" features using Easy Ensemble.**

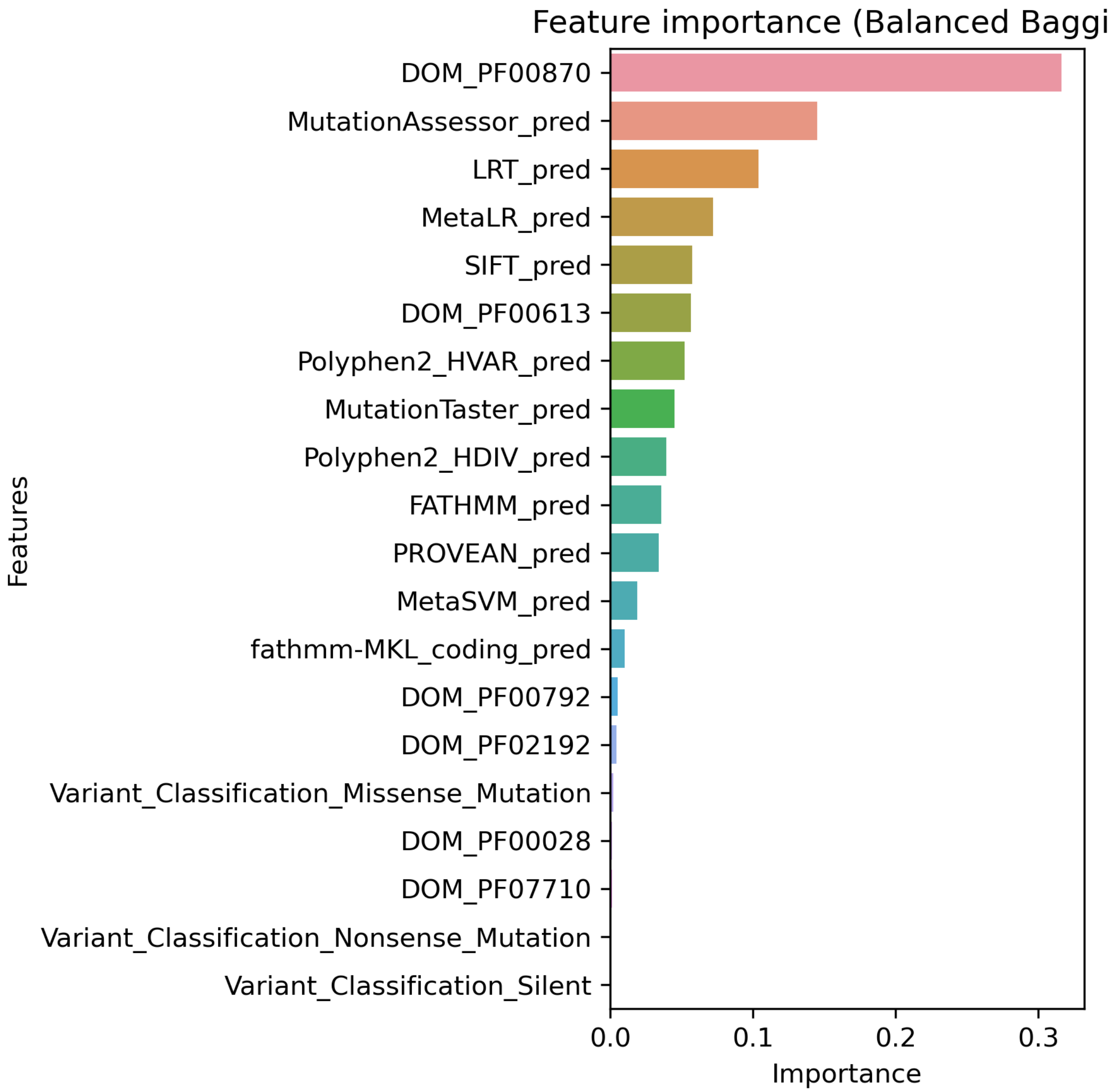


**Figure S8A. Top 20 features for BRCA using SNV data with Marellotto *et al.* labels built on "some" features using Balanced Bagging.**

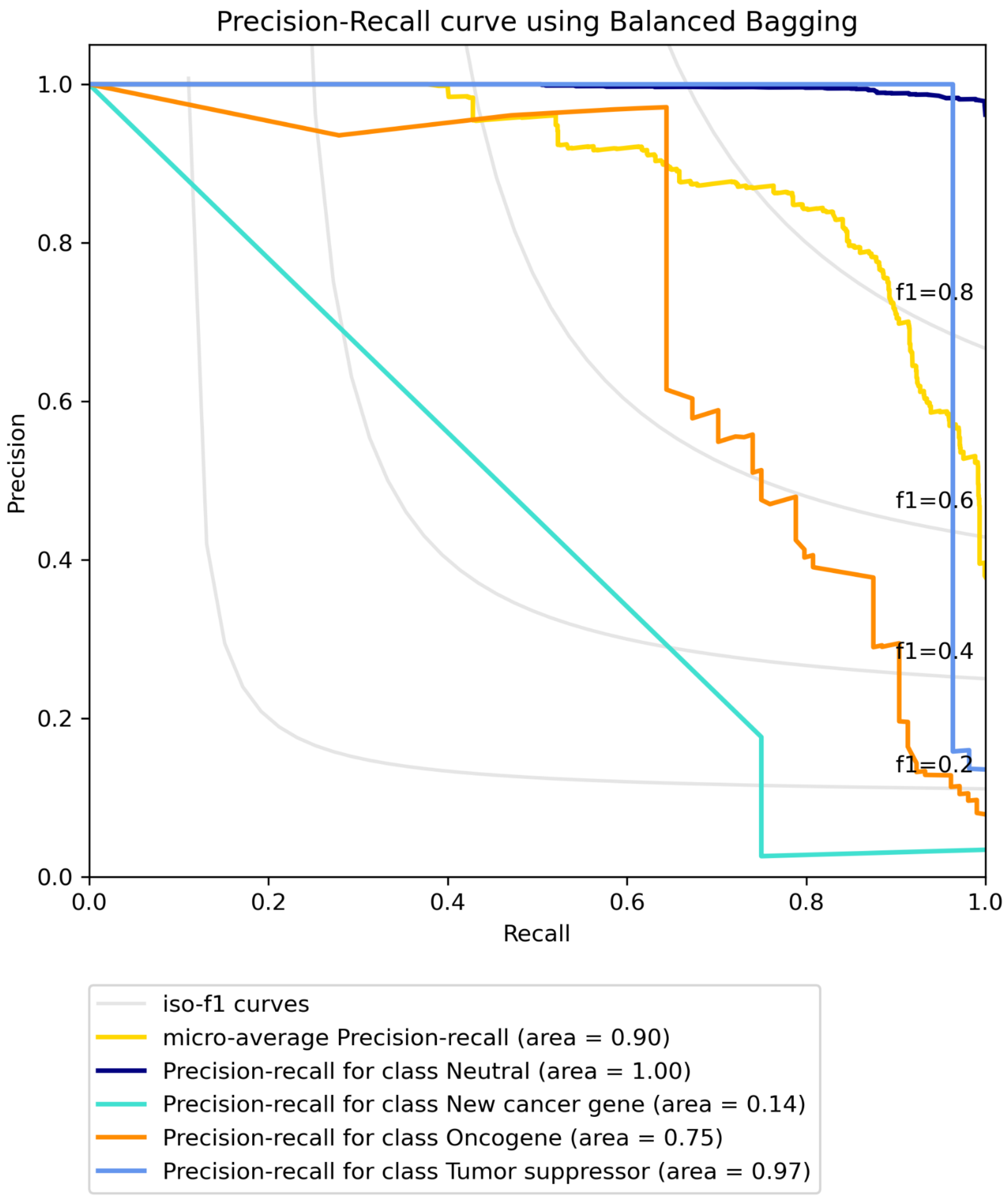


**Figure S8B. Precision-recall curve for BRCA using SNV data with Marellotto *et al.* labels built on "some" features using Balanced Bagging.**

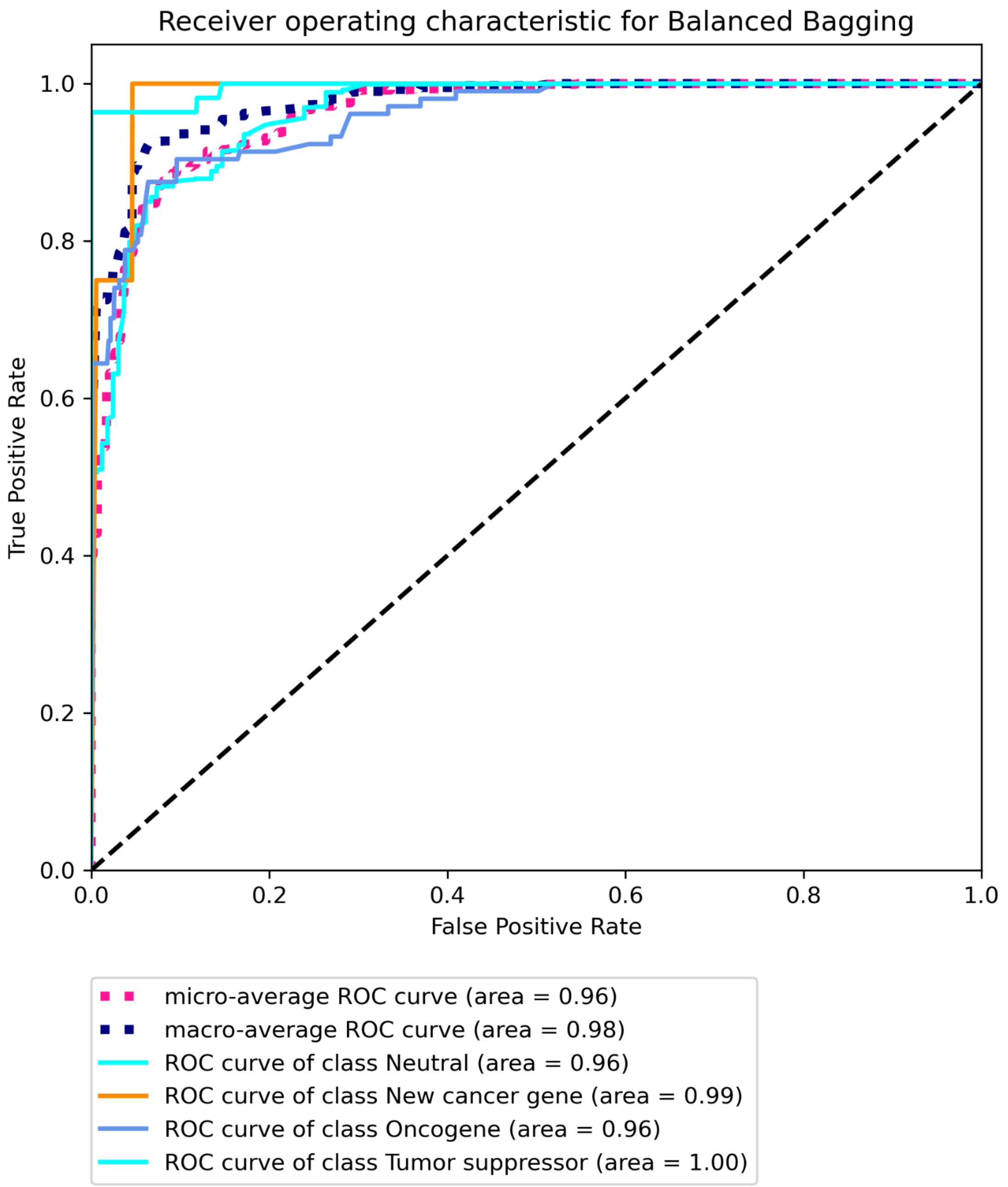


**Figure S8C. Receiver operating characteristic for BRCA using SNV data with Marellotto *et al.* labels built on "some" features using Balanced Bagging.**

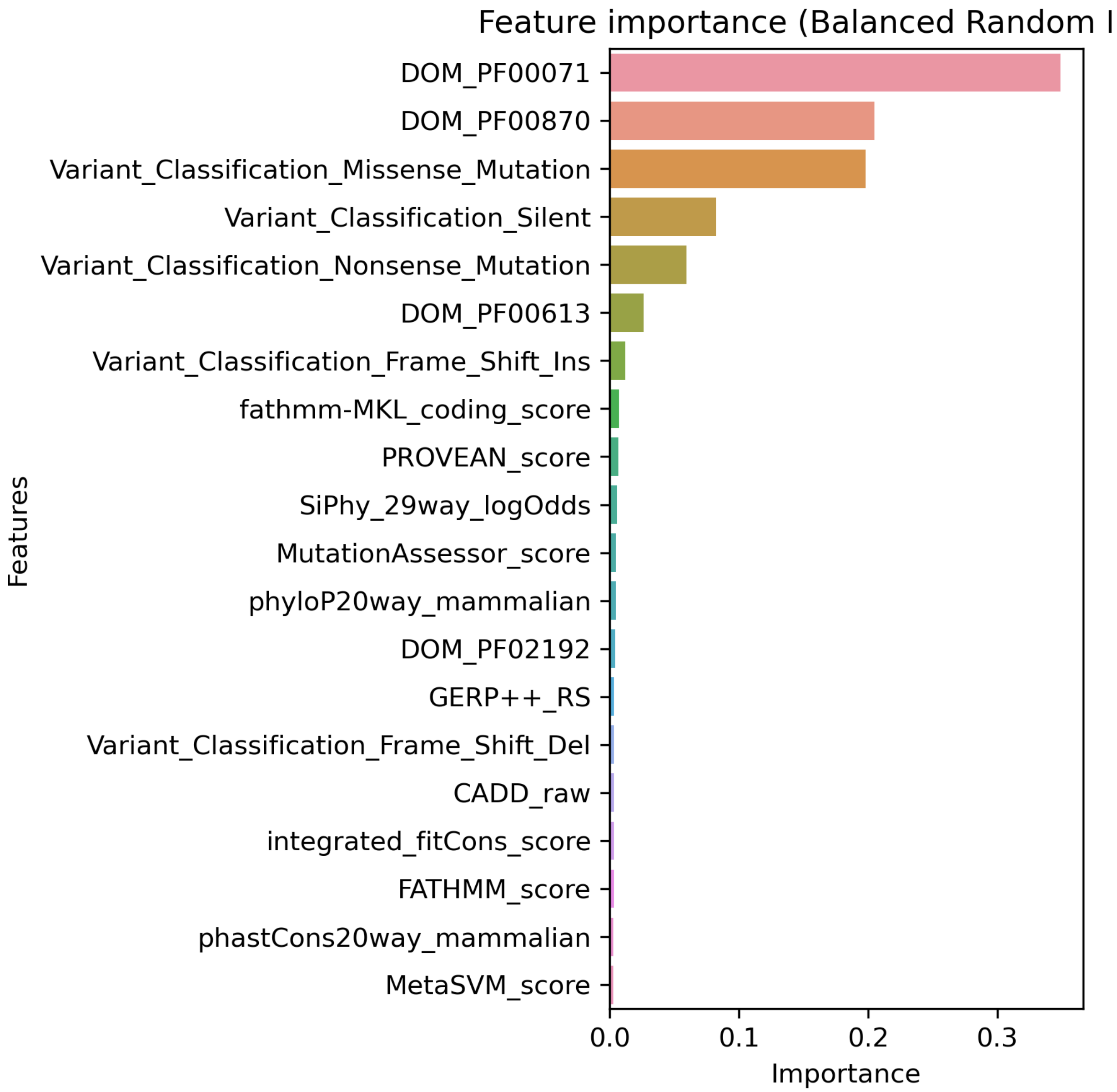


**Figure S9A. Top 20 features for COAD using SNV data with Bailey *et al.* labels built on "all" features using Balanced Random Forest.**

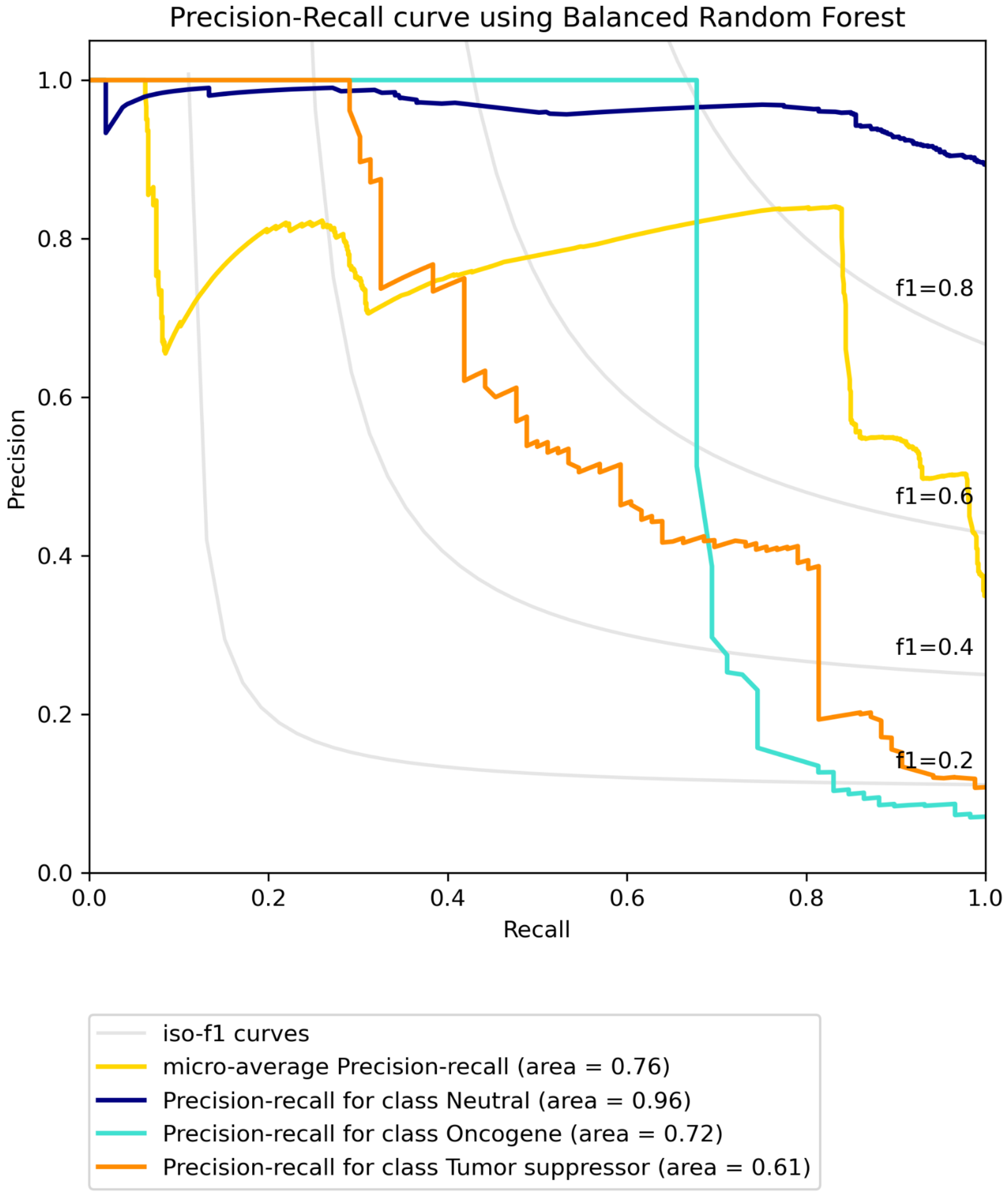


**Figure S9B. Precision-recall curve for COAD using SNV data with Bailey *et al.* labels built on "all" features using Balanced Random Forest.**

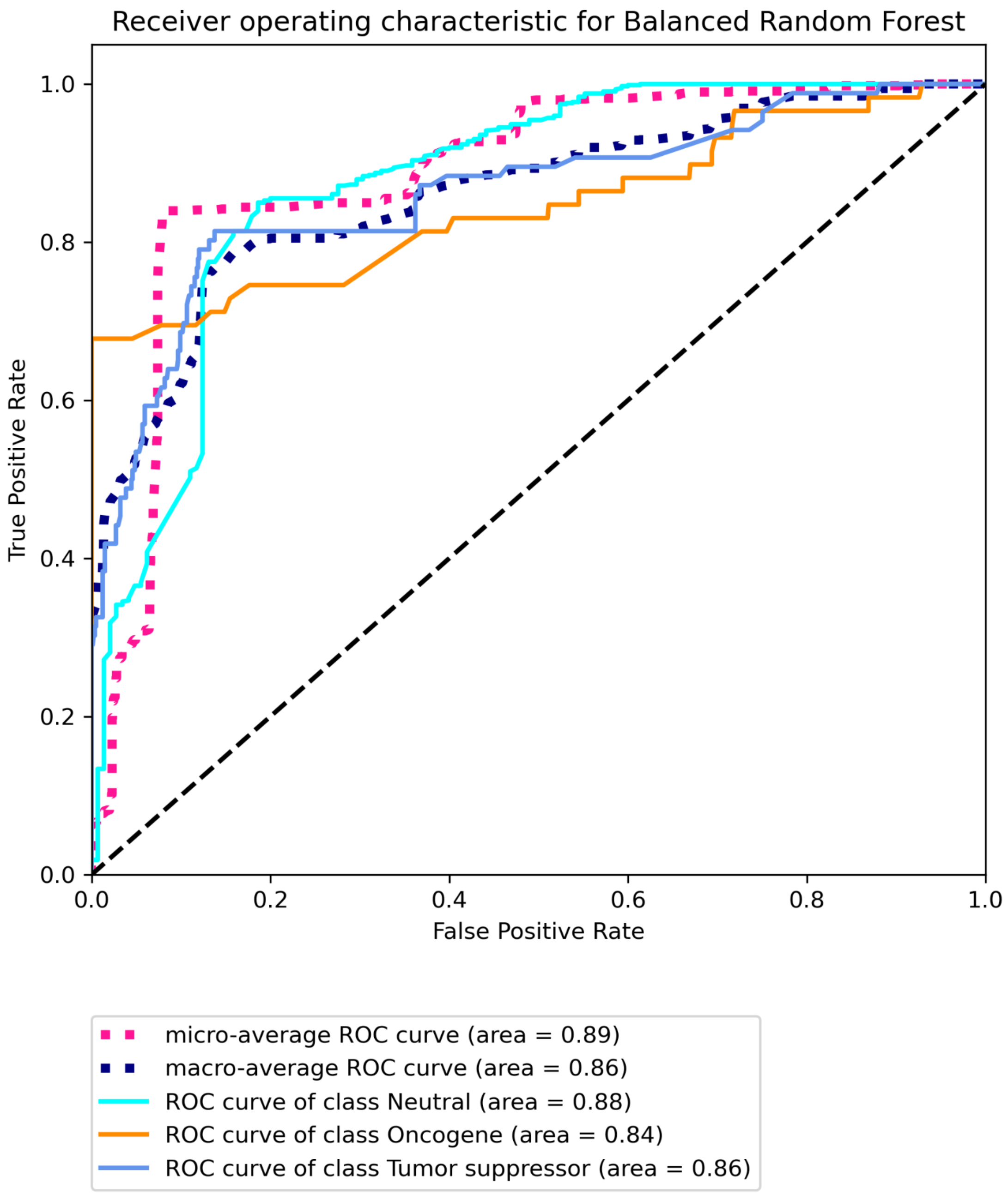


**Figure S9C Receiver operating characteristic for COAD using SNV data with Bailey *et al.* labels built on "all" features using Balanced Random Forest.**

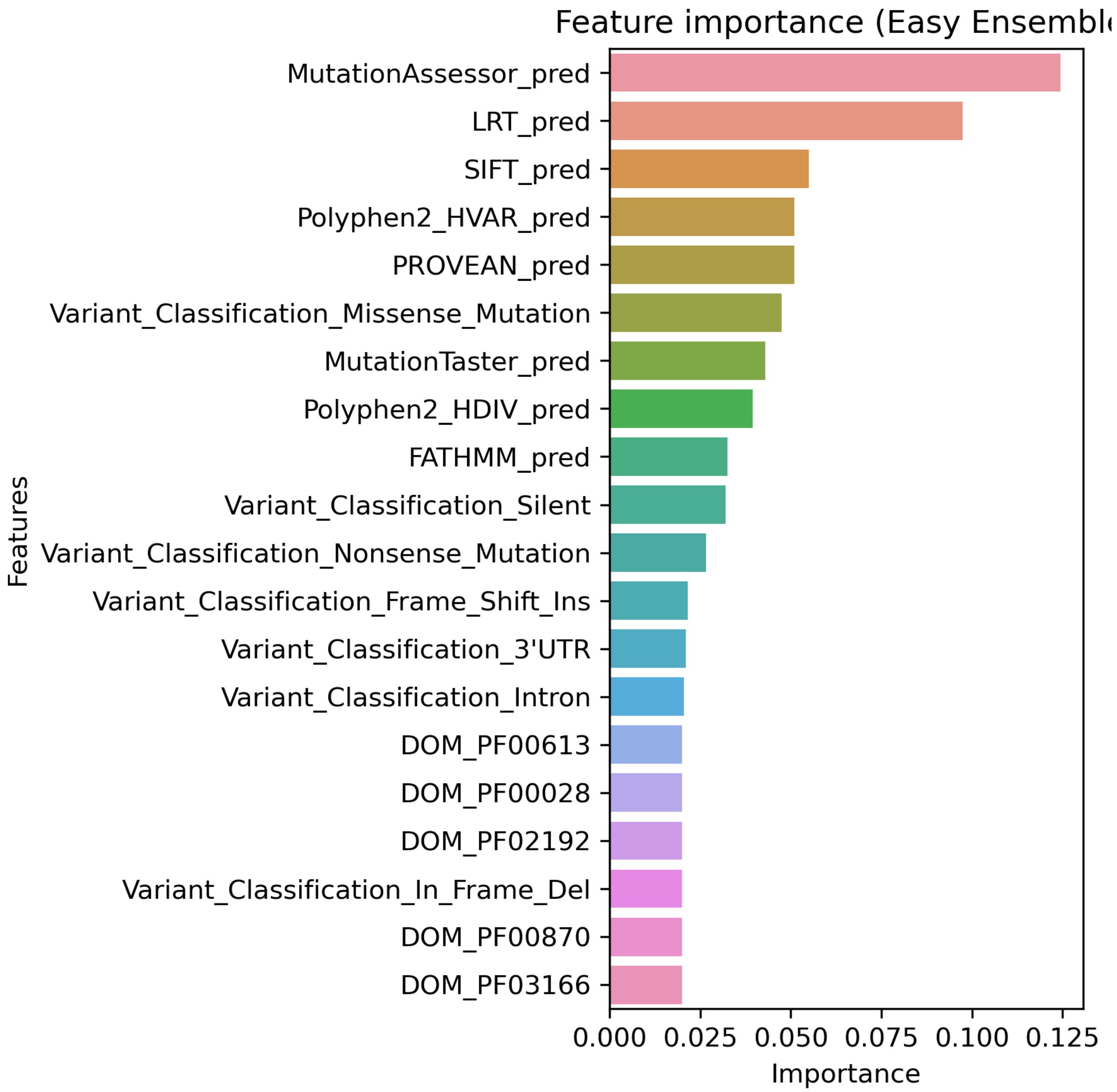


**Figure S10A. Top 20 features for COAD using SNV data with Bailey *et al.* labels built on "some" features using Easy Ensemble.**

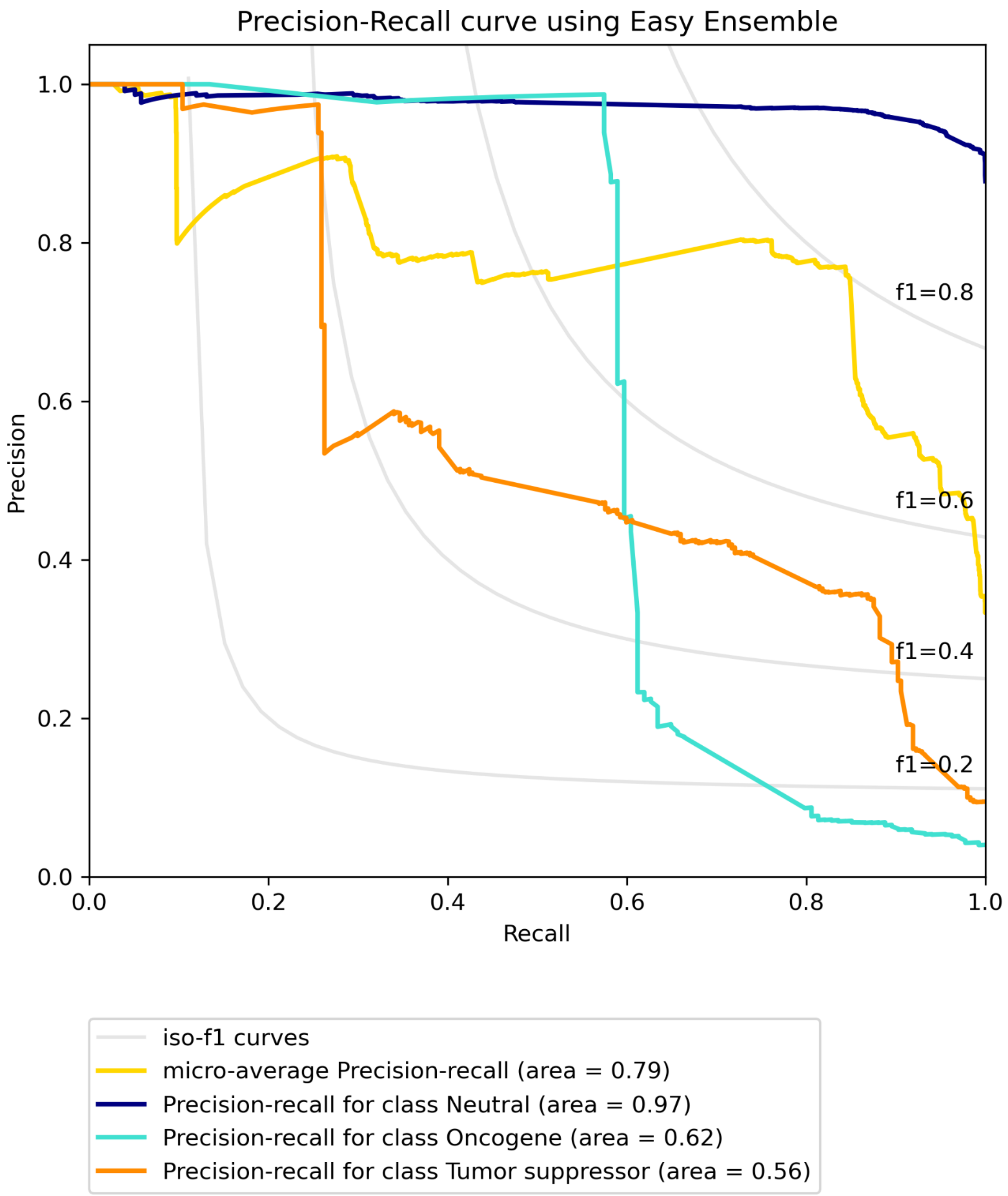


**Figure S10B. Precision-recall curve for COAD using SNV data with Bailey *et al.* labels built on "some" features using Easy Ensemble.**

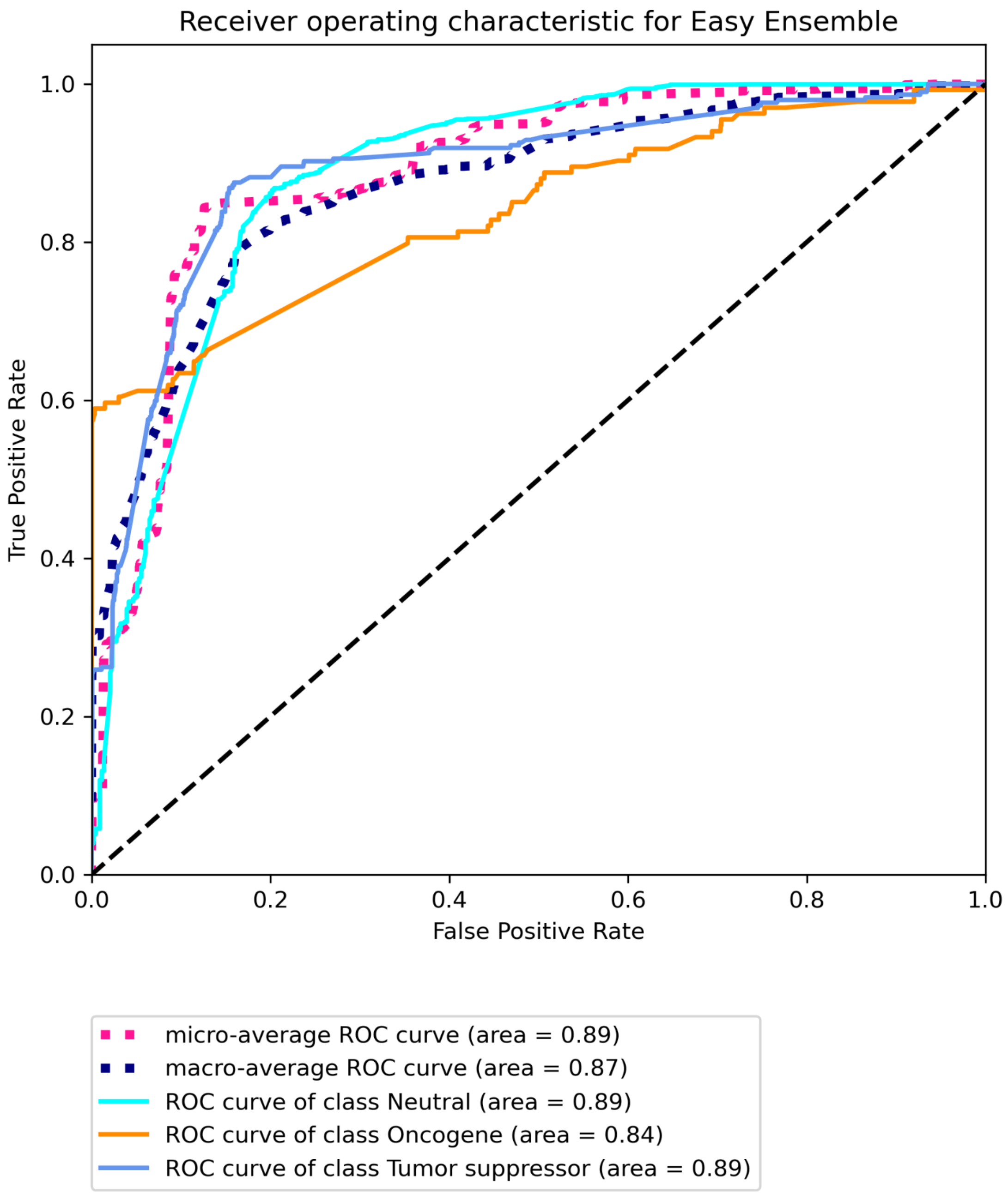


**Figure S10C Receiver operating characteristic for COAD using SNV data with Bailey *et al.* labels built on “some” features using Easy Ensemble.**

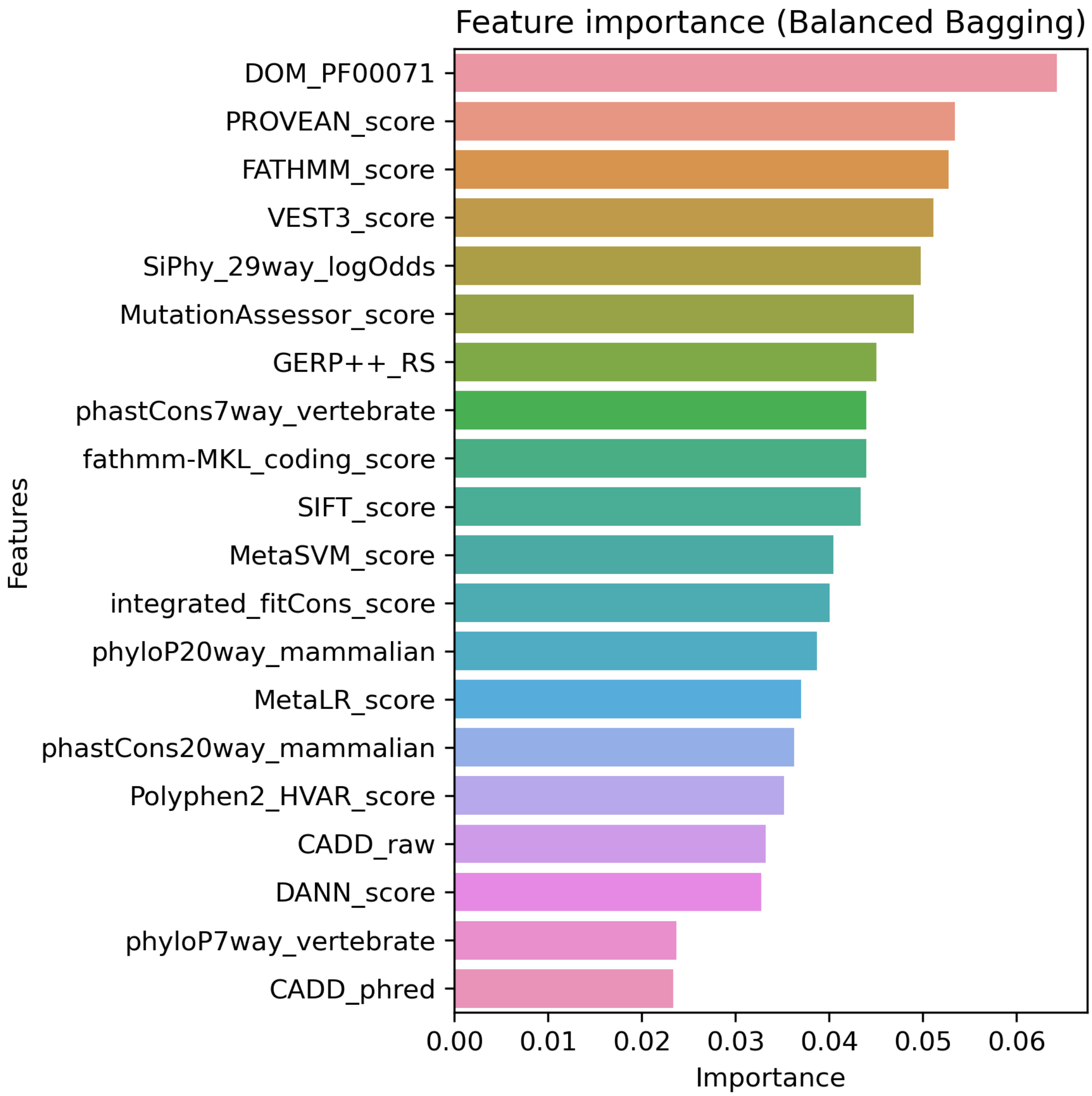


**Figure S11A. Top 20 features for COAD using SNV data with CGC labels built on "all" features using Balanced Bagging.**

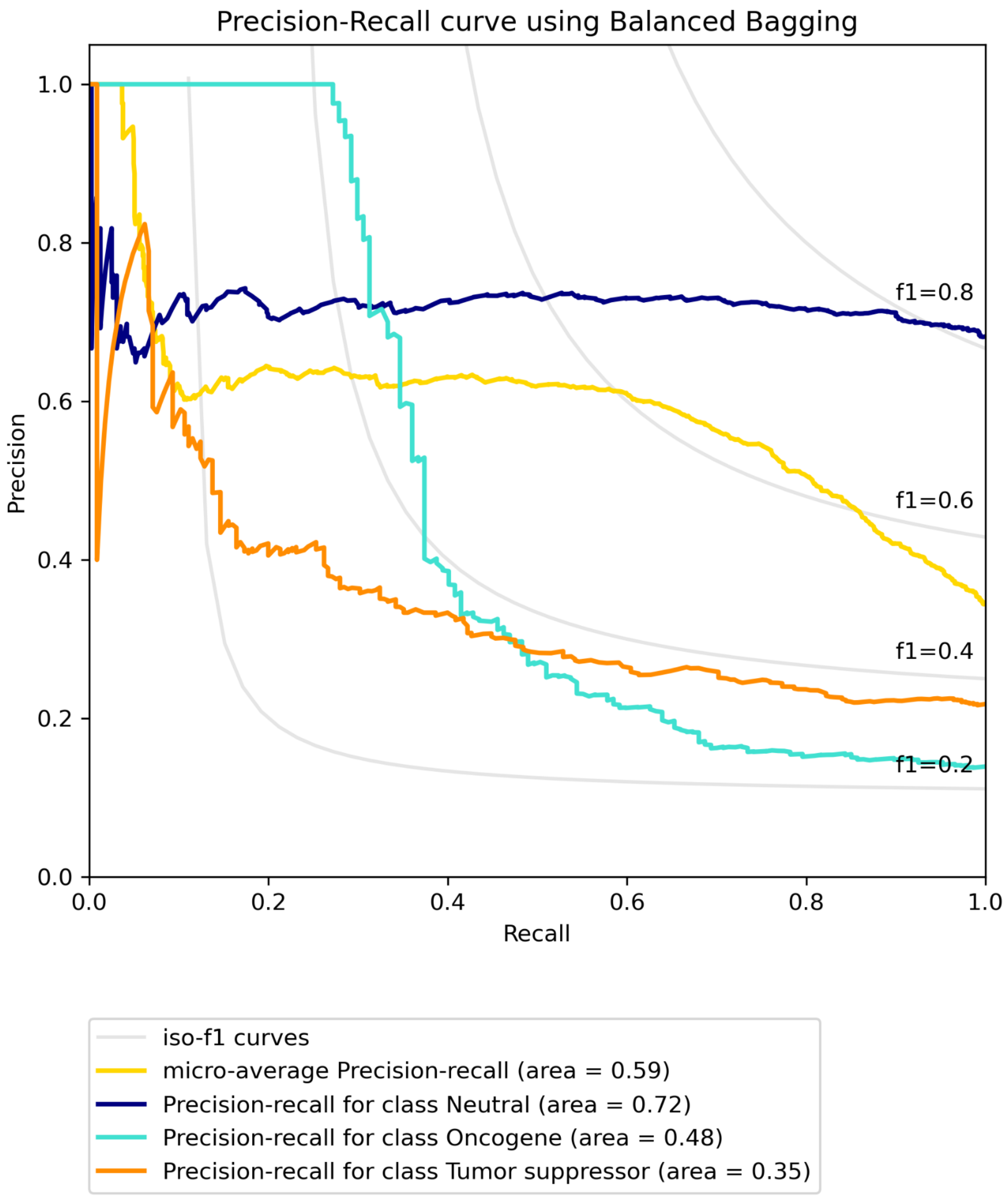


**Figure S11B. Precision-recall curve for COAD using SNV data with CGC labels built on "all" features using Balanced Bagging.**

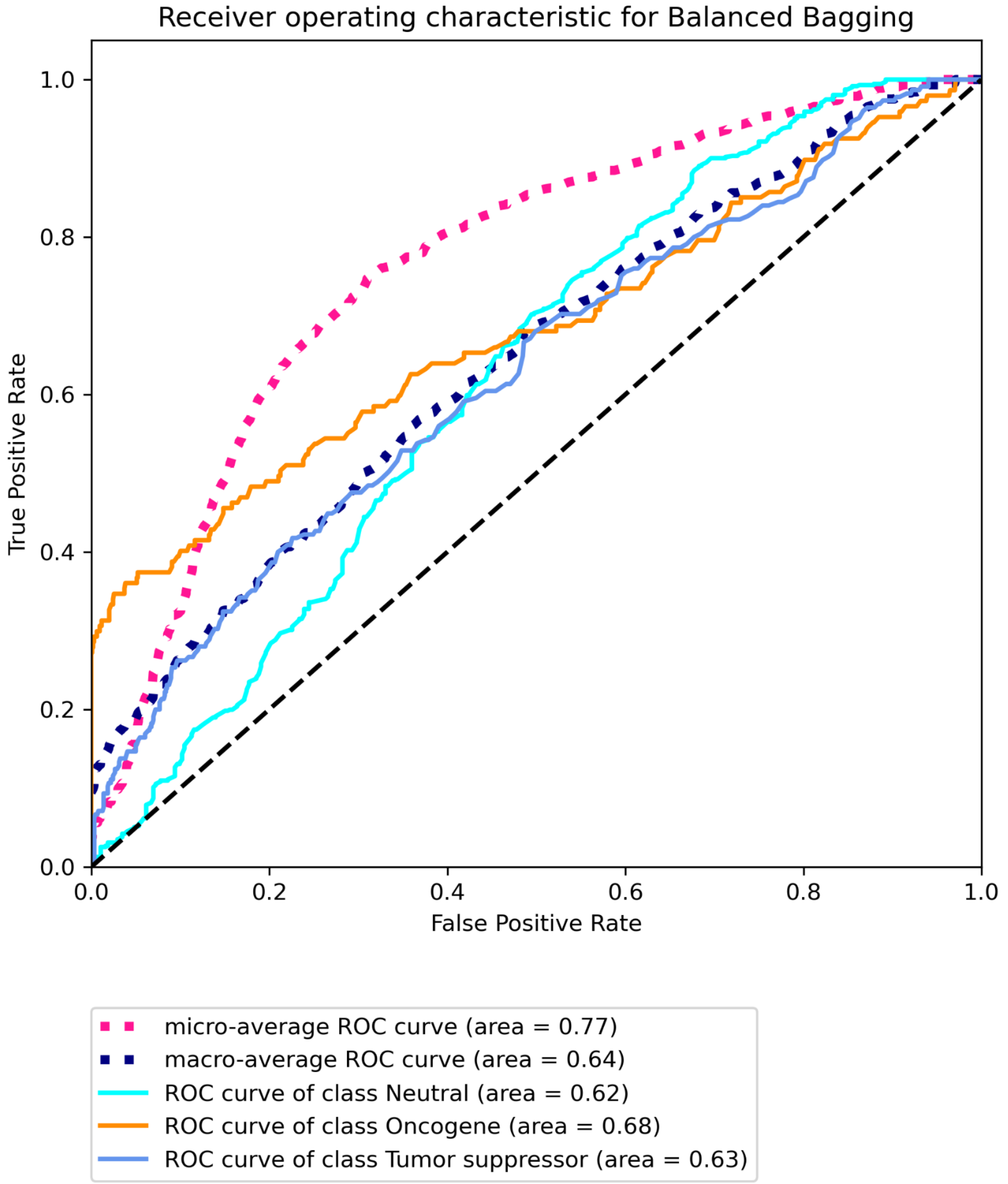


**Figure S11C Receiver operating characteristic for COAD using SNV data with CGC labels built on “all” features using Balanced Bagging.**

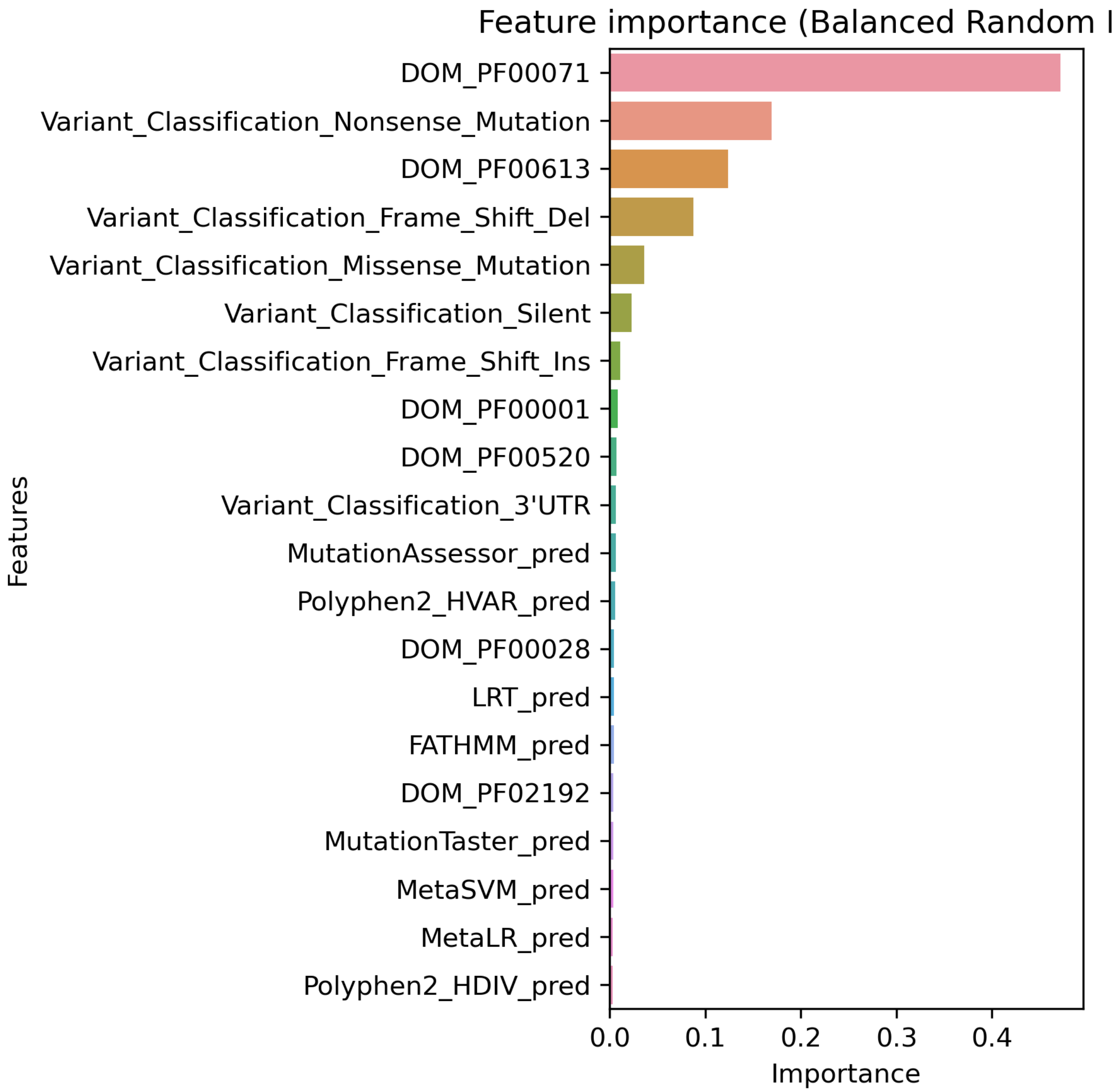


**Figure S12A. Top 20 features for COAD using SNV data with CGC labels built on “some” features using Balanced Random Forest.**

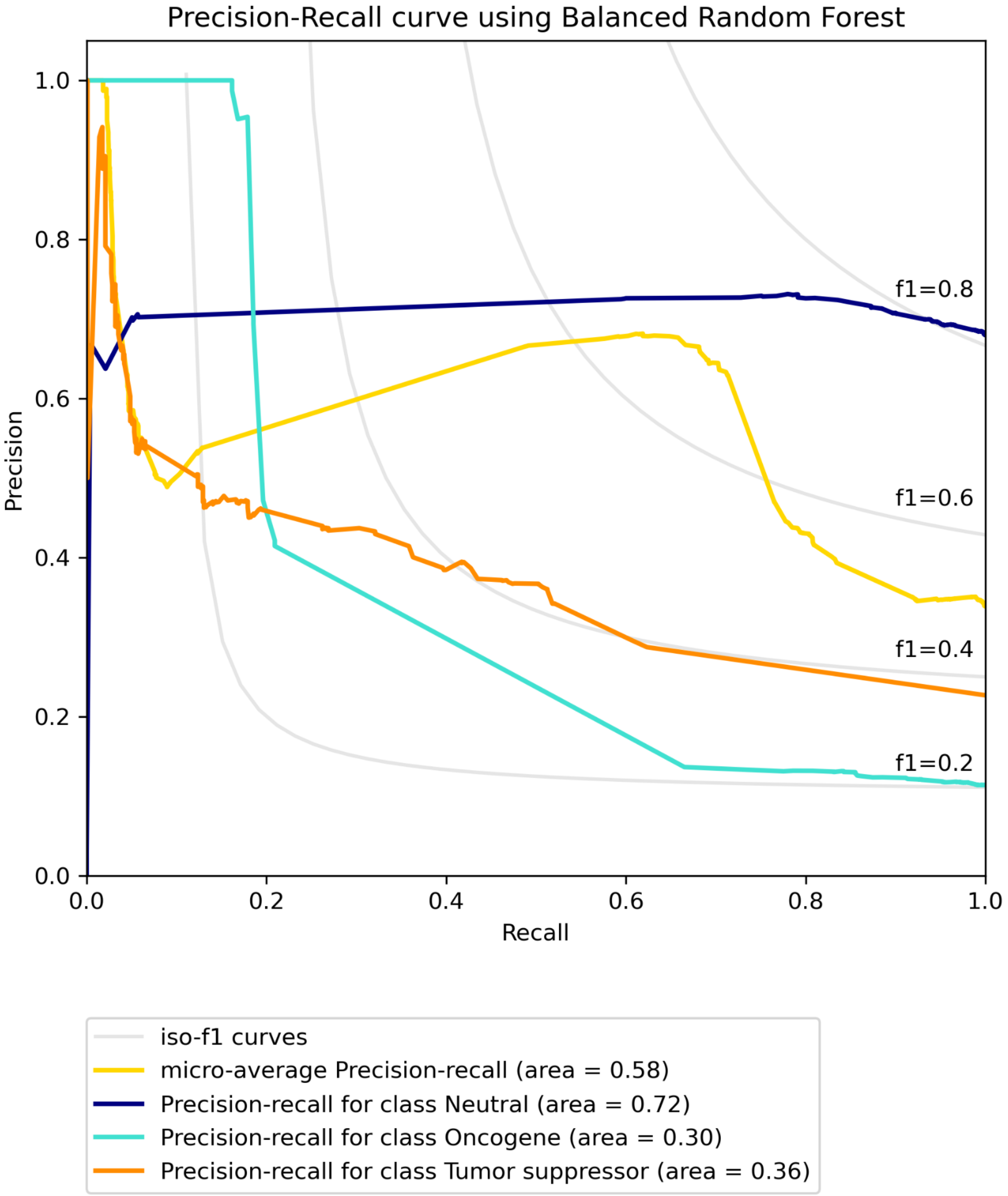


**Figure S12B. Precision-recall curve for COAD using SNV data with CGC labels built on "some" features using Balanced Random Forest.**

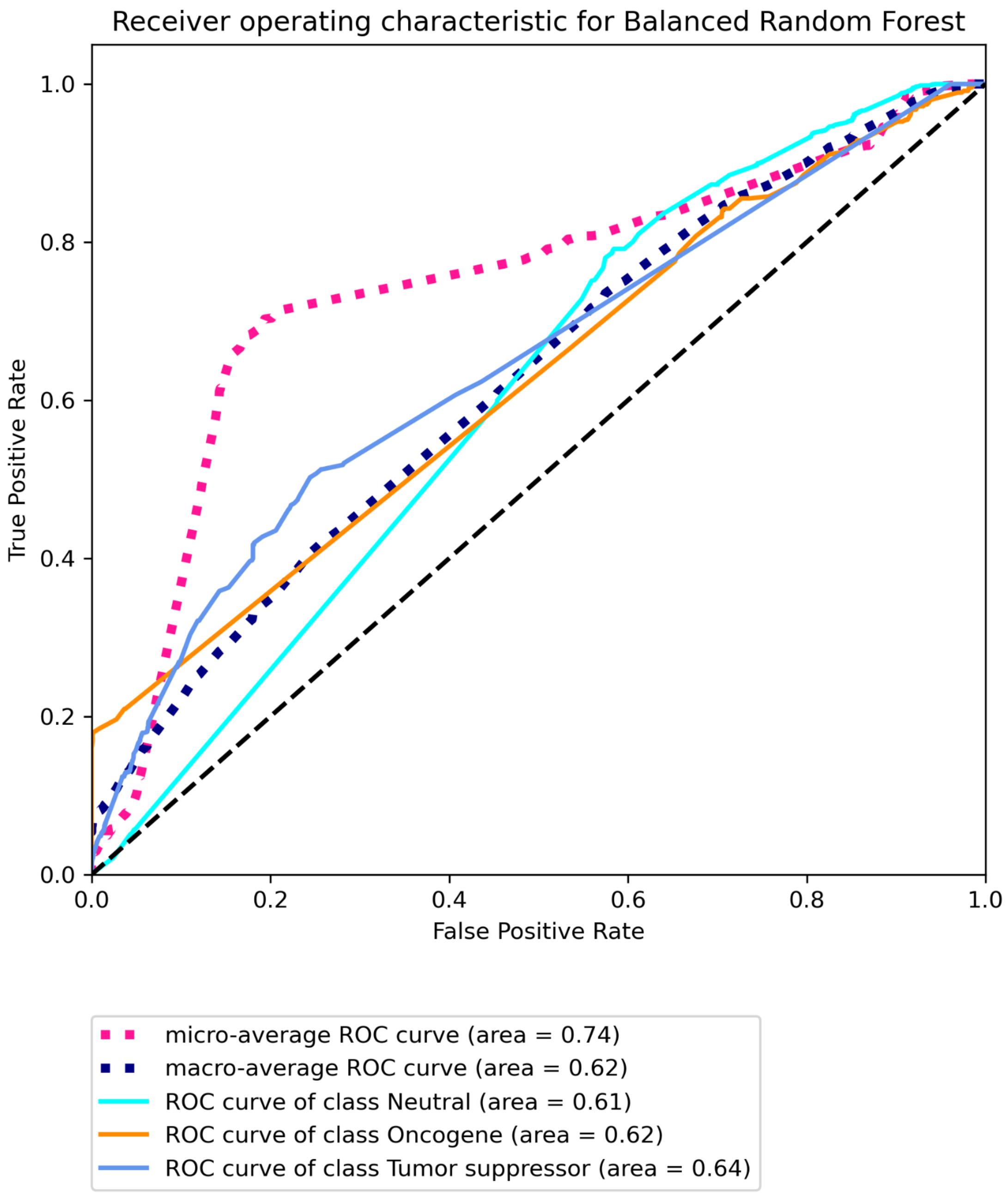


**Figure S12C Receiver operating characteristic for COAD using SNV data with CGC labels built on "some" features using Balanced Random Forest.**

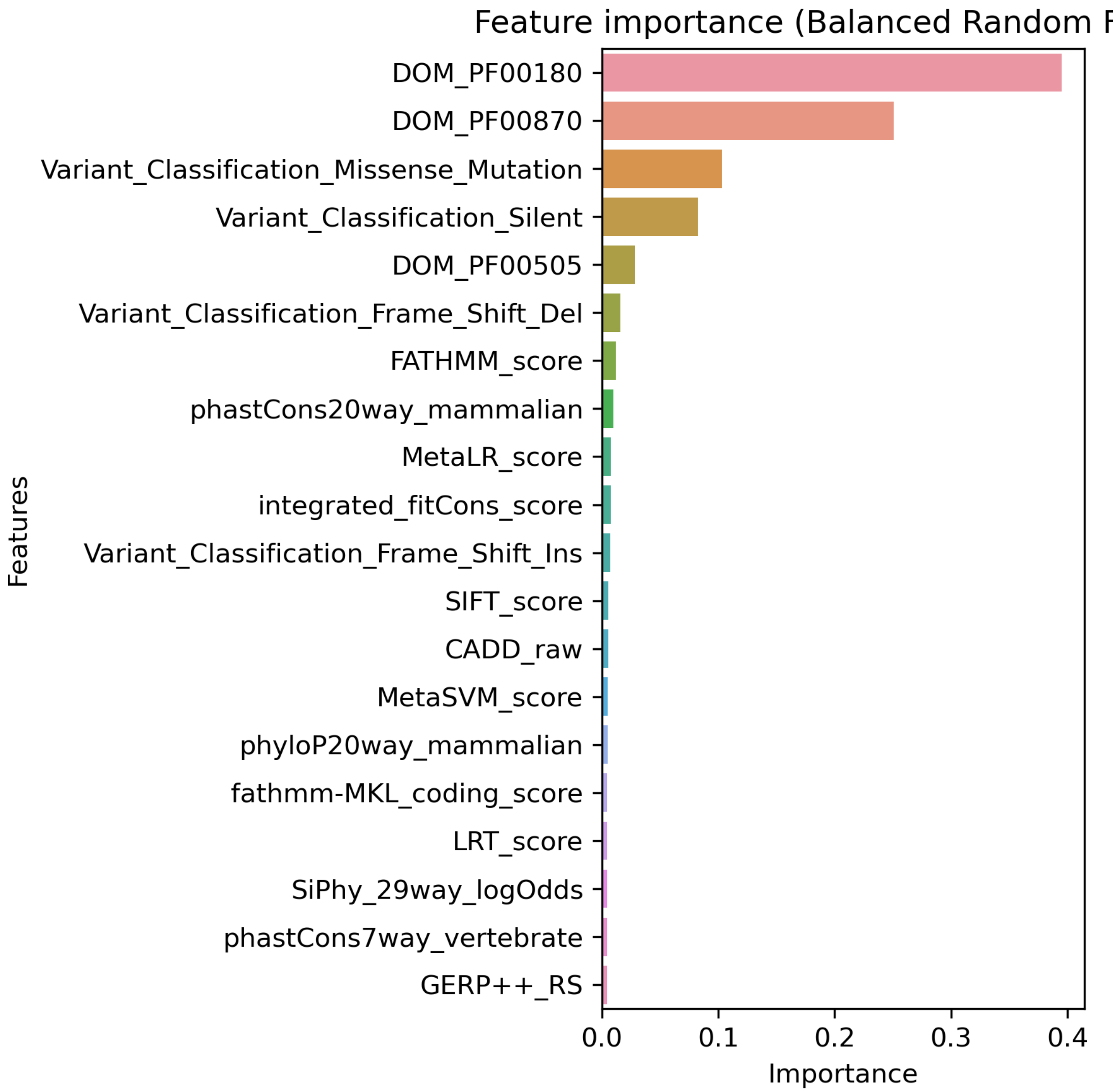


**Figure S13A. Top 20 features for LGG using SNV data with Bailey *et al.* labels built on "all" features using Balanced Random Forest.**

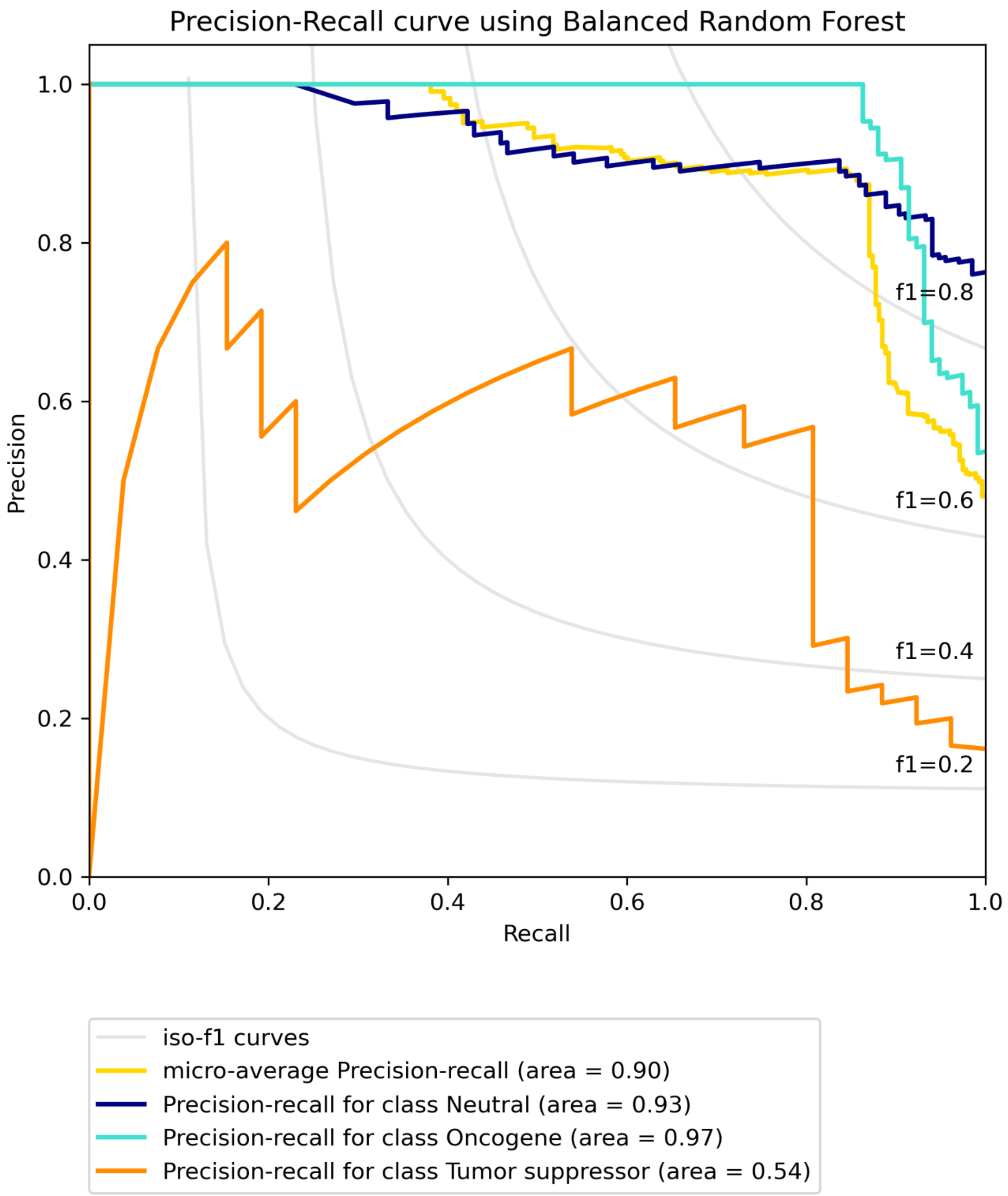


**Figure S13B. Precision-recall curve for LGG using SNV data with Bailey *et al.* labels built on "all" features using Balanced Random Forest.**

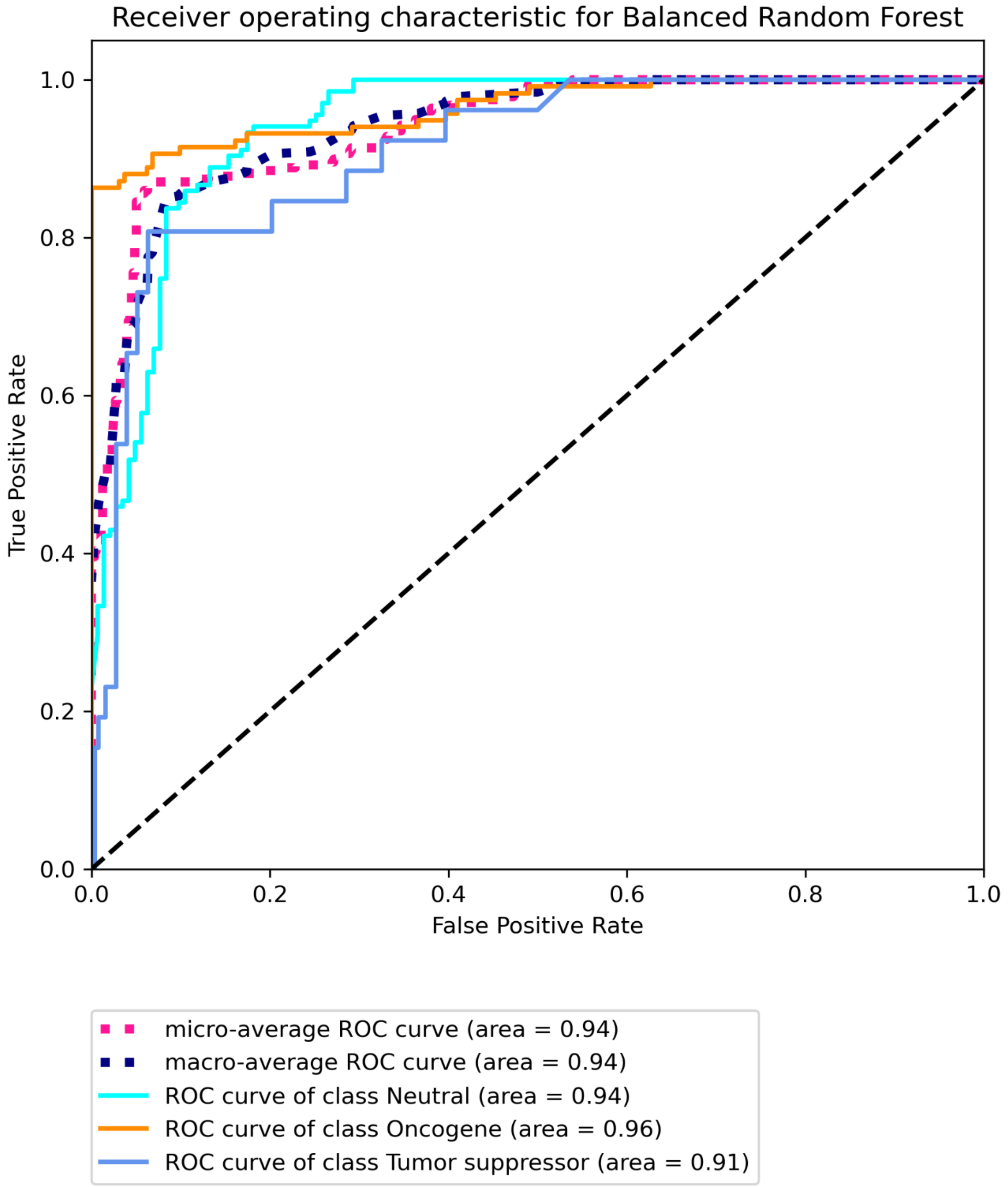


**Figure S13C Receiver operating characteristic for LGG using SNV data with Bailey *et al.* labels built on "all" features using Balanced Random Forest.**

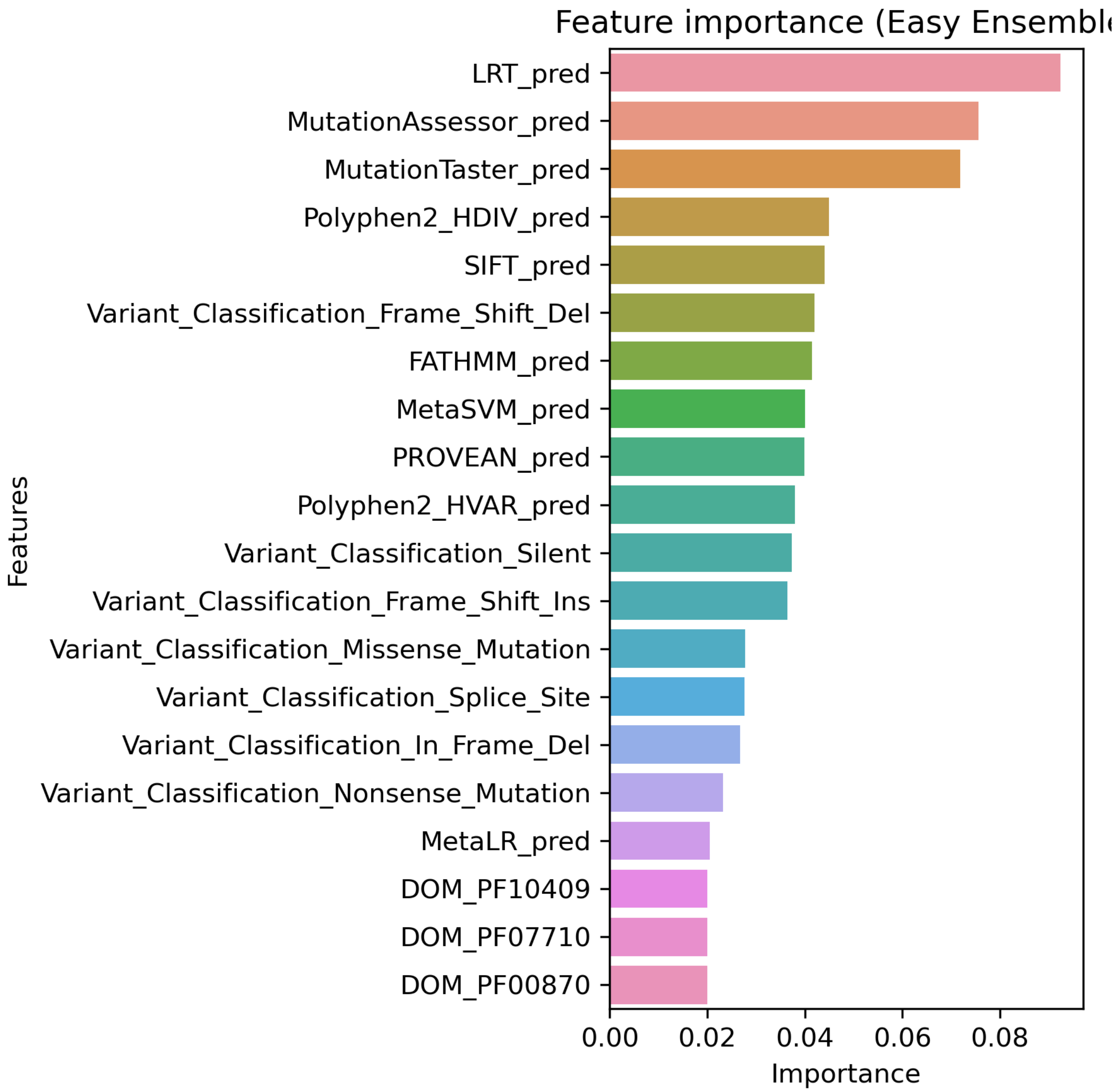


**Figure S14A. Top 20 features for LGG using SNV data with Bailey *et al.* labels built on "some" features using Easy Ensemble.**

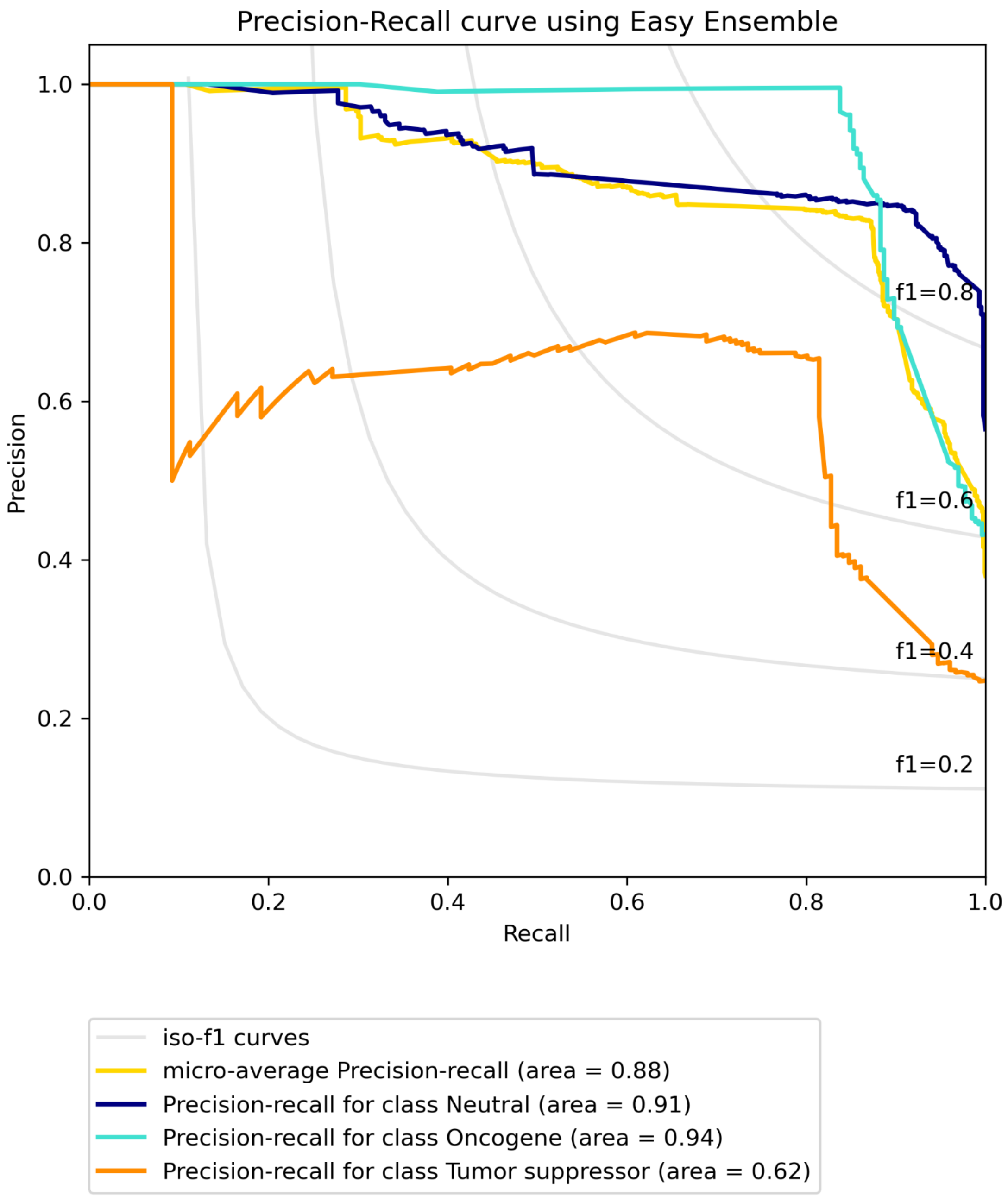


**Figure S14B. Precision-recall curve for LGG using SNV data with Bailey *et al.* labels built on "some" features using Easy Ensemble.**

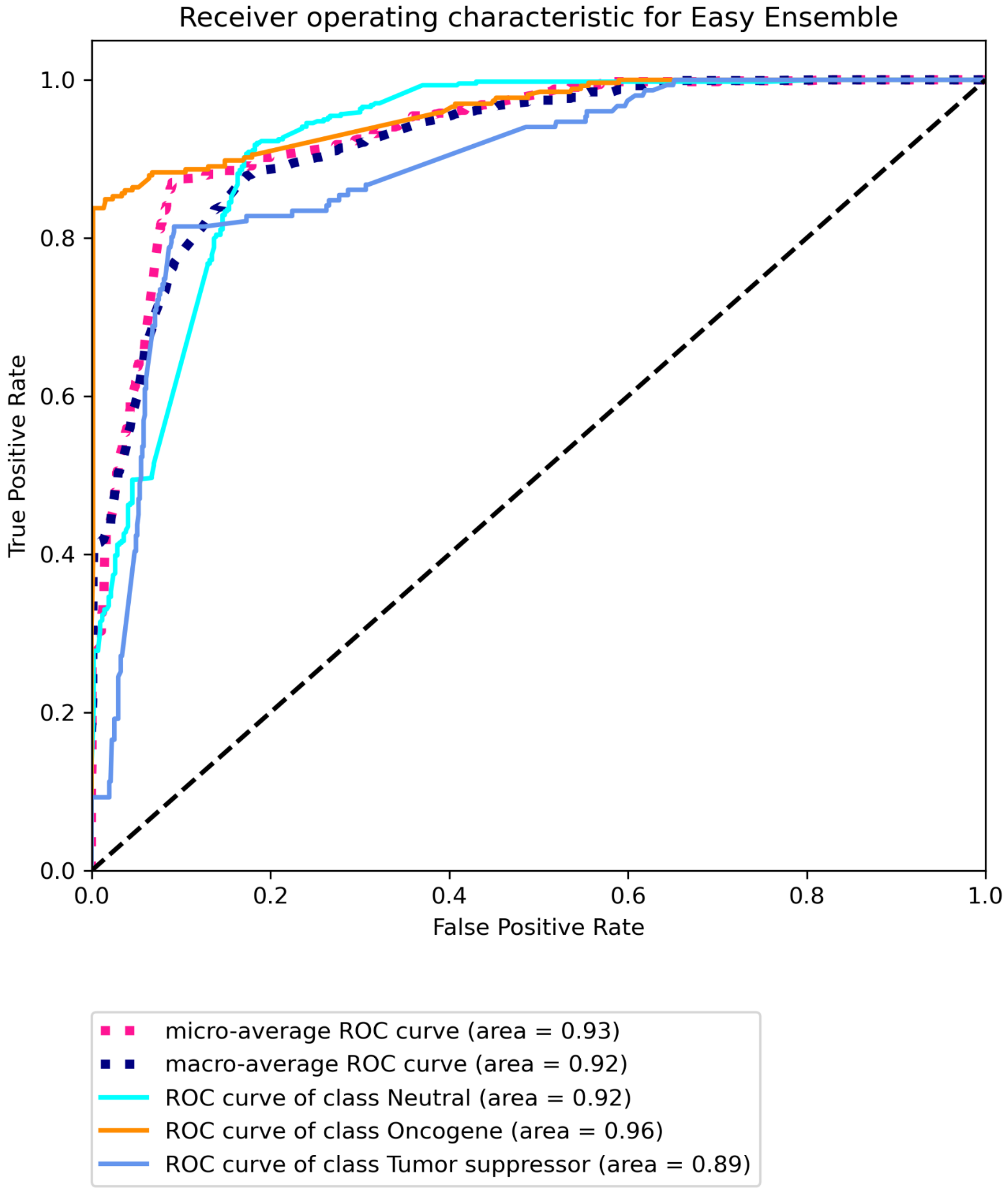


**Figure S14C Receiver operating characteristic for LGG using SNV data with Bailey *et al.* labels built on “some” features using Easy Ensemble.**

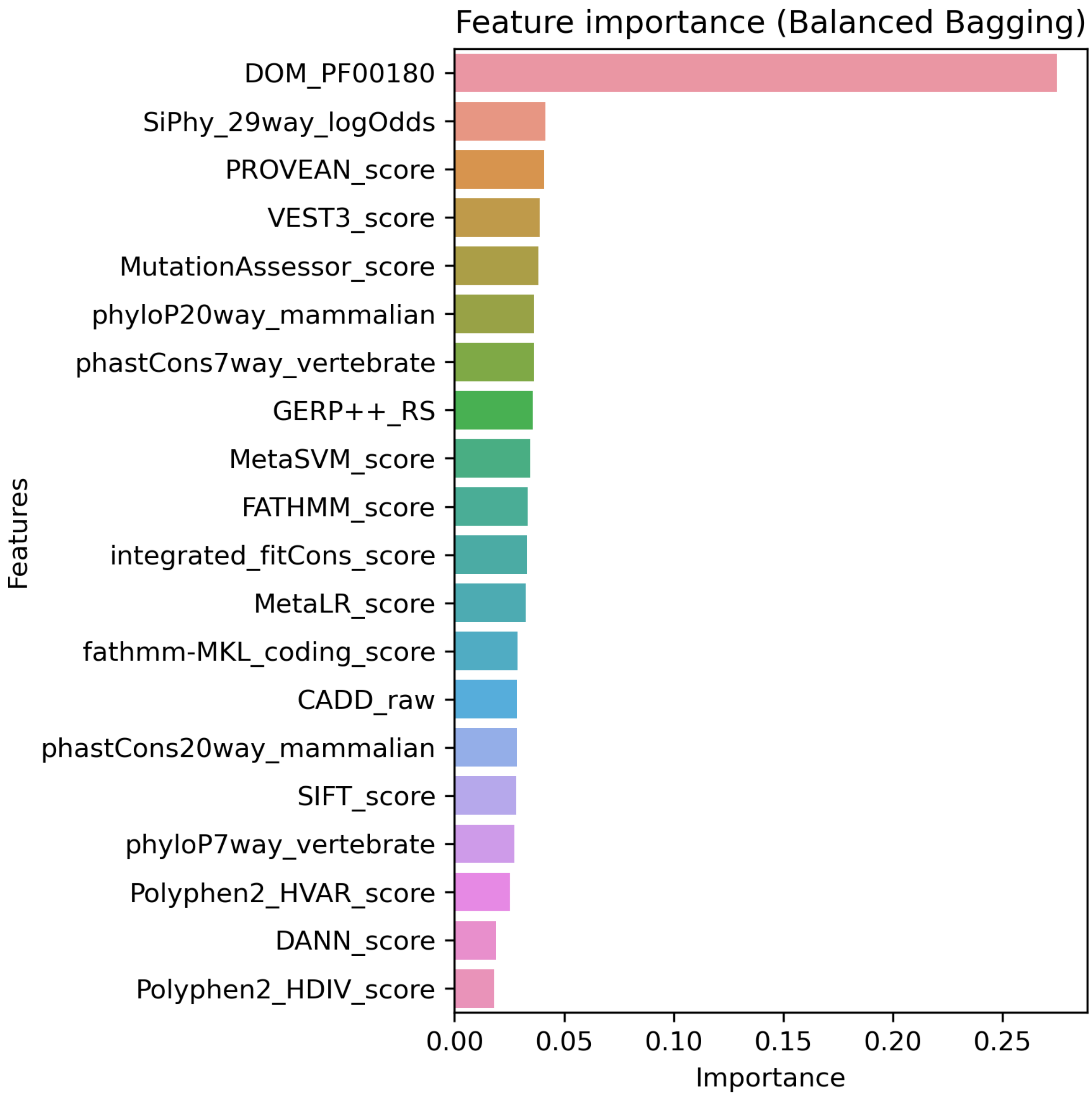


**Figure S15A. Top 20 features for LGG using SNV data with CGC labels built on “all” features using Balanced Bagging.**

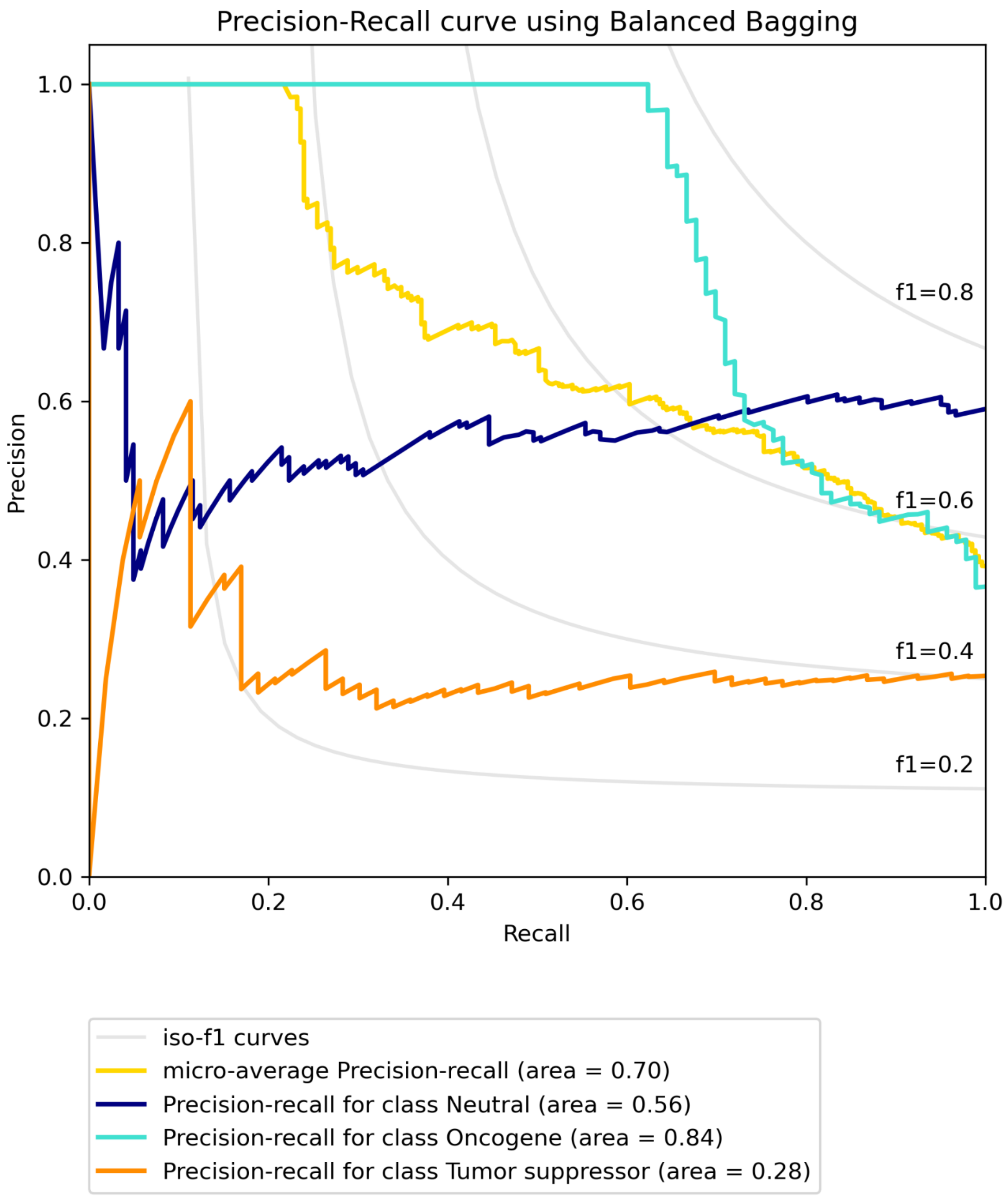


**Figure S15B. Precision-recall curve for LGG using SNV data with CGC labels built on "all" features using Balanced Bagging.**

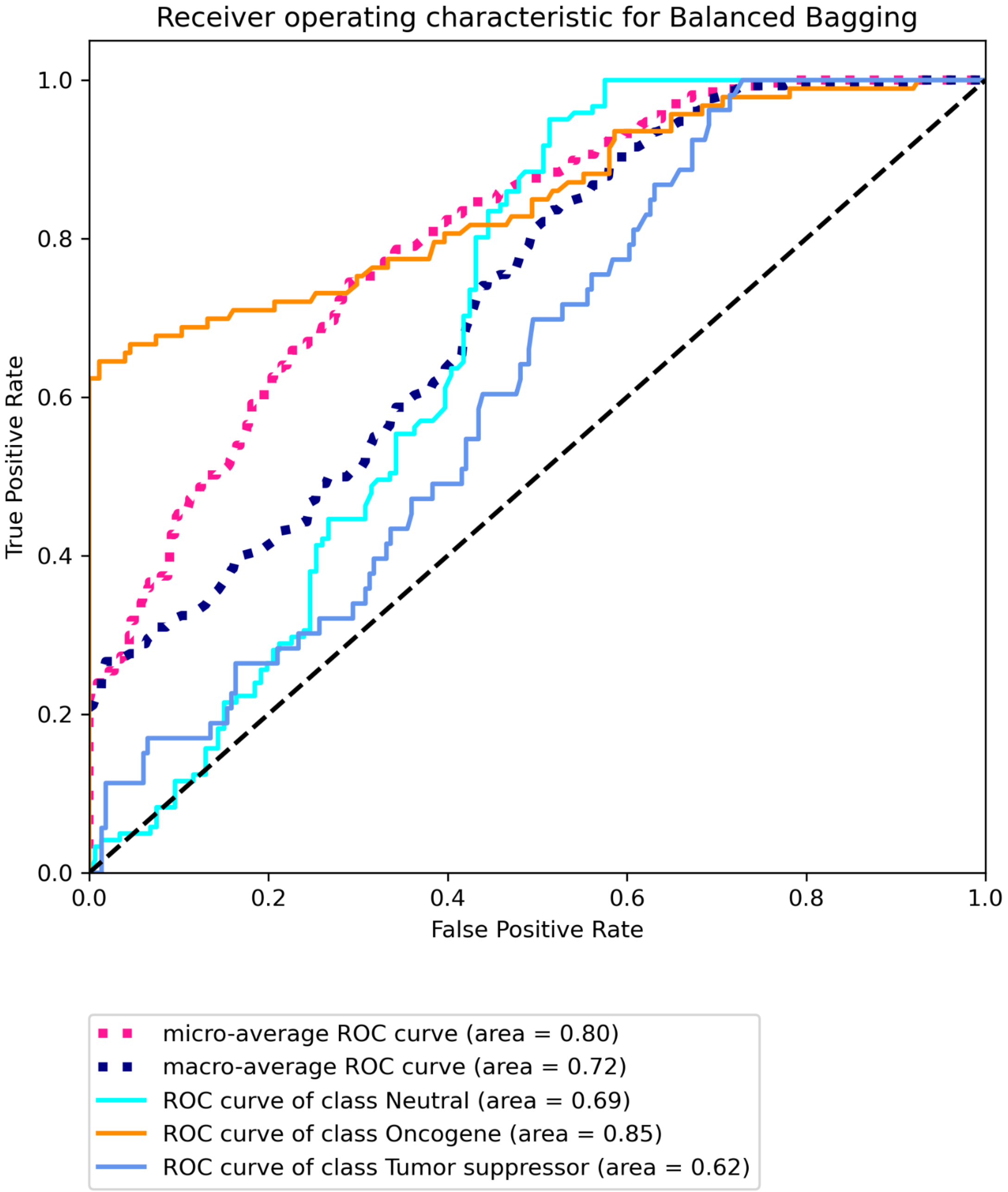


**Figure S15C Receiver operating characteristic for LGG using SNV data with CGC labels built on “all” features using Balanced Bagging.**

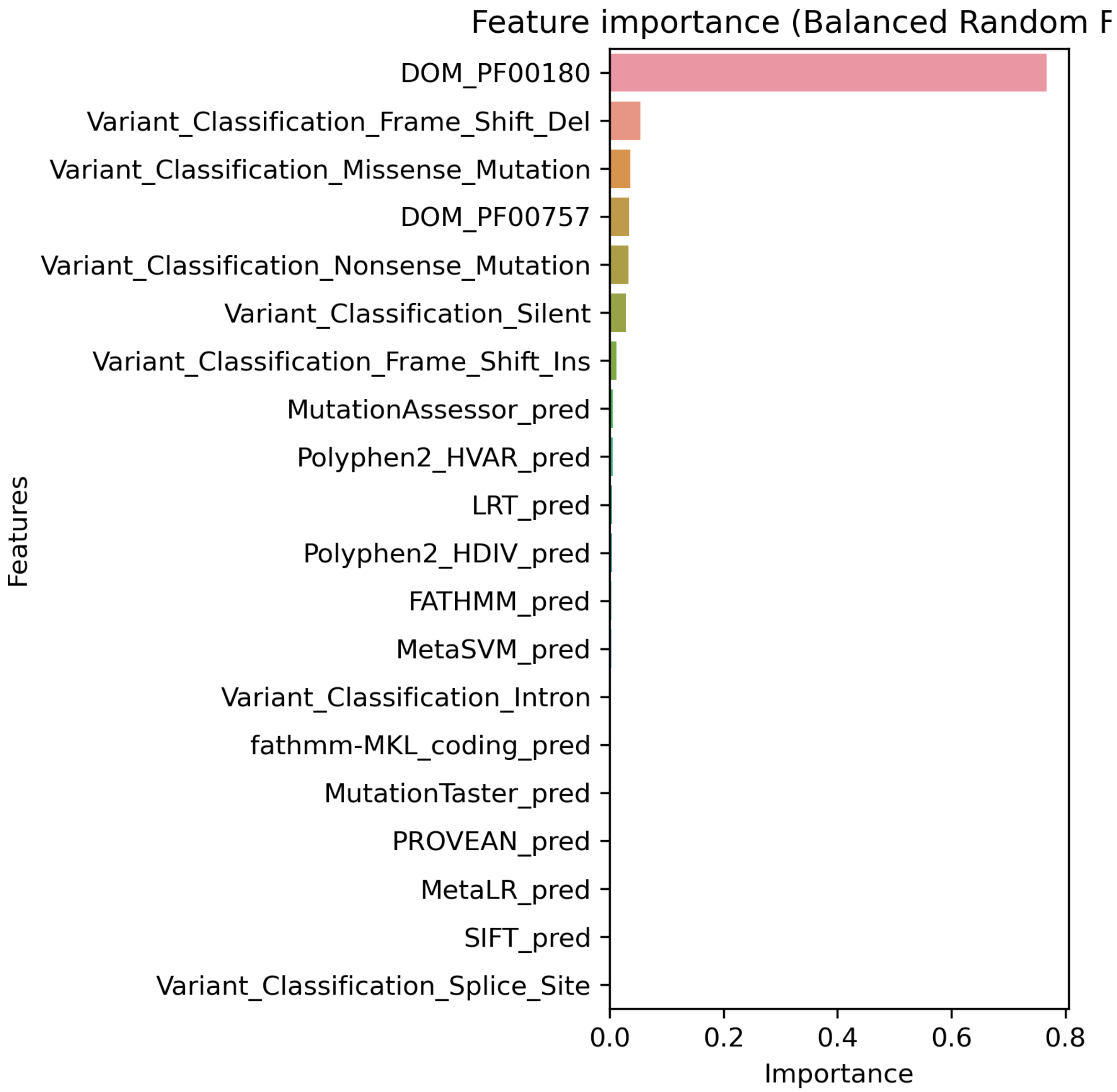


**Figure S16A. Top 20 features for LGG using SNV data with CGC labels built on "some" features using Balanced Random Forest.**

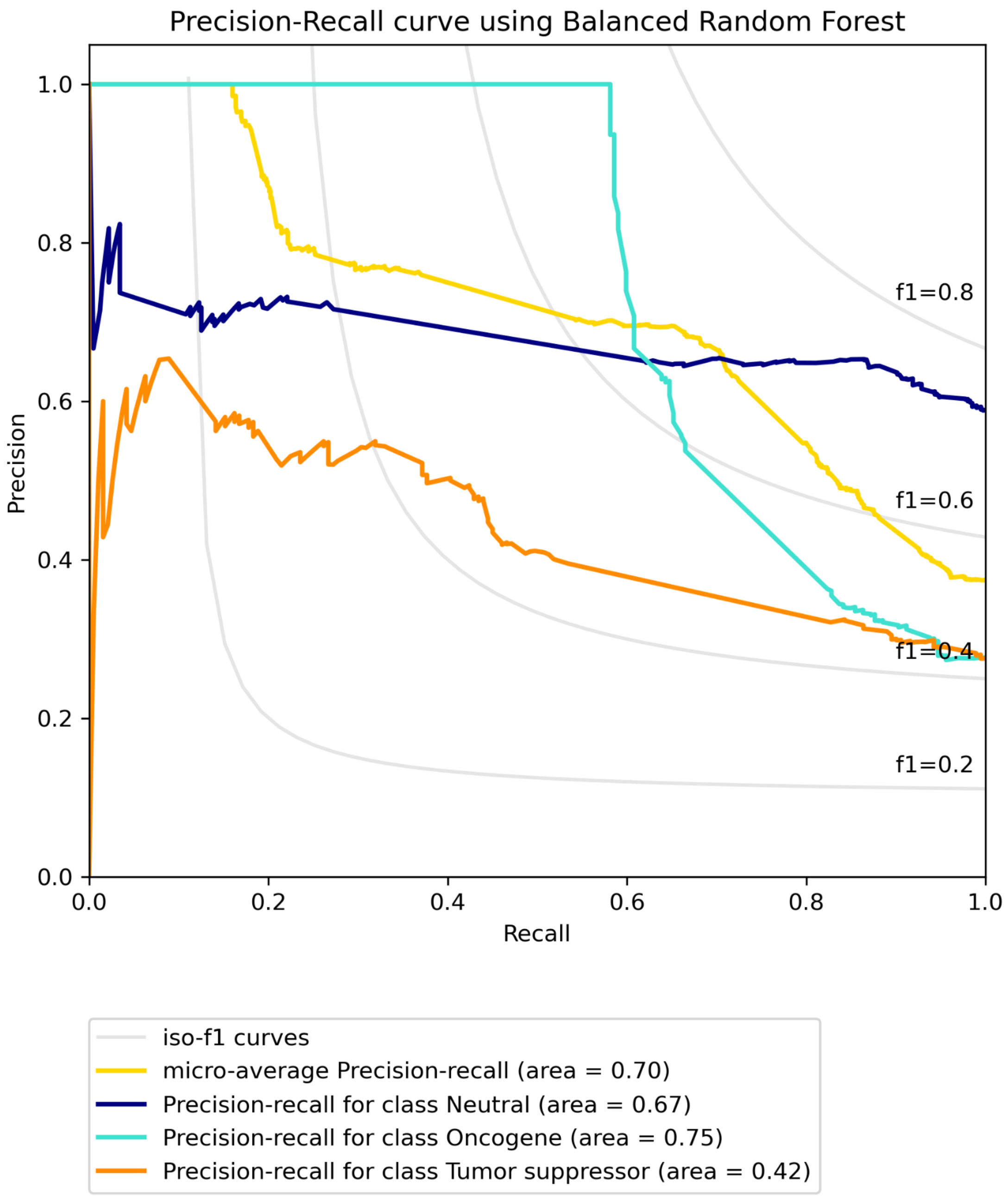


**Figure S16B. Precision-recall curve for LGG using SNV data with CGC labels built on “some” features using Balanced Random Forest.**

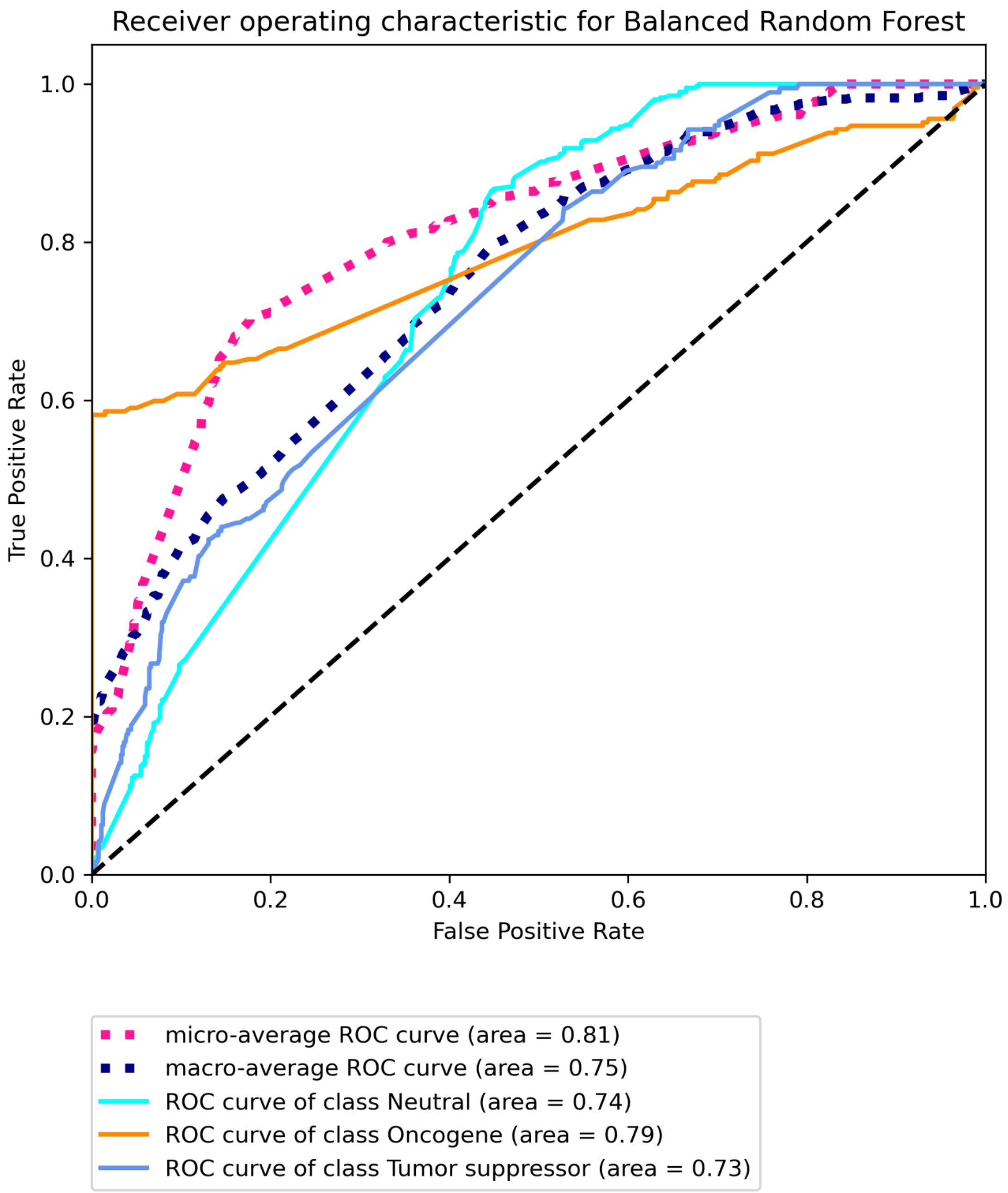


**Figure S16C Receiver operating characteristic for LGG using SNV data with CGC labels built on “some” features using Balanced Random Forest.**

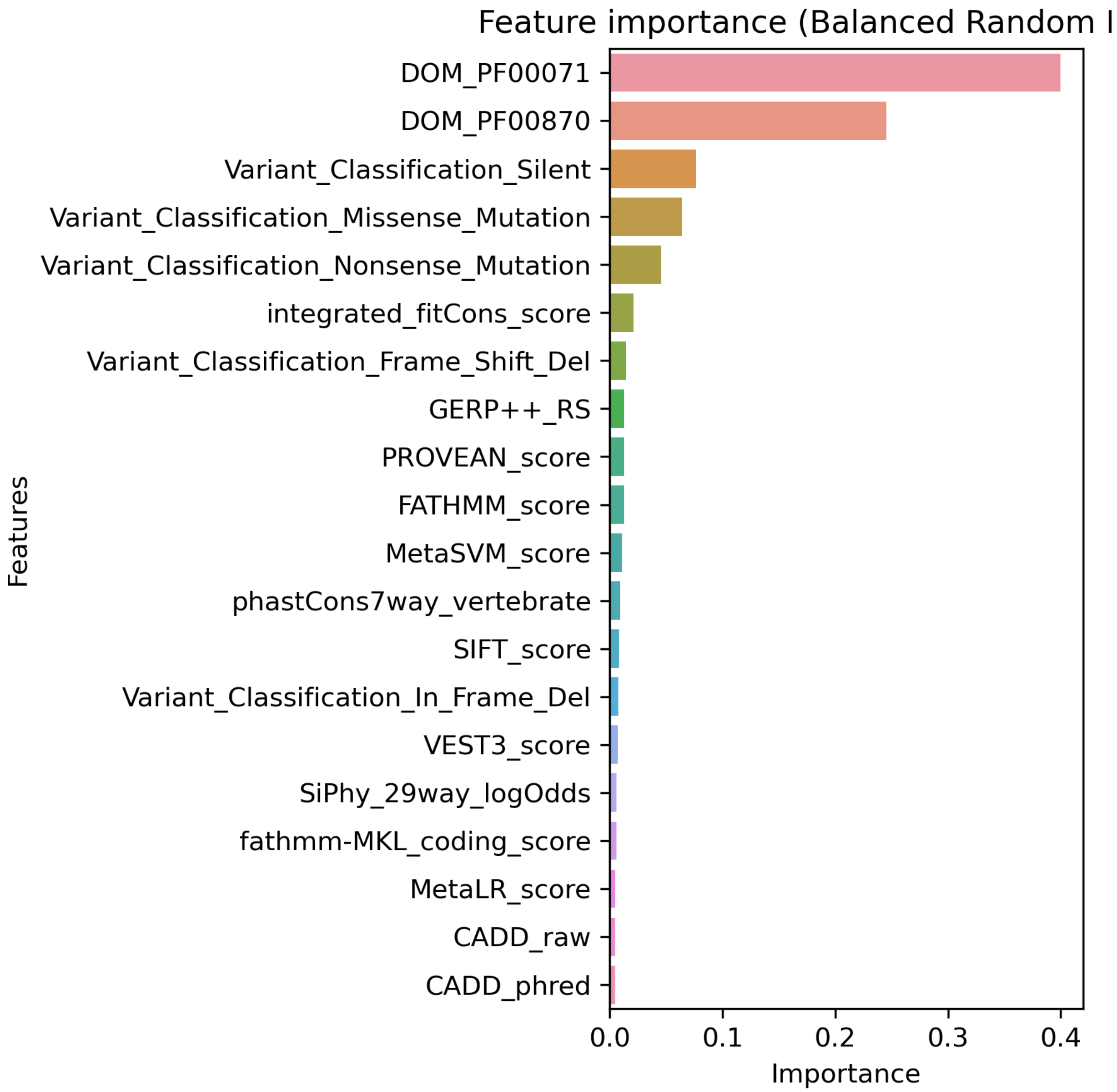


**Figure S17A. Top 20 features for LUAD using SNV data with Bailey *et al.* labels built on "all" features using Balanced Random Forest.**

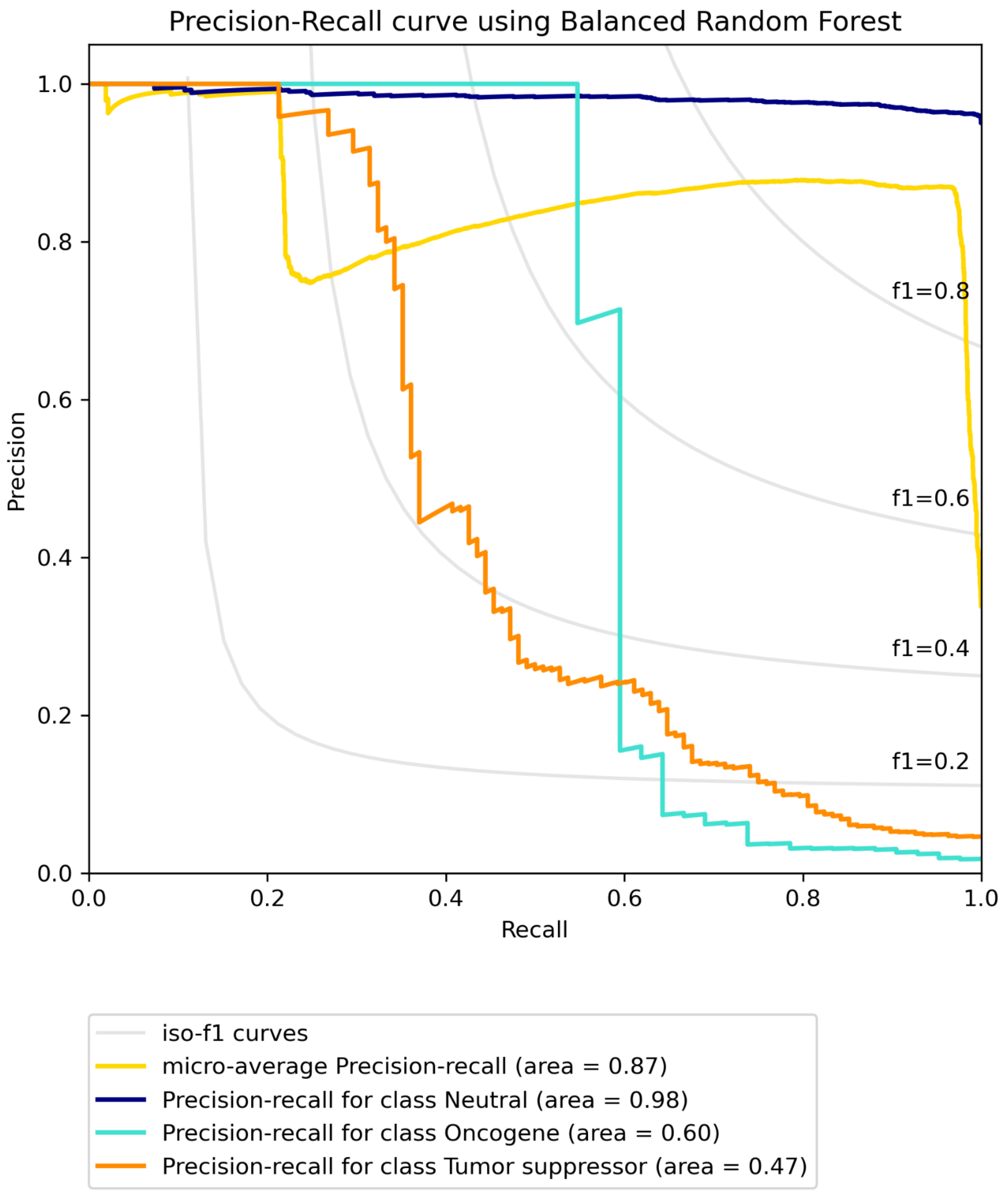


**Figure S17B. Precision-recall curve for LUAD using SNV data with Bailey *et al.* labels built on "all" features using Balanced Random Forest.**

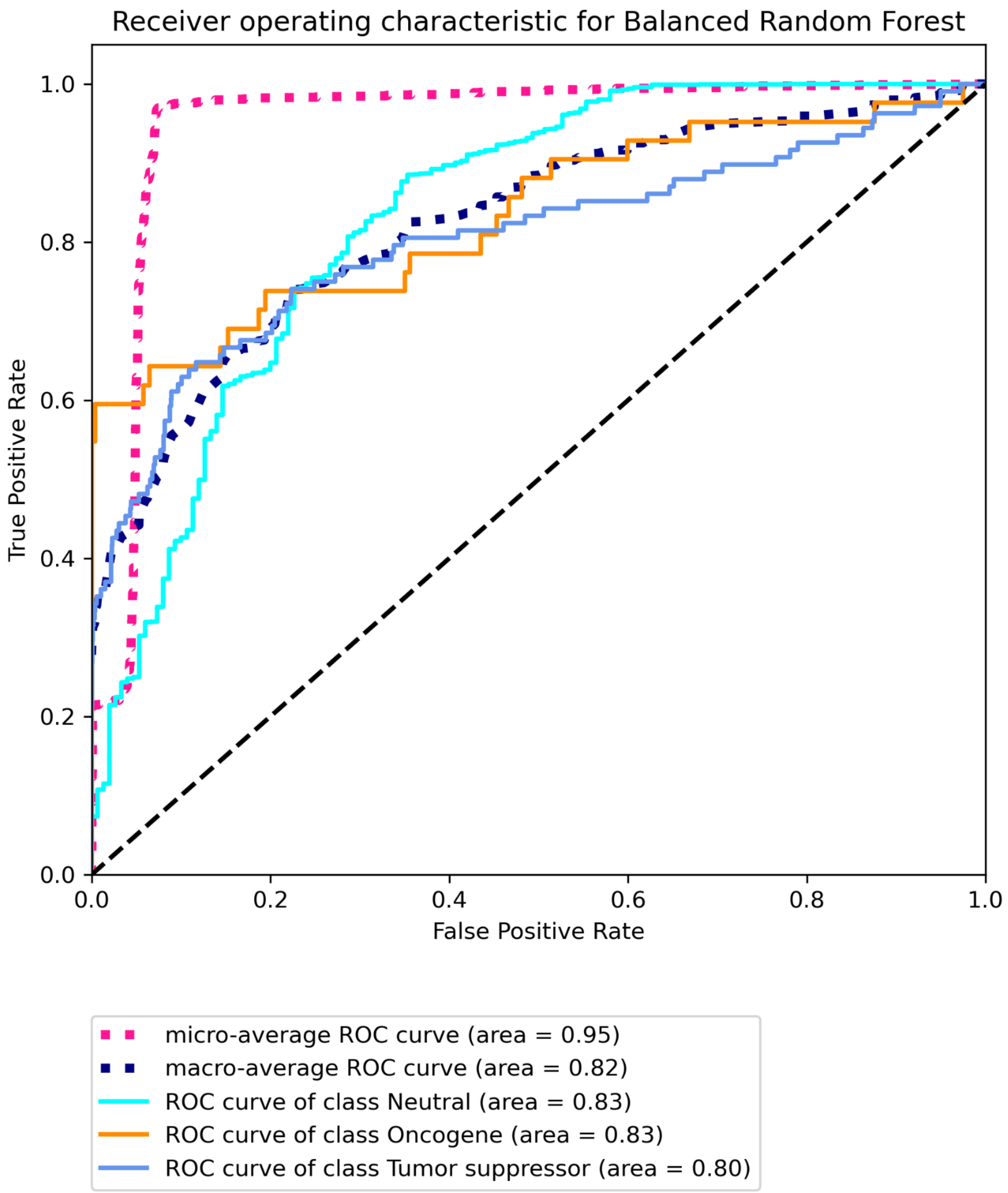


**Figure S17C Receiver operating characteristic for LUAD using SNV data with Bailey *et al.* labels built on "all" features using Balanced Random Forest.**

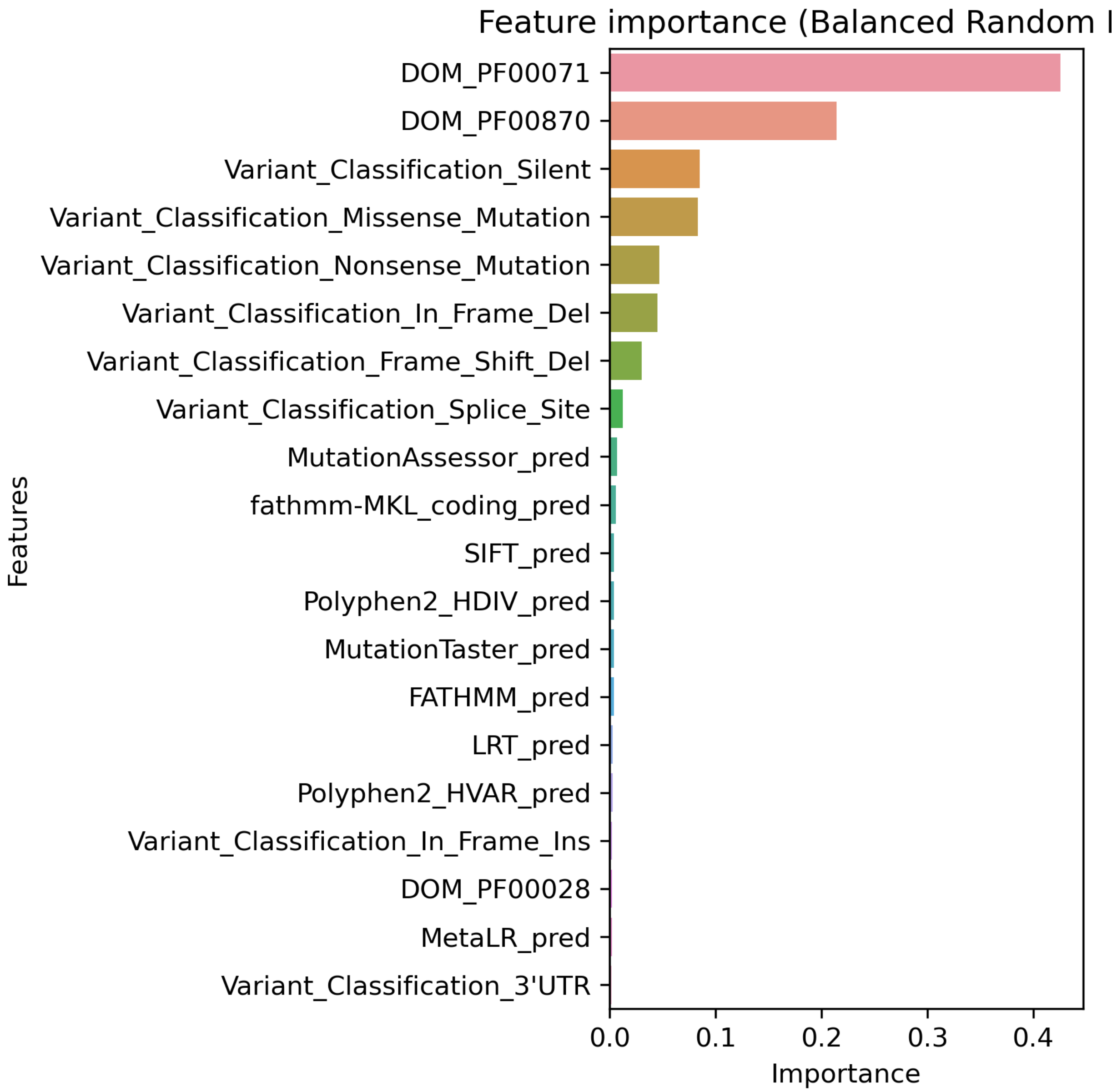


**Figure S18A. Top 20 features for LUAD using SNV data with Bailey *et al.* labels built on "some" features using Balanced Random Forest.**

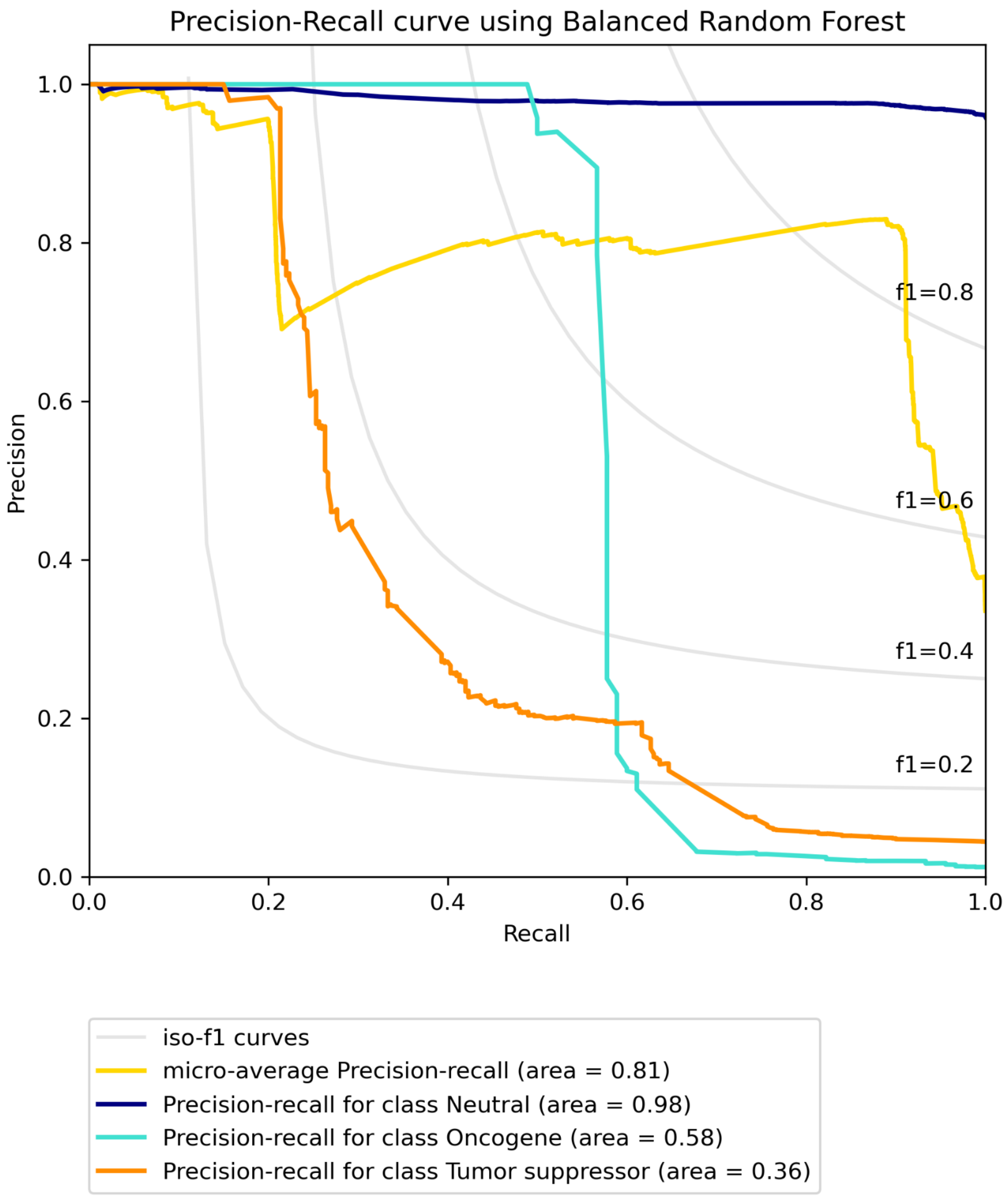


**Figure S18B. Precision-recall curve for LUAD using SNV data with Bailey *et al.* labels built on "some" features using Balanced Random Forest.**

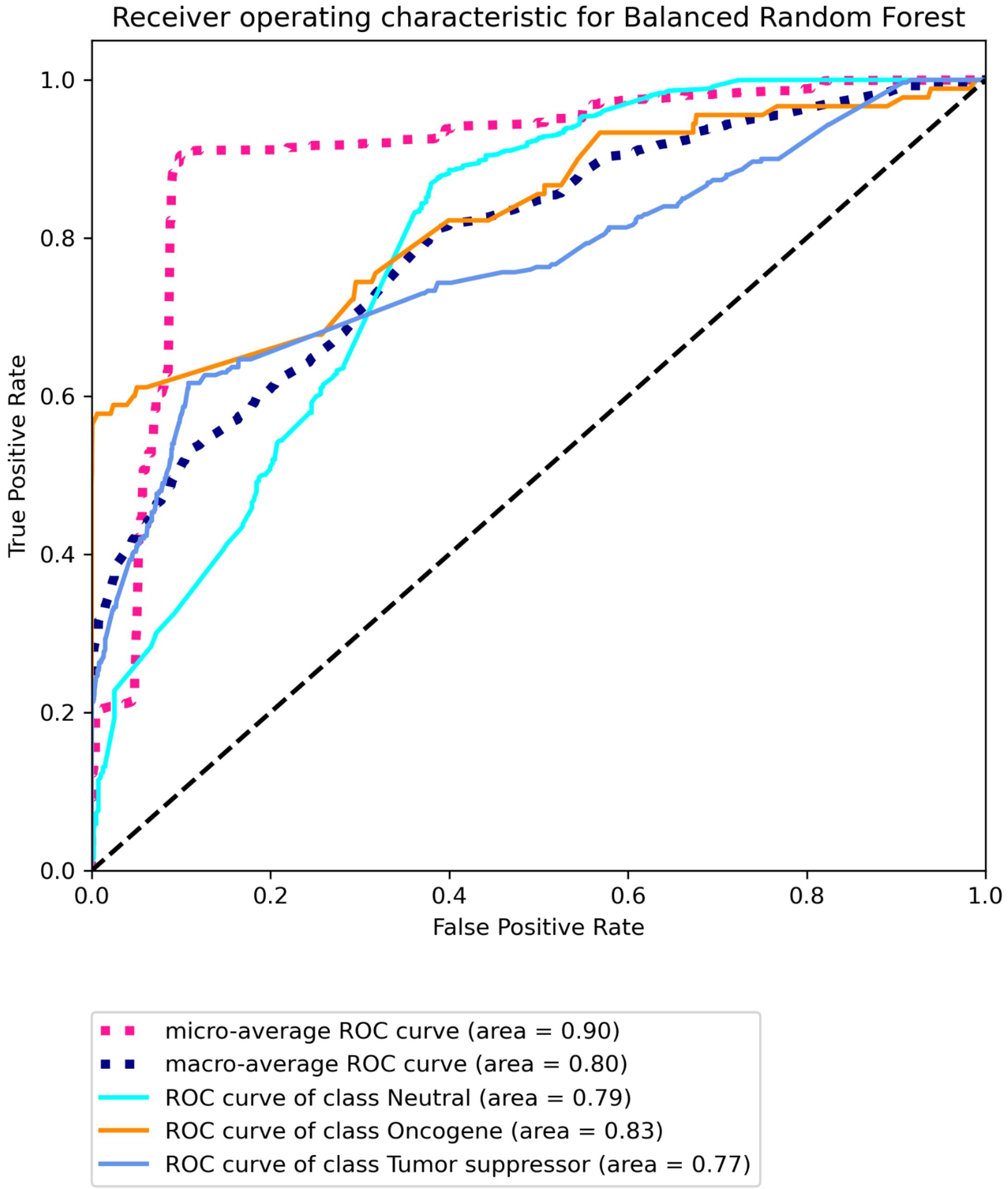


**Figure S18C Receiver operating characteristic for LUAD using SNV data with Bailey *et al.* labels built on "some" features using Balanced Random Forest.**

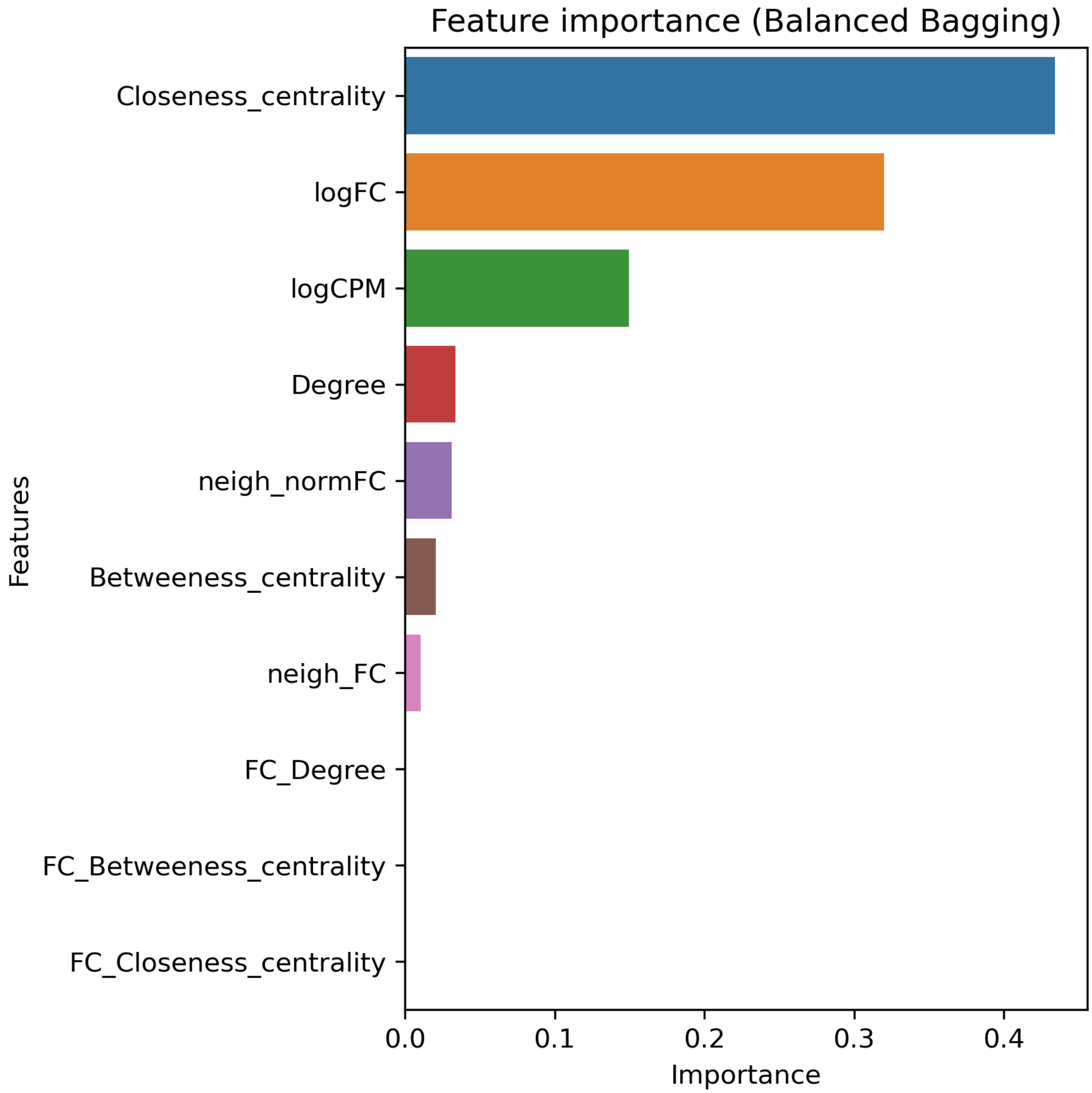


**Figure S19A. Top 20 features for BRCA using RNA data with Bailey *et al.* labels built using Balanced bagging.**

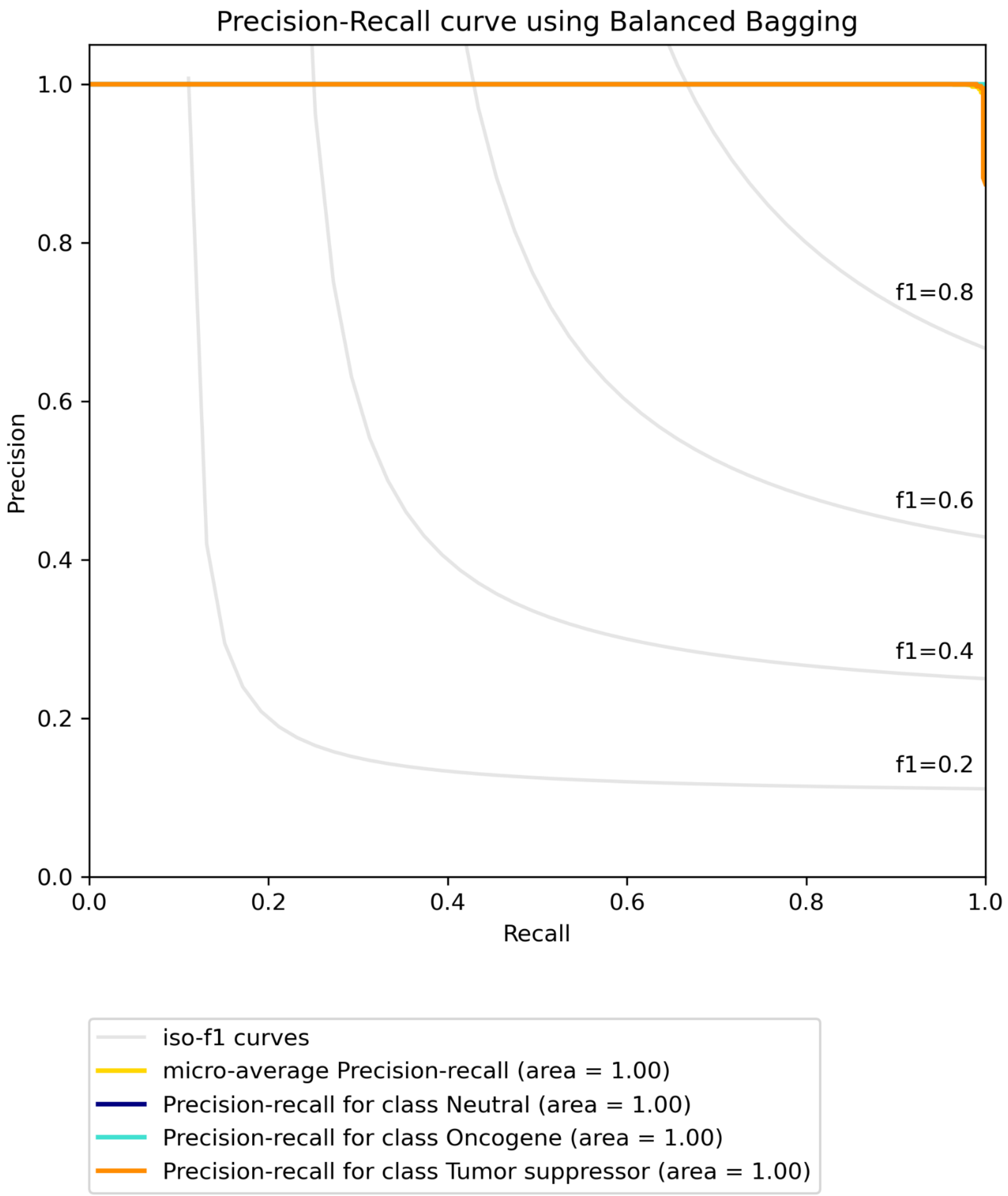


**Figure S19B. Precision-recall curve for BRCA using RNA data with Bailey *et al.* labels built using Balanced bagging.**

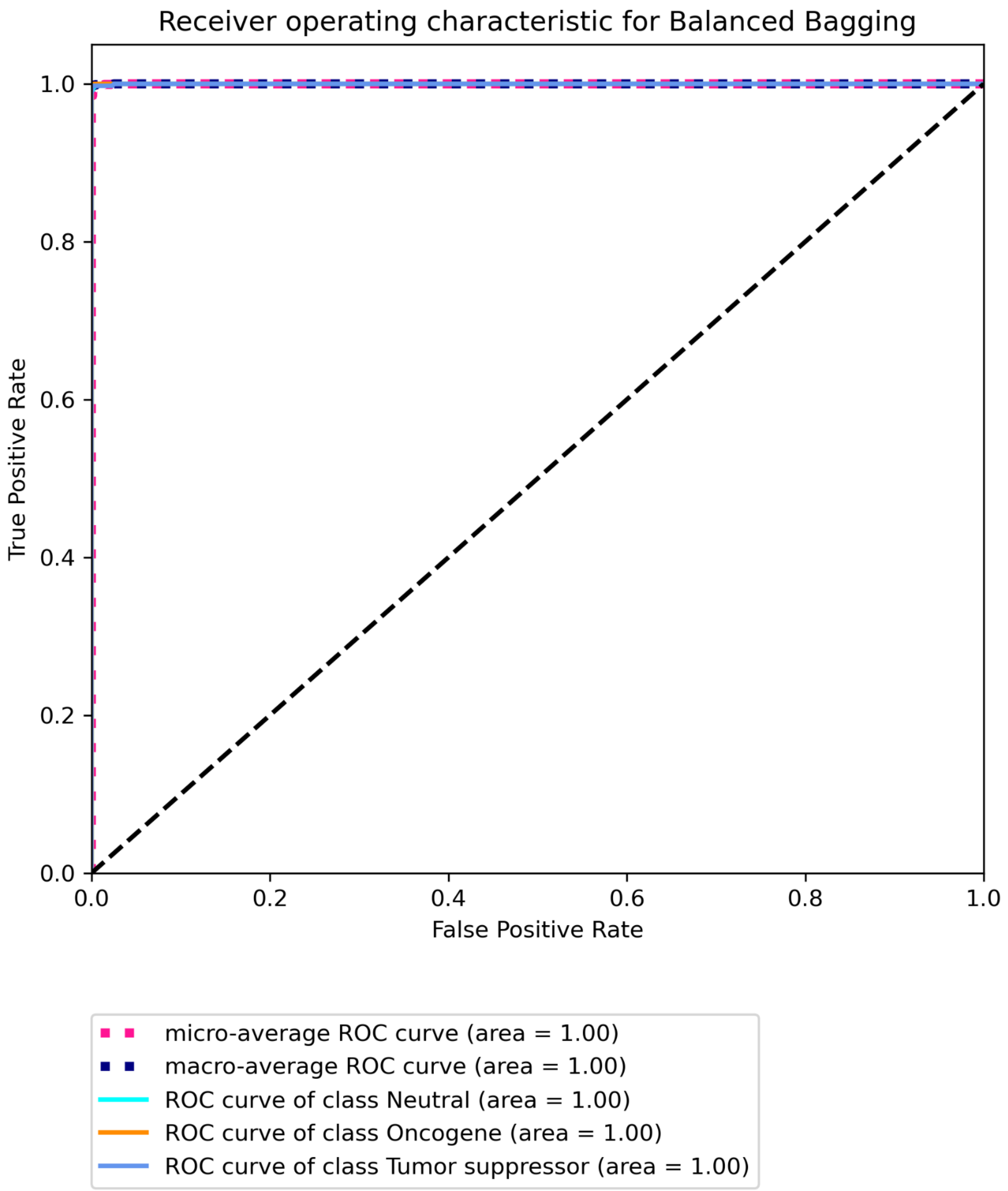


**Figure S19C Receiver operating characteristic for BRCA using RNA data with Bailey *et al.* labels built using Balanced bagging.**

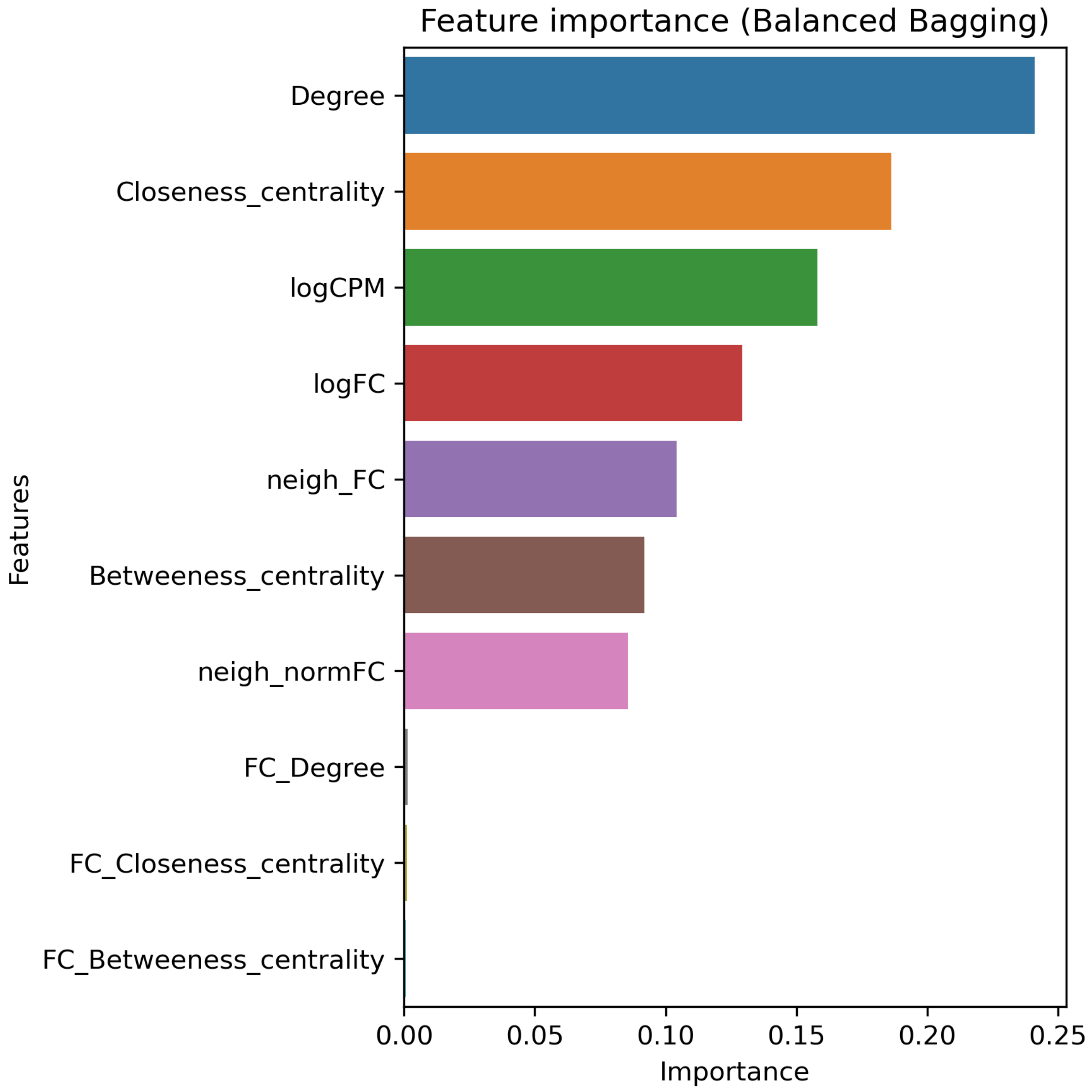


**Figure S20A. Top 20 features for BRCA using RNA data with CGC labels built using Balanced bagging.**

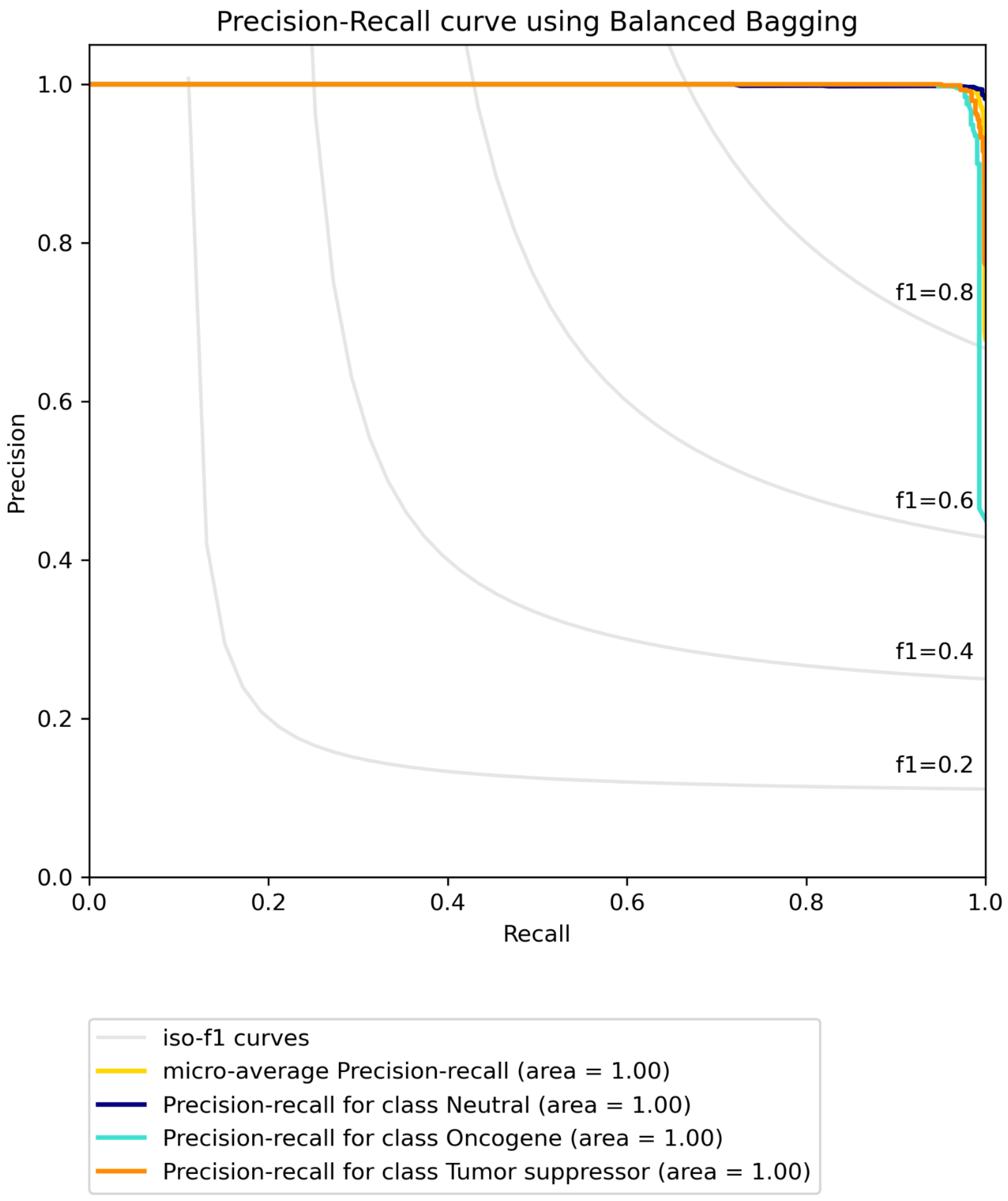


**Figure S20B. Precision-recall curve for BRCA using RNA data with CGC labels built using Balanced bagging.**

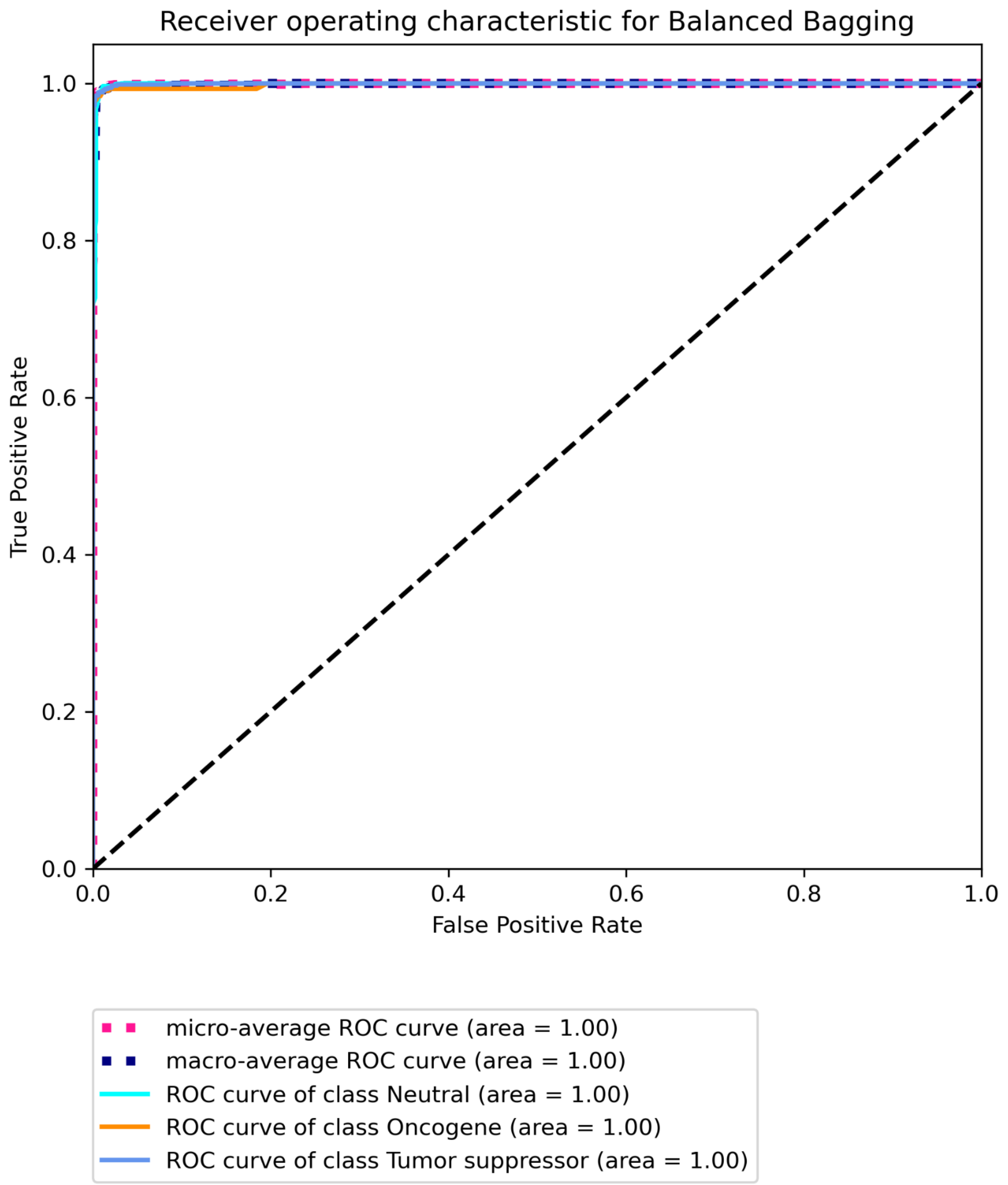


**Figure S20C Receiver operating characteristic for BRCA using RNA data with CGC labels built using Balanced bagging.**

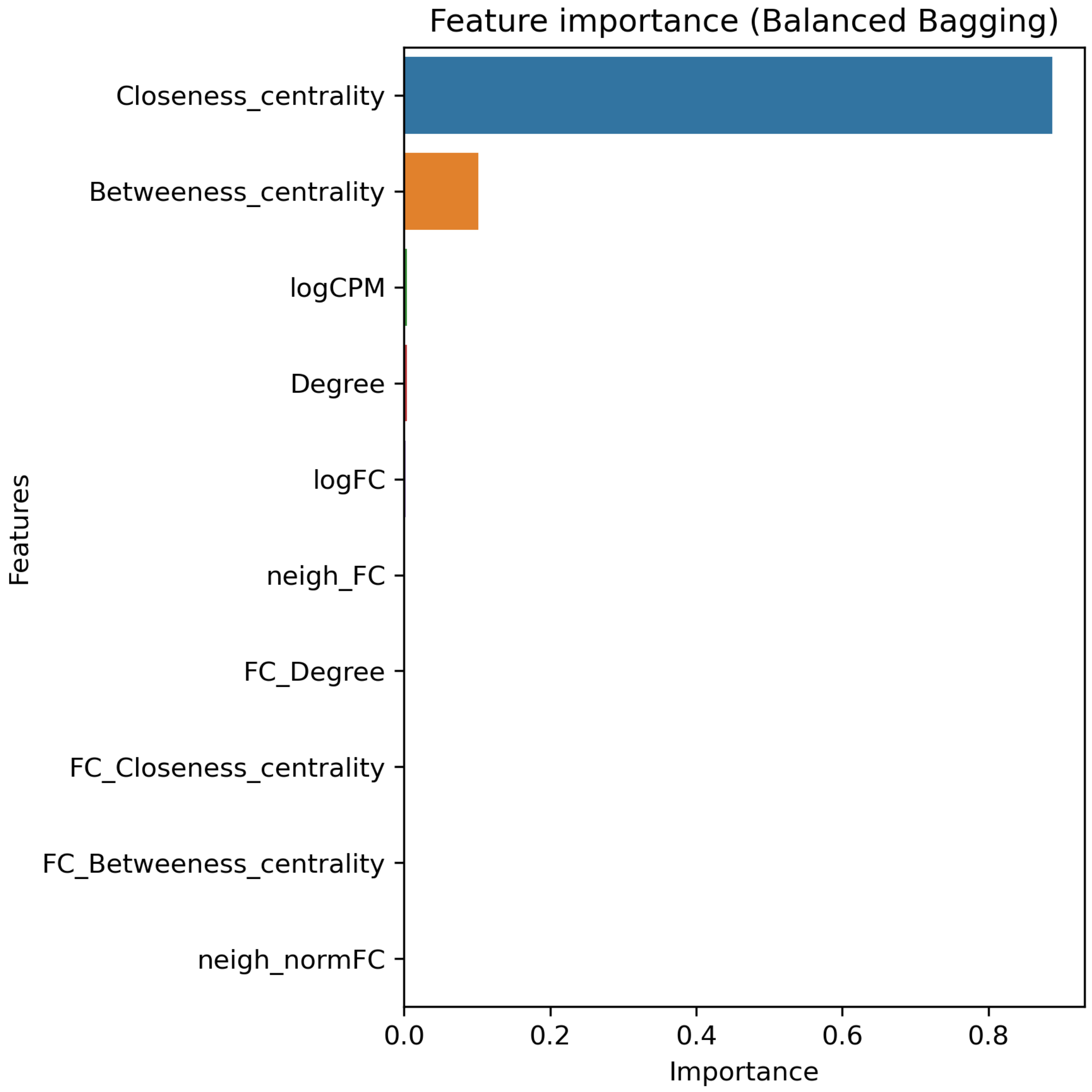


**Figure S21A. Top 20 features for BRCA using RNA data with CIViC labels built using Balanced bagging.**

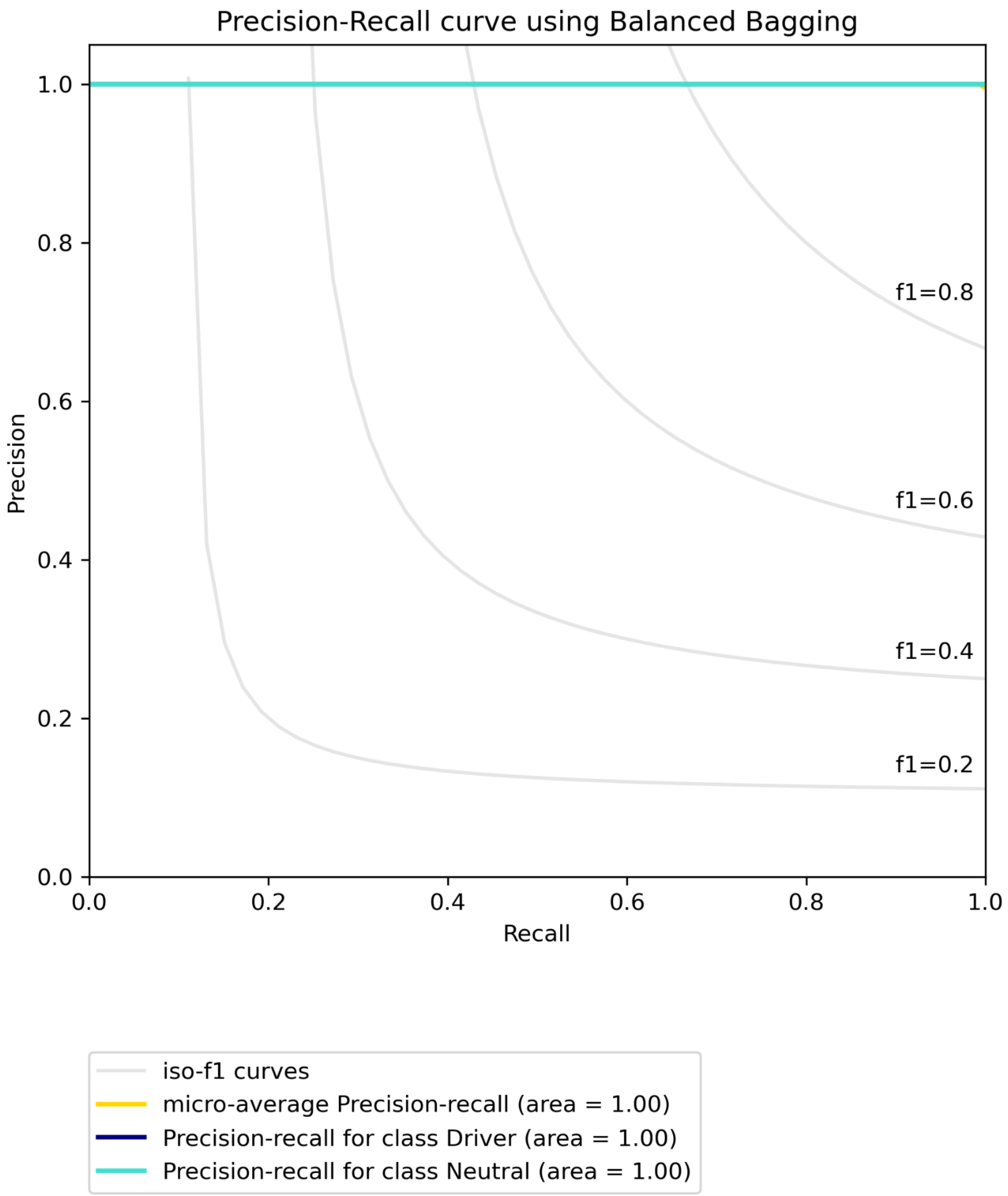


**Figure S21B. Precision-recall curve for BRCA using RNA data with CIViC labels built using Balanced bagging.**

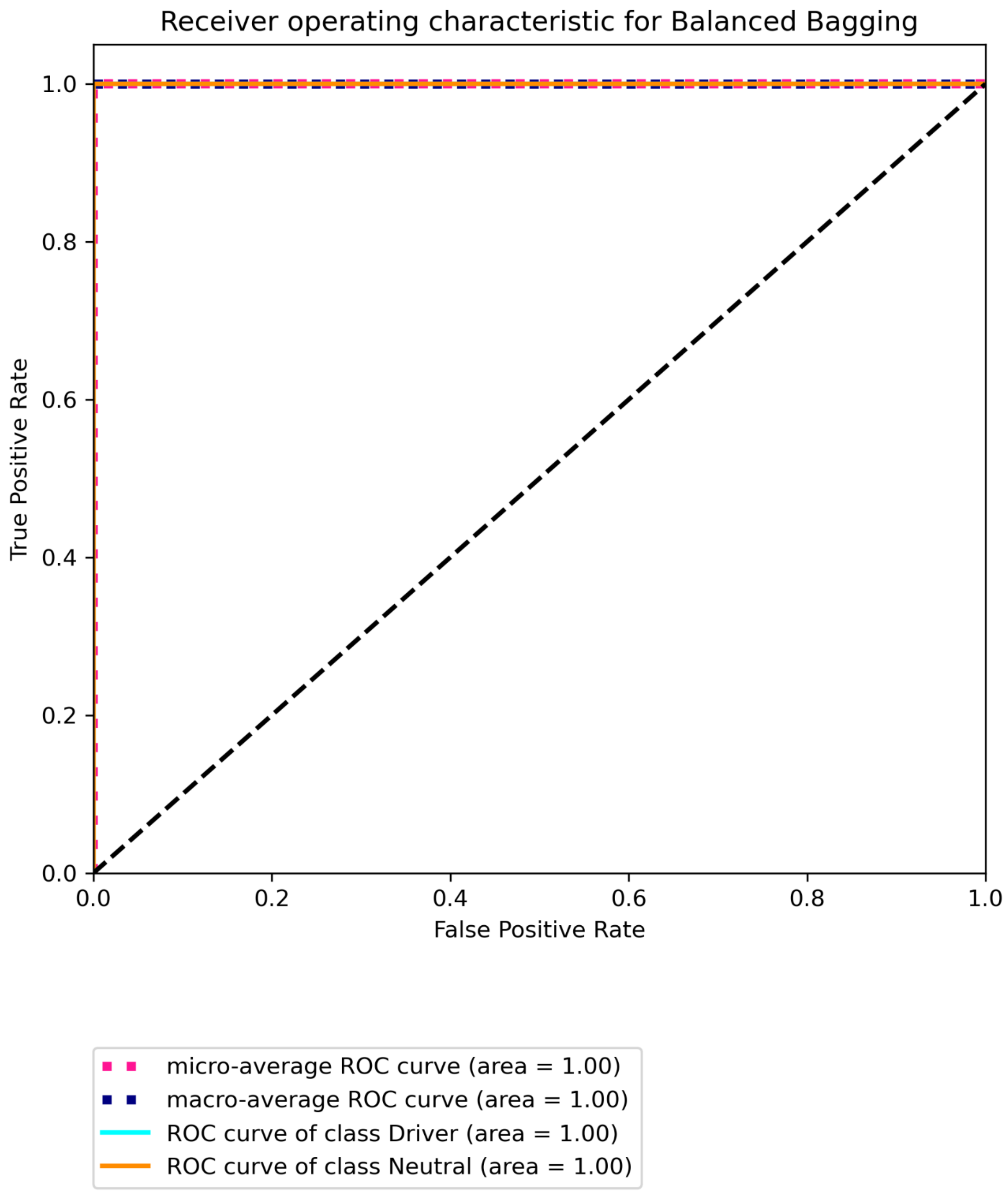


**Figure S21C Receiver operating characteristic for BRCA using RNA data with CIViC labels built using Balanced bagging.**

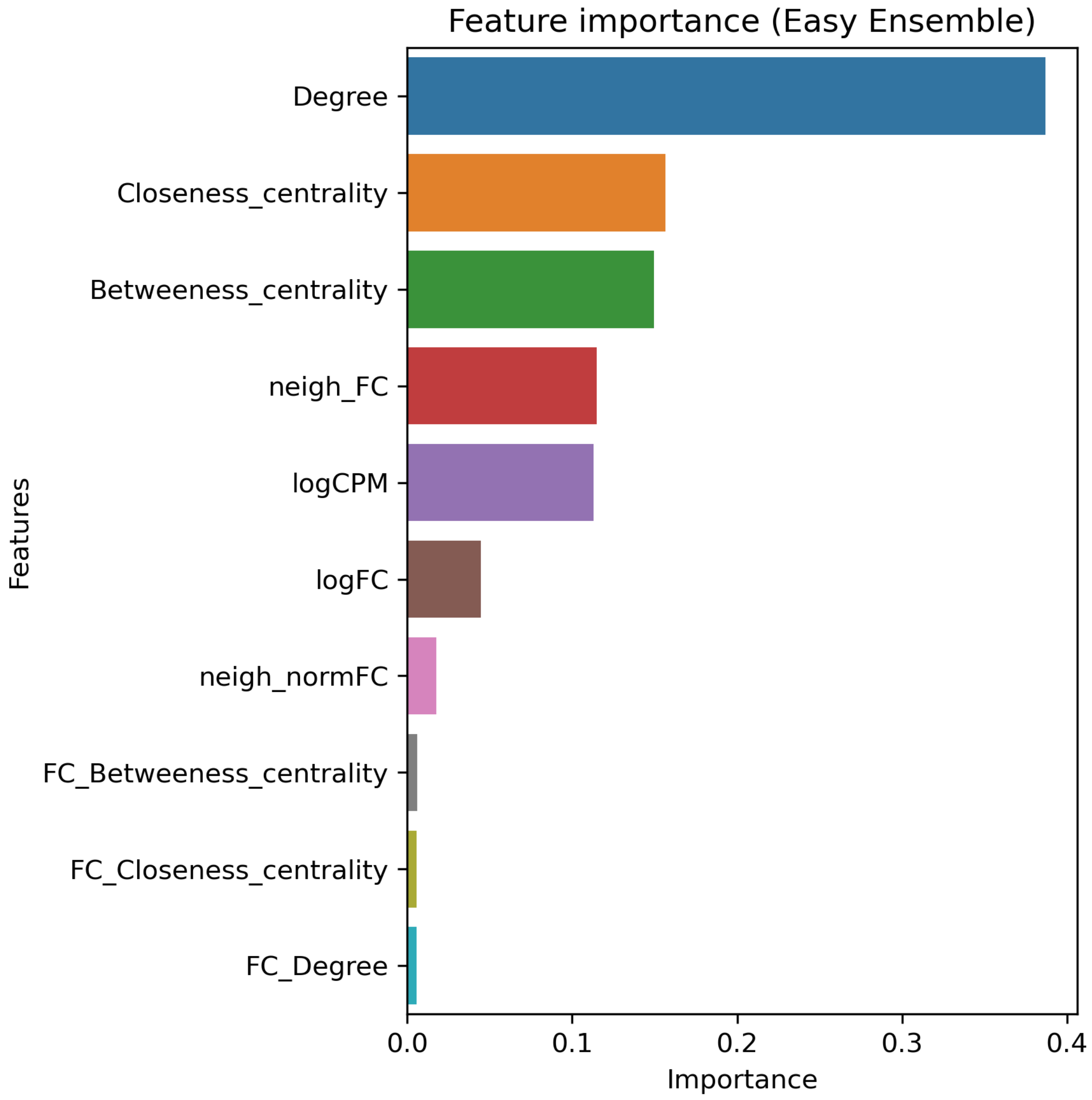


**Figure S22A. Top 20 features for BRCA using RNA data with Marellotto *et al.* labels built using Easy Ensemble.**

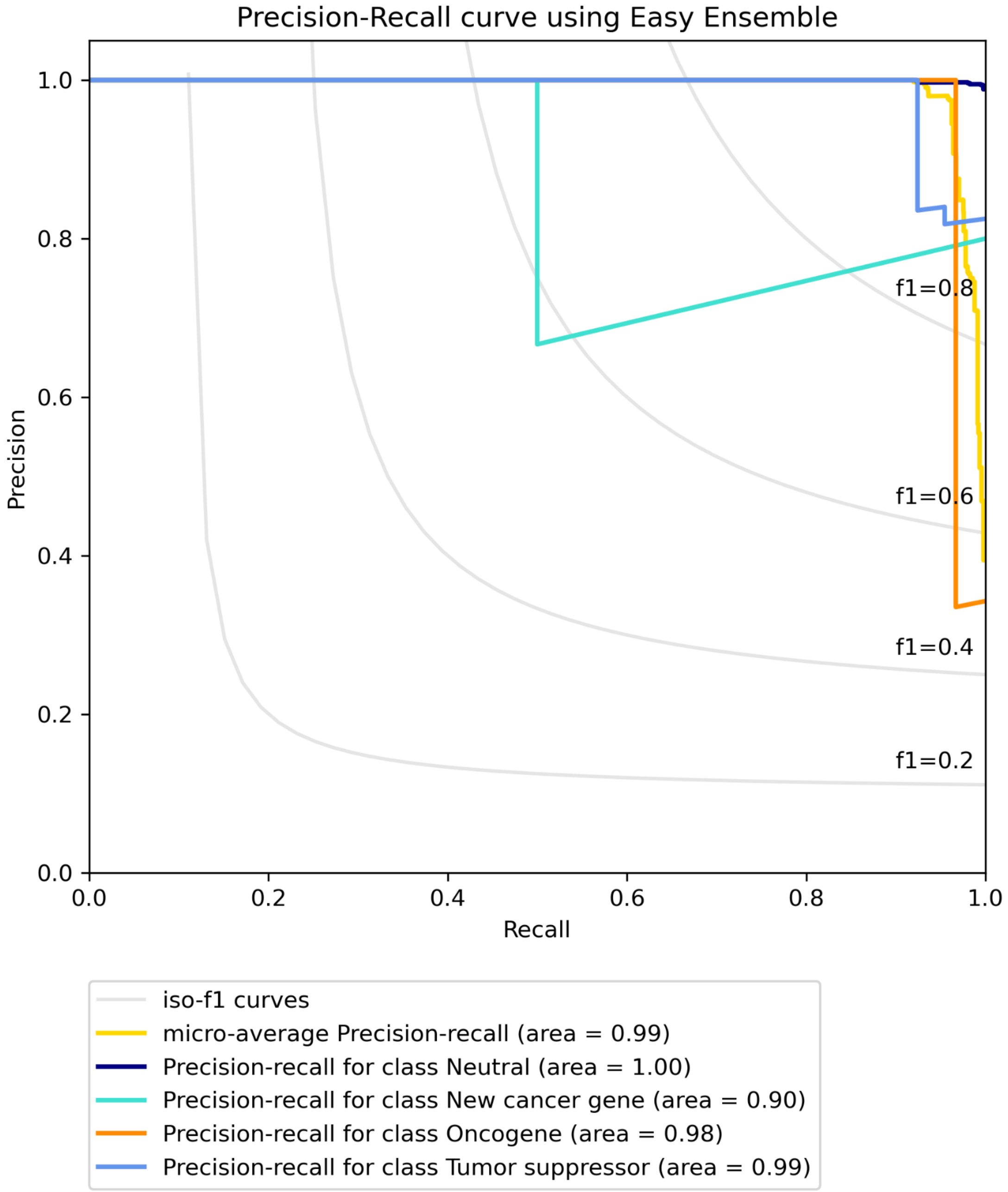


**Figure S22B. Precision-recall curve for BRCA using RNA data with Marellotto *et al.* labels built using Easy Ensemble.**

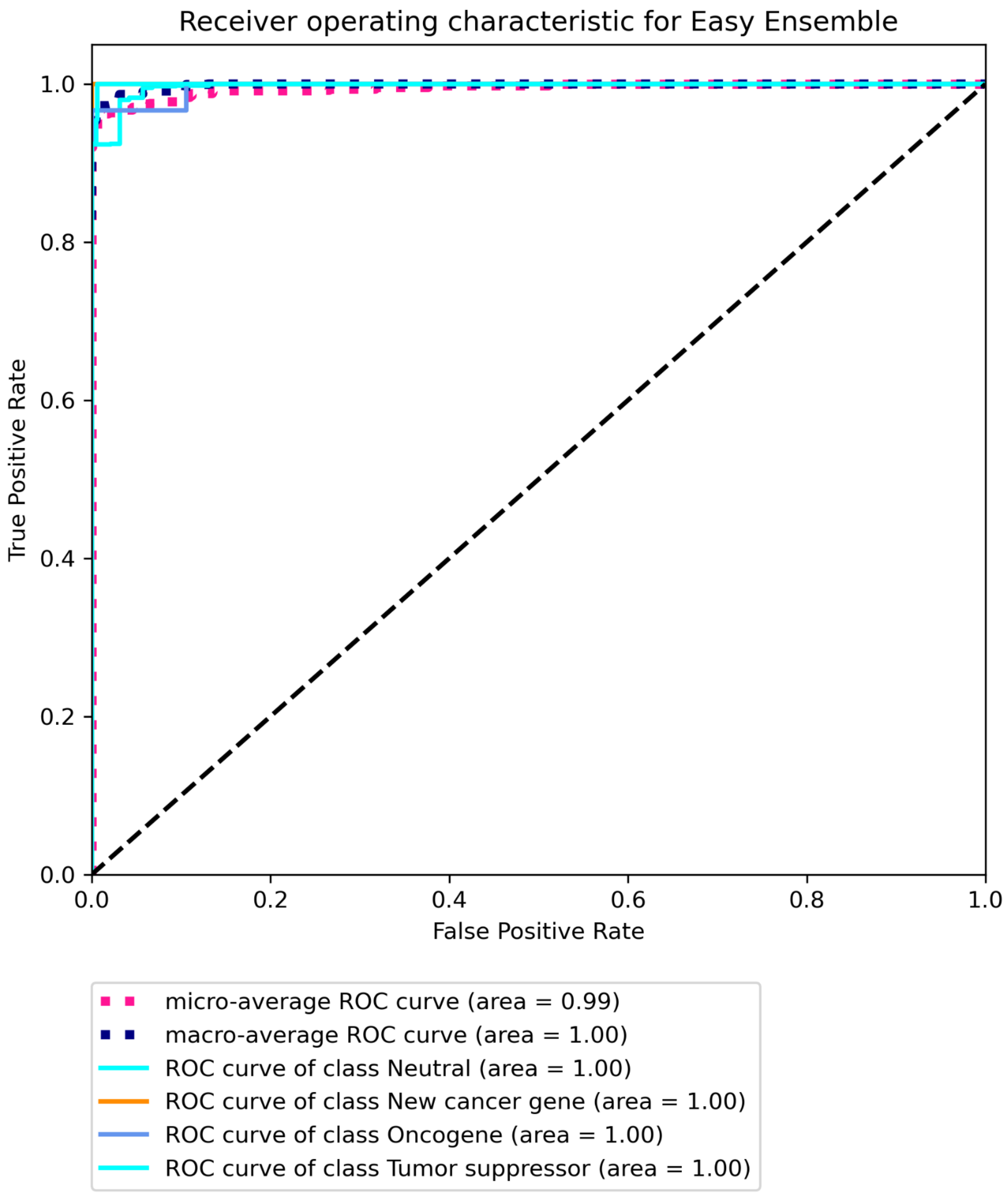


**Figure S22C Receiver operating characteristic for BRCA using RNA data with Marellotto *et al.* labels built using Easy Ensemble.**

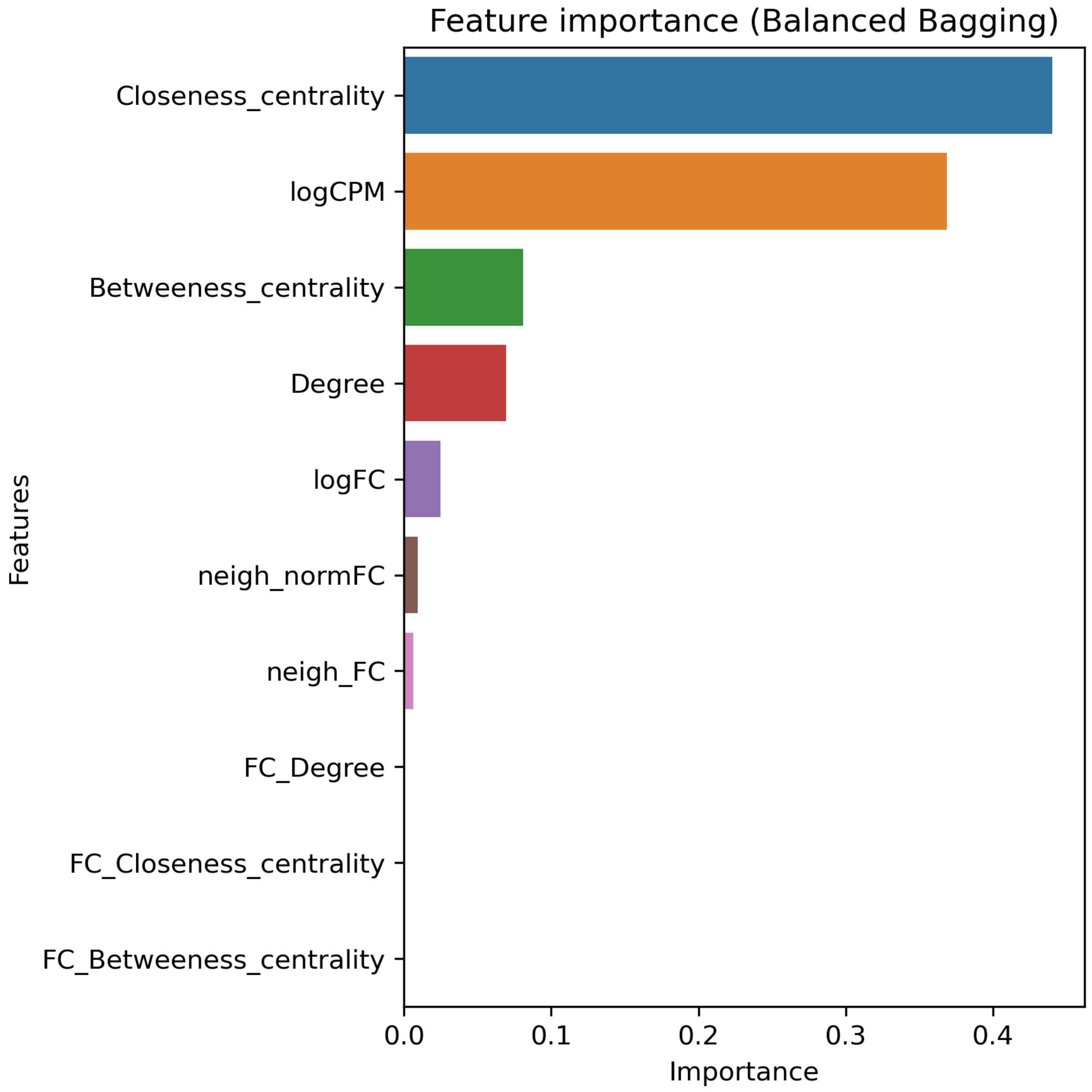


**Figure S23A. Top 20 features for COAD using RNA data with Bailey *et al.* labels built using Balanced bagging.**

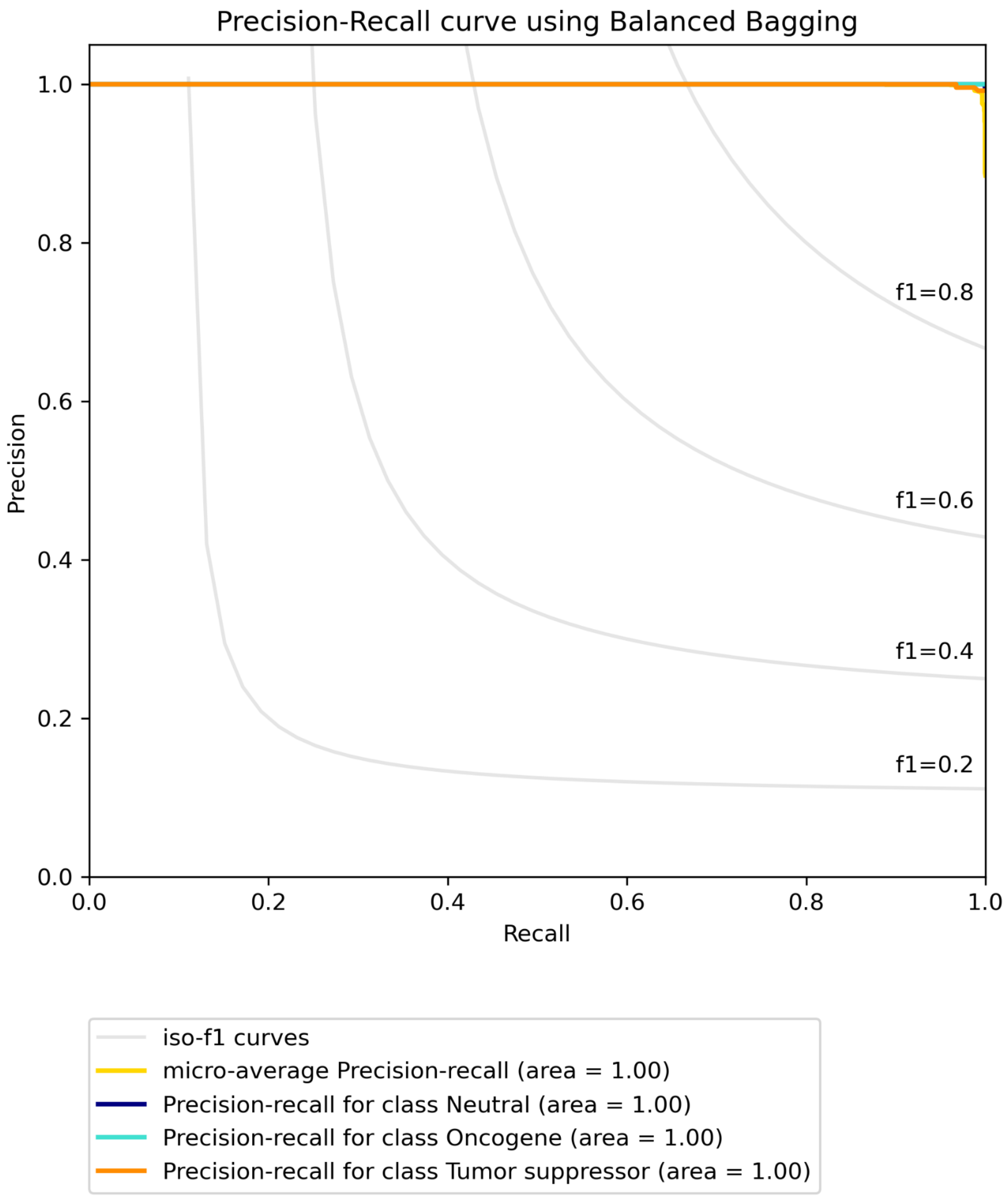


**Figure S23B. Precision-recall curve for COAD using RNA data with Bailey *et al.* labels built using Balanced bagging.**

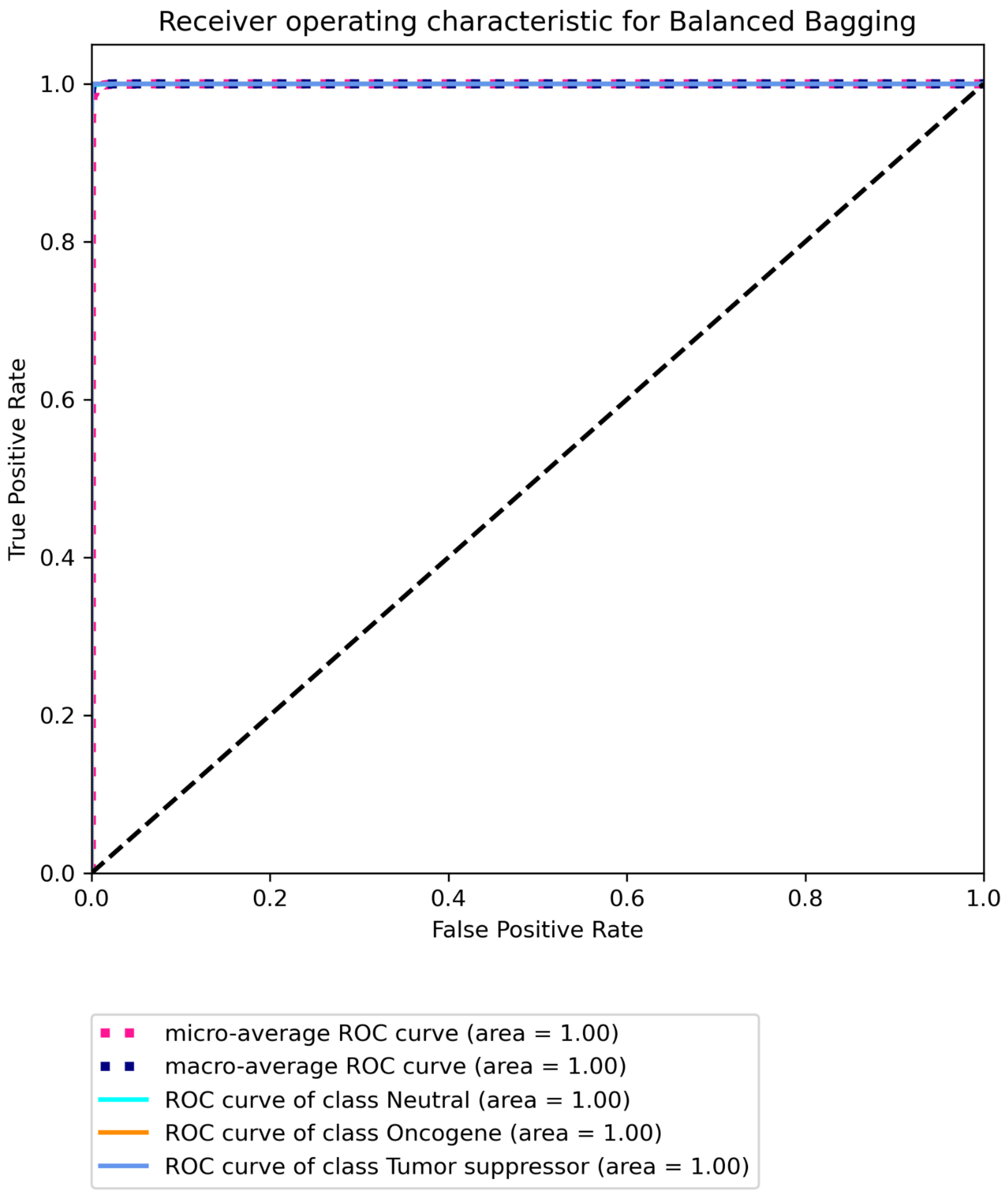


**Figure S23C Receiver operating characteristic for COAD using RNA data with Bailey *et al.* labels built using Balanced bagging.**

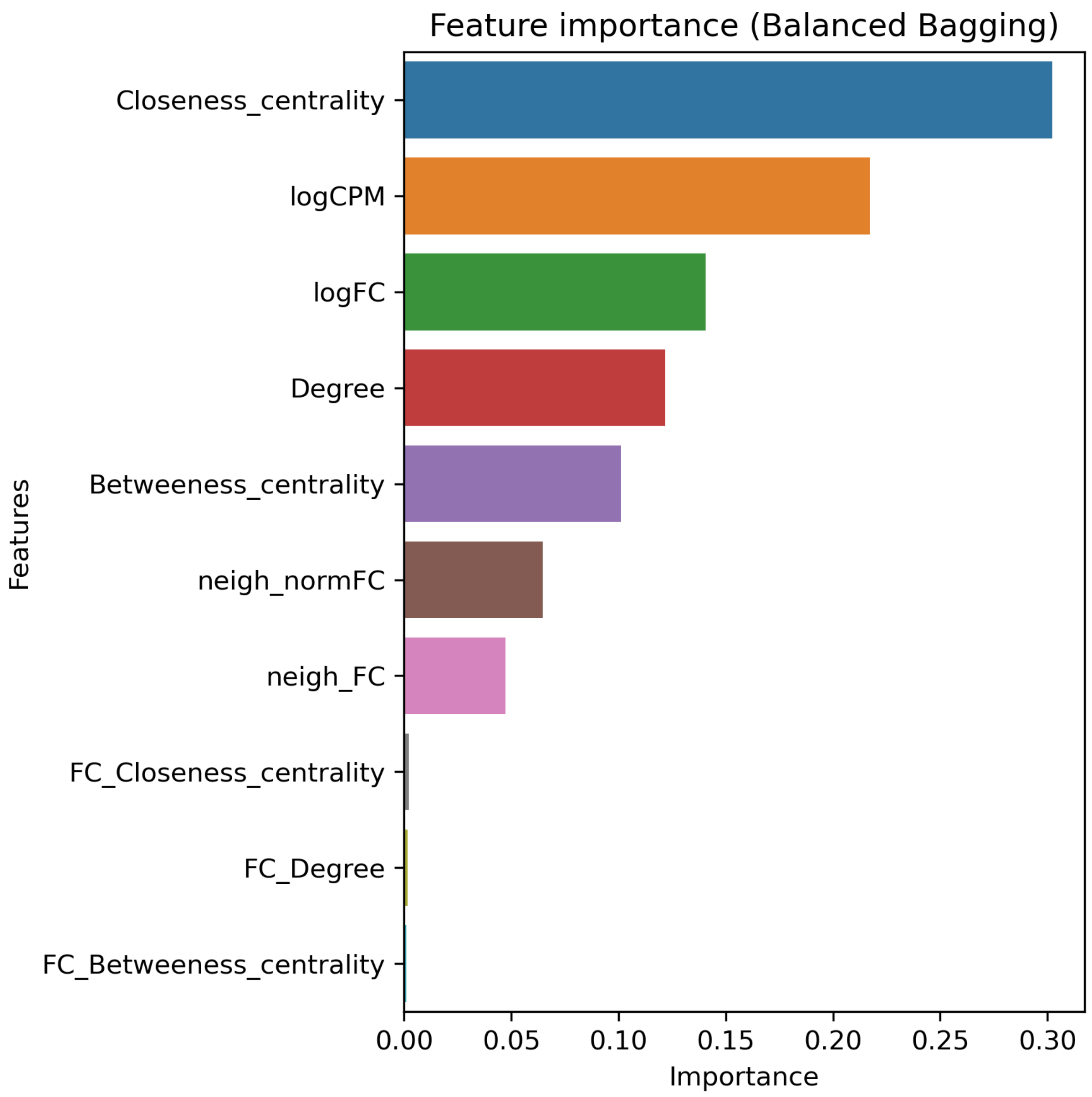


**Figure S24A. Top 20 features for COAD using RNA data with CGC labels built using Balanced bagging.**

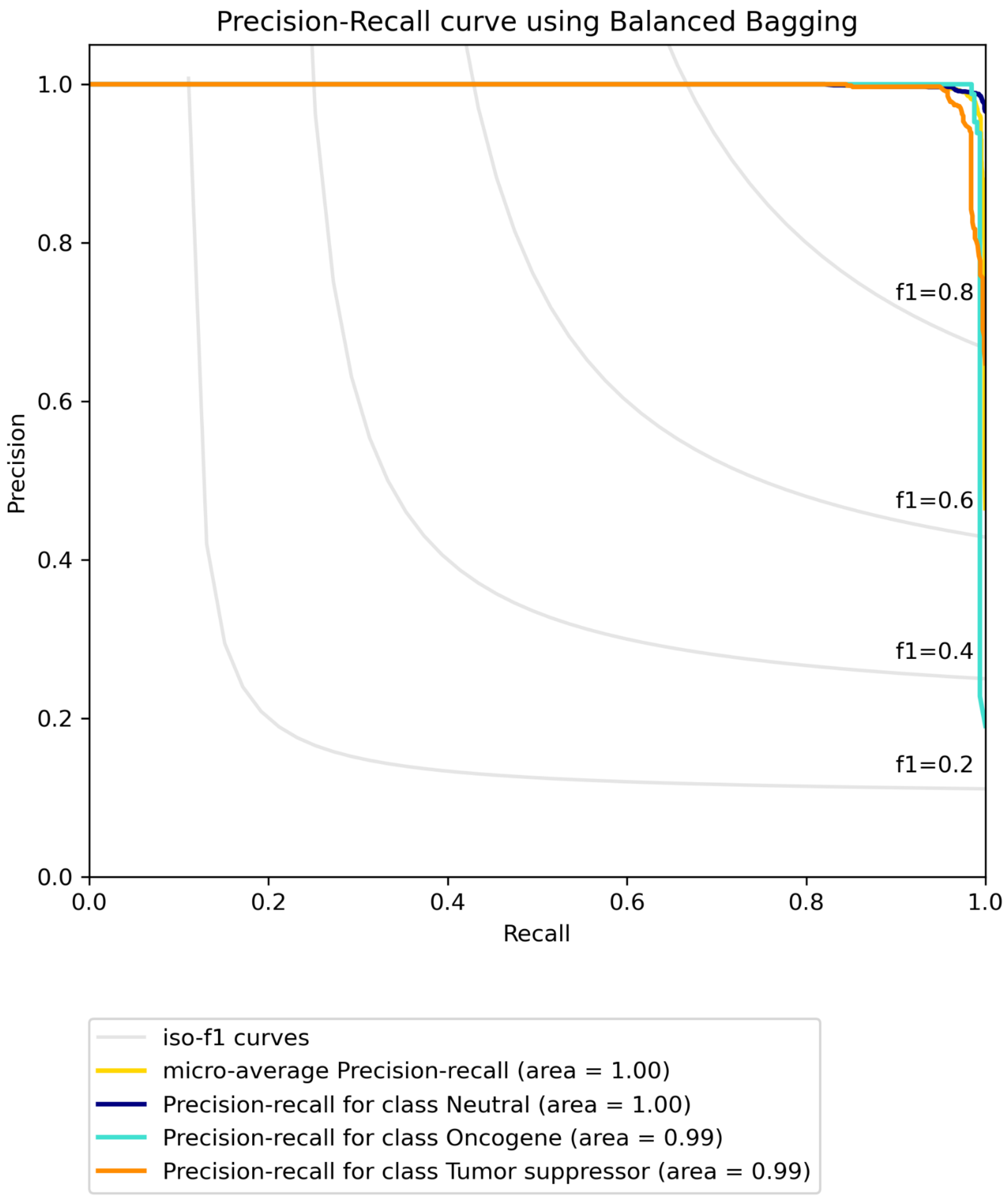


**Figure S24B. Precision-recall curve for COAD using RNA data with CGC labels built using Balanced bagging.**

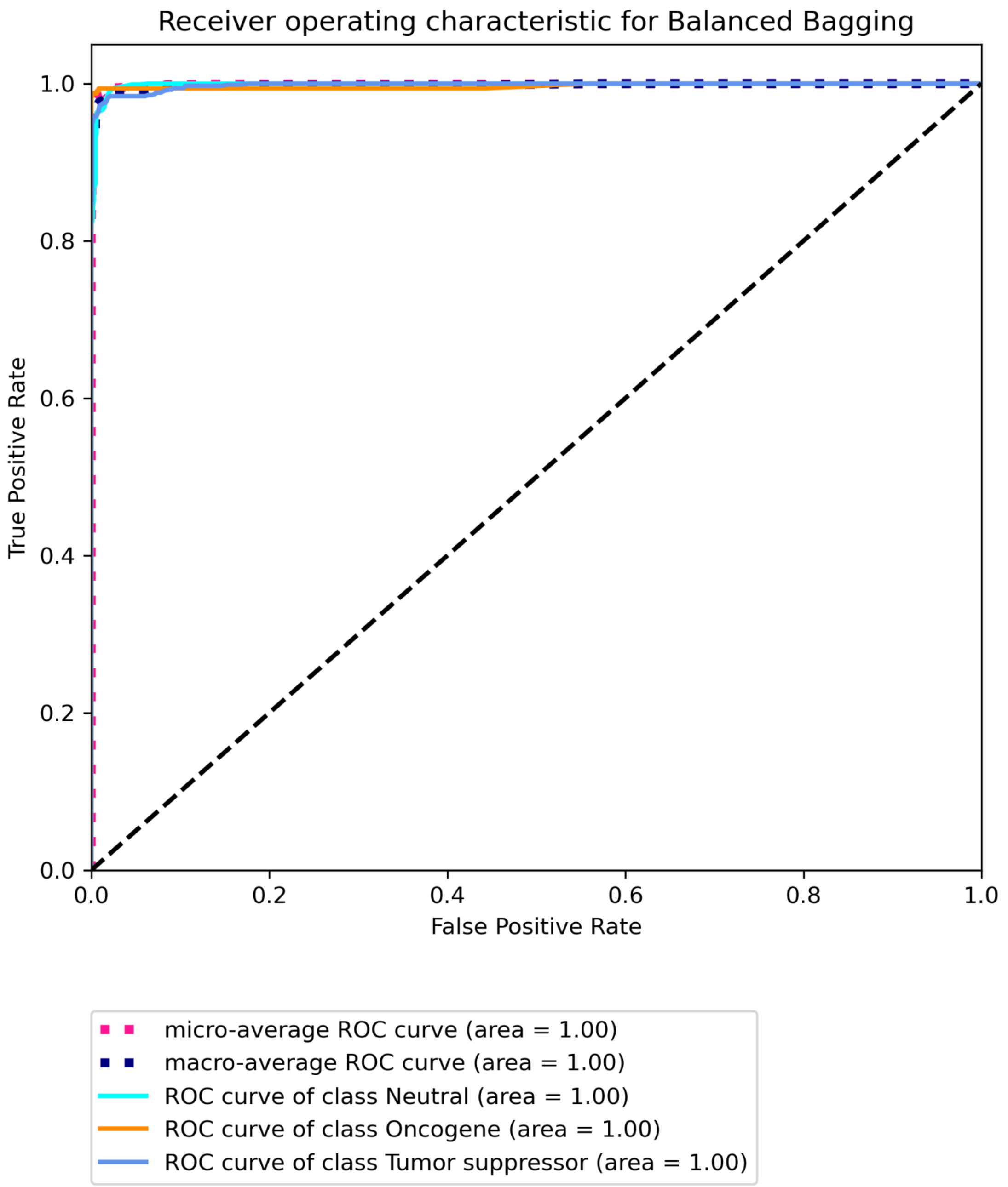


**Figure S24C Receiver operating characteristic for COAD using RNA data with CGC labels built using Balanced bagging.**

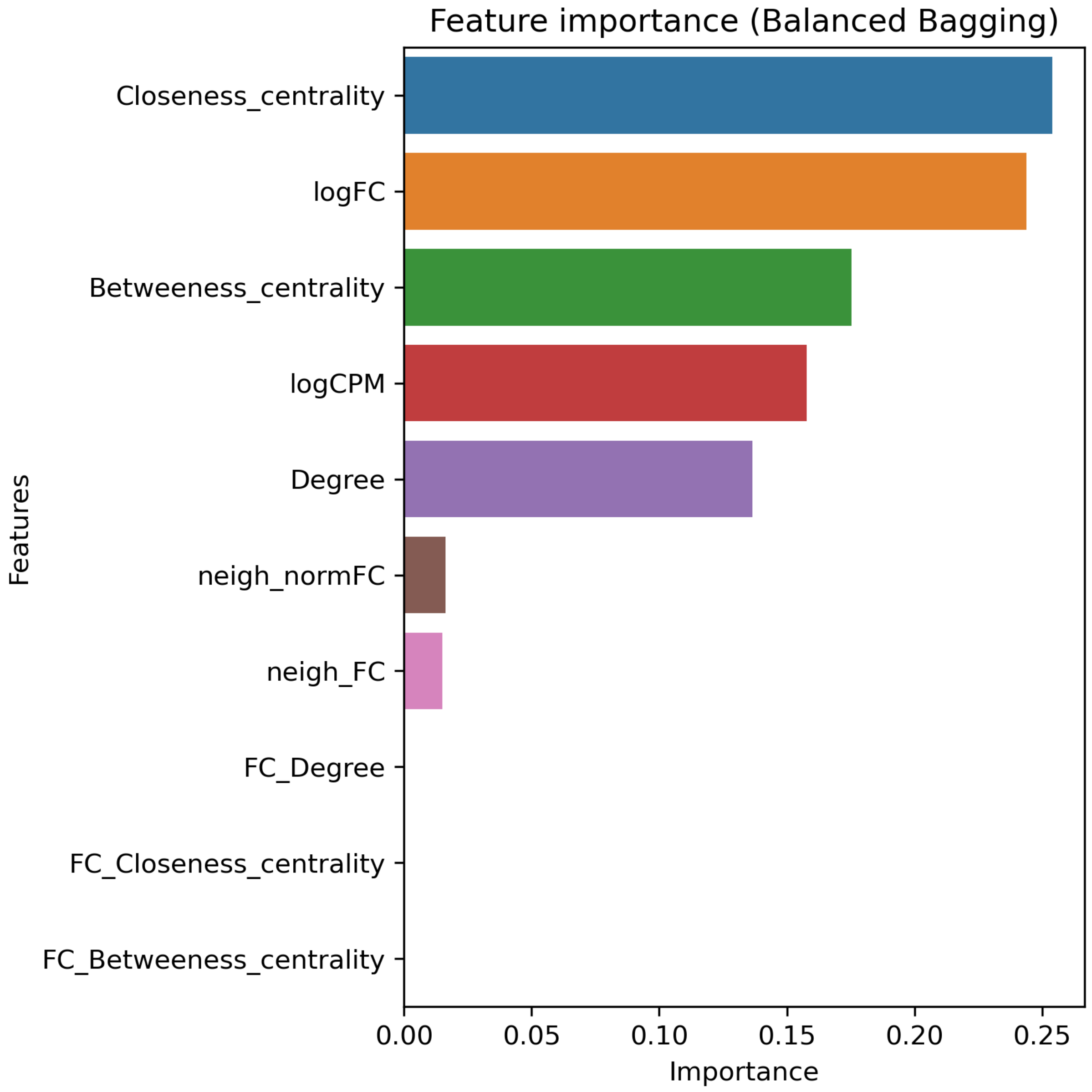


**Figure S25A. Top 20 features for LUAD using RNA data with Bailey *et al.* labels built using Balanced bagging.**

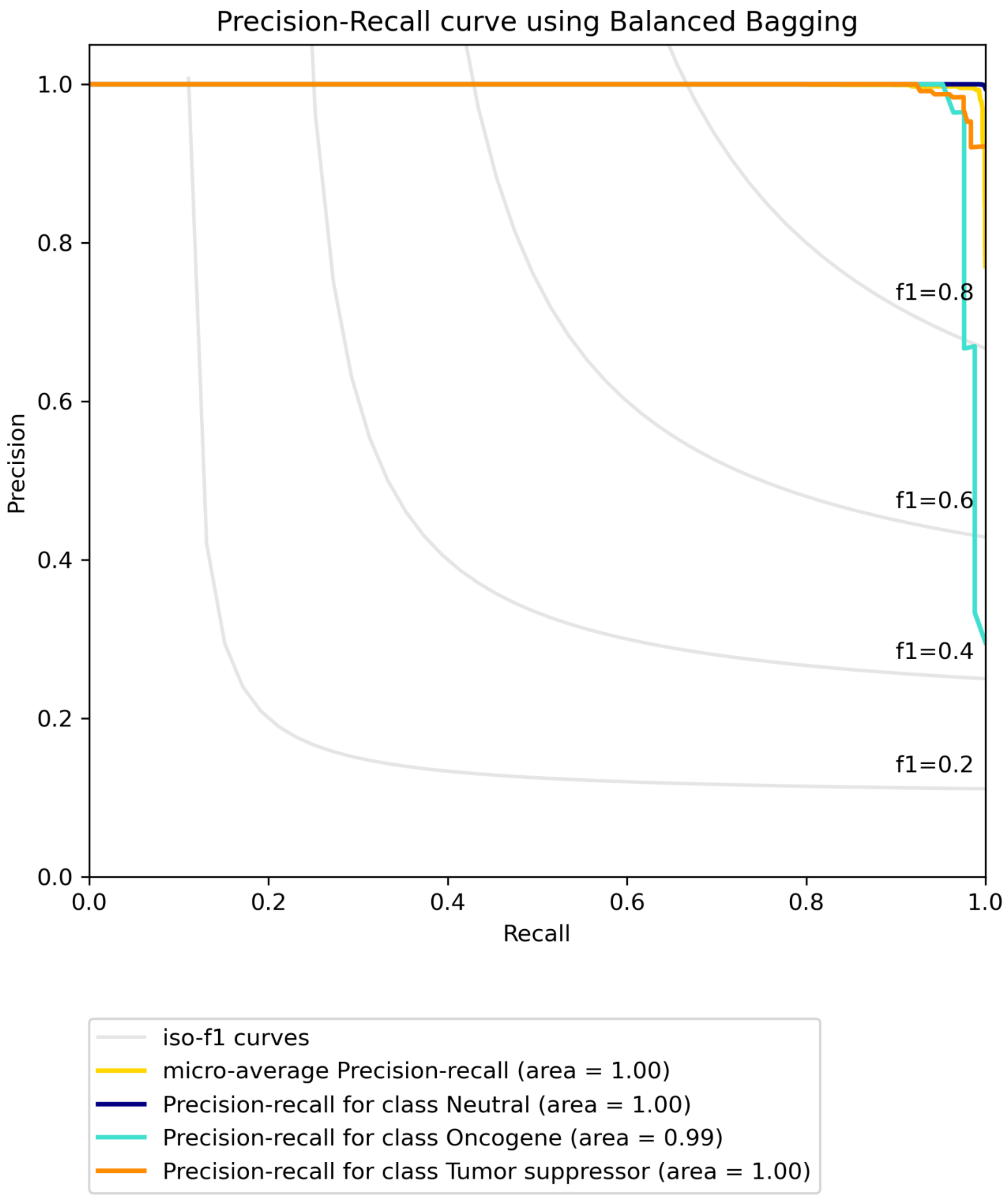


**Figure S25B. Precision-recall curve for LUAD using RNA data with Bailey *et al.* labels built using Balanced bagging.**

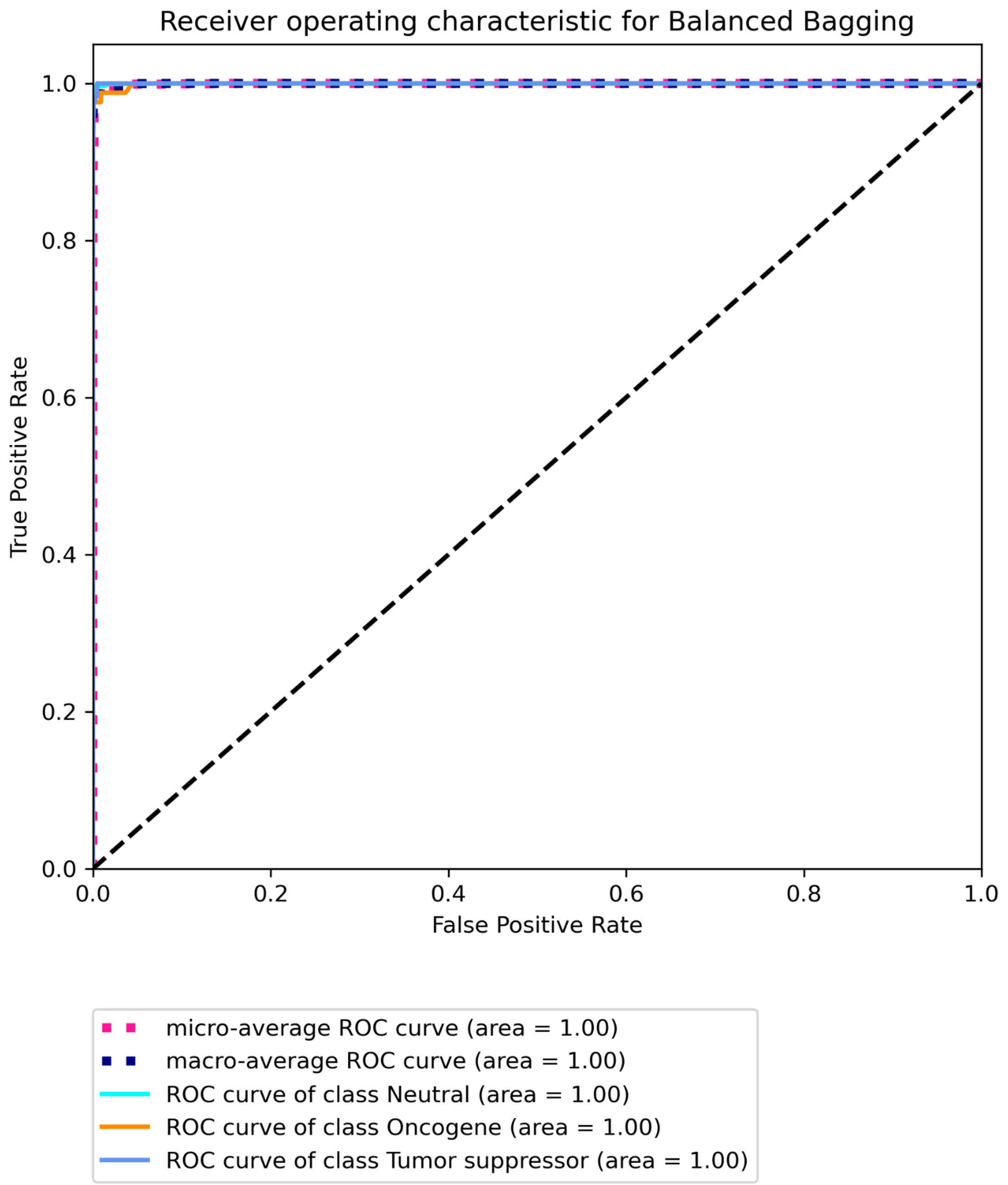


**Figure S25C Receiver operating characteristic for LUAD using RNA data with Bailey *et al.* labels built using Balanced bagging.**

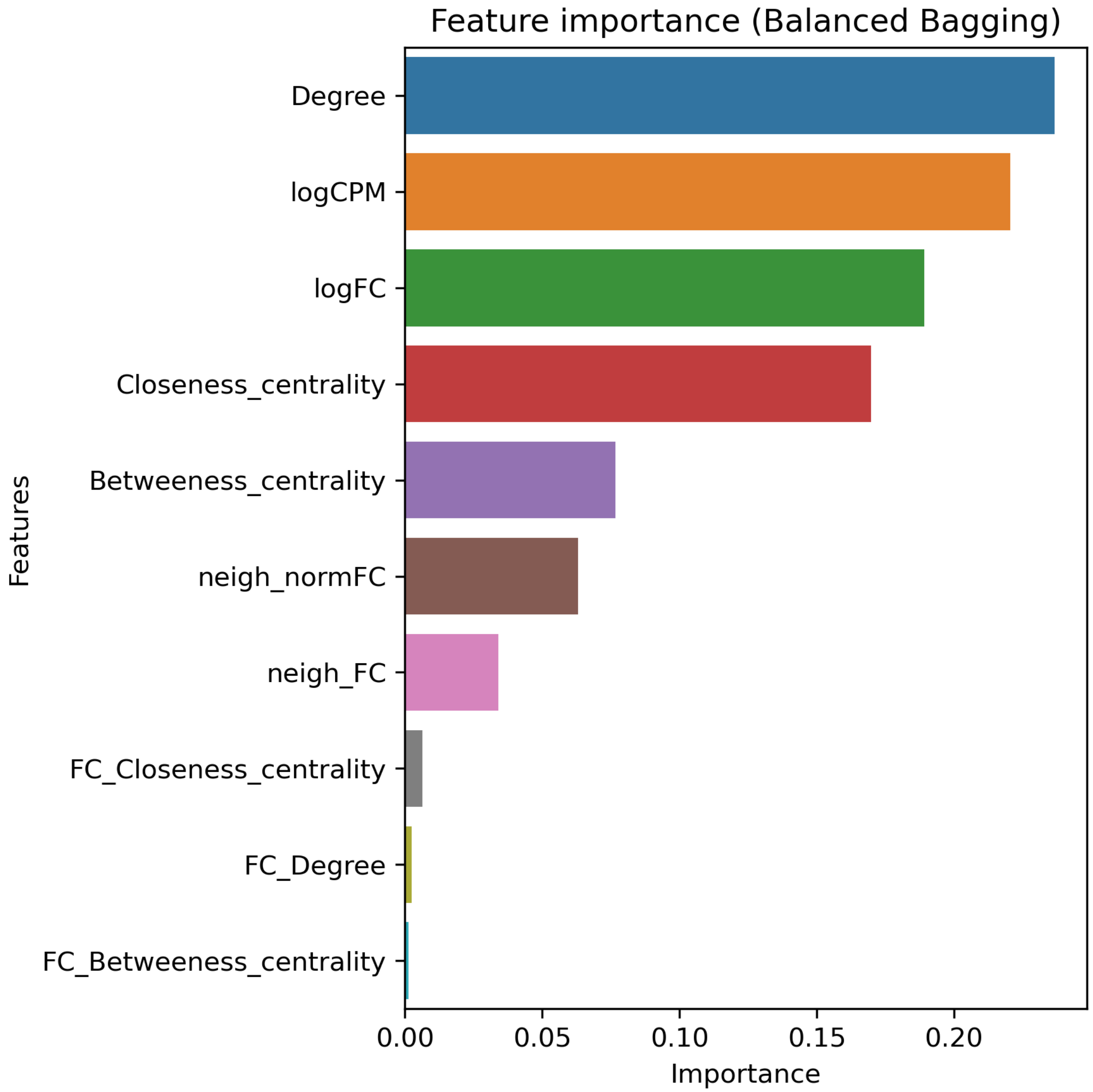


**Figure S26A. Top 20 features for LUAD using RNA data with CGC labels built using Balanced bagging.**

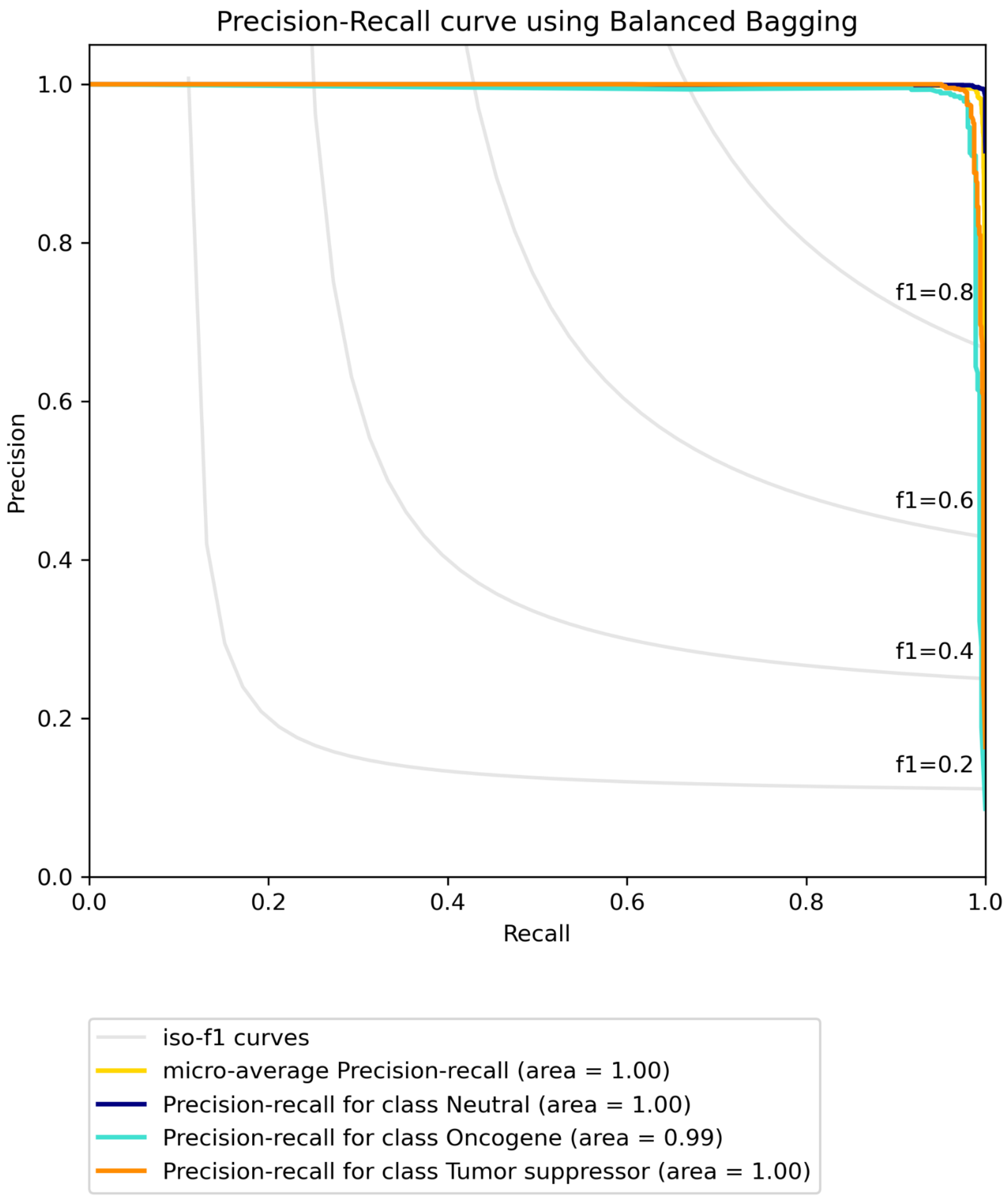


**Figure S26B. Precision-recall curve for LUAD using RNA data with CGC labels built using Balanced bagging.**

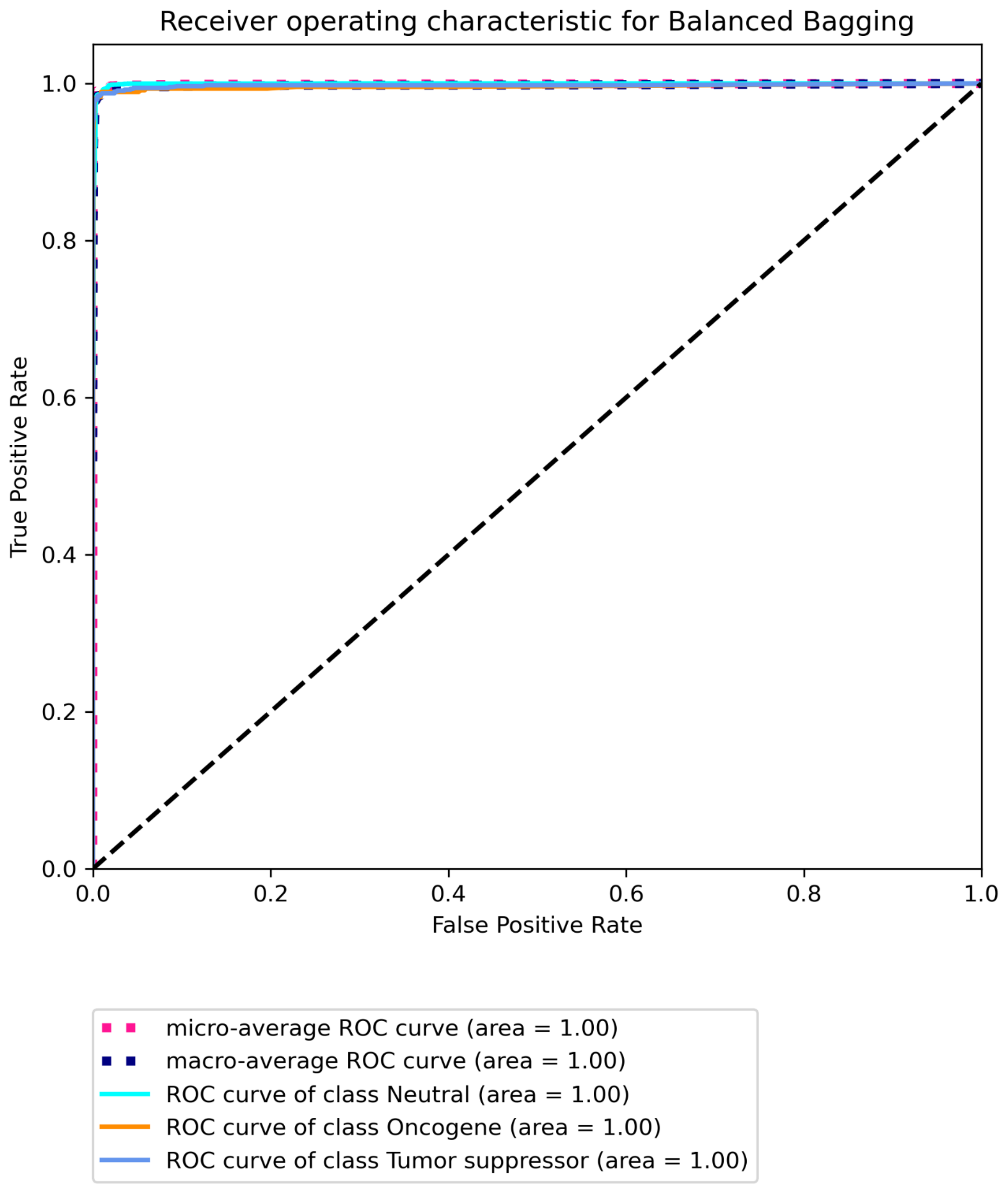


**Figure S26C Receiver operating characteristic for LUAD using RNA data with CGC labels built using Balanced bagging.**

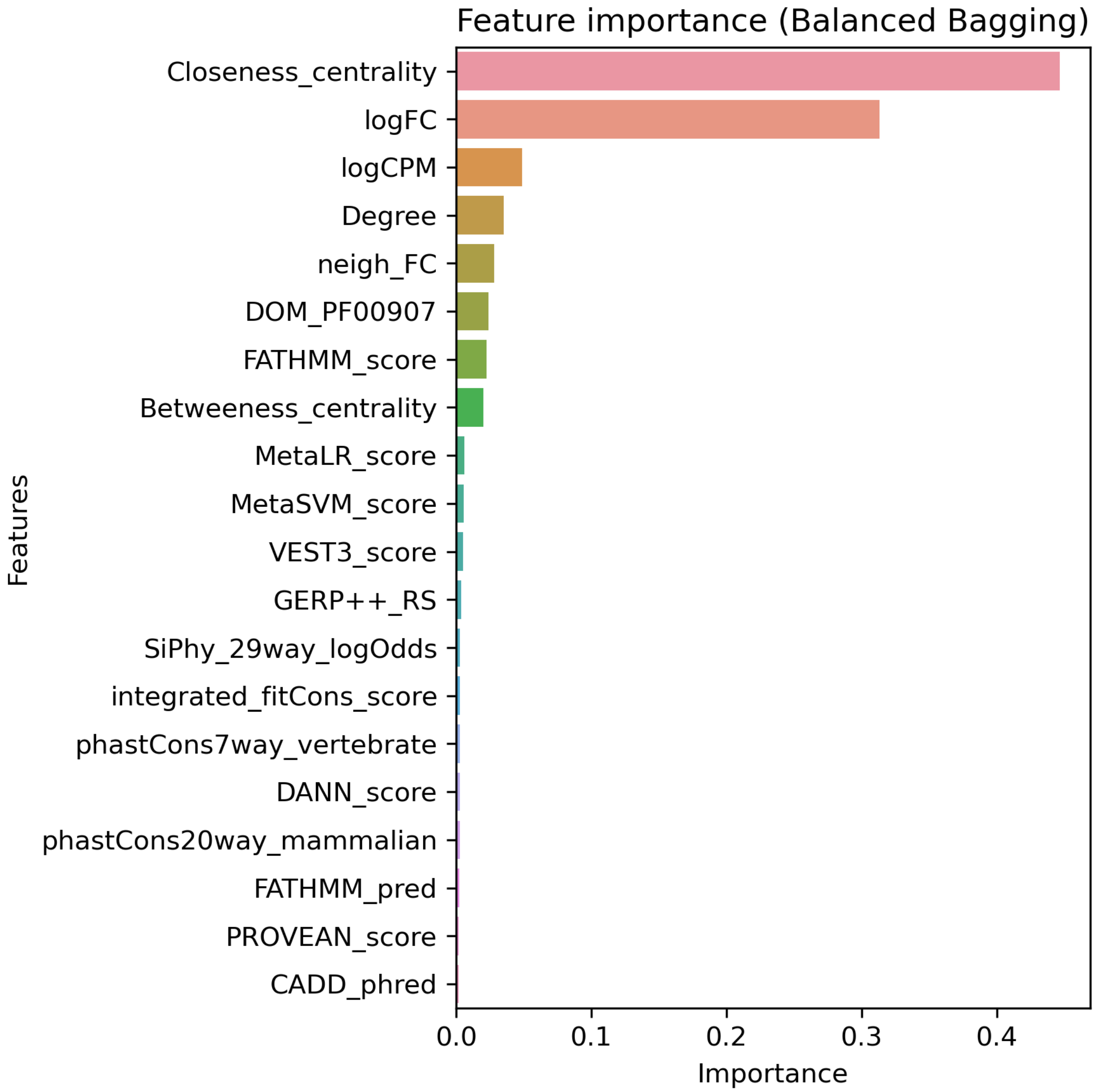


**Figure S27A. Top 20 features for BRCA using multi-omic data with Bailey *et al.* labels built on "all" features using Balanced bagging.**

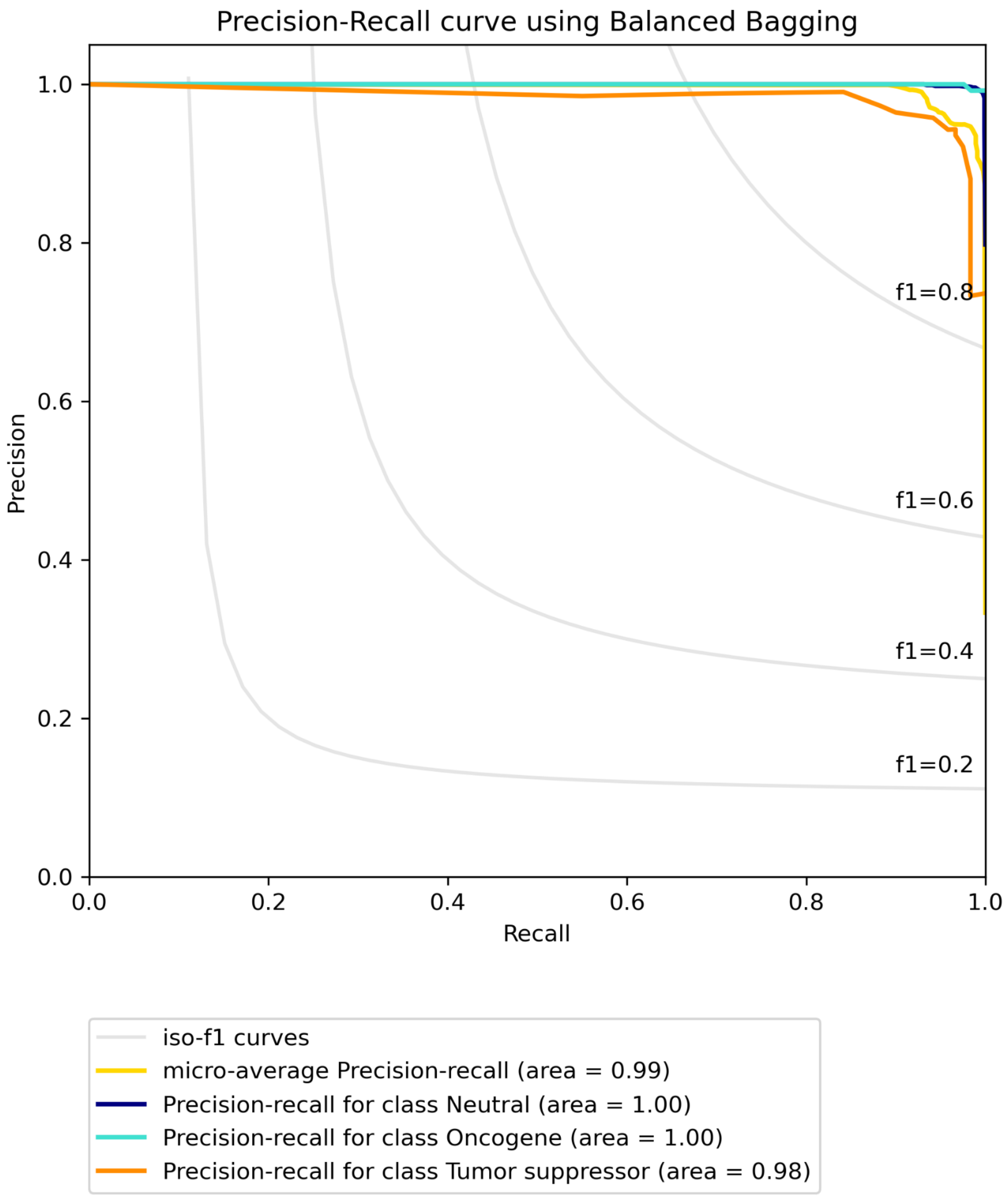


**Figure S27B. Precision-recall curve for BRCA using multi-omic data with Bailey *et al.* labels built on "all" features using Balanced bagging.**

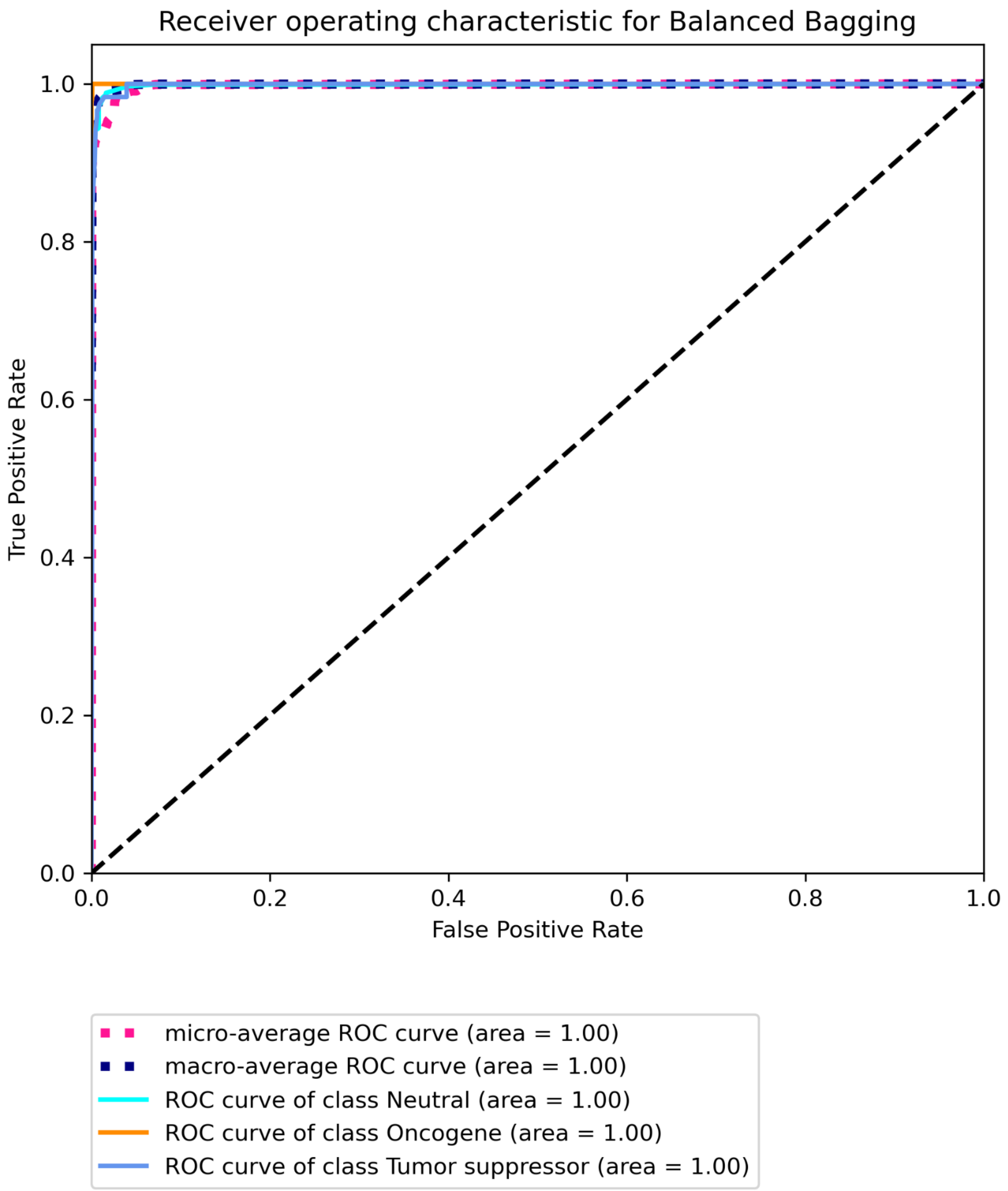


**Figure S27C Receiver operating characteristic for BRCA using multi-omic data with Bailey *et al.* labels built on "all" features using Balanced bagging.**

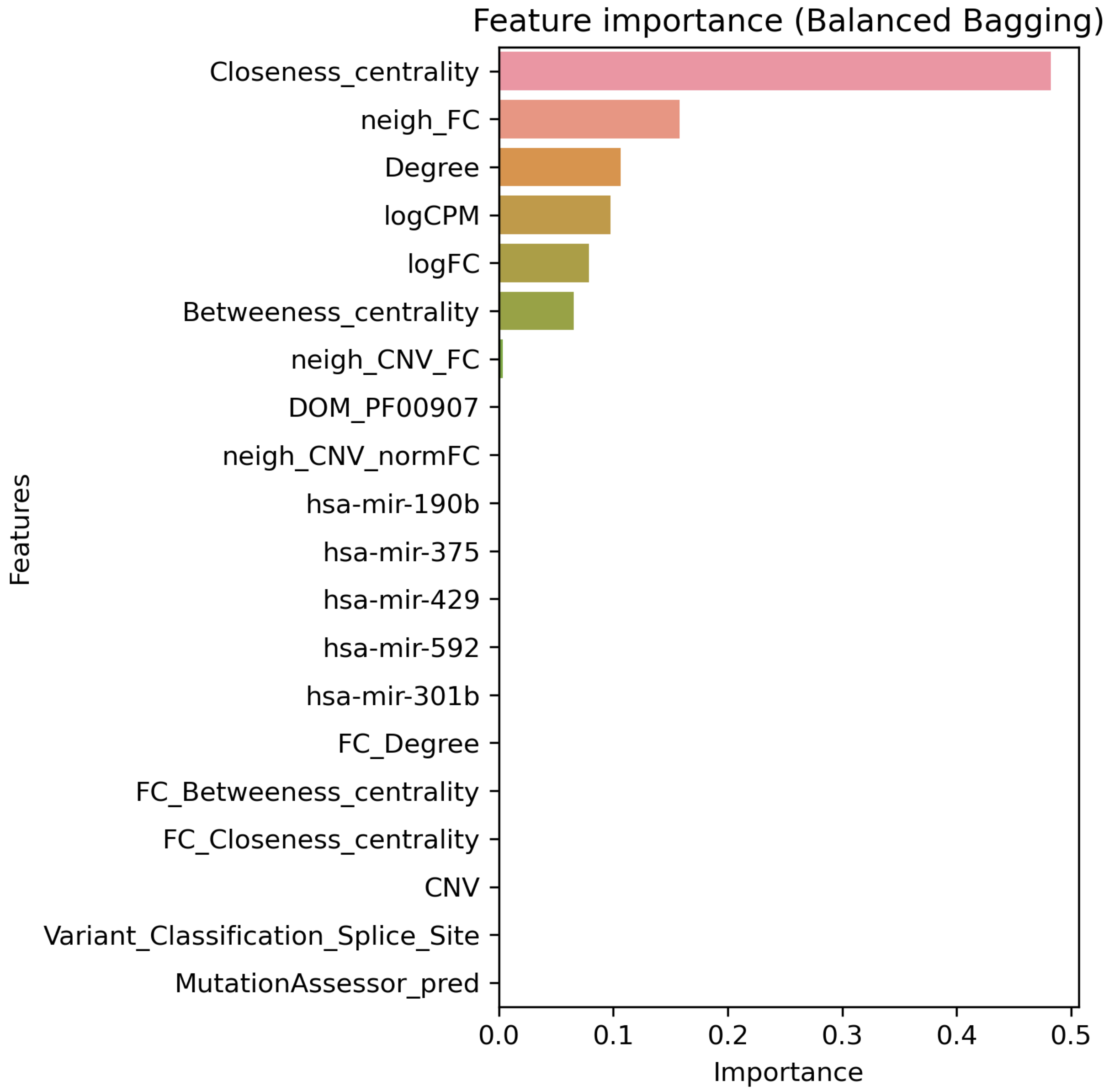


**Figure S28A. Top 20 features for BRCA using multi-omic data with Bailey *et al.* labels built on "some" features using Balanced bagging.**

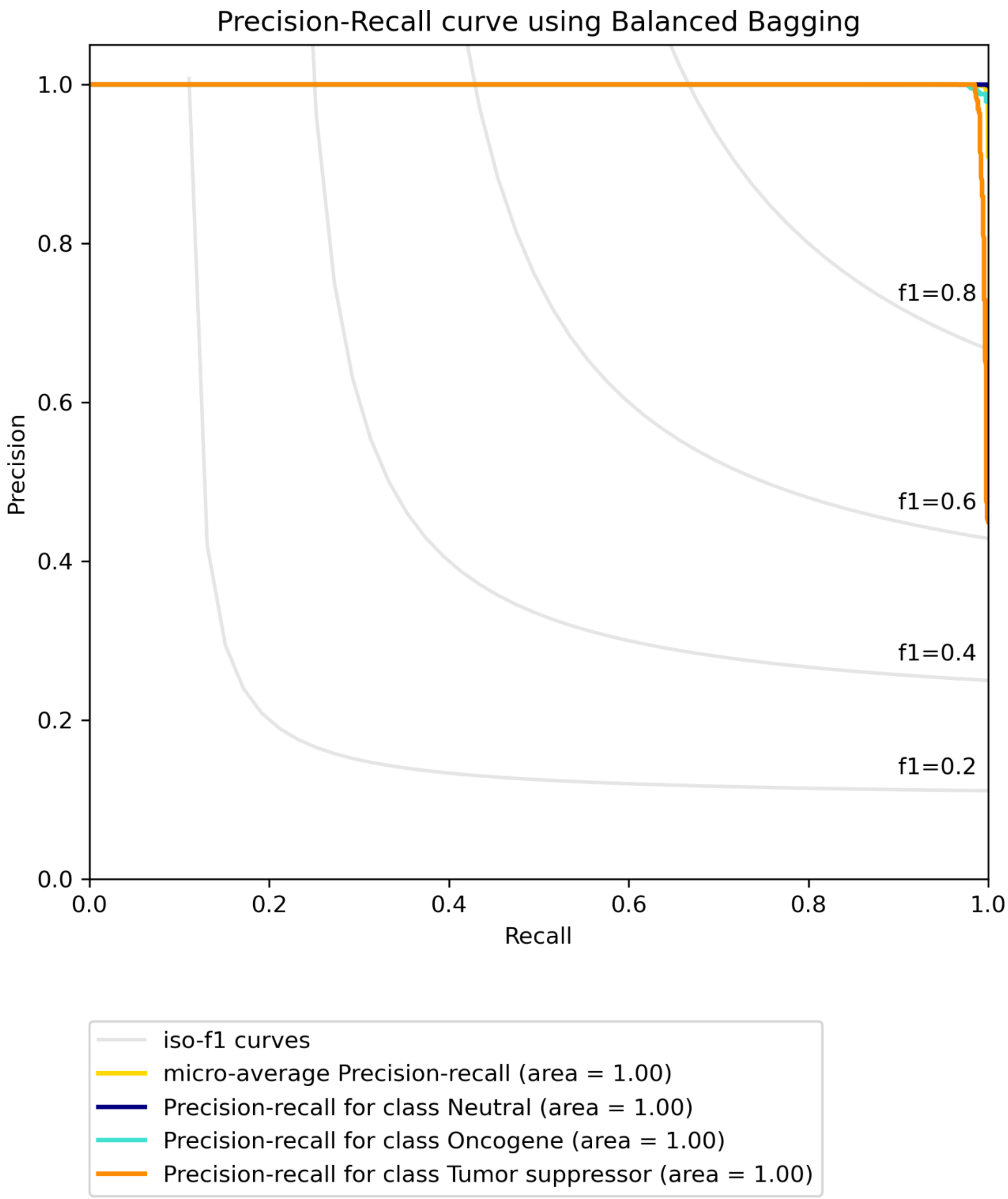


**Figure S28B. Precision-recall curve for BRCA using multi-omic data with Bailey *et al.* labels built on "some" features using Balanced bagging.**

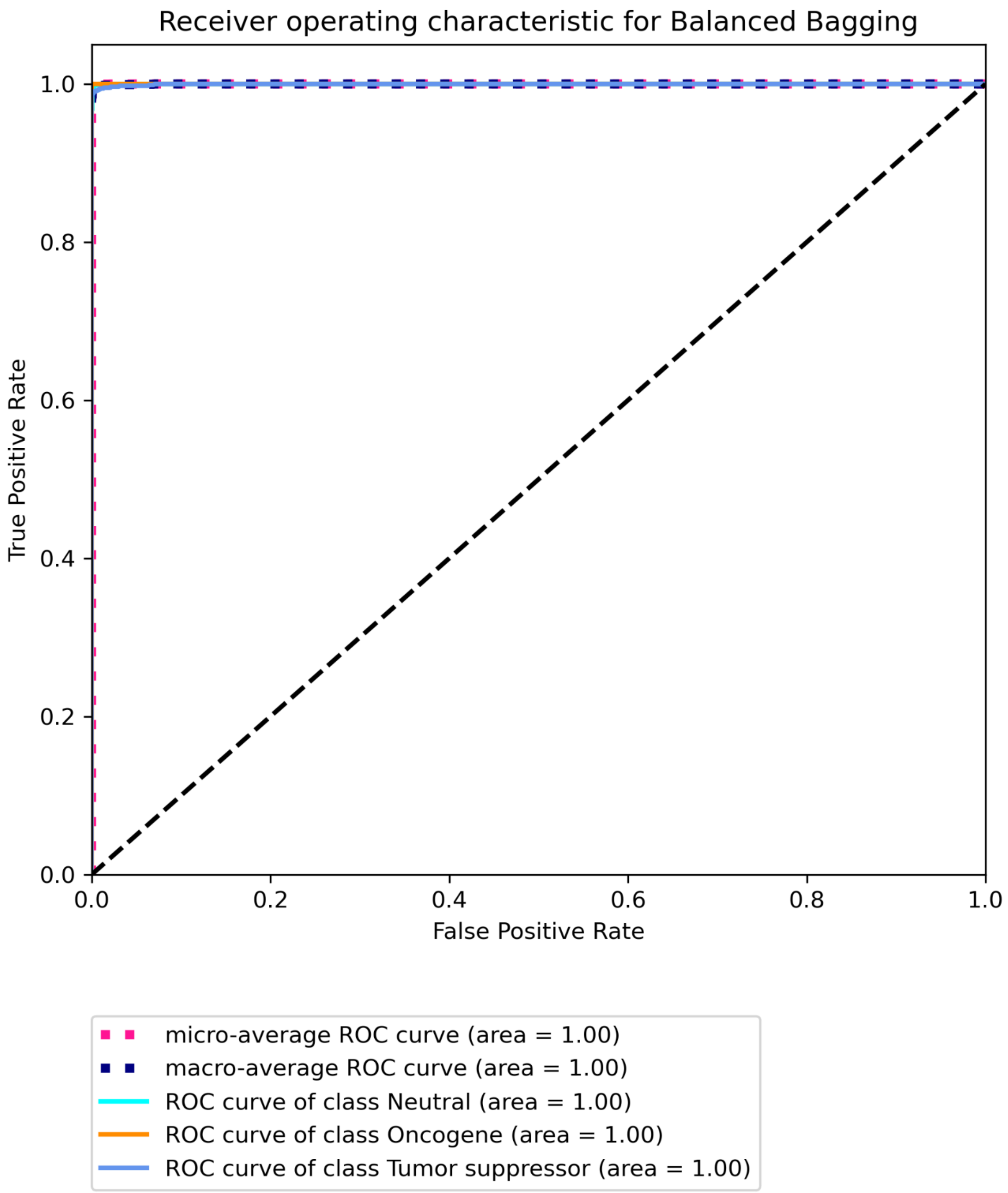


**Figure S28C Receiver operating characteristic for BRCA using multi-omic data with Bailey *et al.* labels built on “some” features using Balanced bagging.**

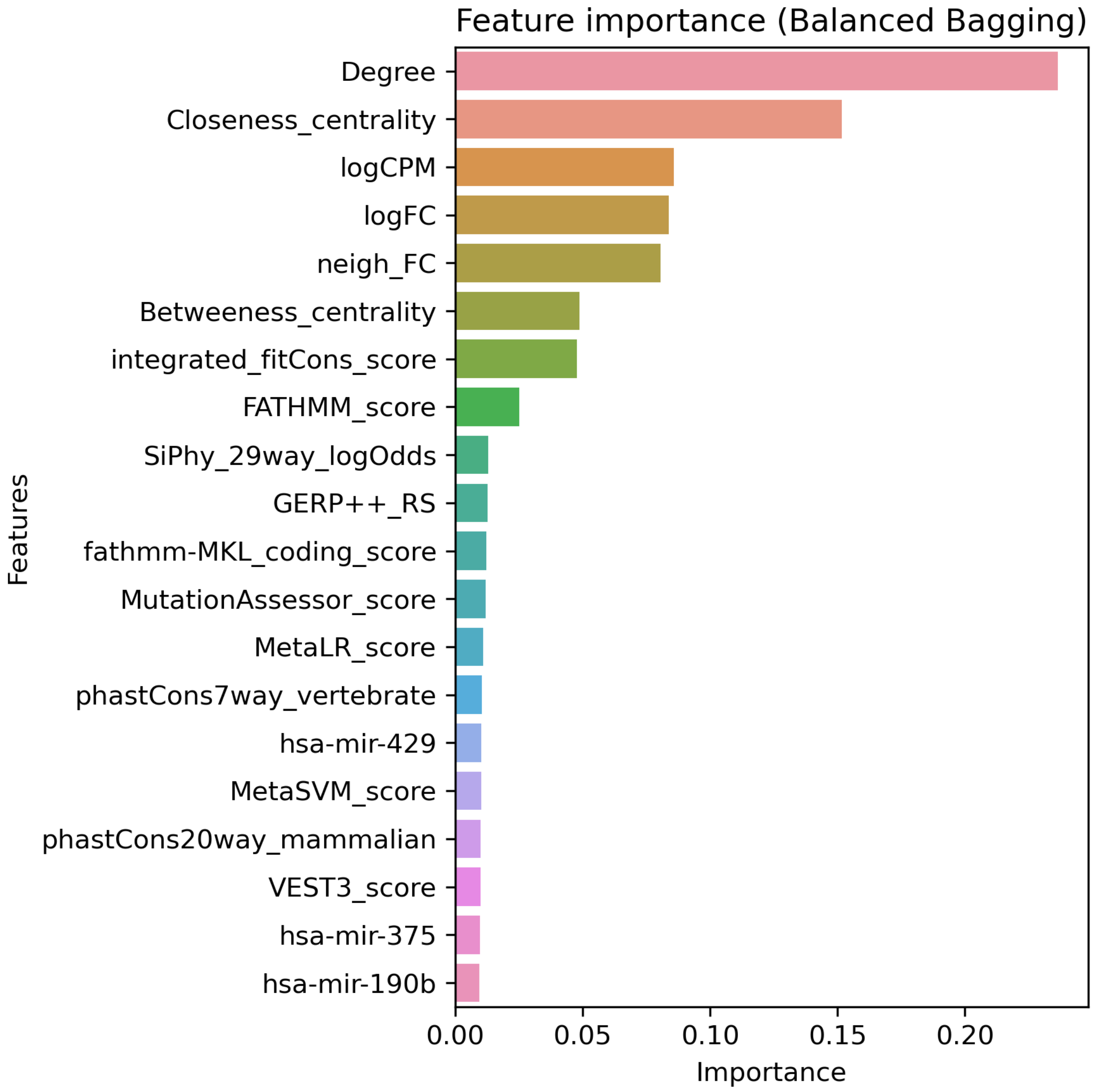


**Figure S29A. Top 20 features for BRCA using multi-omic data with CGC labels built on "all" features using Balanced bagging.**

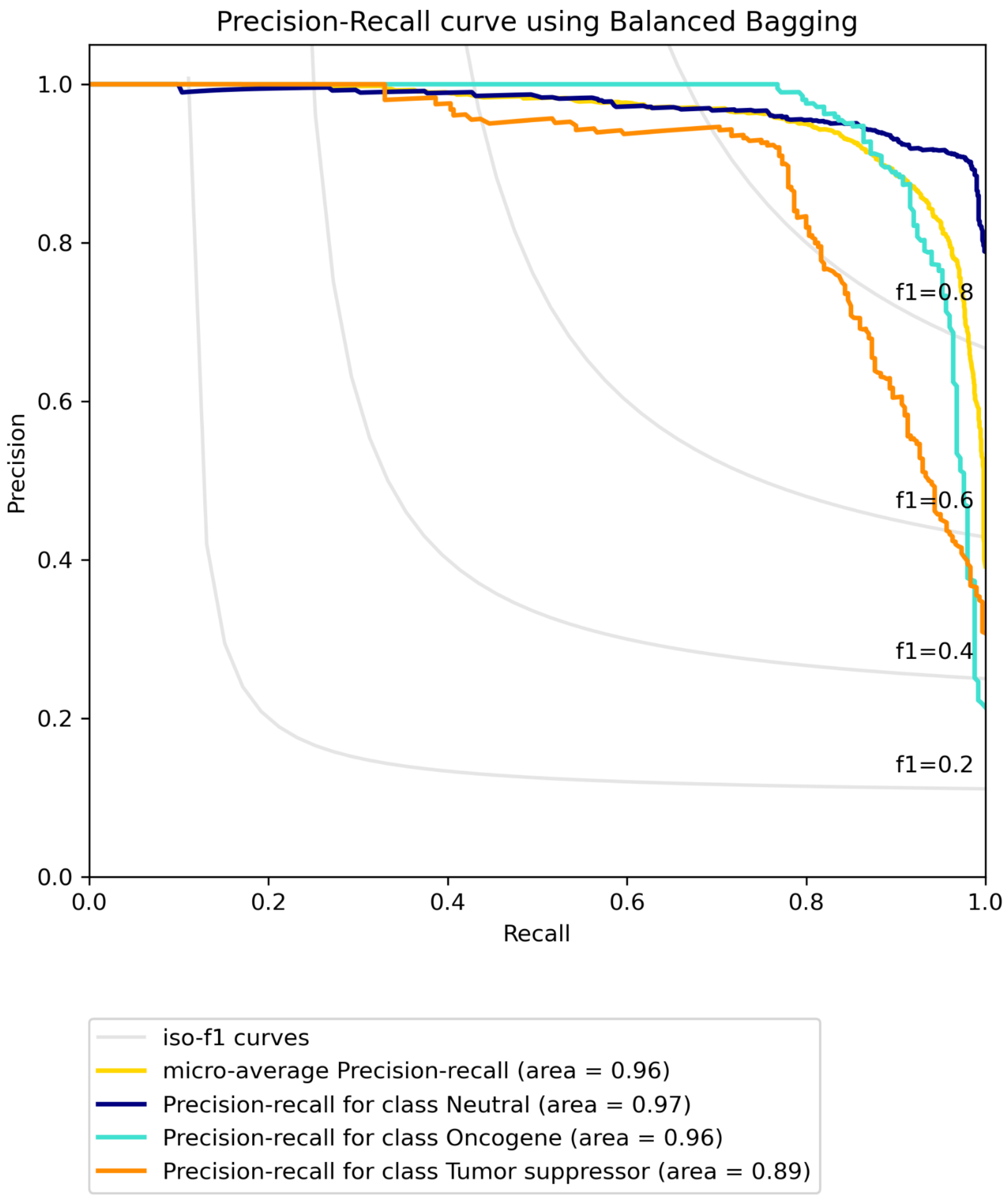


**Figure S29B. Precision-recall curve for BRCA using multi-omic data with CGC labels built on “all” features using Balanced bagging.**

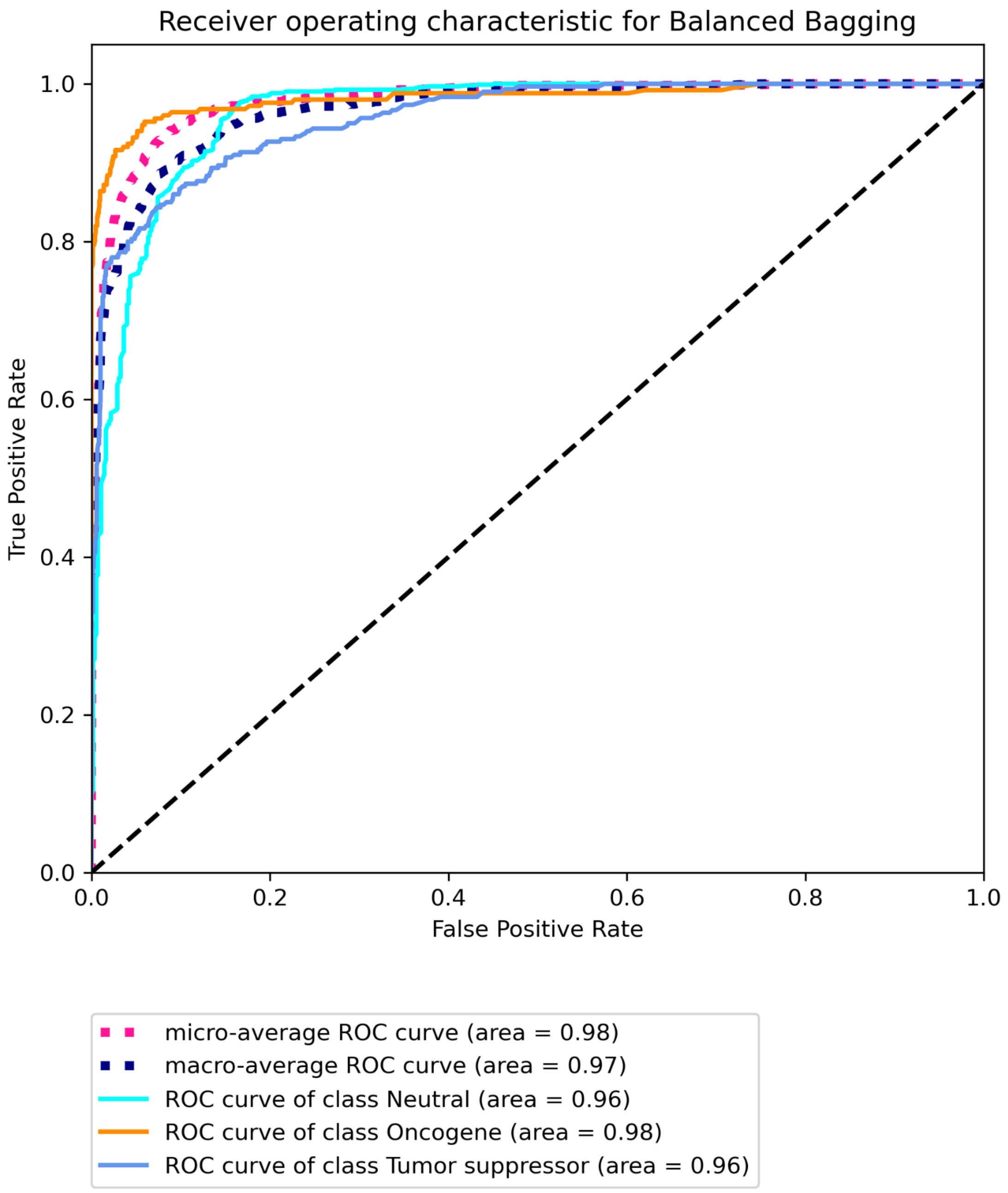


**Figure S29C. Receiver operating characteristic for BRCA using multi-omic data with CGC labels built on "all" features using Balanced Bagging.**

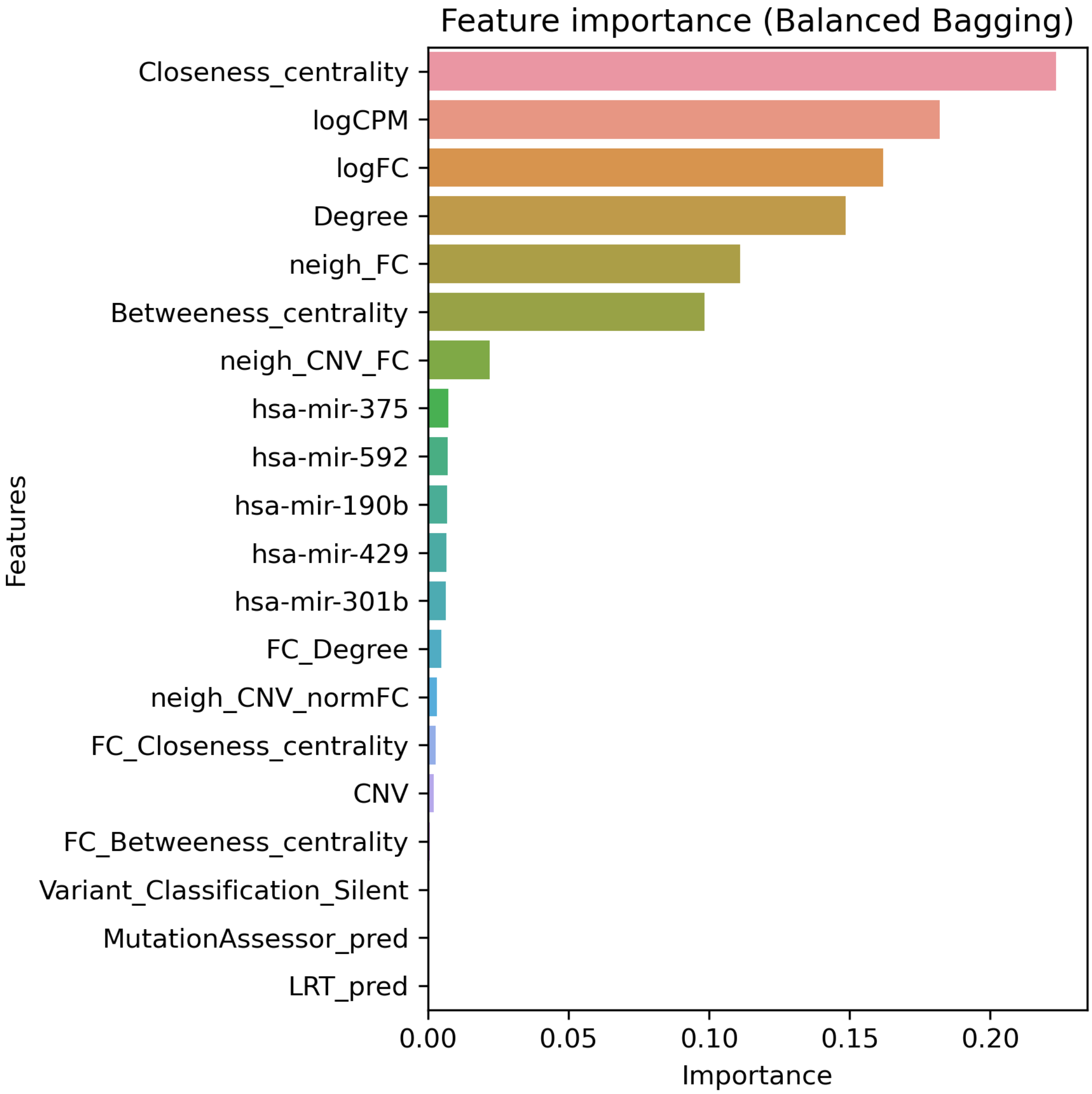


**Figure S30A. Top 20 features for BRCA using multi-omic data with CGC labels built on "some" features using Balanced Bagging.**

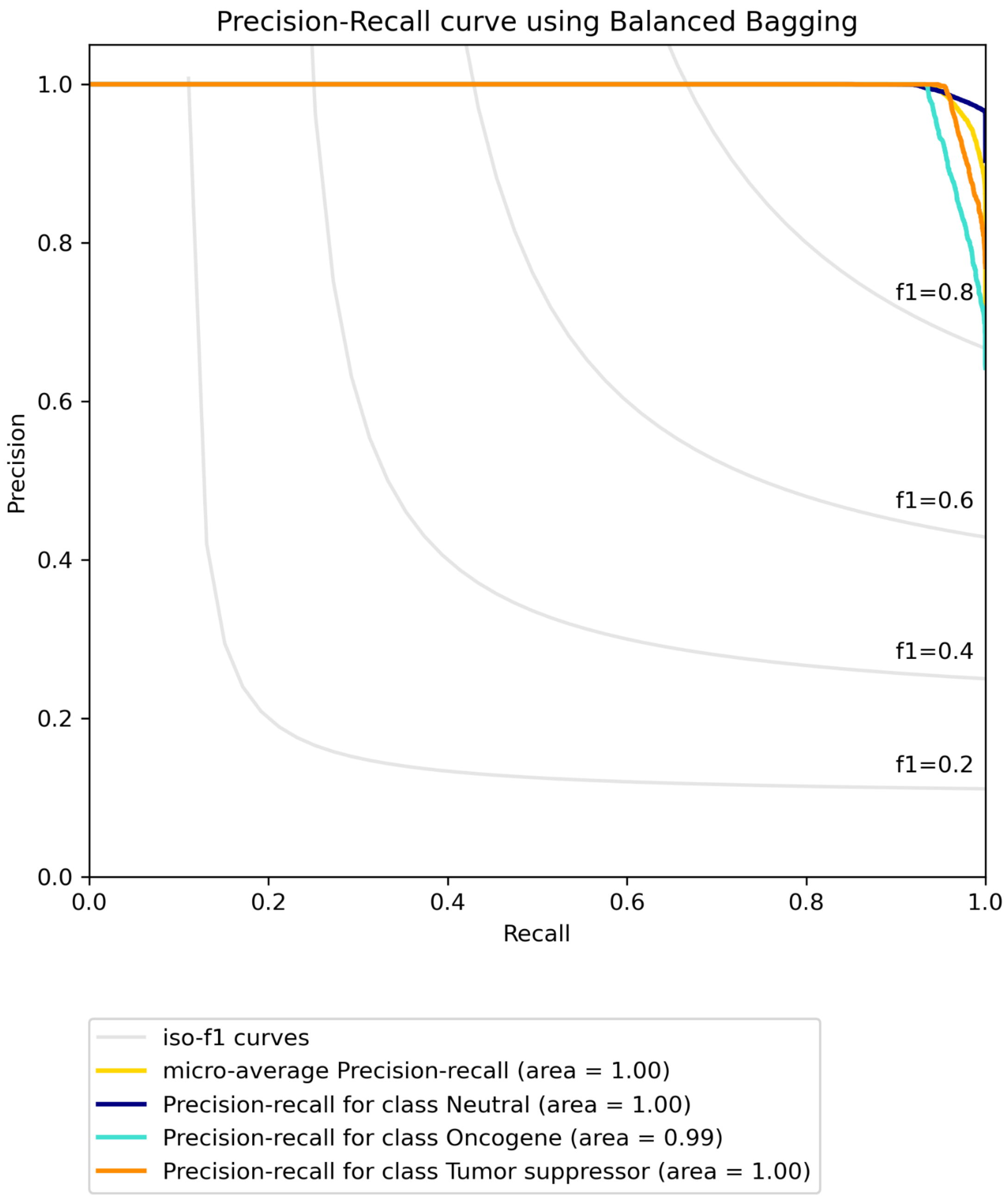


**Figure S30B. Precision-recall curve for BRCA using multi-omic data with CGC labels built on "some" features using Balanced Bagging.**

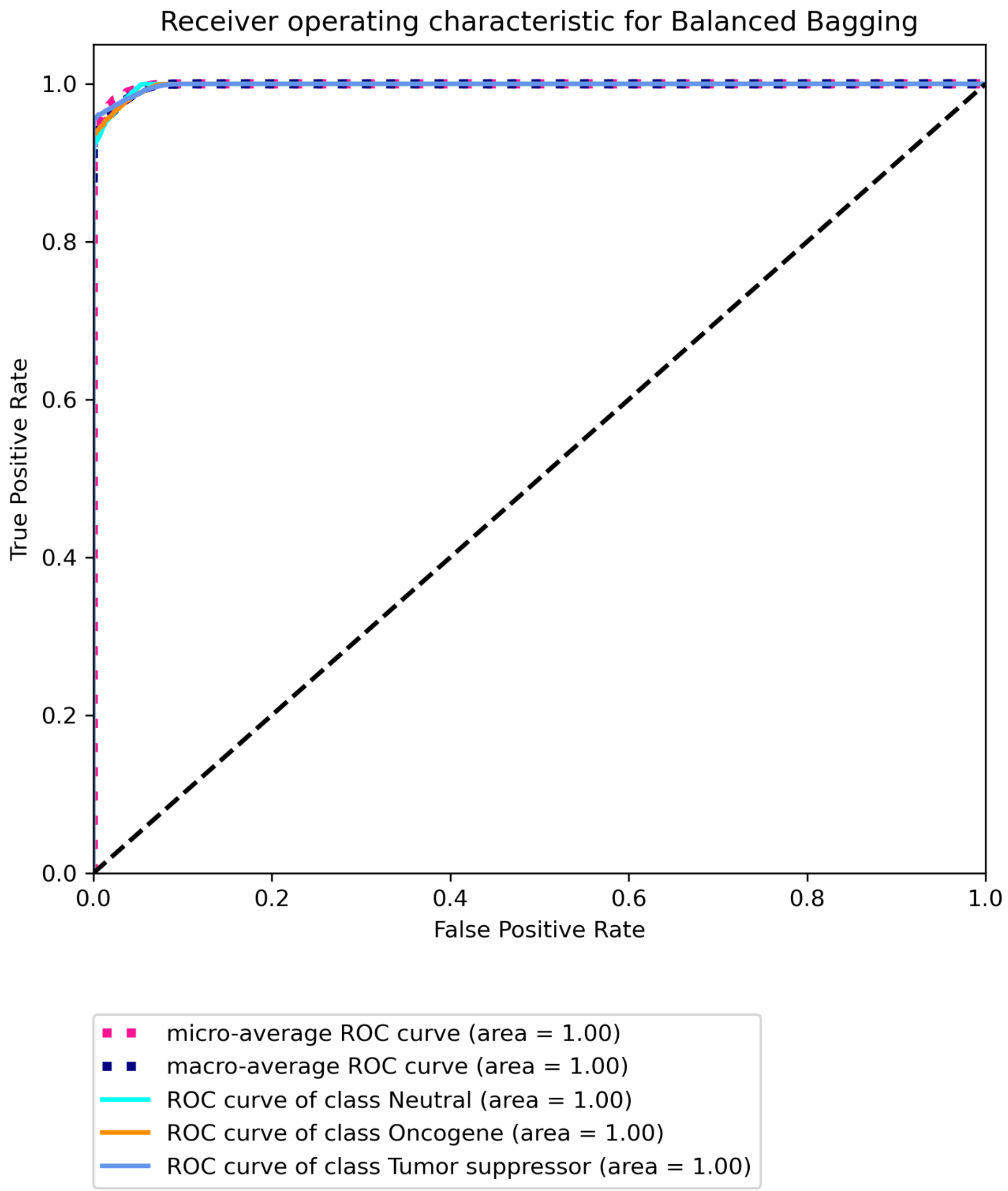


**Figure S30C. Receiver operating characteristic for BRCA using multi-omic data with CGC labels built on "some" features using Balanced Bagging.**

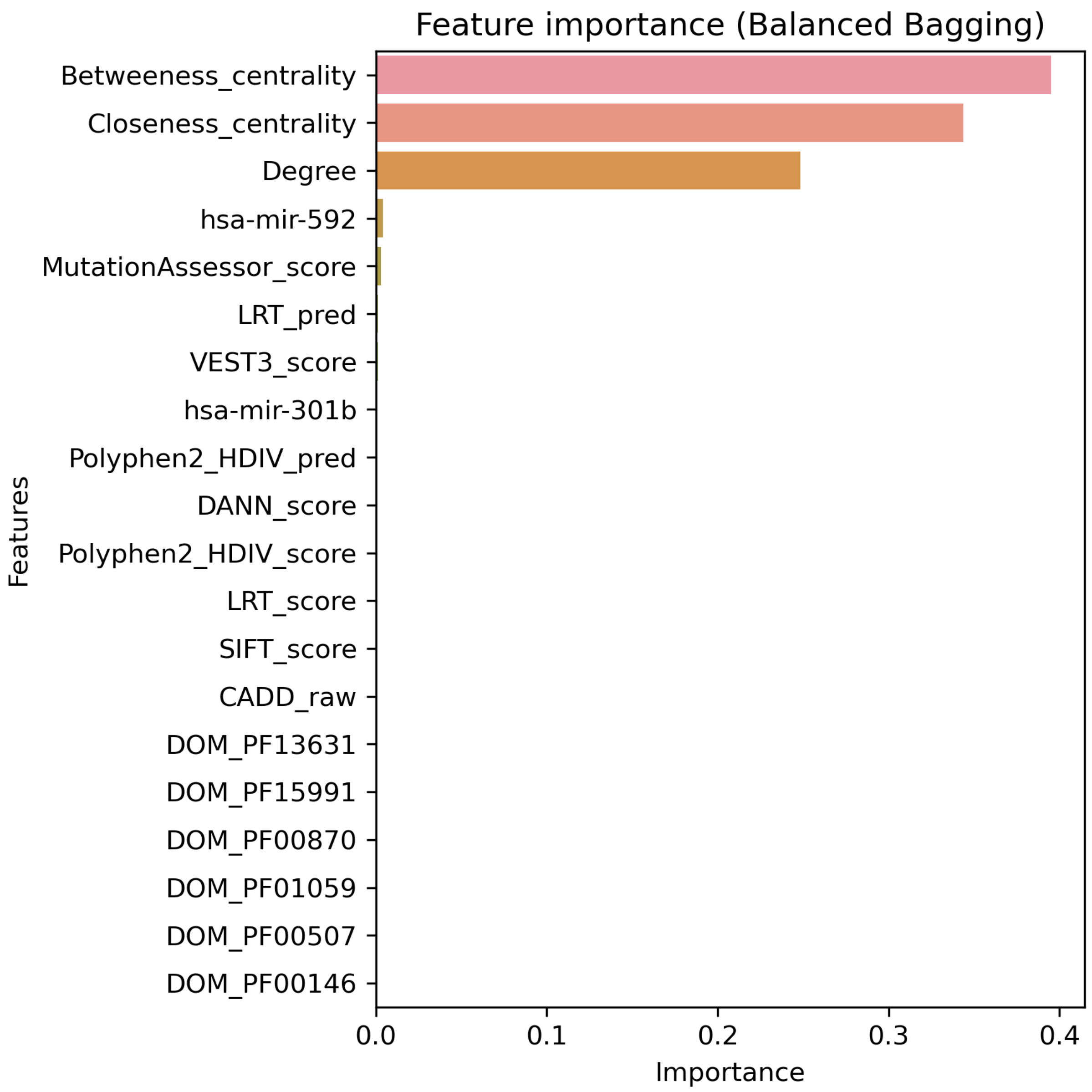


**Figure S31A. Top 20 features for BRCA using multi-omic data with CIViC labels built on "all" features using Balanced Bagging.**

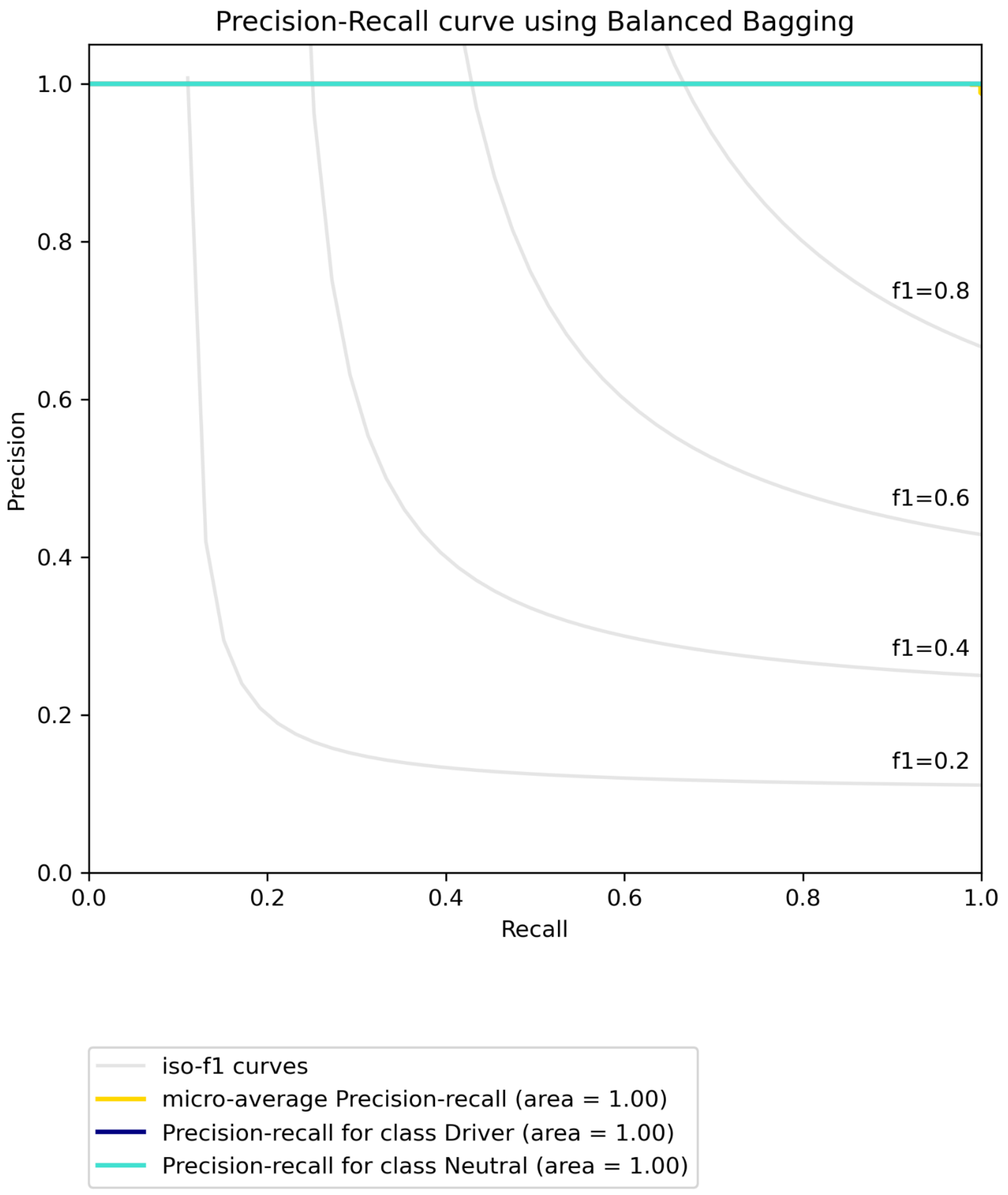


**Figure S31B. Precision-recall curve for BRCA using multi-omic data with CIViC labels built on “all” features using Balanced Bagging.**

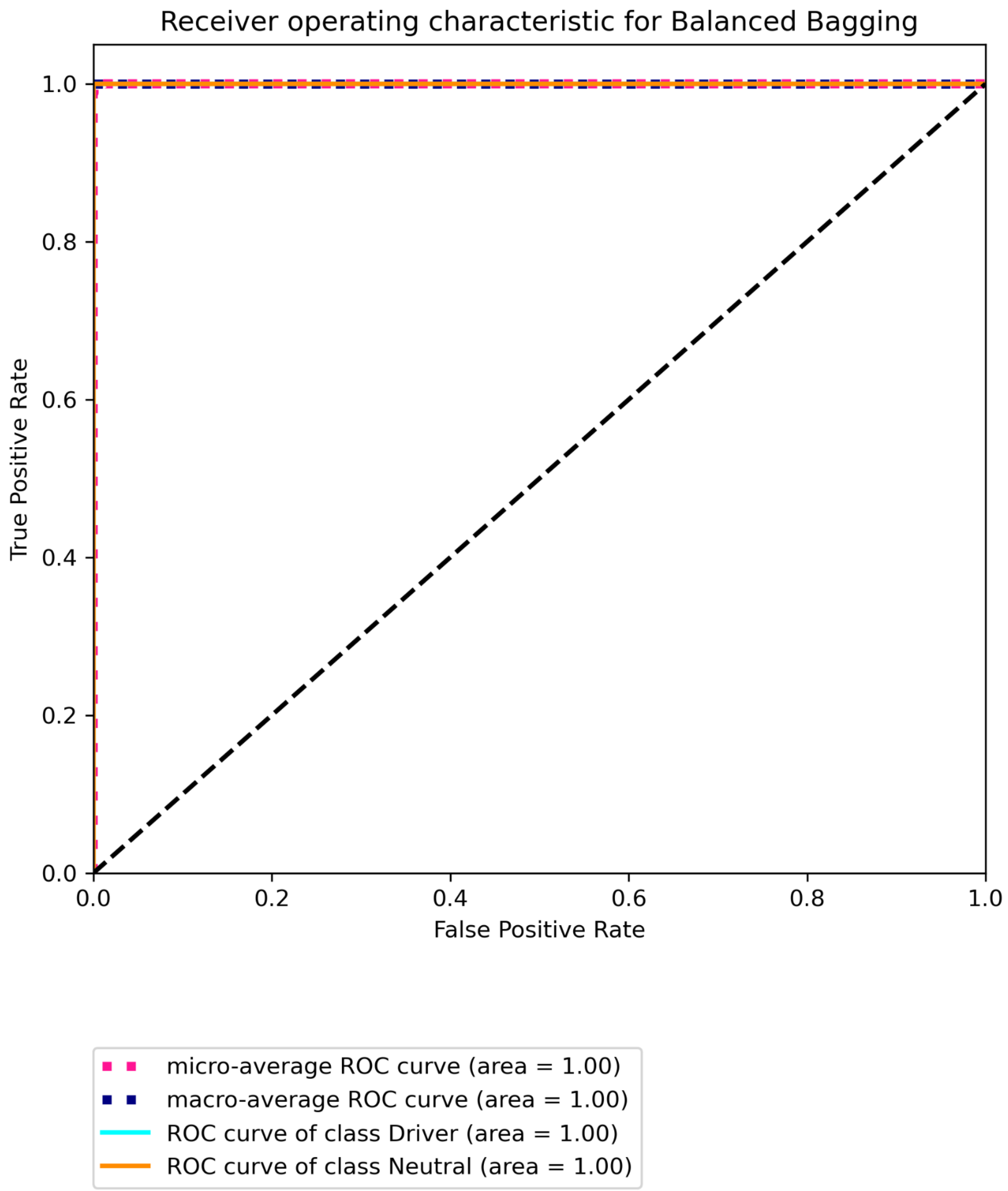


**Figure S31C. Receiver operating characteristic for BRCA using multi-omic data with CIViC labels built on "all" features using Balanced Bagging.**

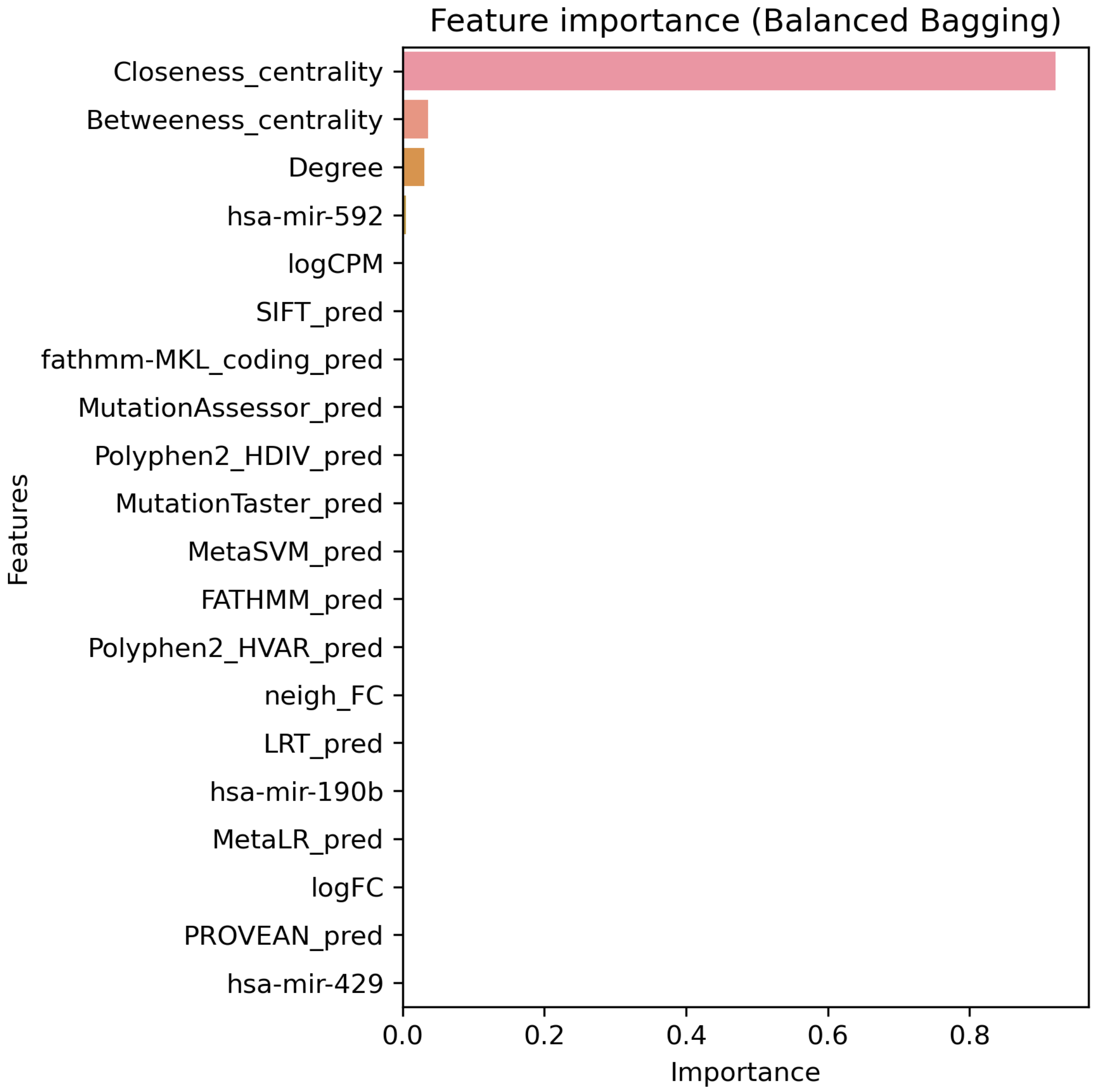


**Figure S32A. Top 20 features for BRCA using multi-omic data with CIViC labels built on “some” features using Balanced Bagging.**

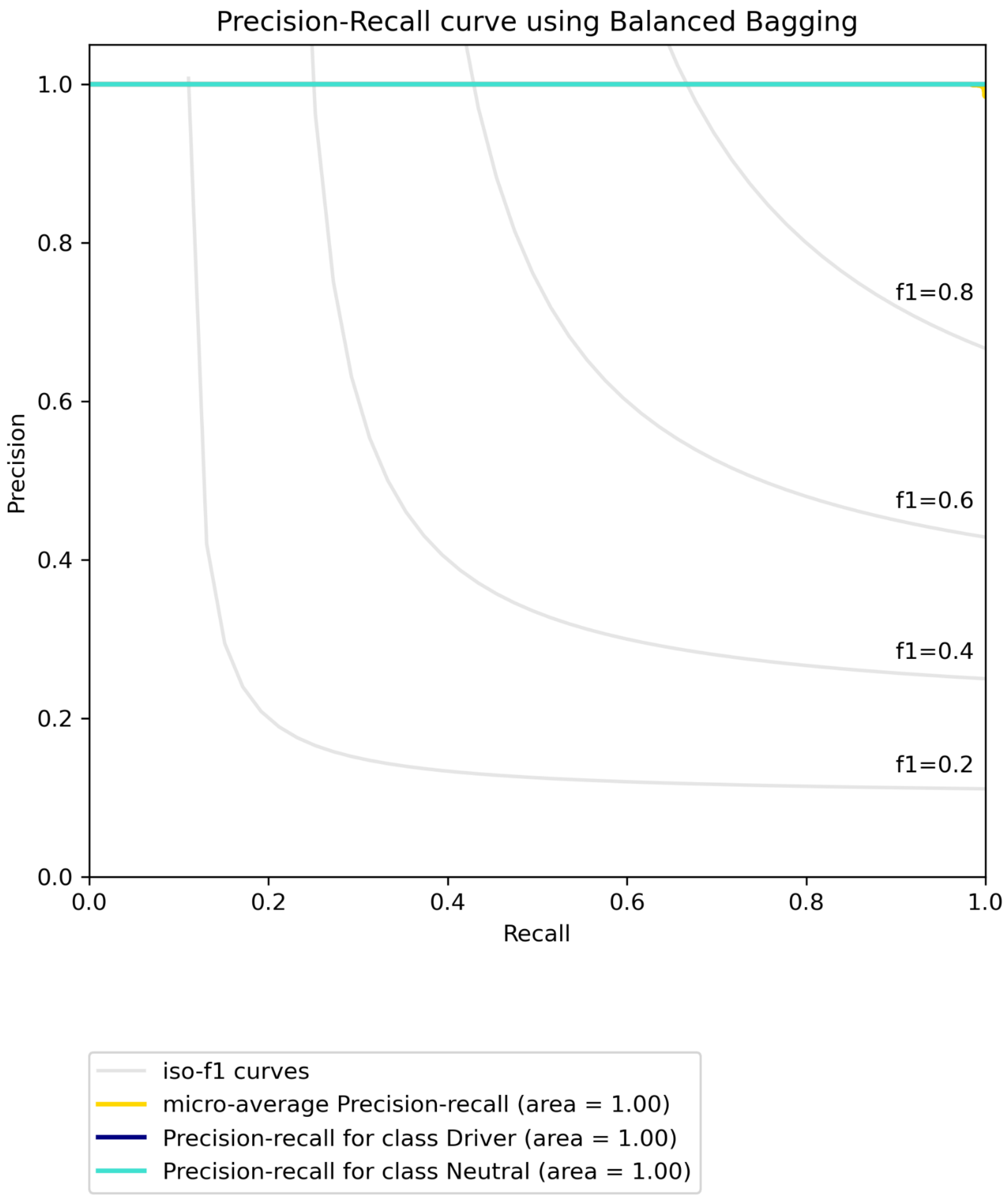


**Figure S32B. Precision-recall curve for BRCA using multi-omic data with CIViC labels built on "some" features using Balanced Bagging.**

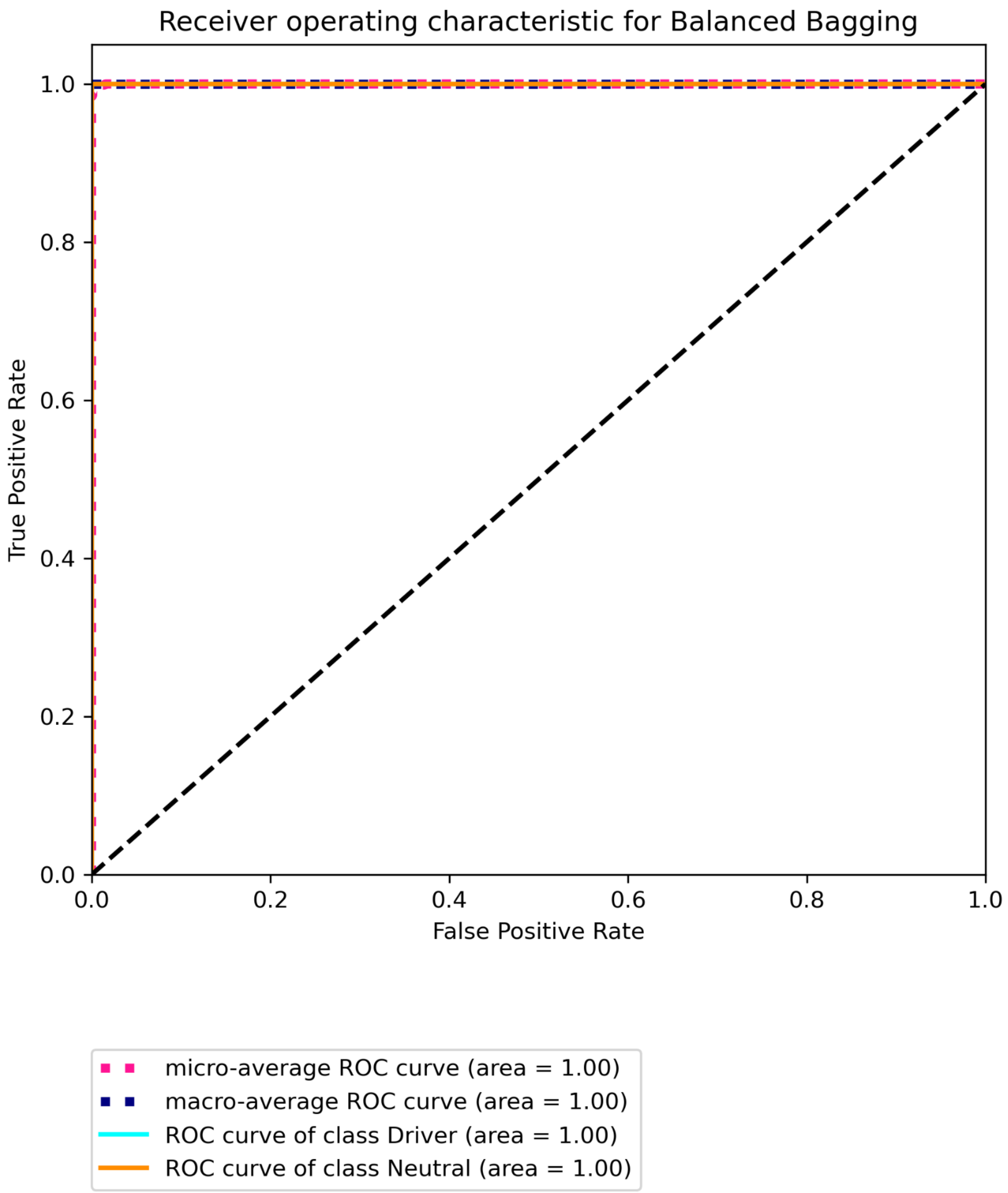


**Figure S32C. Receiver operating characteristic for BRCA using multi-omic data with CIViC labels built on "some" features using Balanced Bagging.**

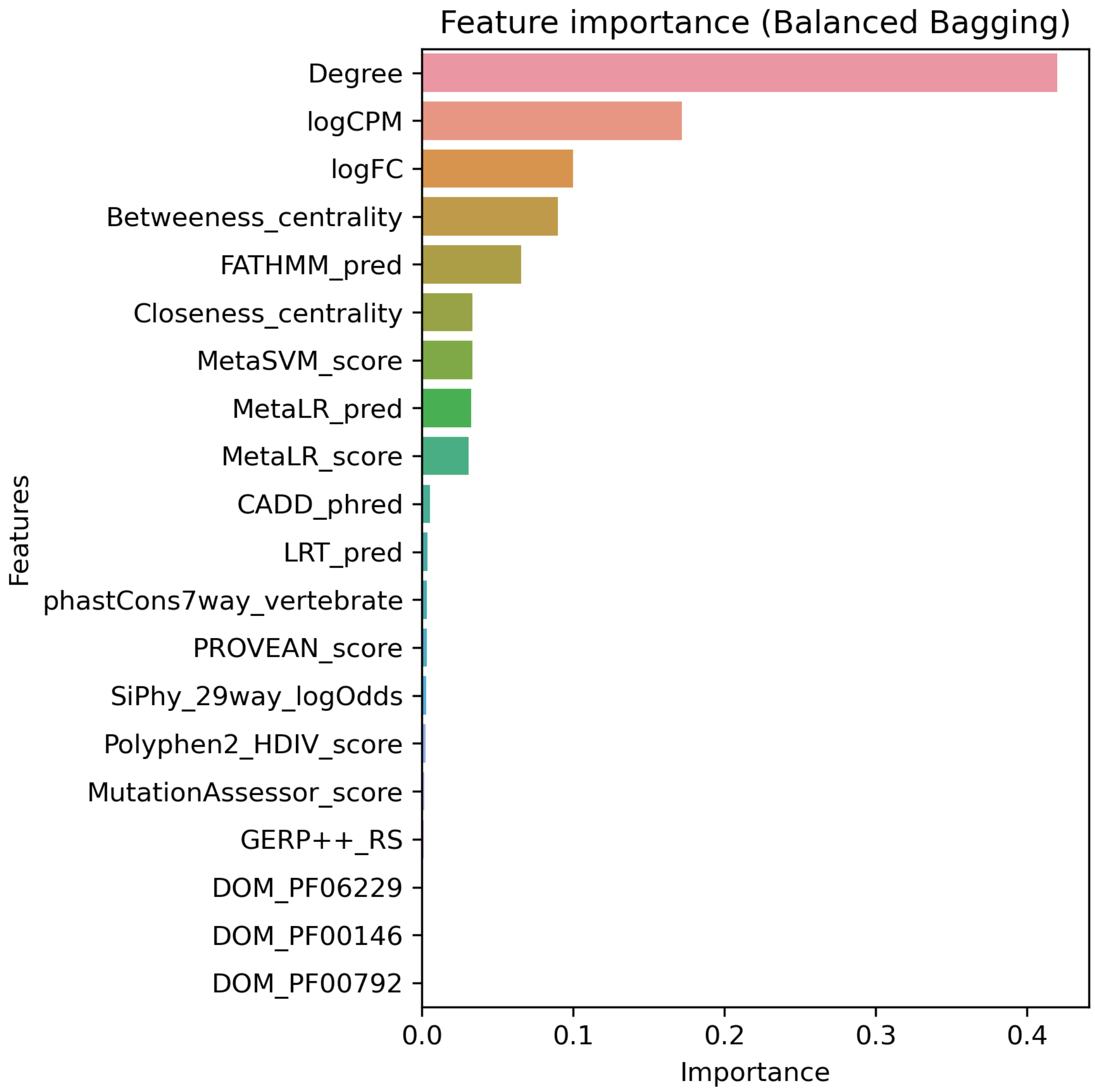


**Figure S33A. Top 20 features for BRCA using multi-omic data with Marellotto *et al.* labels built on "all" features using Balanced Bagging.**

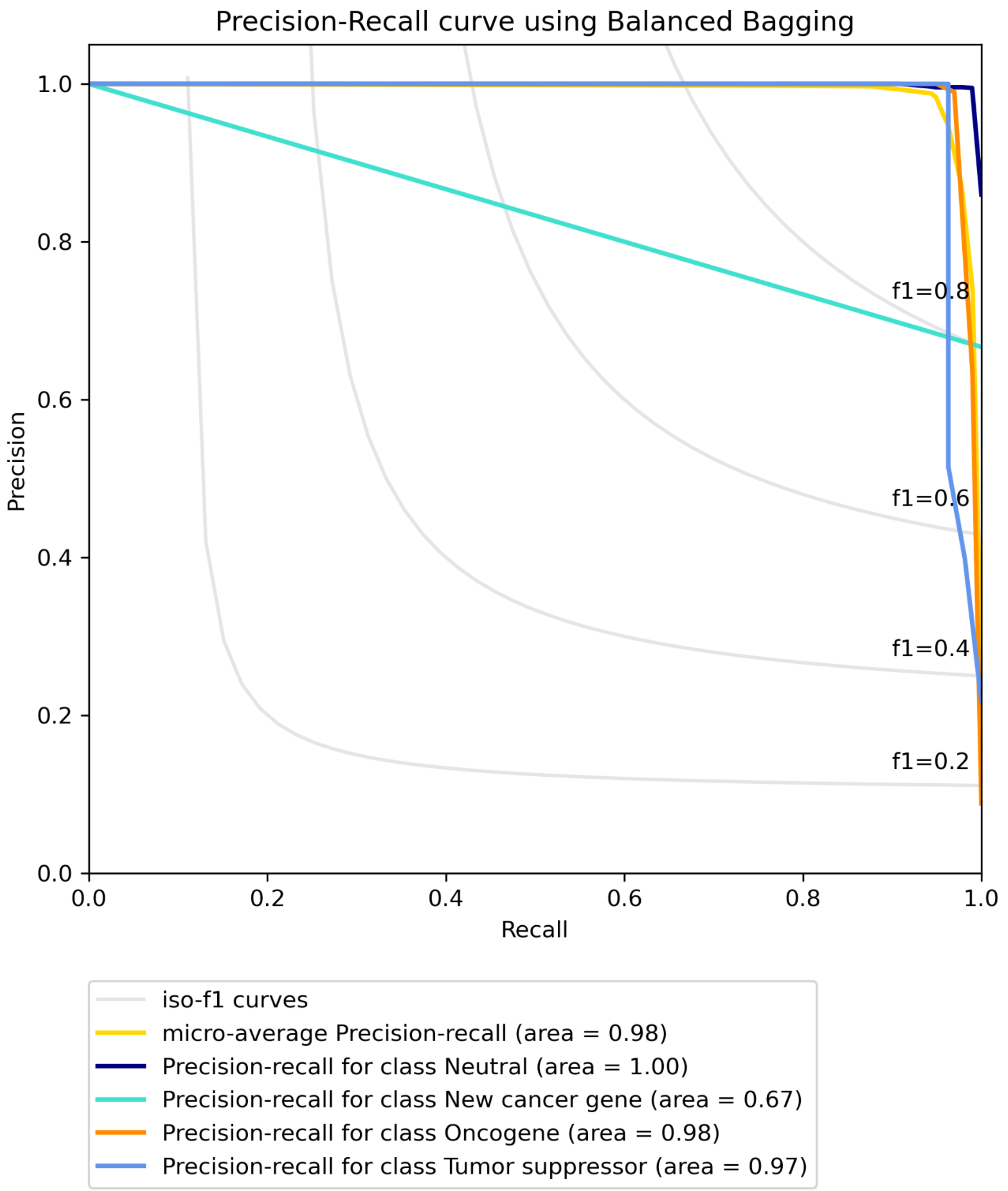


**Figure S33B. Precision-recall curve for BRCA using multi-omic data with Marellotto *et al.* labels built on "all" features using Balanced Bagging.**

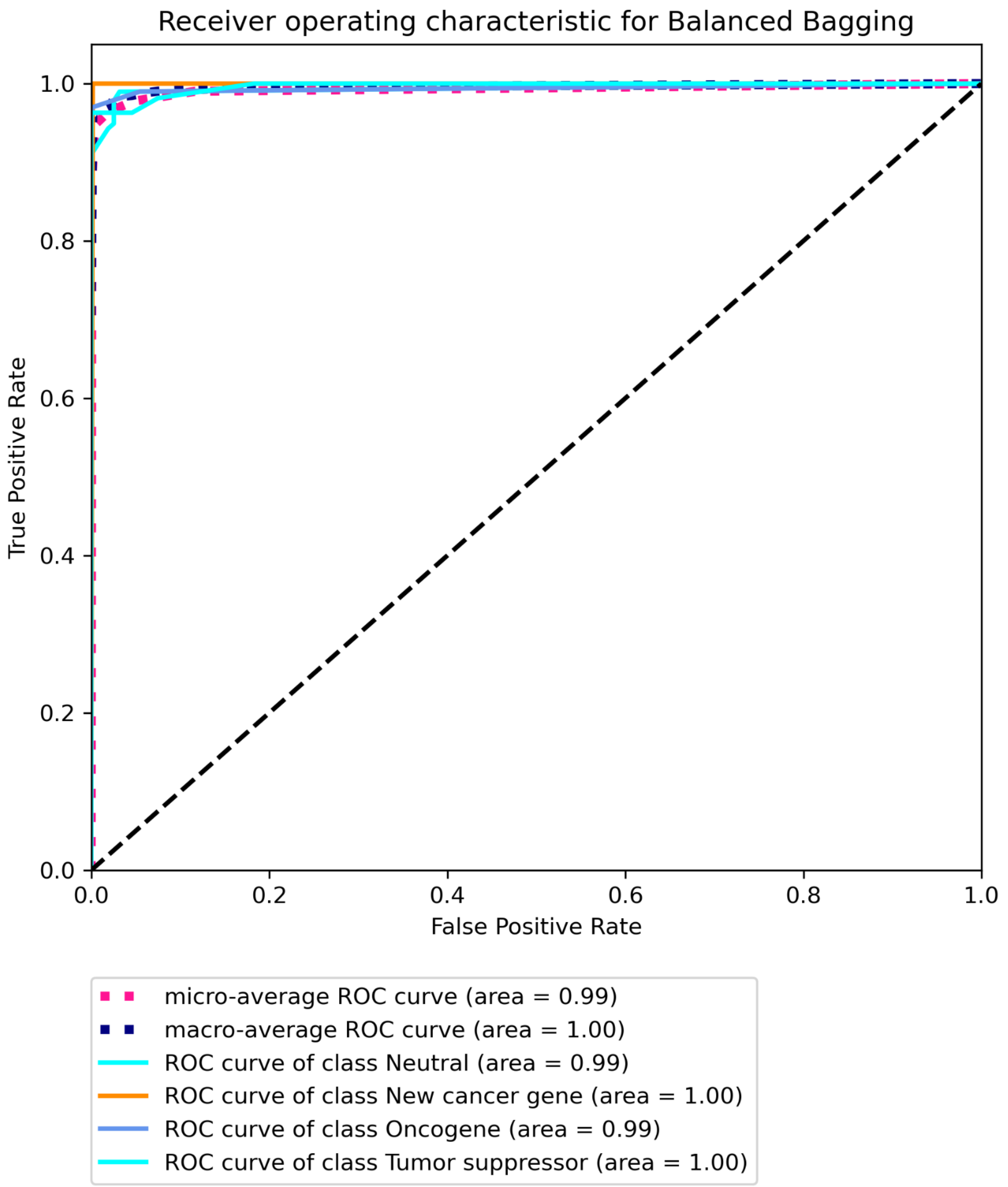


**Figure S33C. Receiver operating characteristic for BRCA using multi-omic data with Marellotto *et al.* labels built on "all" features using Balanced Bagging.**

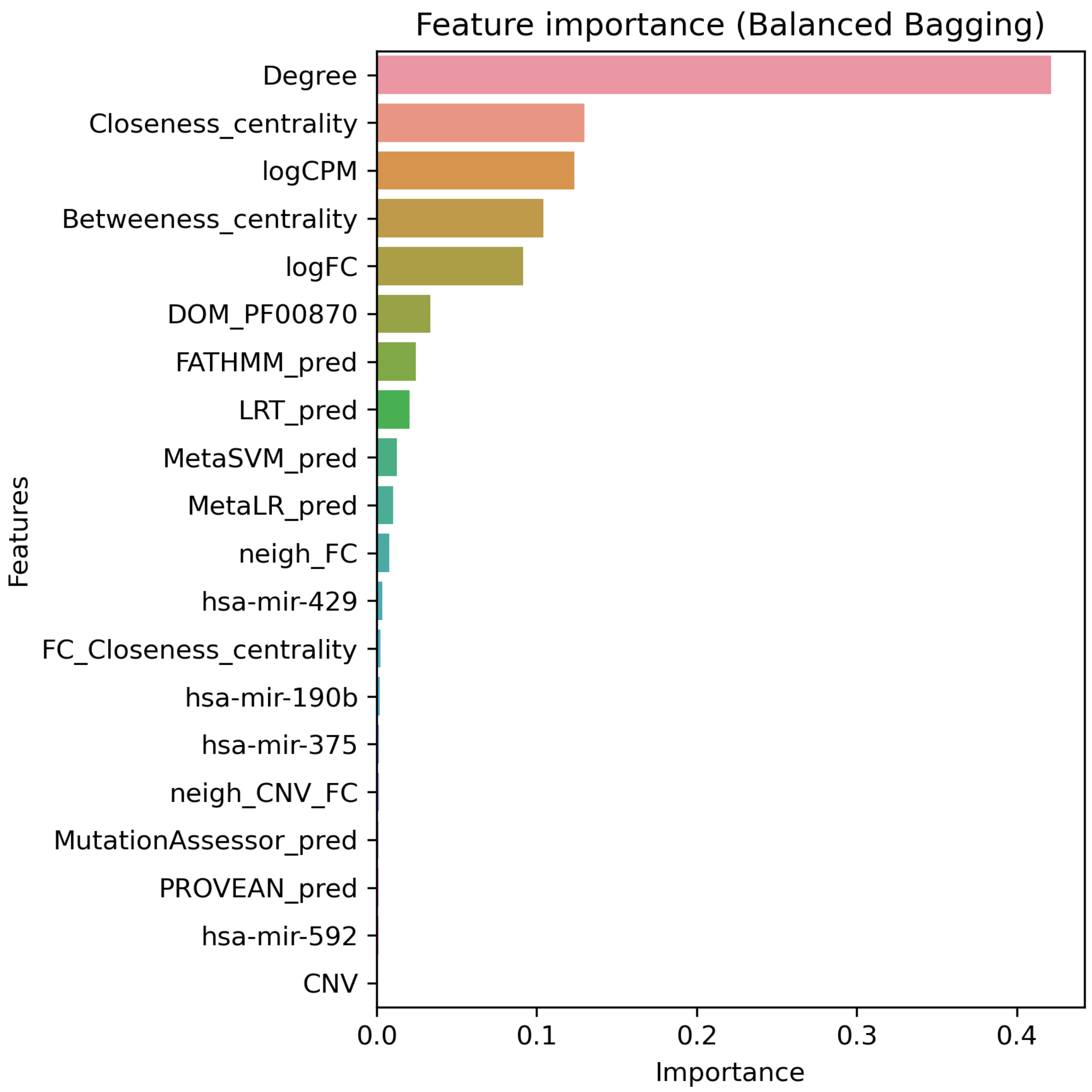


**Figure S34A. Top 20 features for BRCA using multi-omic data with Marellotto *et al.* labels built on "some" features using Balanced Bagging.**

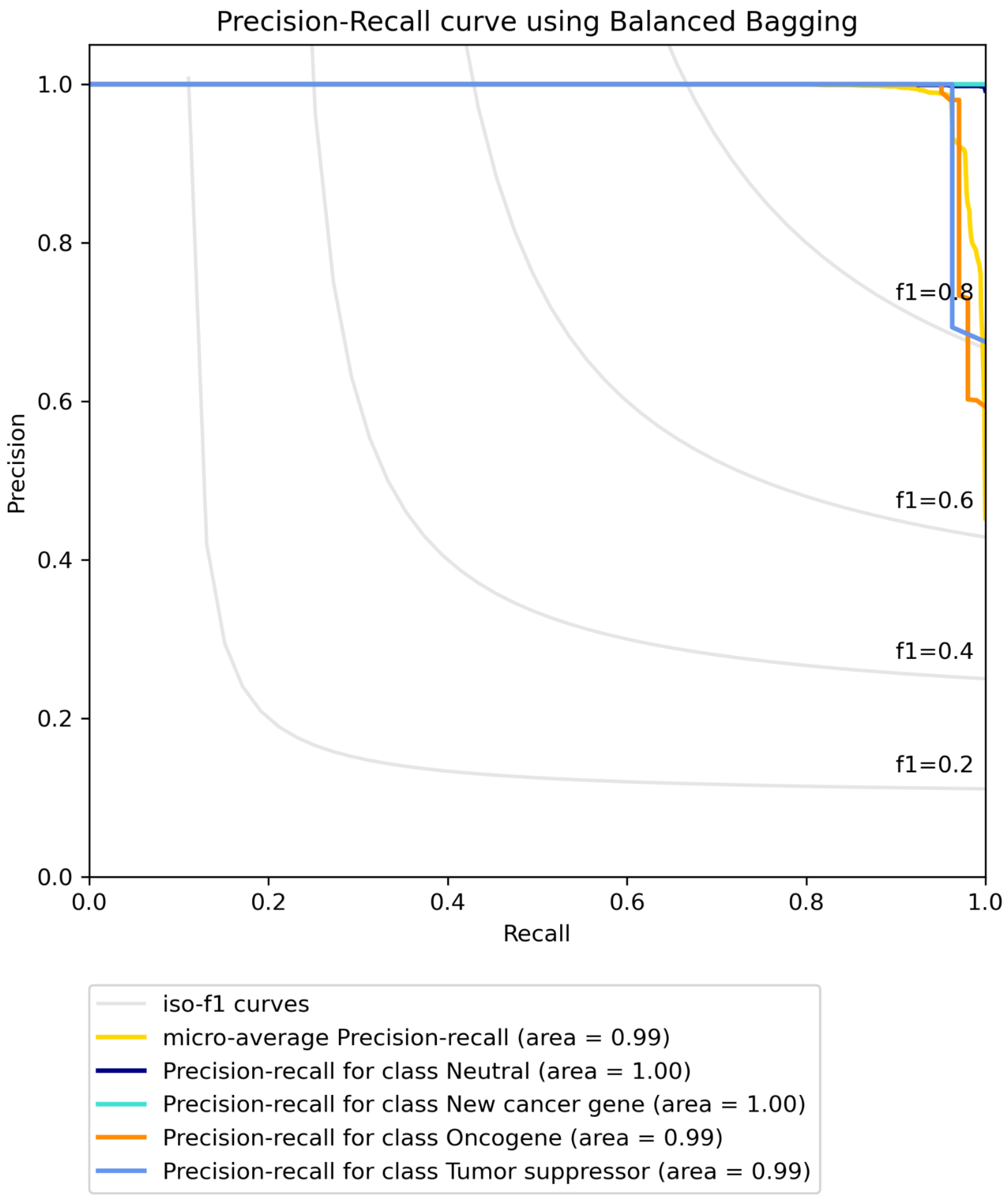


**Figure S34B. Precision-recall curve for BRCA using multi-omic data with Marellotto *et al.* labels built on "some" features using Balanced Bagging.**

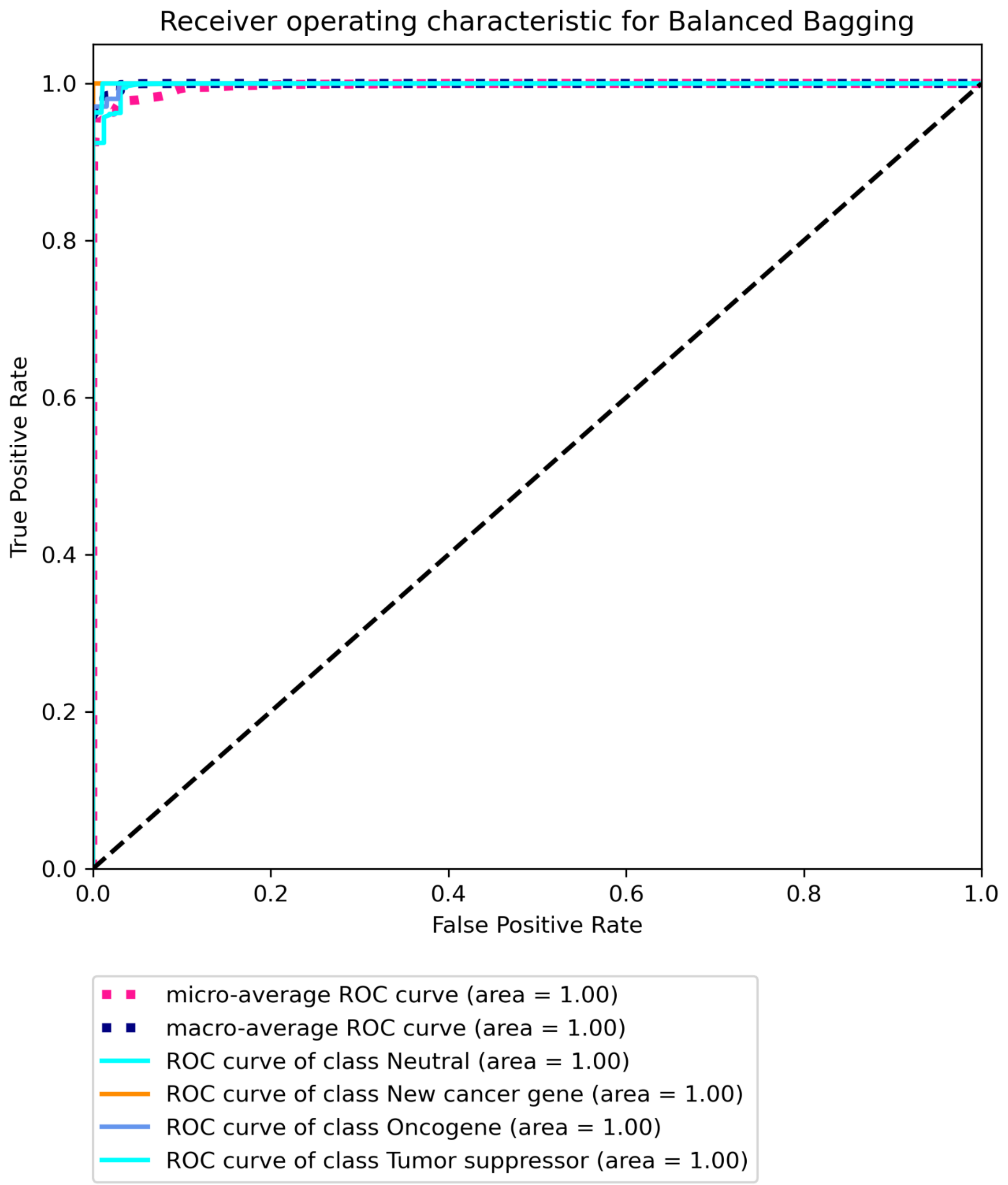


**Figure S34C. Receiver operating characteristic for BRCA using multi-omic data with Marellotto *et al.* labels built on "some" features using Balanced Bagging.**

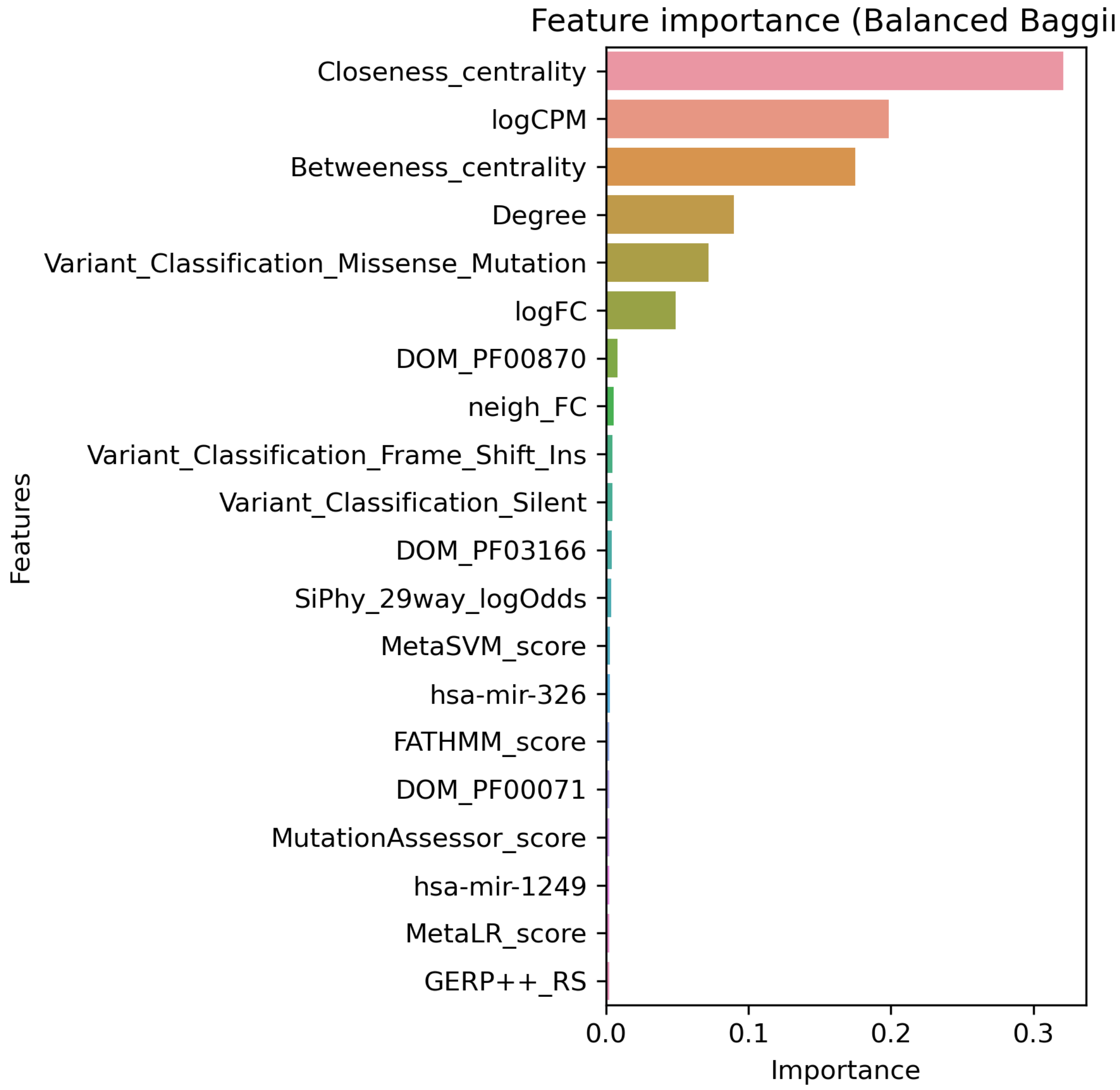


**Figure S35A. Top 20 features for COAD using multi-omic data with Bailey *et al.* labels built on "all" features using Balanced bagging.**

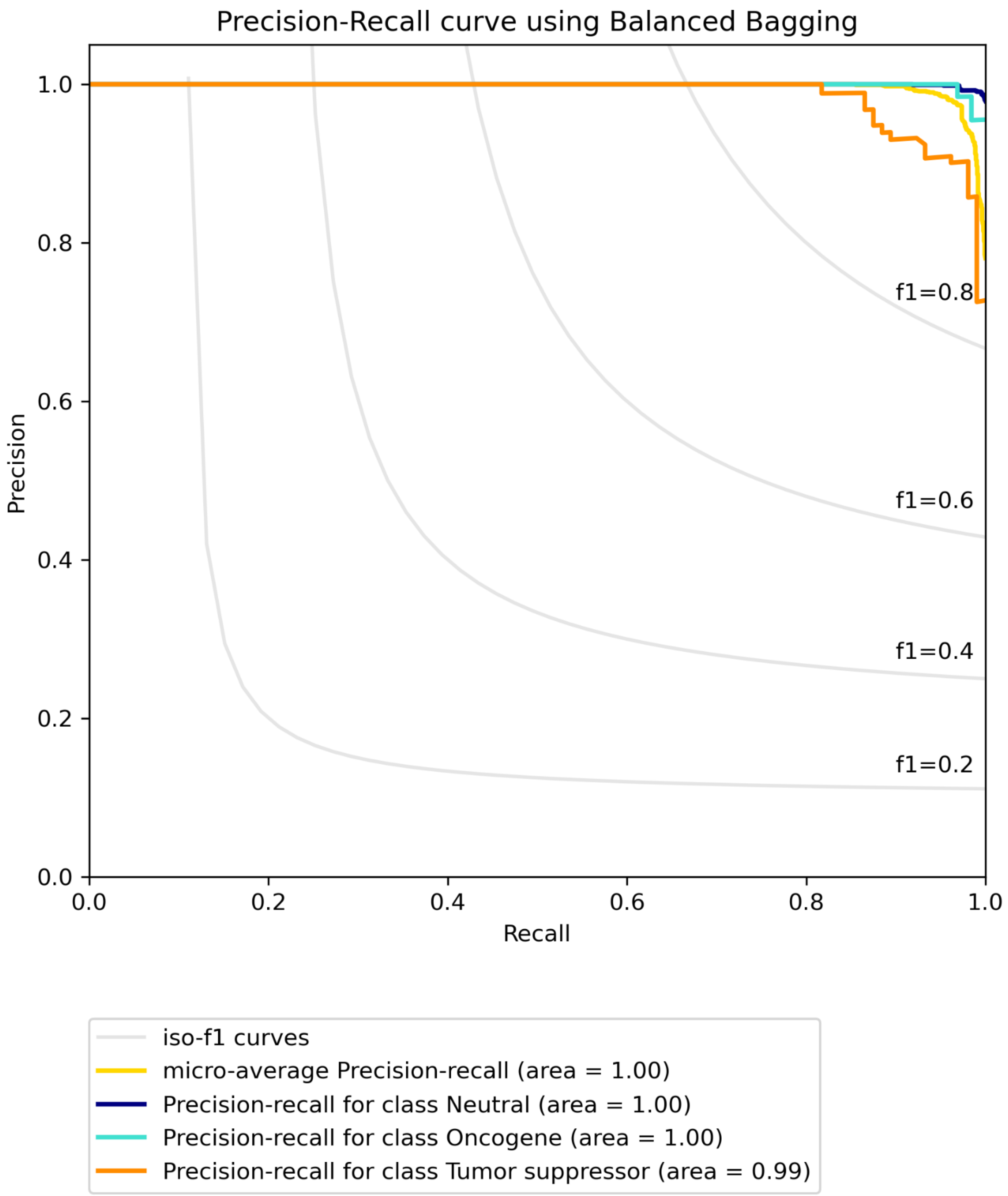


**Figure S35B. Precision-recall curve for COAD using multi-omic data with Bailey *et al.* labels built on "all" features using Balanced bagging.**

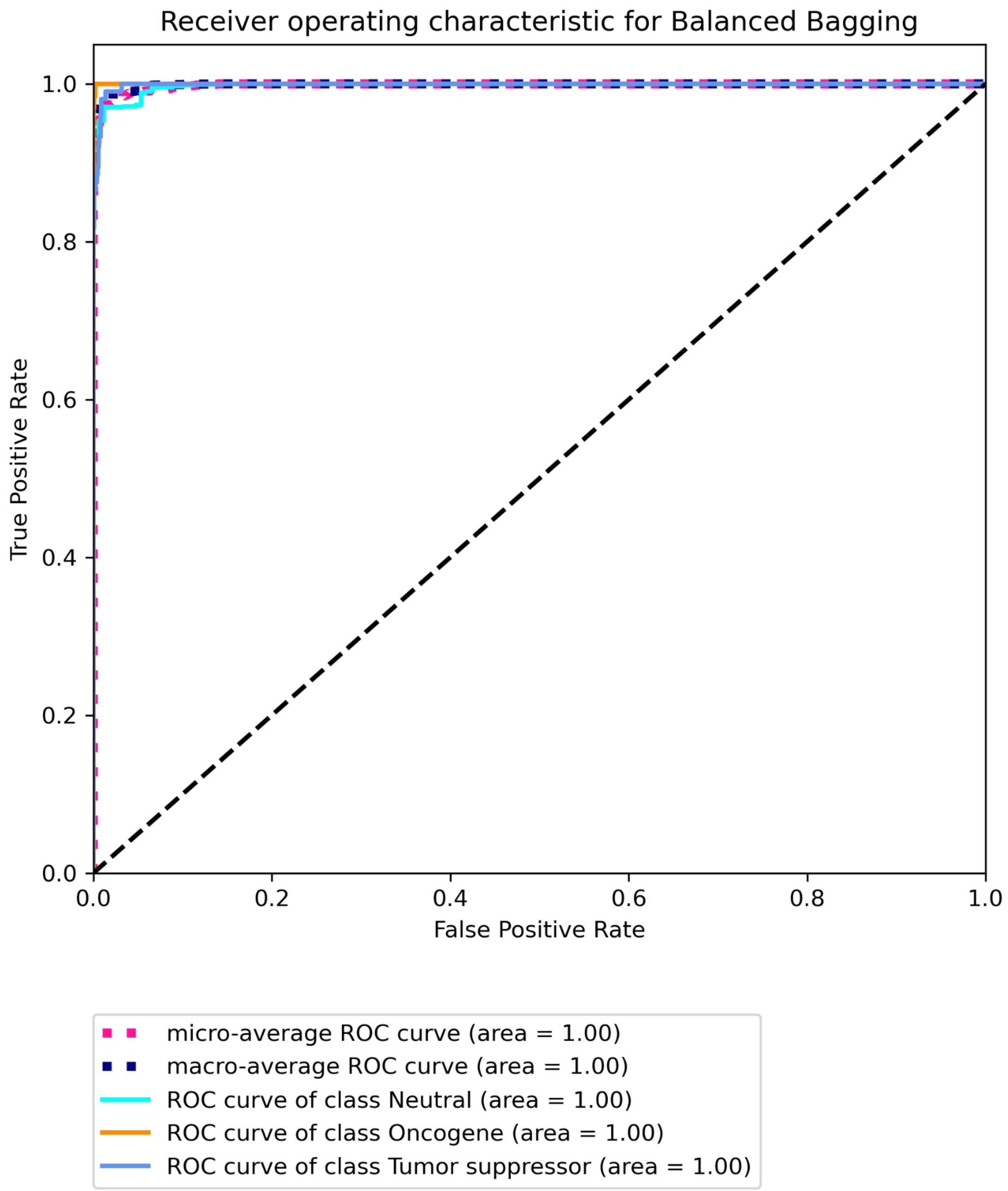


**Figure S35C Receiver operating characteristic for COAD using multi-omic data with Bailey *et al.* labels built on "all" features using Balanced bagging.**

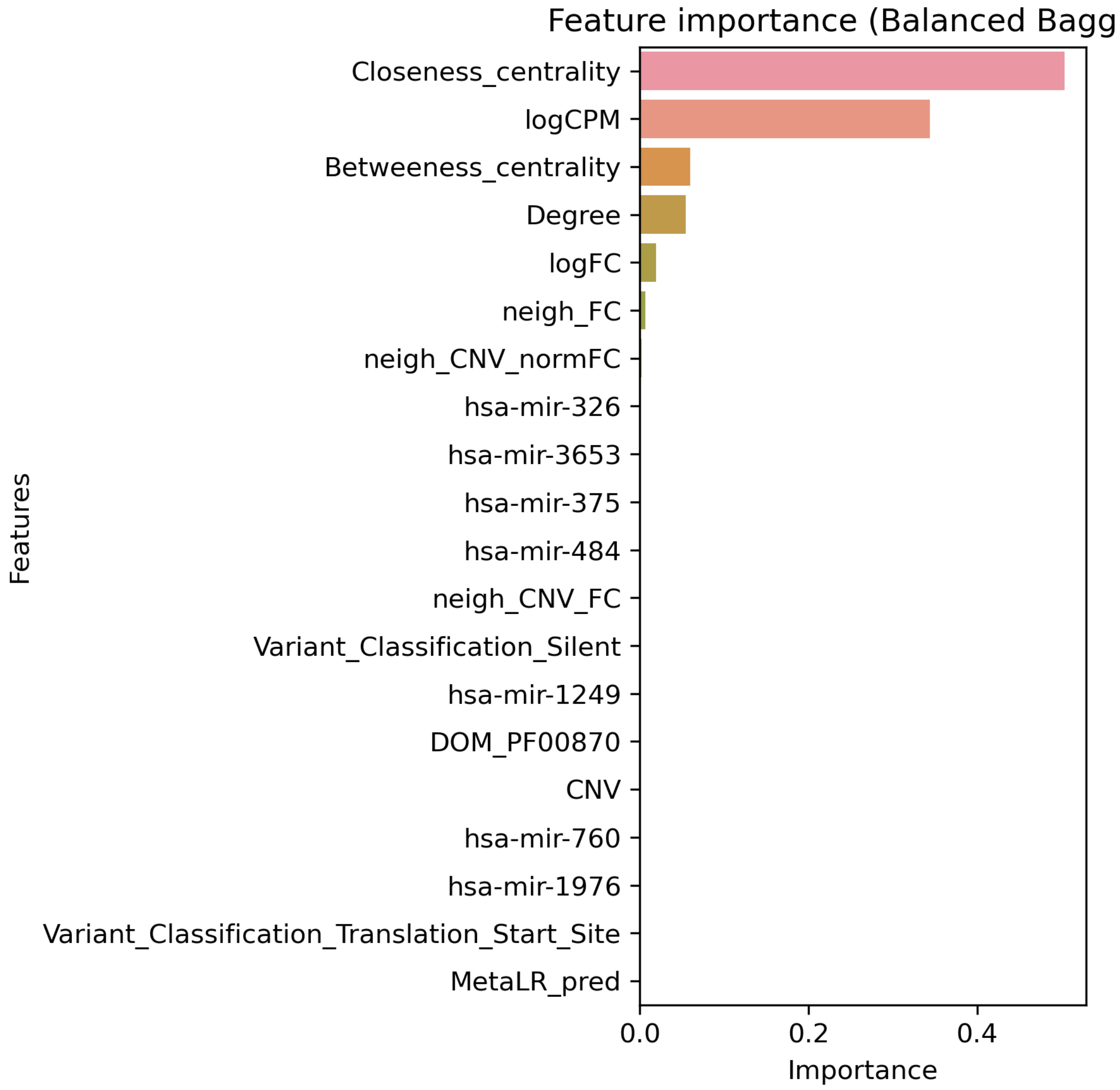


**Figure S36A. Top 20 features for COAD using multi-omic data with Bailey *et al.* labels built on "some" features using Balanced bagging.**

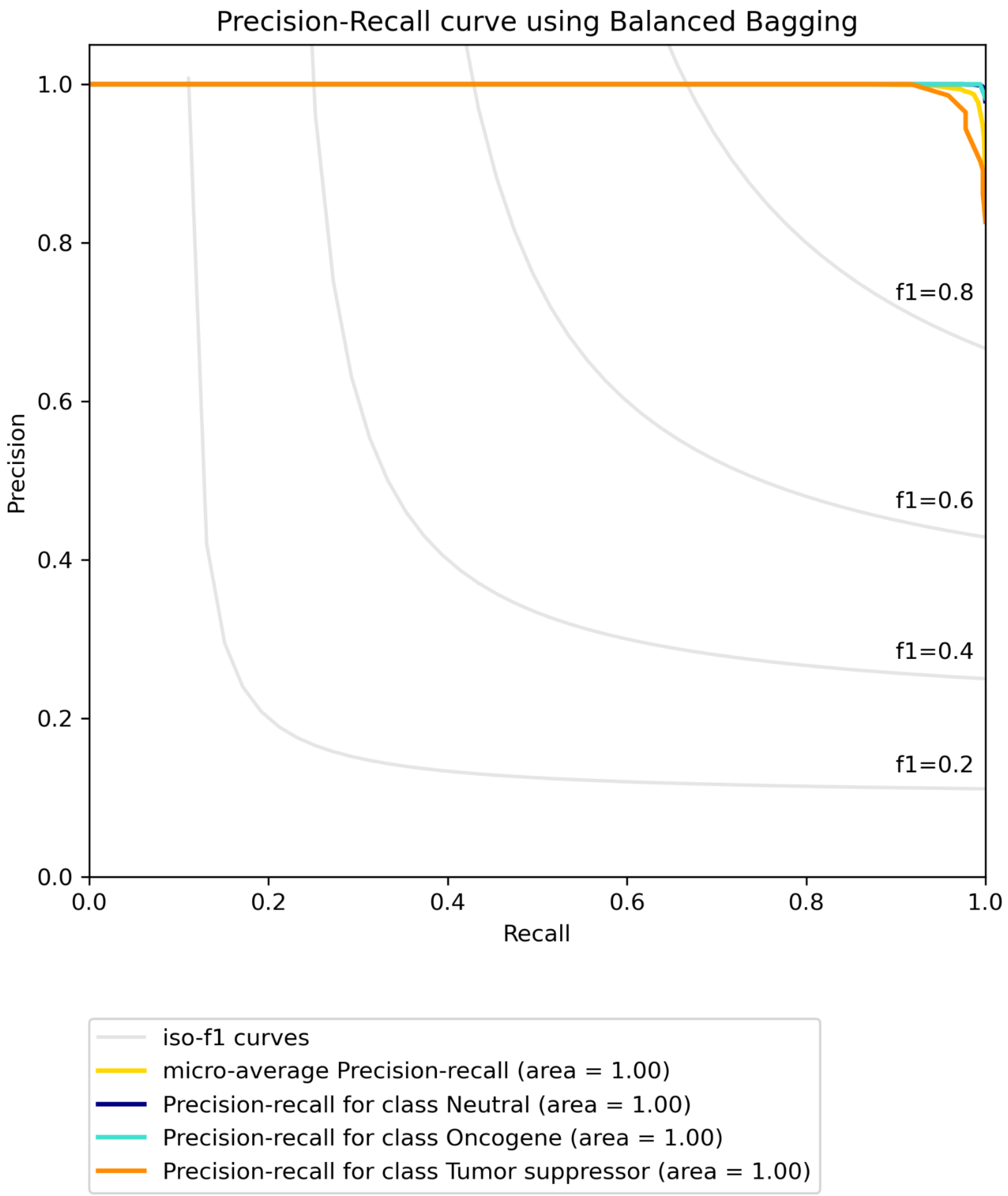


**Figure S36B. Precision-recall curve for COAD using multi-omic data with Bailey *et al.* labels built on "some" features using Balanced bagging.**

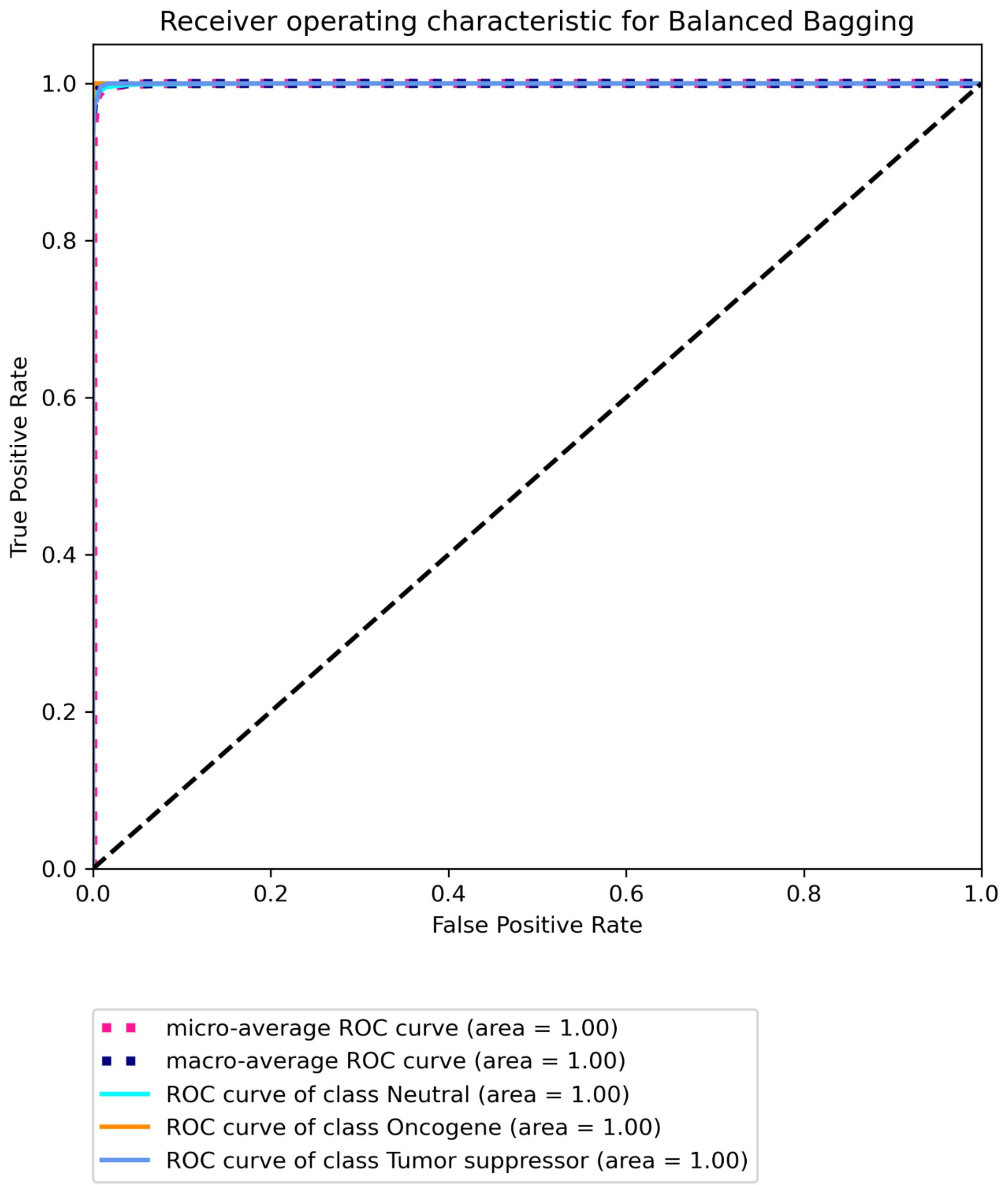


**Figure S36C Receiver operating characteristic for COAD using multi-omic data with Bailey *et al.* labels built on "some" features using Balanced bagging.**

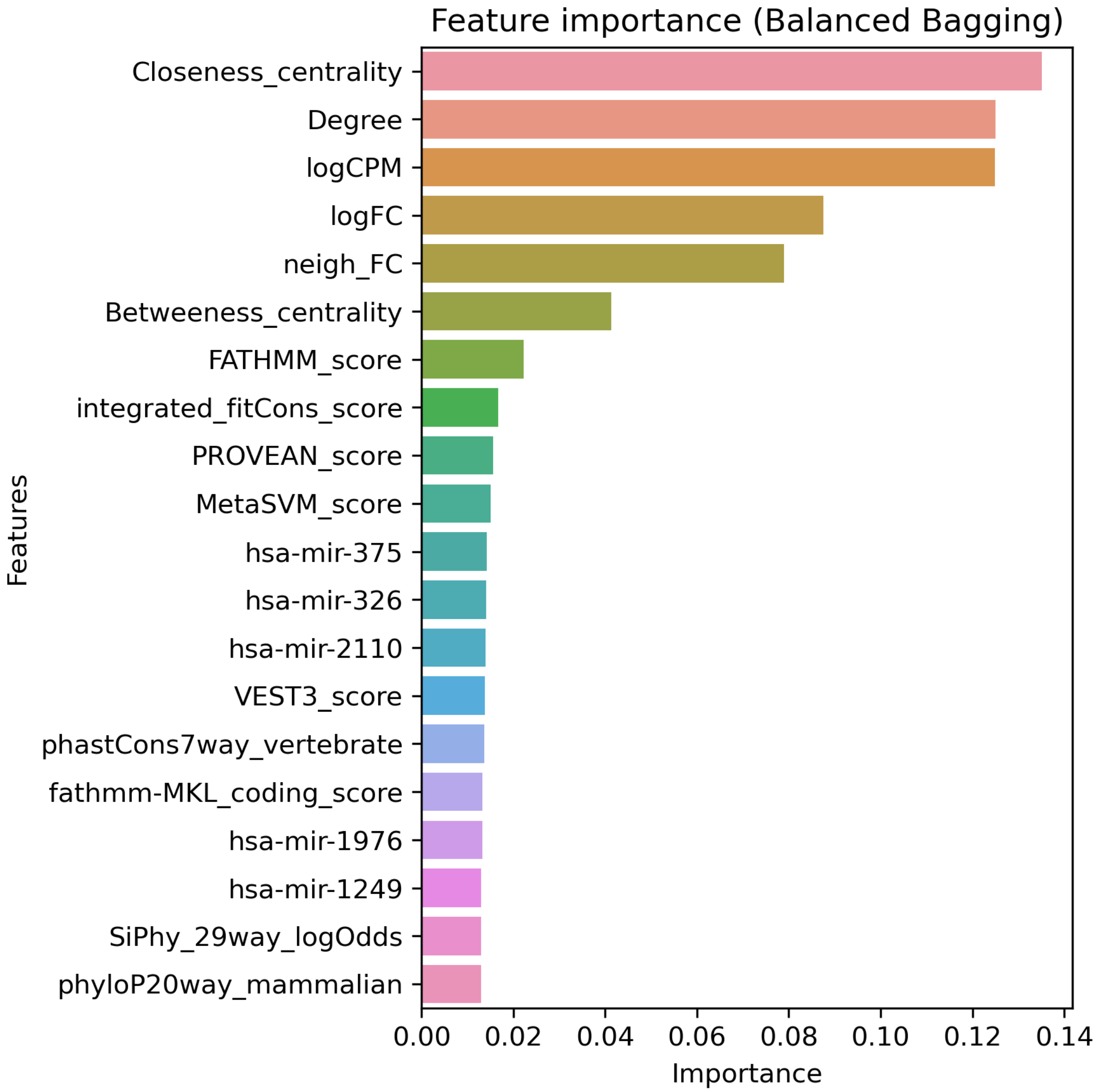


**Figure S37A. Top 20 features for COAD using multi-omic data with CGC labels built on “all” features using Balanced bagging.**

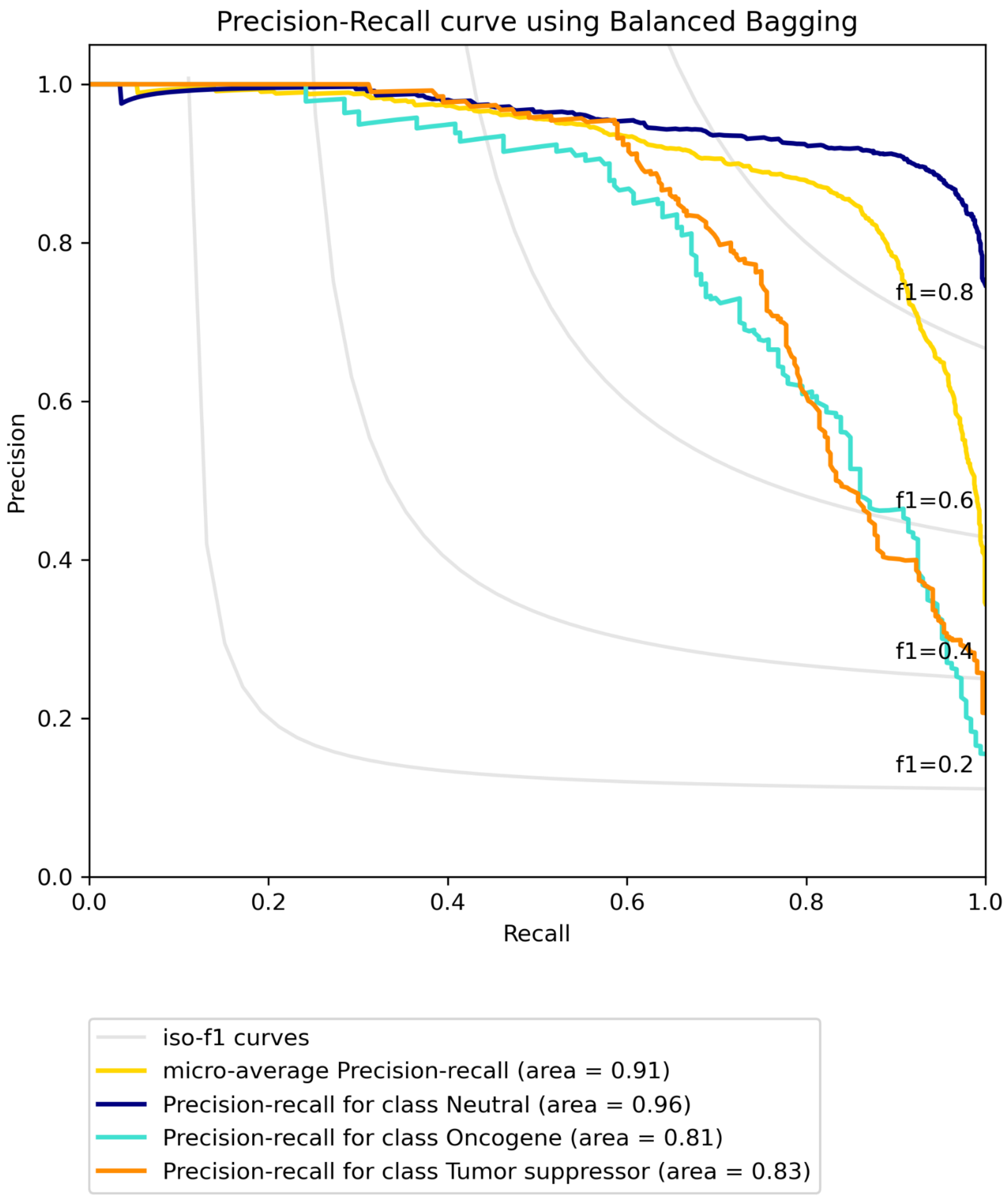


**Figure S37B. Precision-recall curve for COAD using multi-omic data with CGC labels built on "all" features using Balanced bagging.**

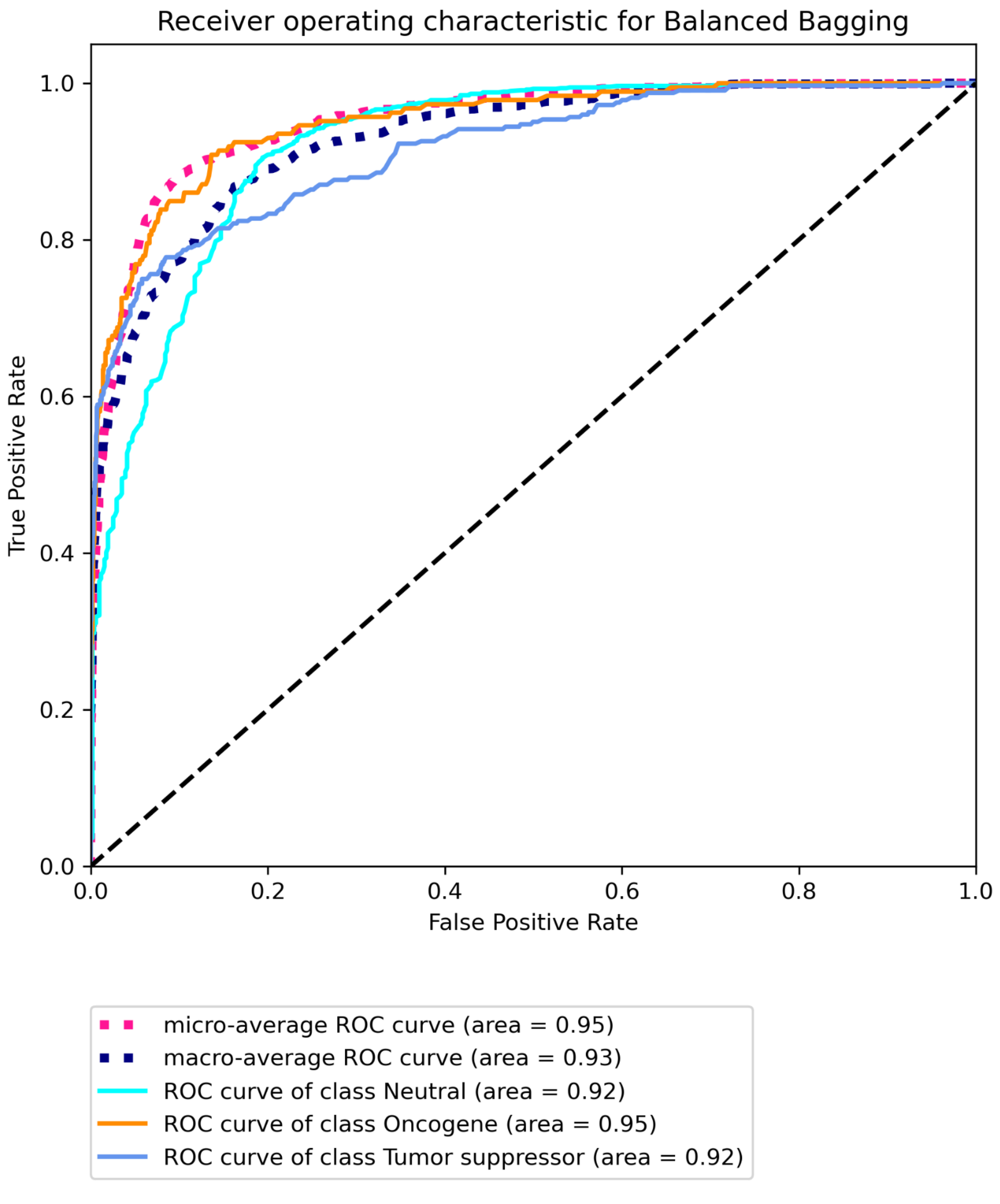


**Figure S37C. Receiver operating characteristic for COAD using multi-omic data with CGC labels built on "all" features using Balanced Bagging.**

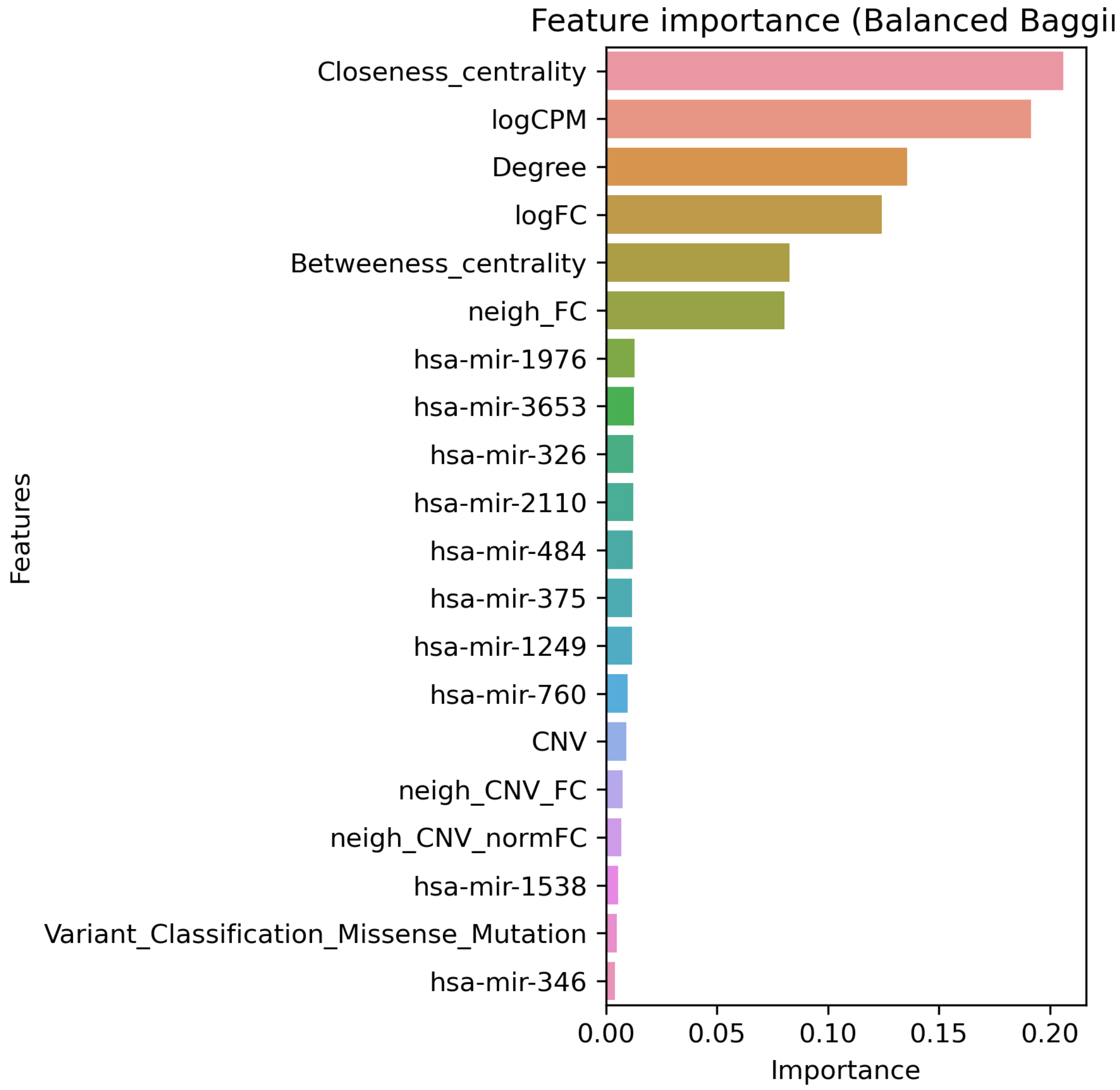


**Figure S38A. Top 20 features for COAD using multi-omic data with CGC labels built on "some" features using Balanced Bagging.**

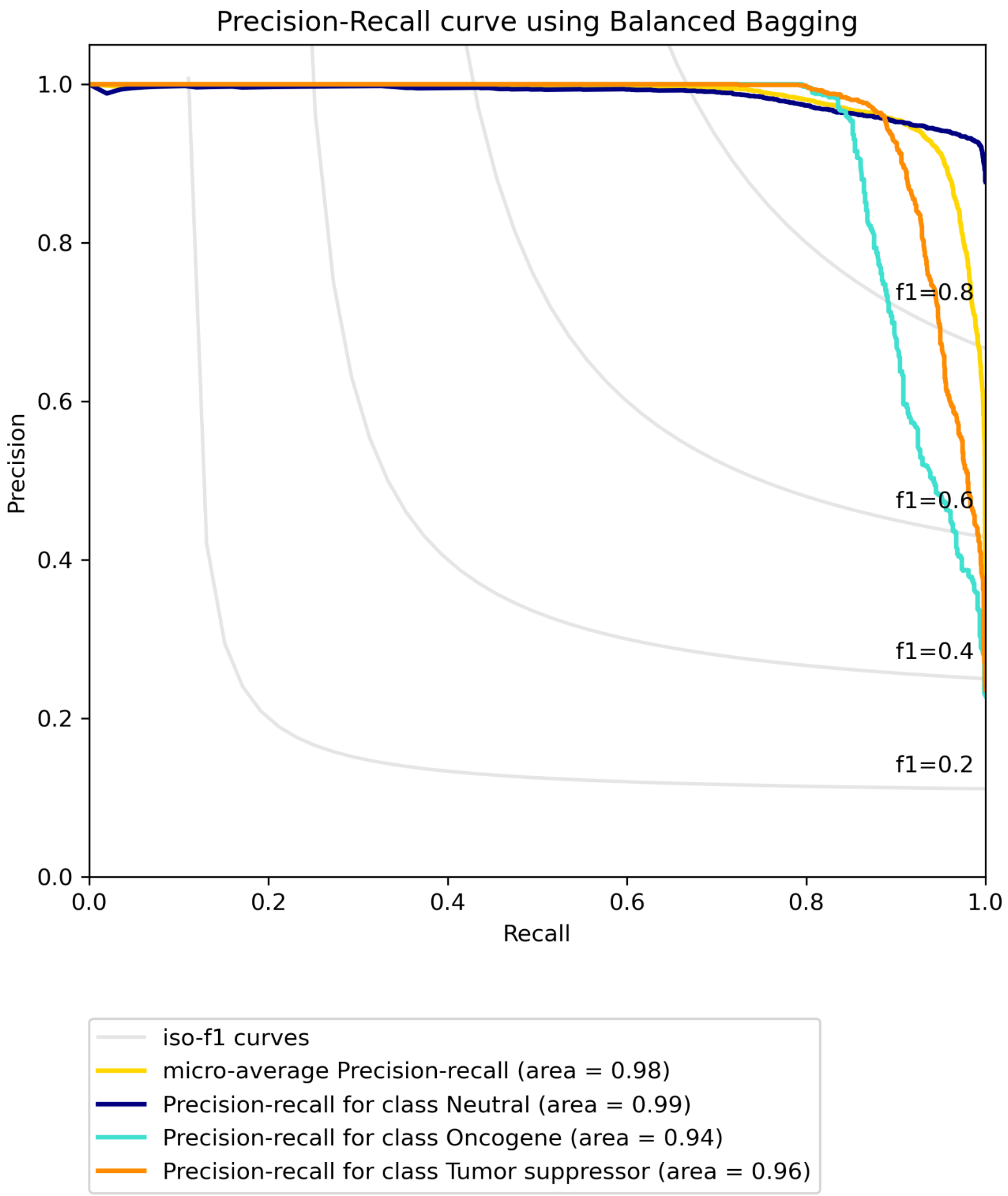


**Figure S38B. Precision-recall curve for COAD using multi-omic data with CGC labels built on “some” features using Balanced Bagging.**

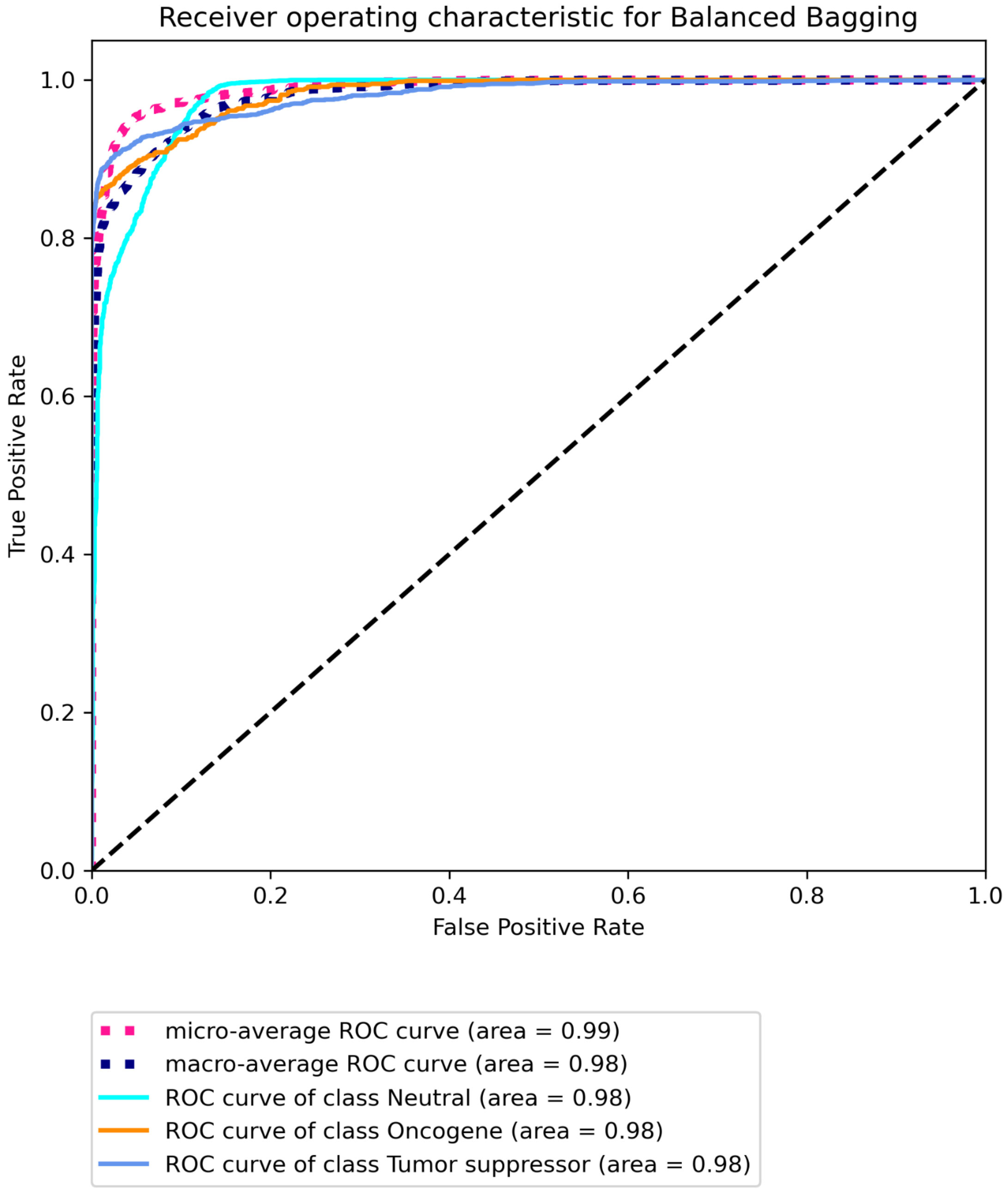


**Figure S38C. Receiver operating characteristic for COAD using multi-omic data with CGC labels built on "some" features using Balanced Bagging.**

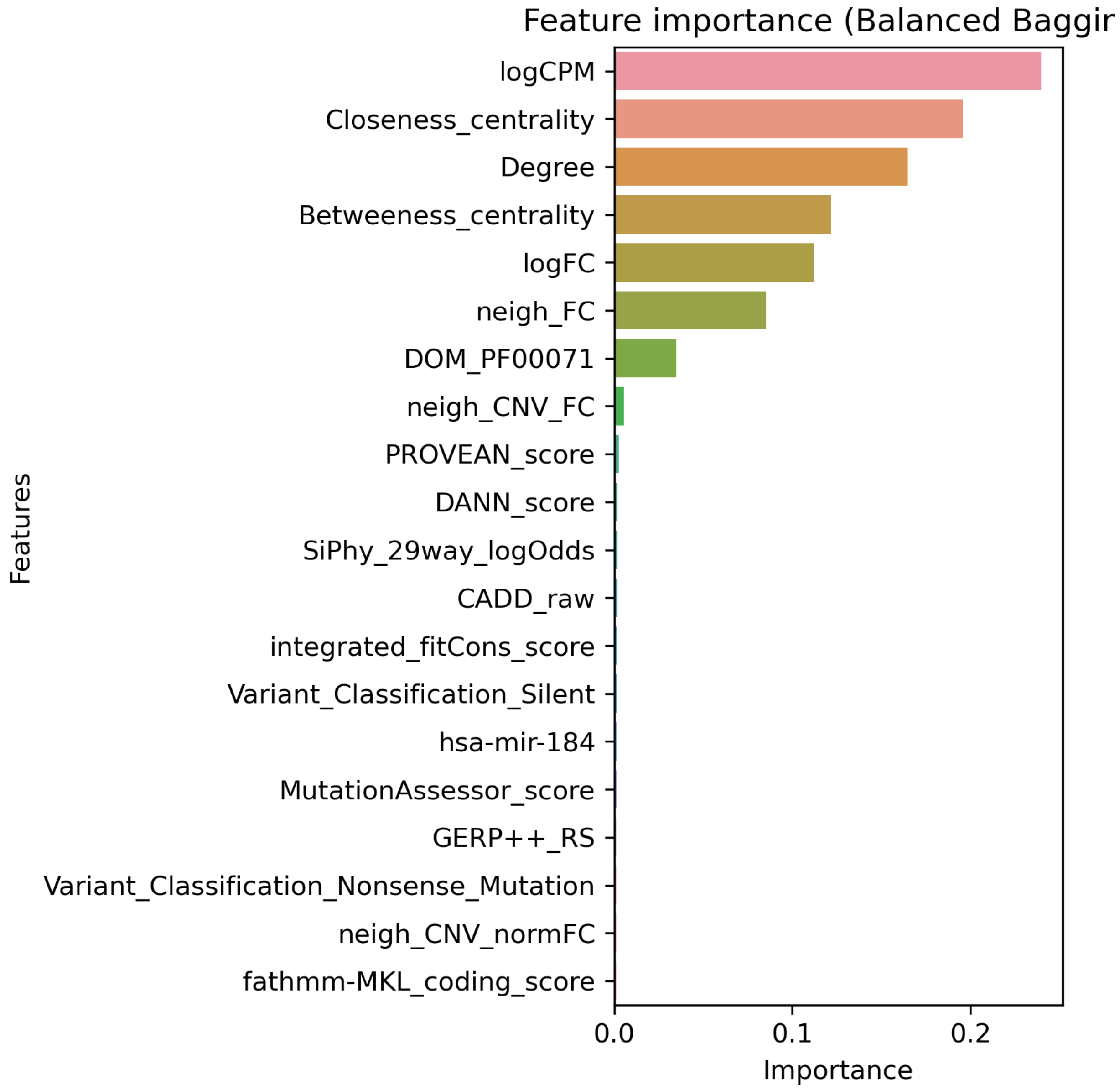


**Figure S39A. Top 20 features for LUAD using multi-omic data with Bailey *et al.* labels built on "all" features using Balanced bagging.**

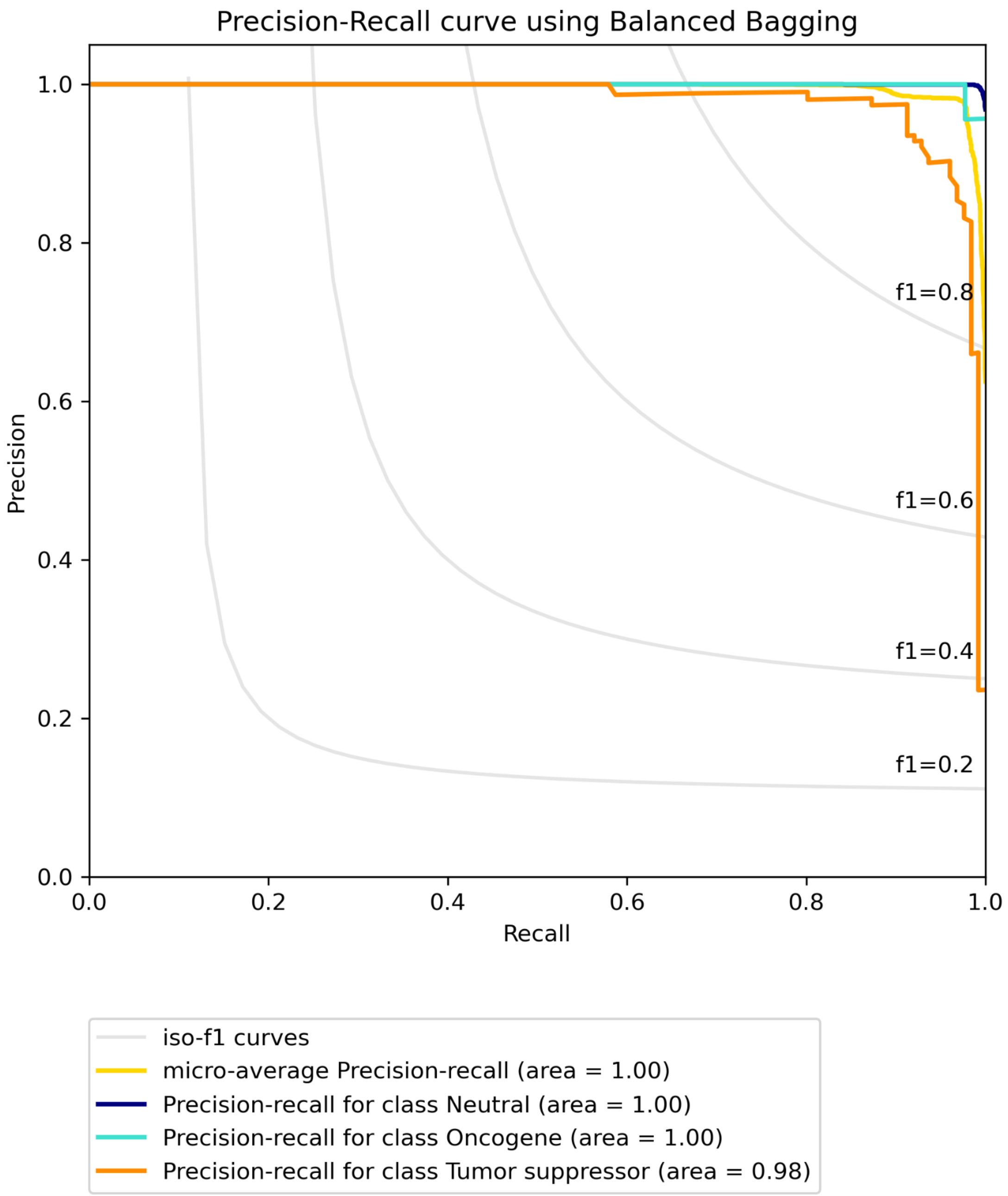


**Figure S39B. Precision-recall curve for LUAD using multi-omic data with Bailey *et al.* labels built on "all" features using Balanced bagging.**

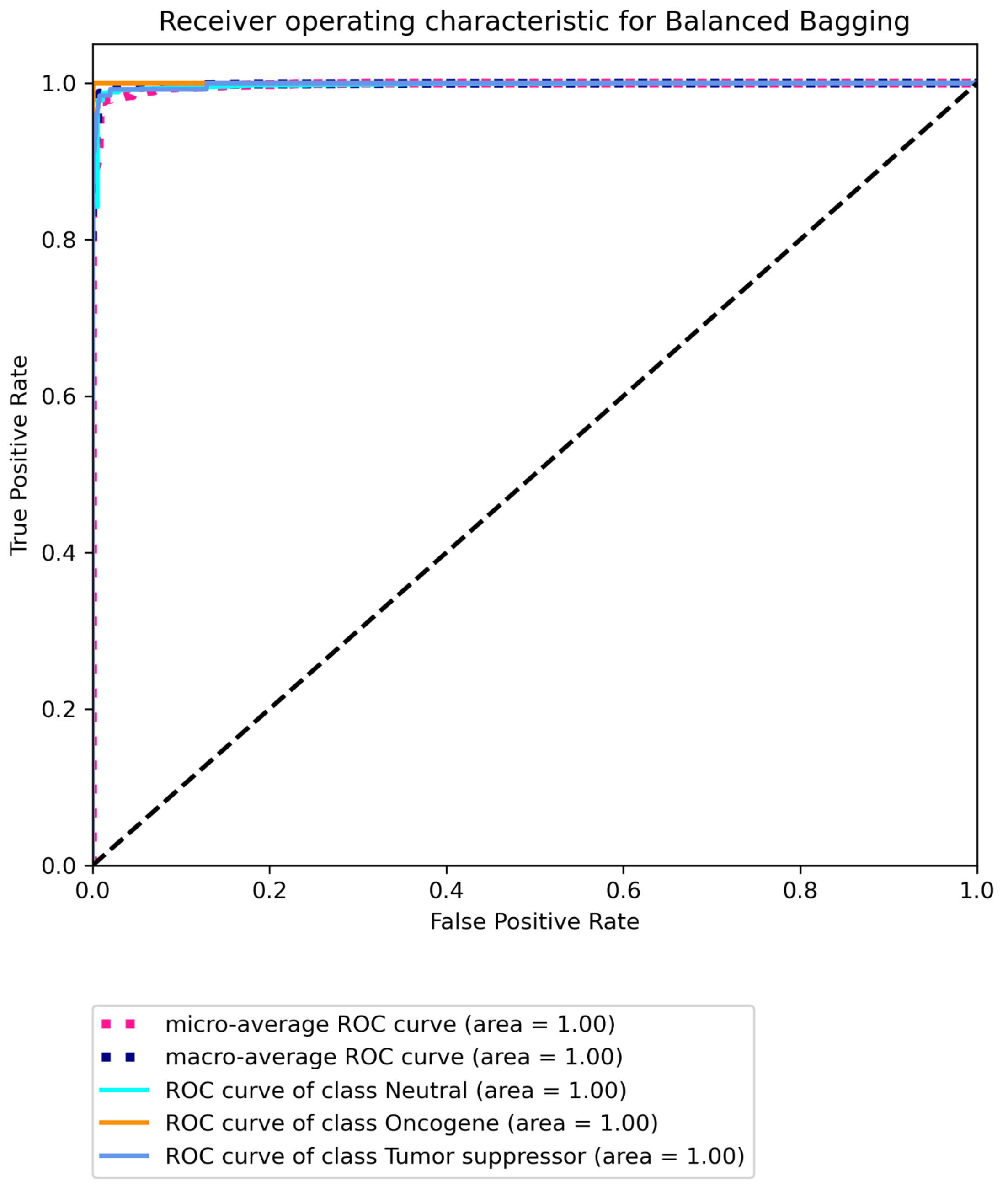


**Figure S39C Receiver operating characteristic for LUAD using multi-omic data with Bailey *et al.* labels built on "all" features using Balanced bagging.**

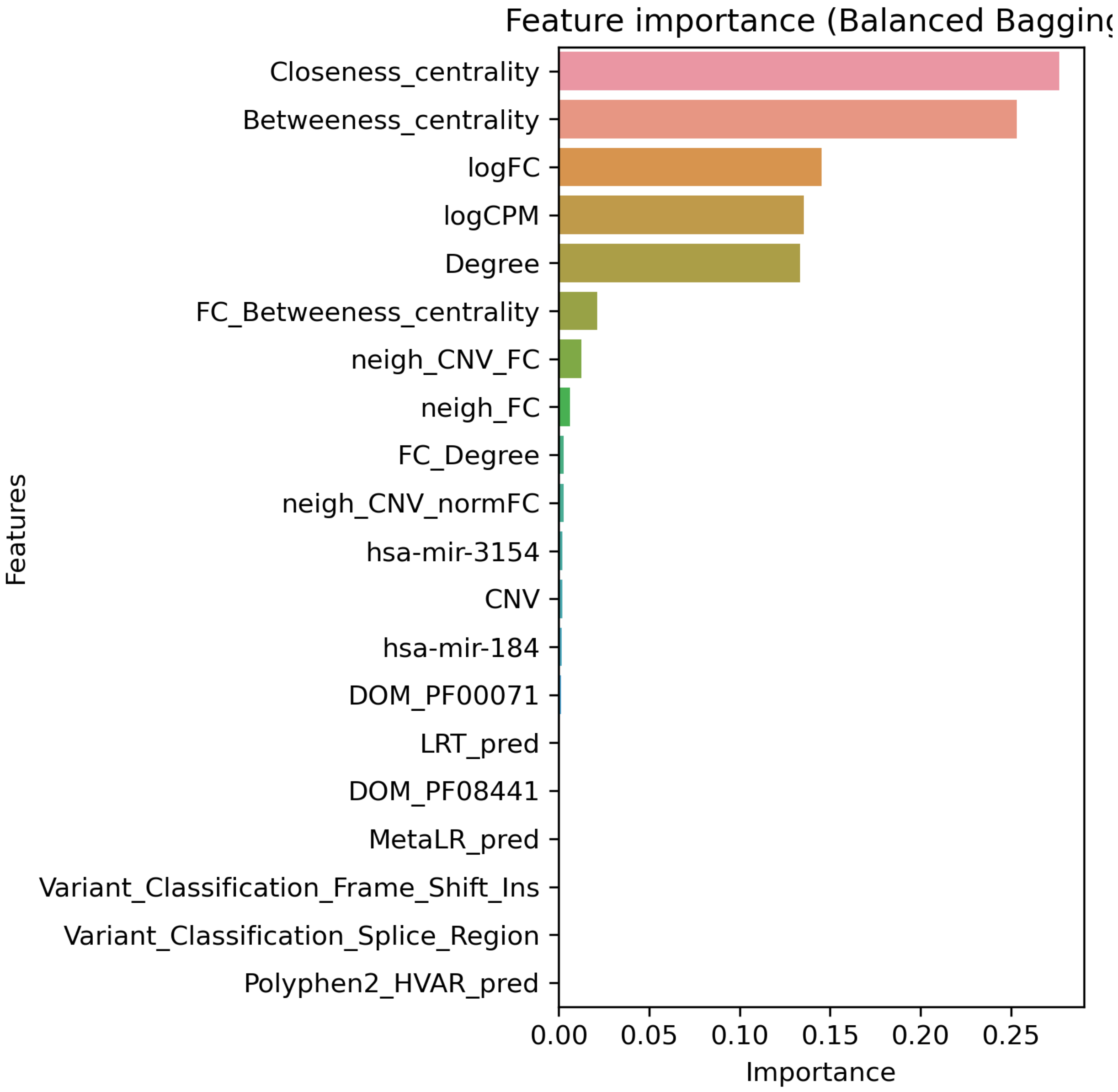


**Figure S40A. Top 20 features for LUAD using multi-omic data with Bailey *et al.* labels built on “some” features using Balanced bagging.**

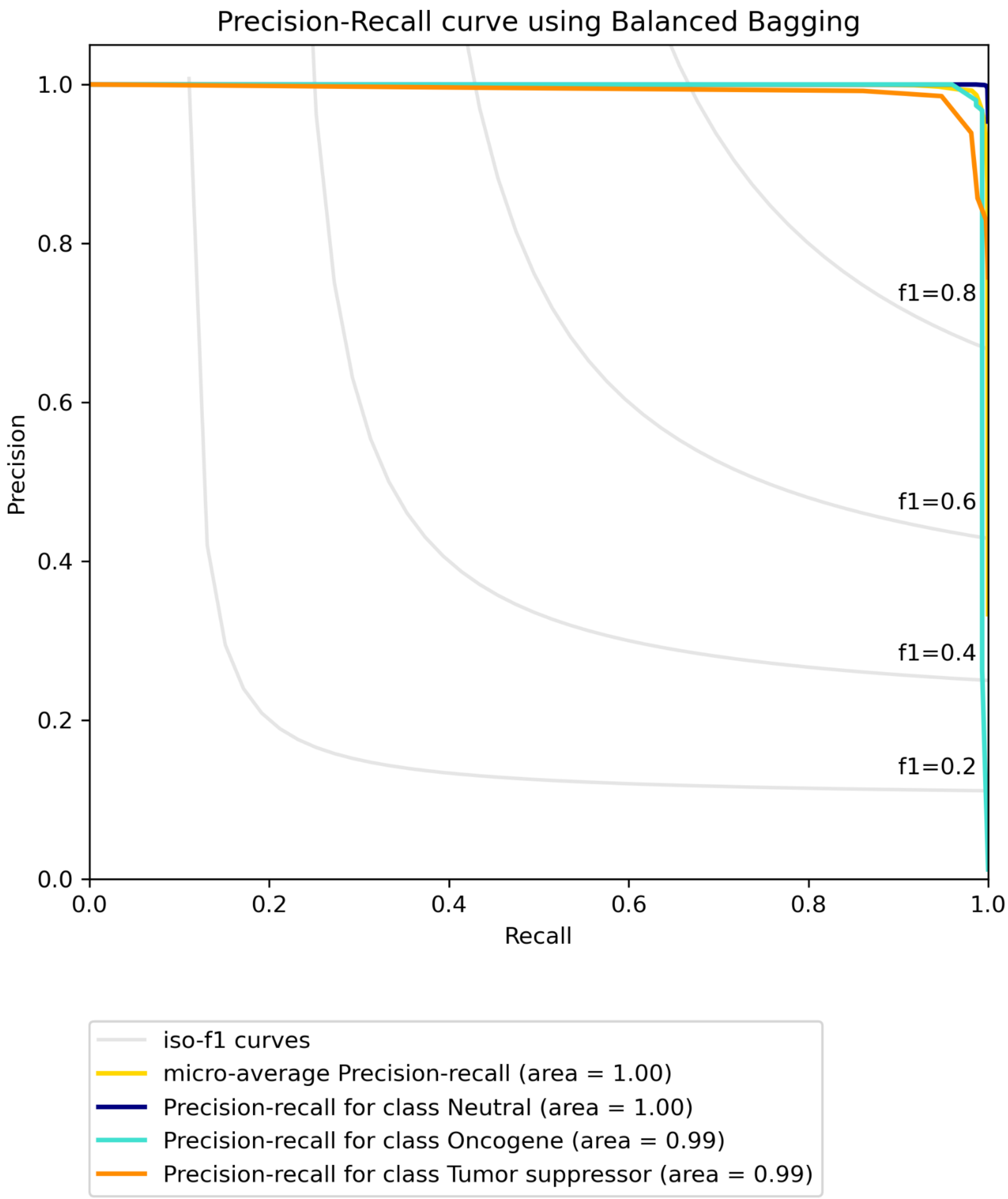


**Figure S40B. Precision-recall curve for LUAD using multi-omic data with Bailey *et al.* labels built on "some" features using Balanced bagging.**

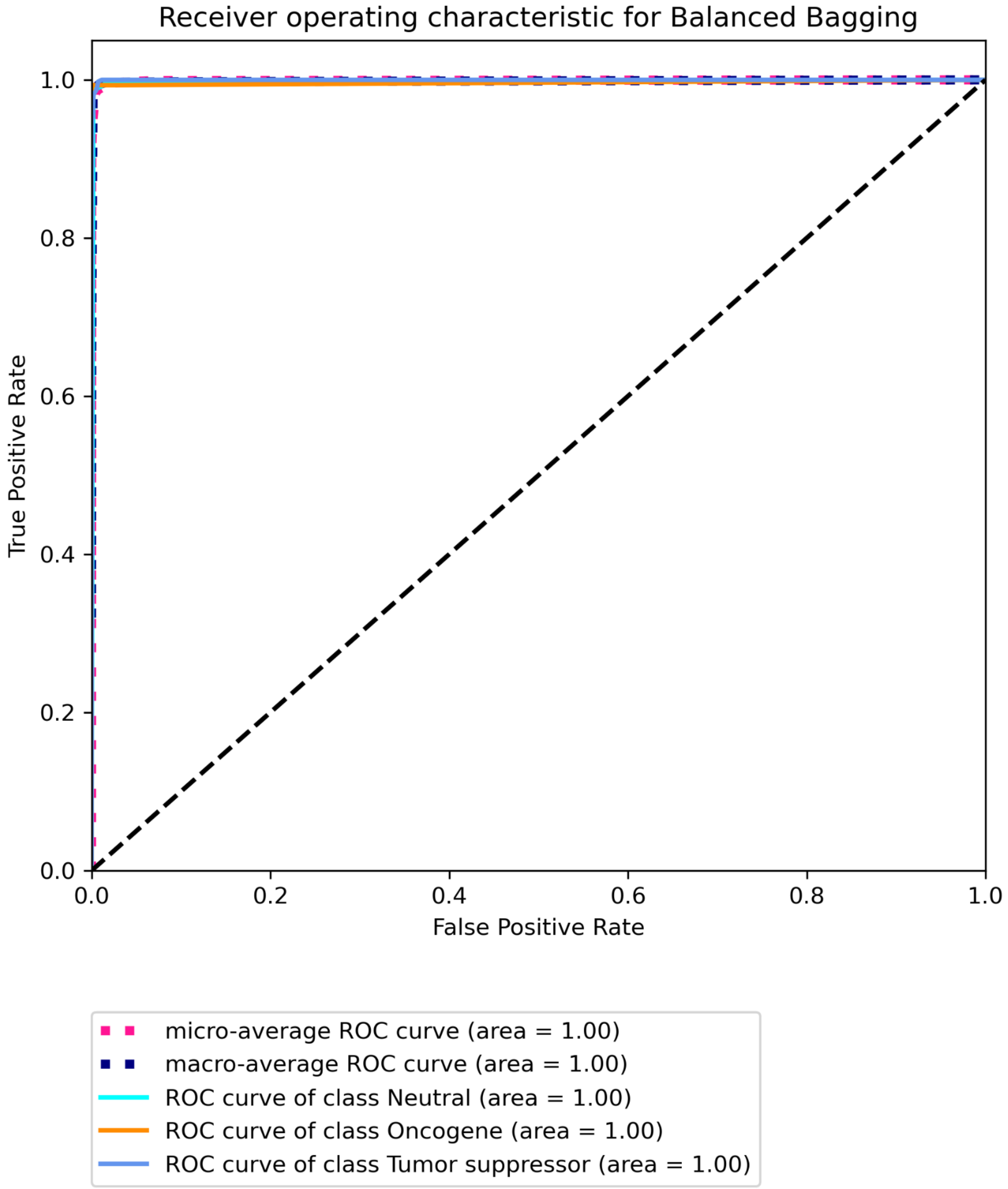


**Figure S40C Receiver operating characteristic for LUAD using multi-omic data with Bailey *et al.* labels built on "some" features using Balanced bagging.**

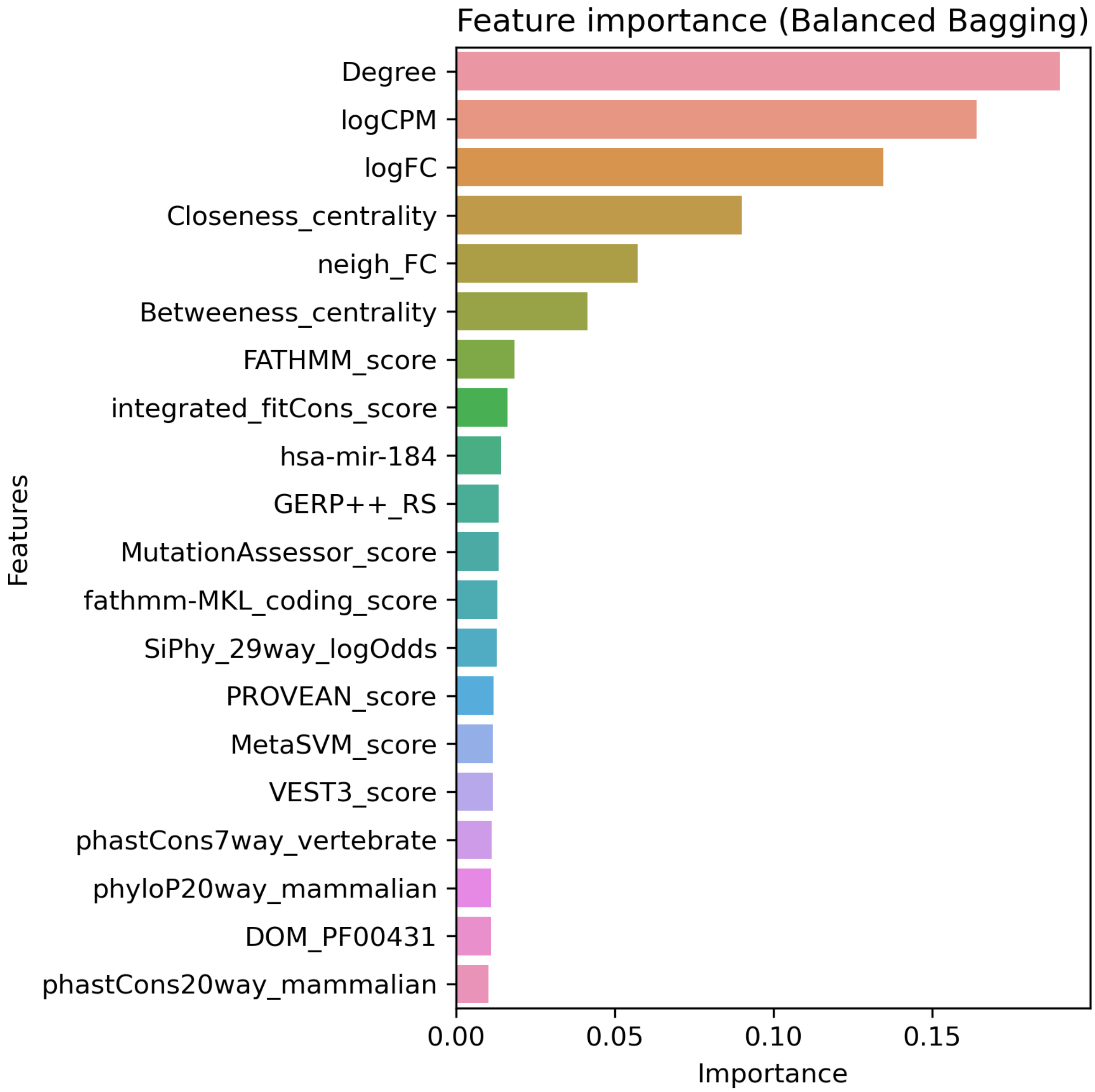


**Figure S41A. Top 20 features for LUAD using multi-omic data with CGC labels built on "all" features using Balanced bagging.**

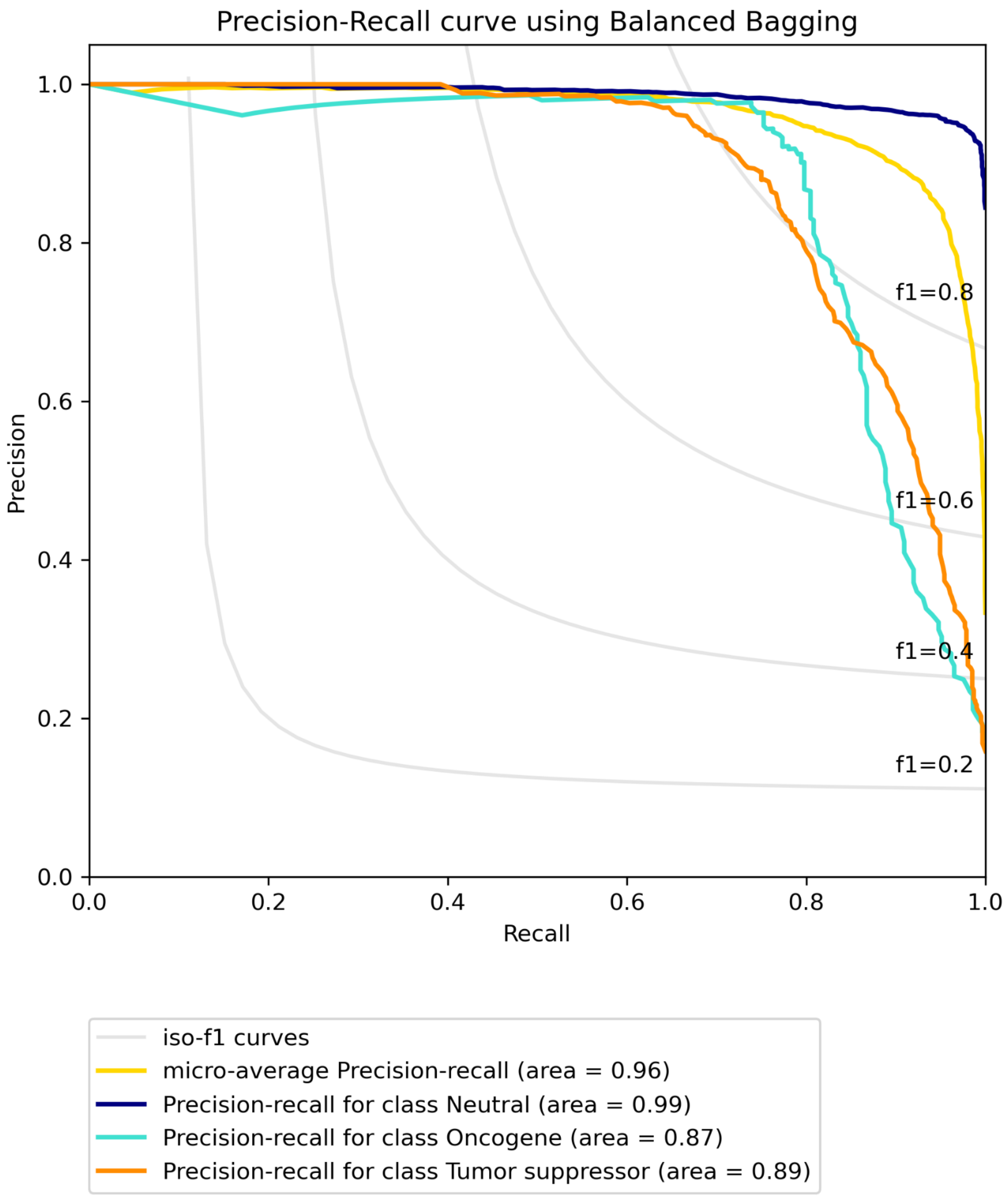


**Figure S41B. Precision-recall curve for LUAD using multi-omic data with CGC labels built on "all" features using Balanced bagging.**

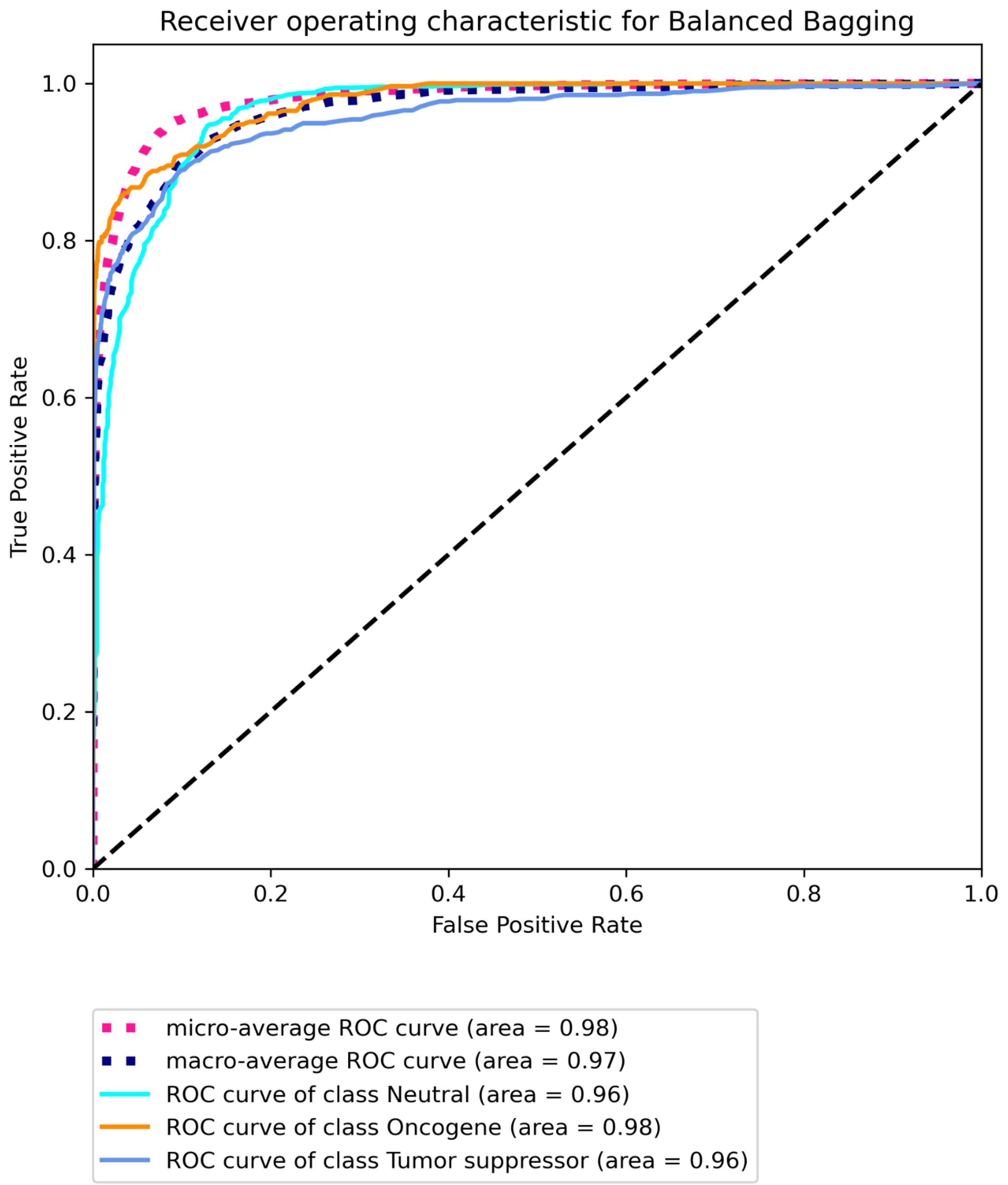


**Figure S41C. Receiver operating characteristic for LUAD using multi-omic data with CGC labels built on "all" features using Balanced Bagging.**

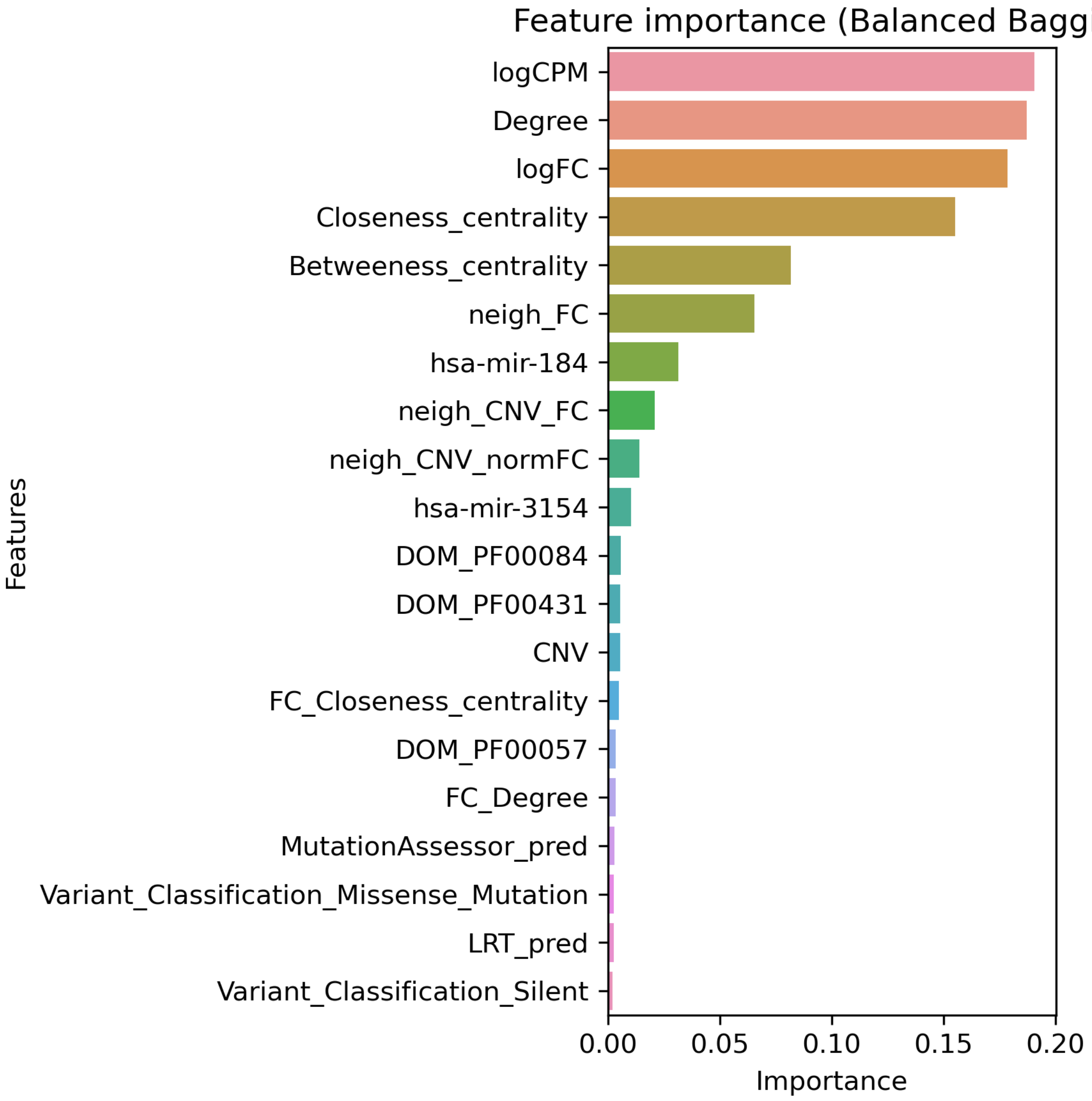


**Figure S42A. Top 20 features for LUAD using multi-omic data with CGC labels built on "some" features using Balanced Bagging.**

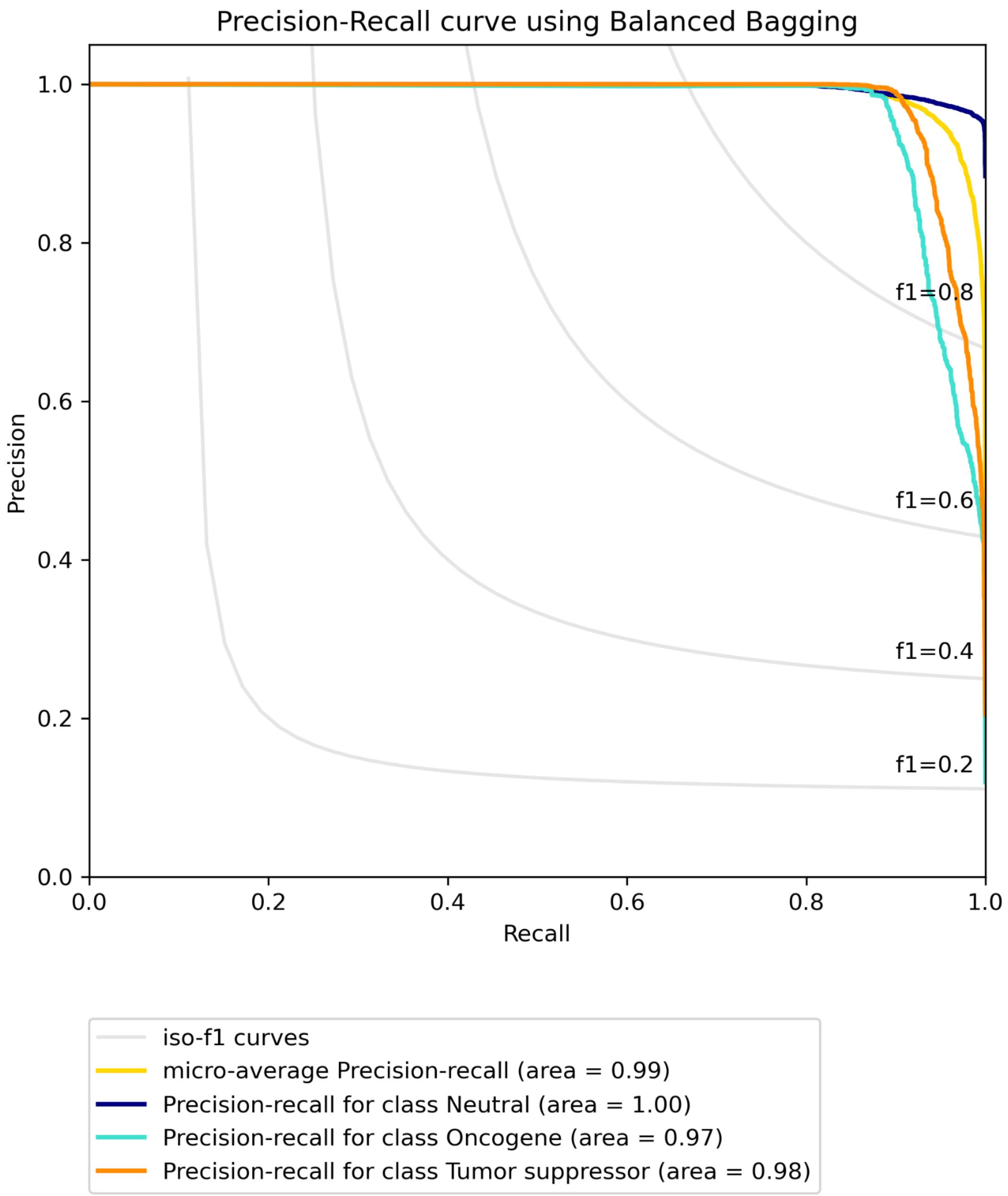


**Figure S42B. Precision-recall curve for LUAD using multi-omic data with CGC labels built on "some" features using Balanced Bagging.**

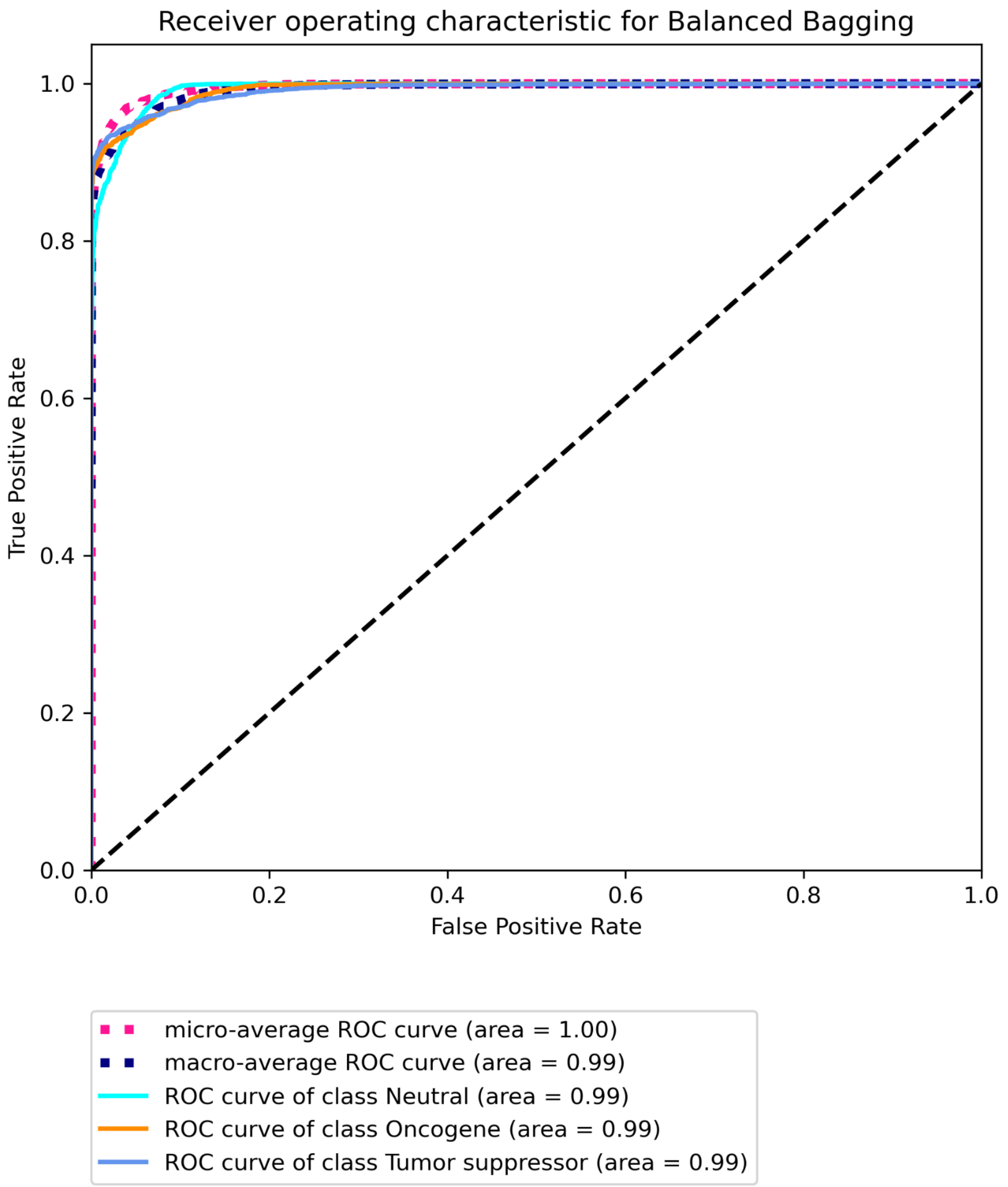


**Figure S42C. Receiver operating characteristic for LUAD using multi-omic data with CGC labels built on "some" features using Balanced Bagging.**

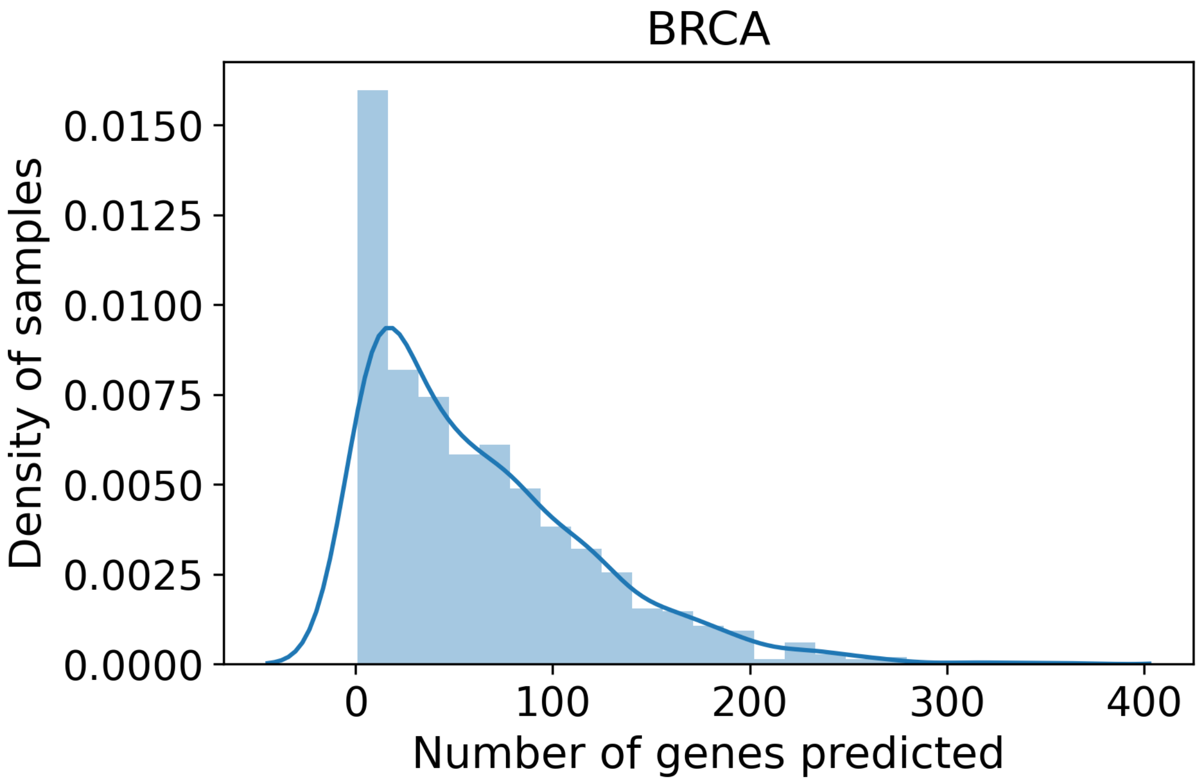


**Figure S43. Distribution of genes predicted for samples in BRCA.**

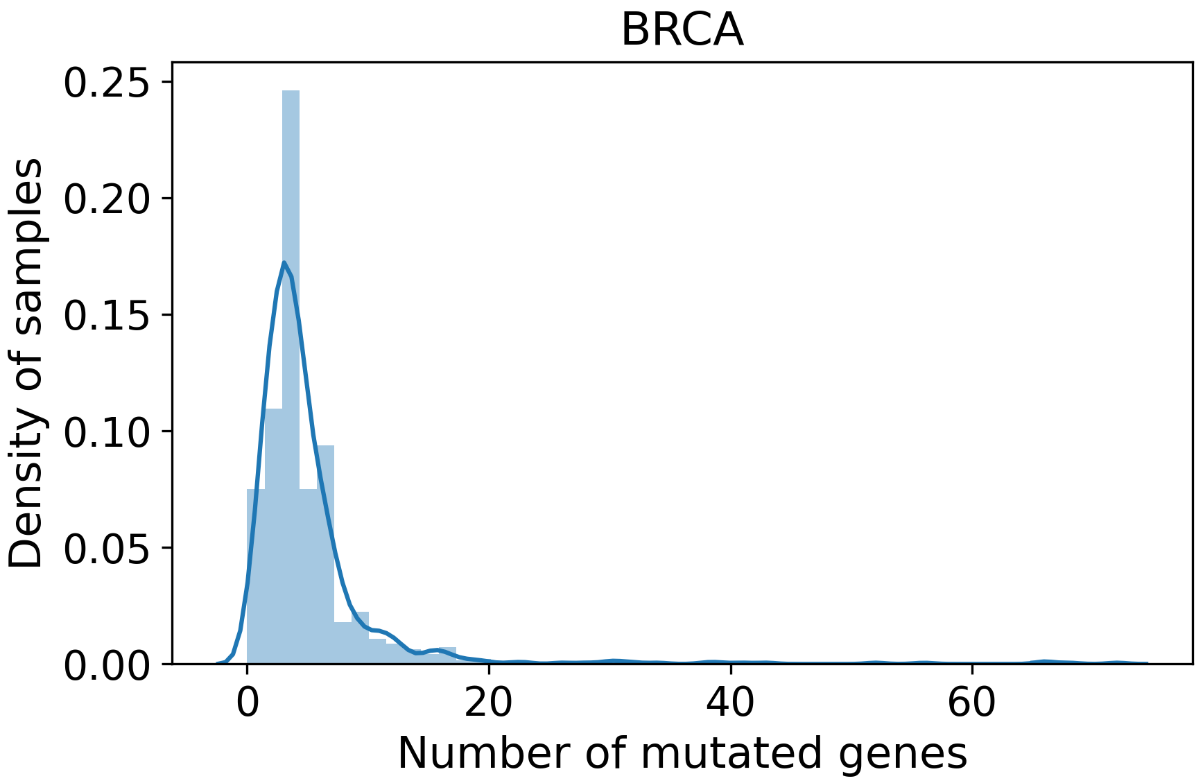


**Figure S44. Distribution of mutated genes predicted for samples in BRCA.**

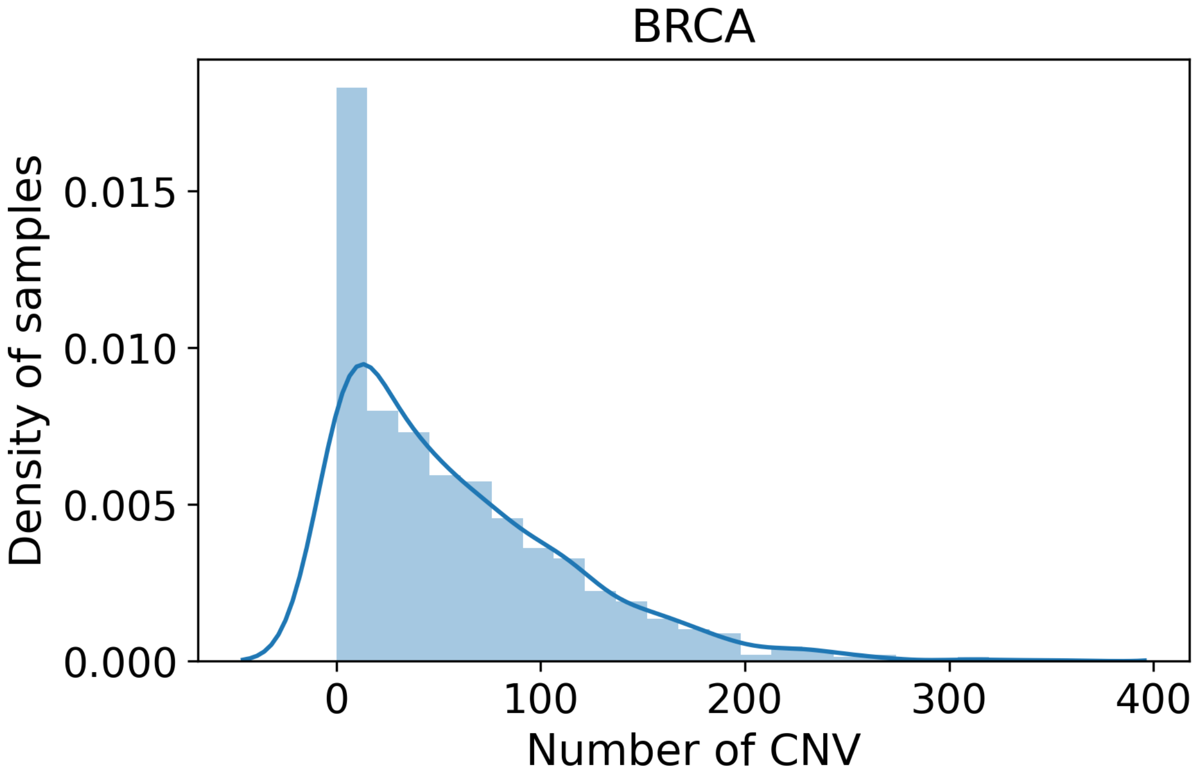


**Figure S45. Distribution of CNV altered genes predicted for samples in BRCA.**

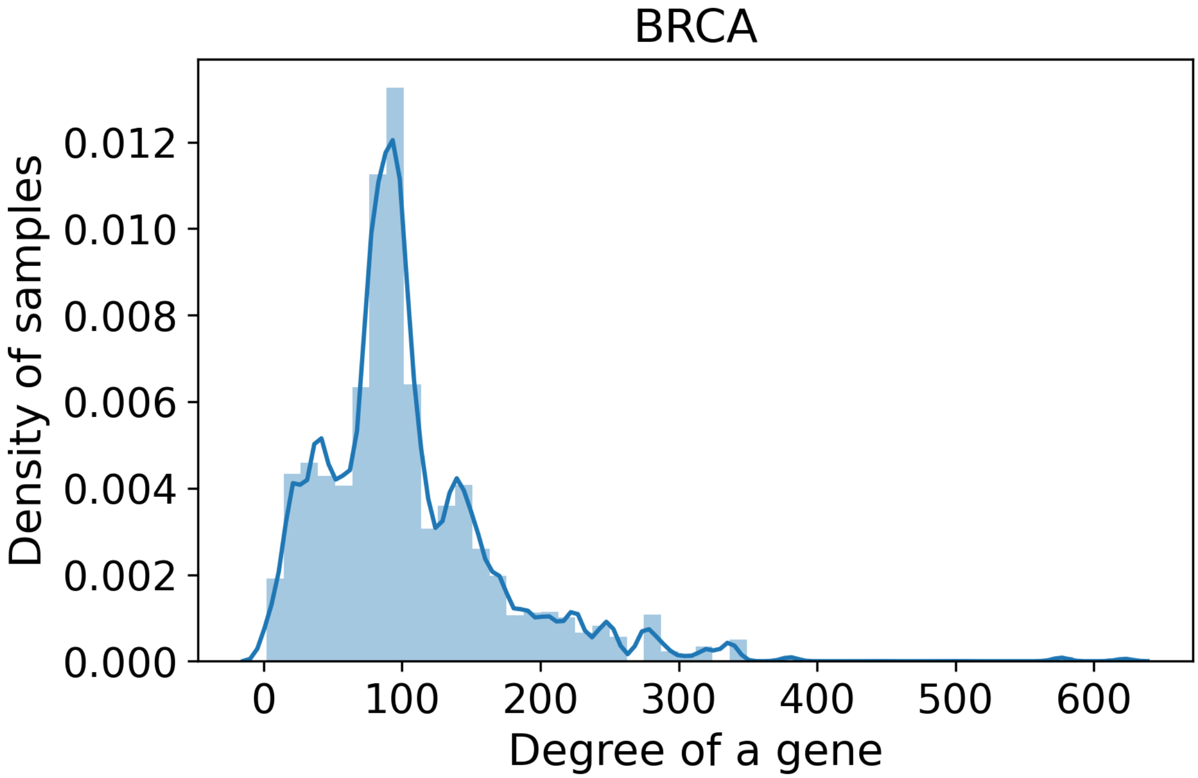


**Figure S46. Distribution of degree of genes predicted for samples in BRCA.**

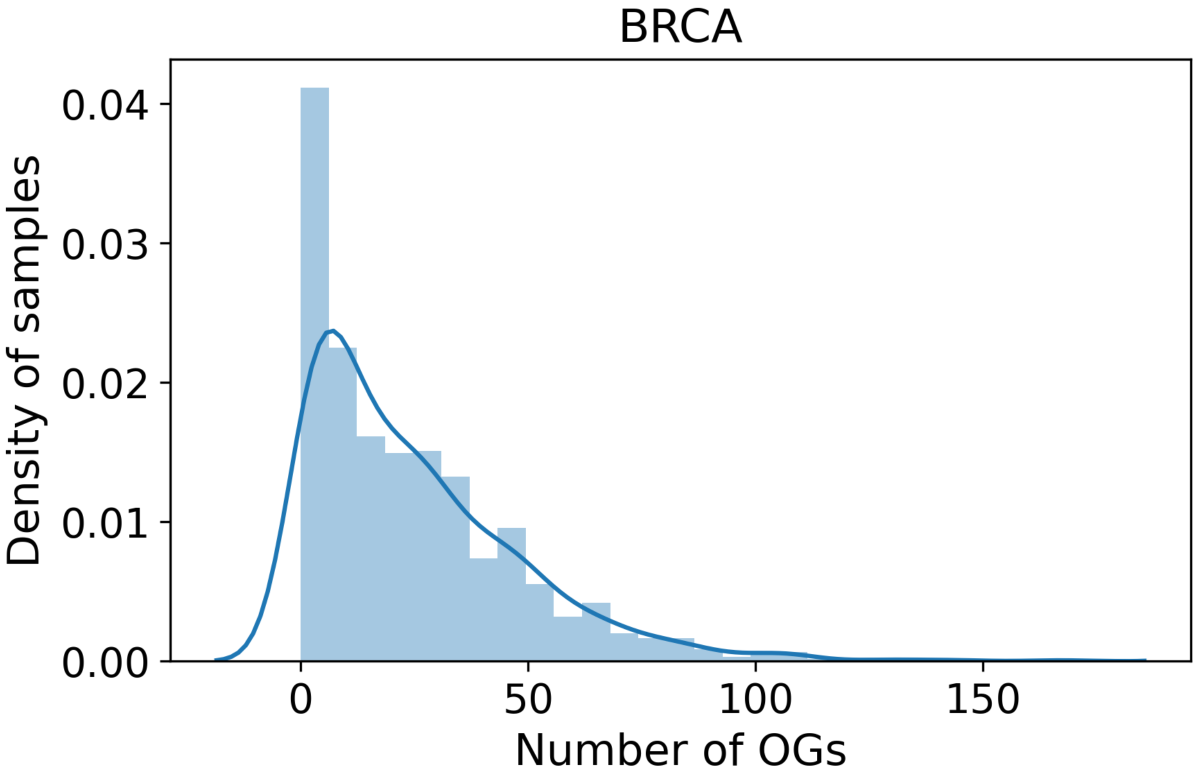


**Figure S47. Distribution of predicted oncogenes for samples in BRCA.**

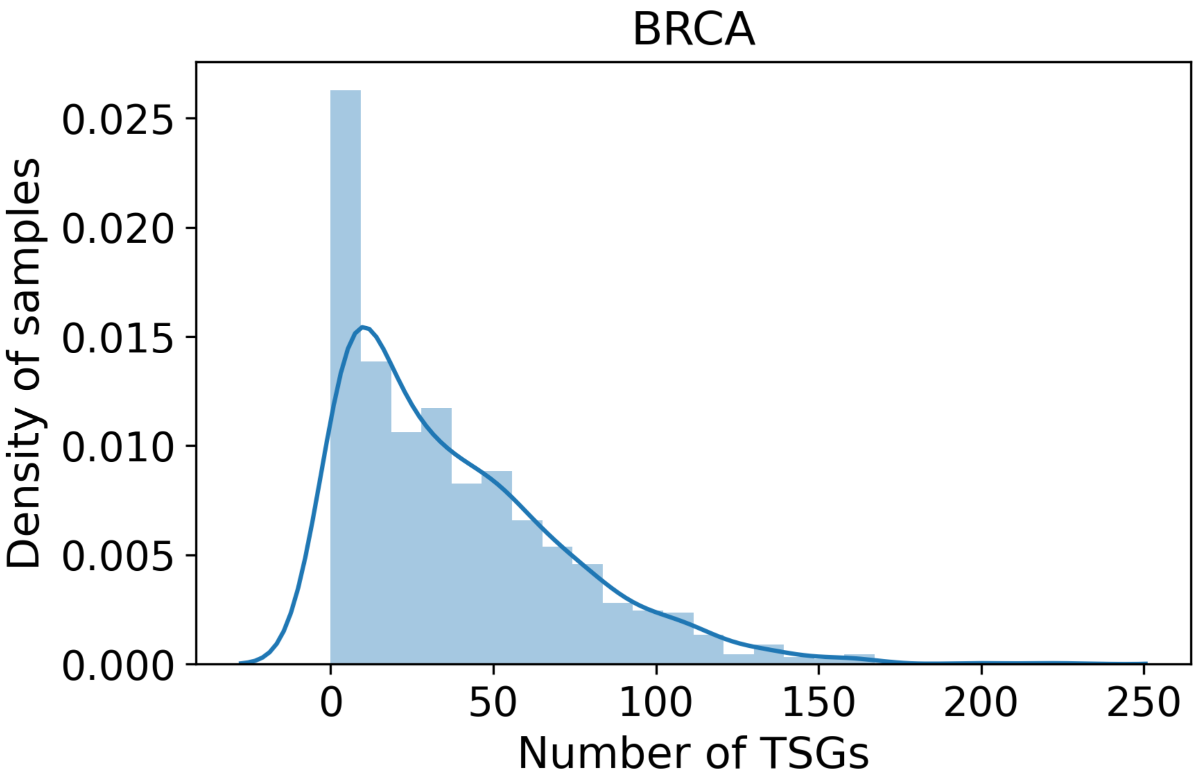


**Figure S48. Distribution of predicted tumour suppressor genes in BRCA.**

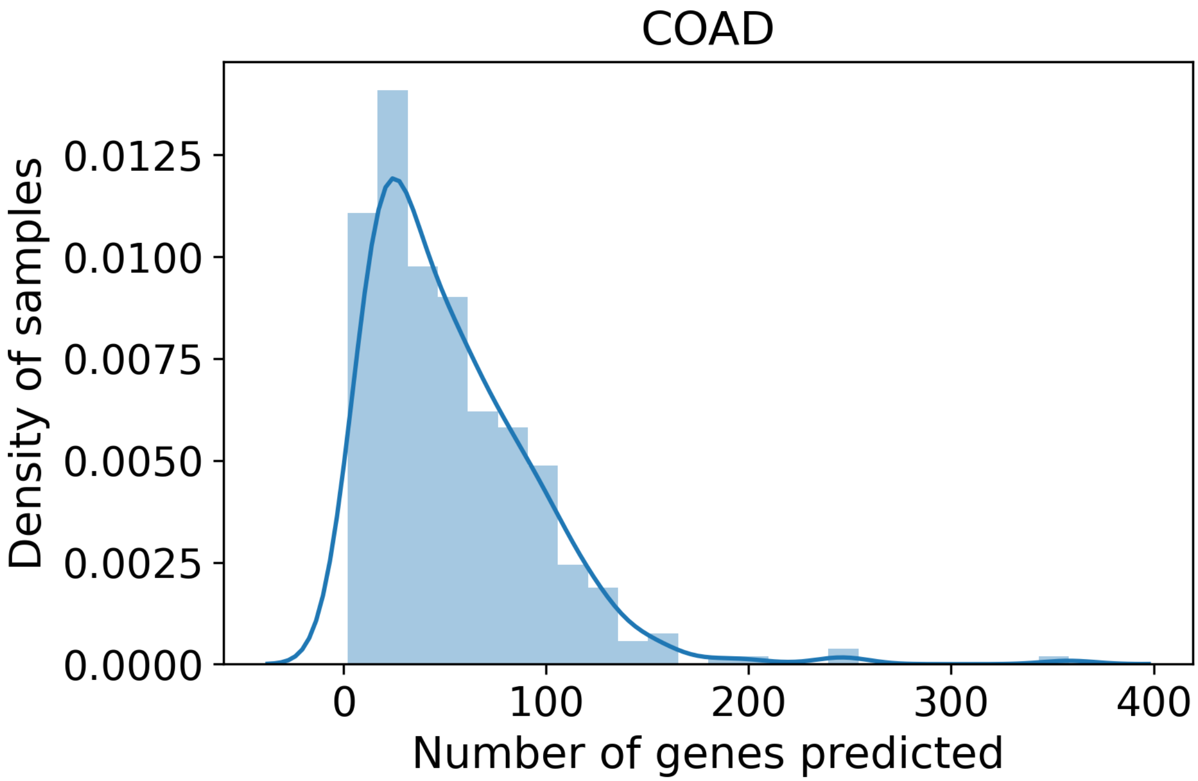


**Figure S49. Distribution of genes predicted for samples in COAD.**

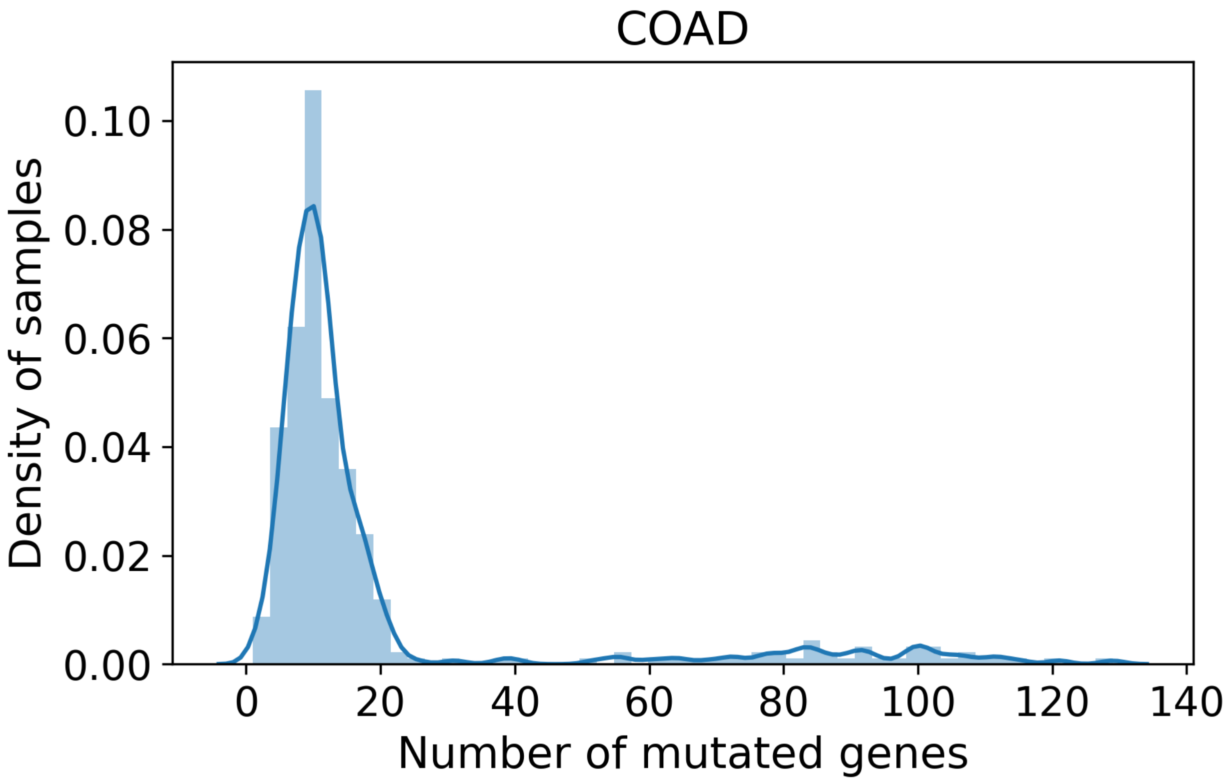


**Figure S50. Distribution of mutated genes predicted for samples in COAD.**

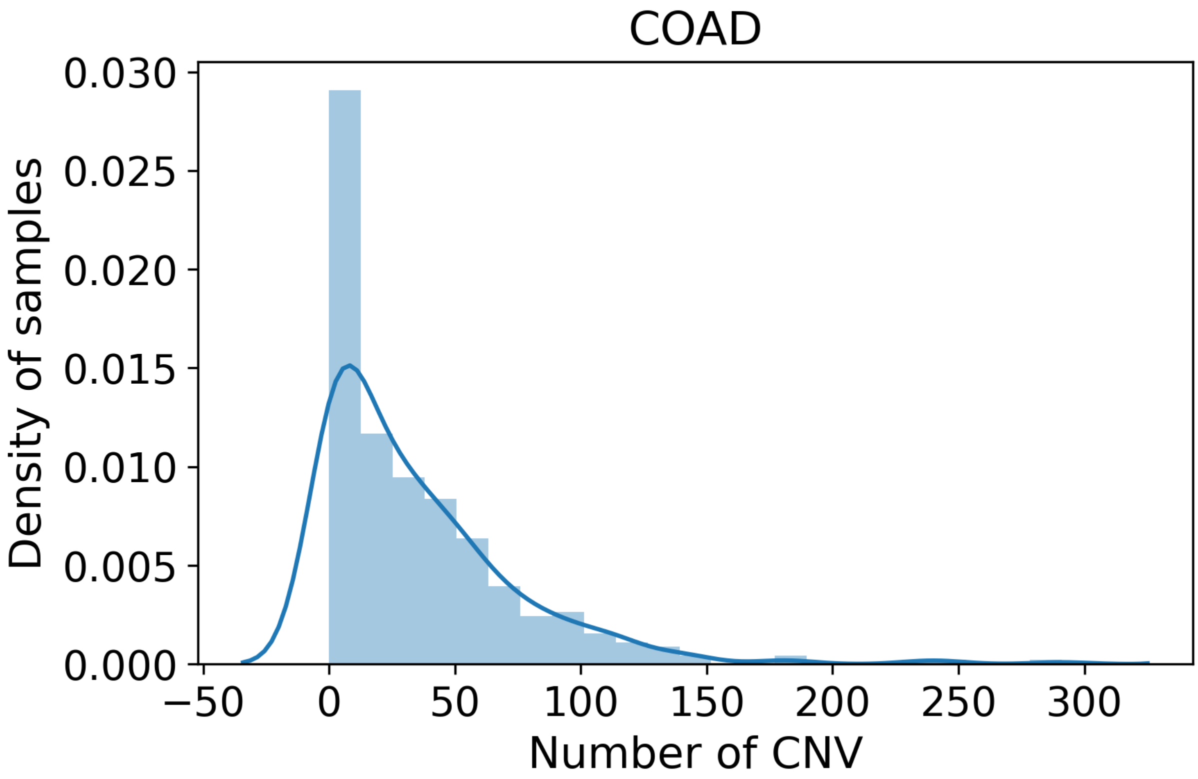


**Figure S51. Distribution of CNV altered genes predicted for samples in COAD.**

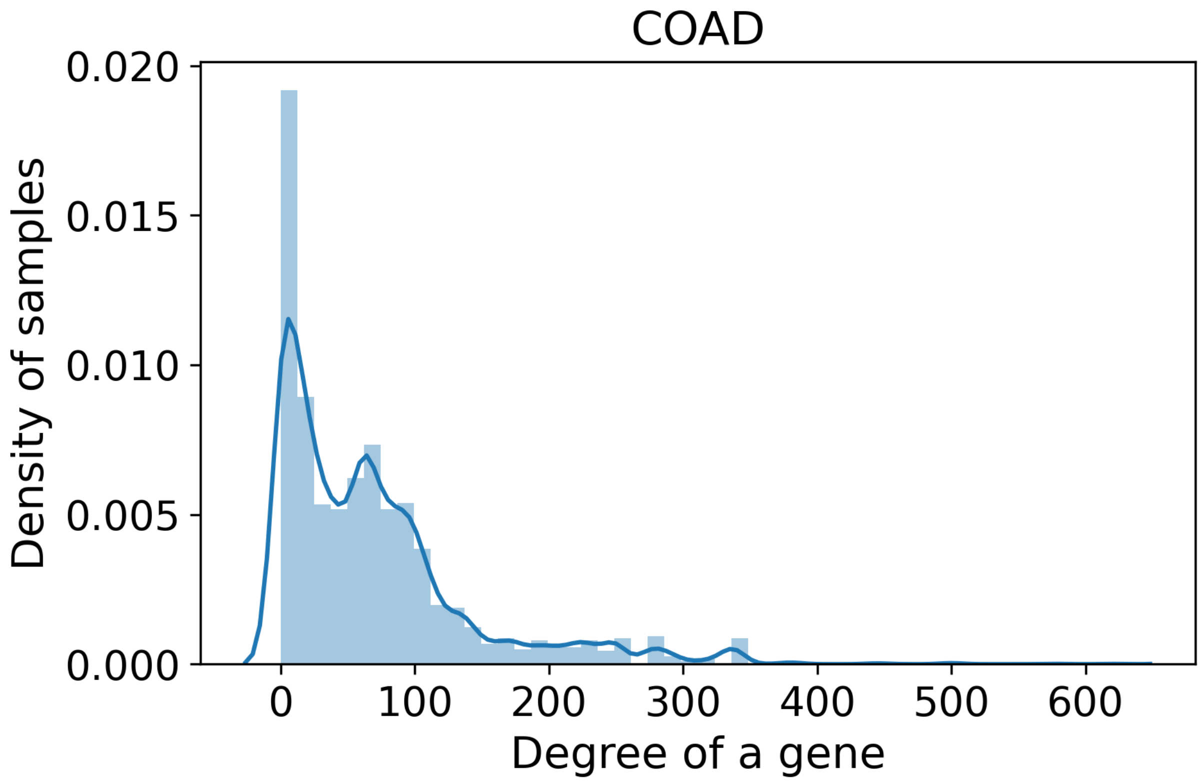


**Figure S52. Distribution of degree of genes predicted for samples in COAD.**

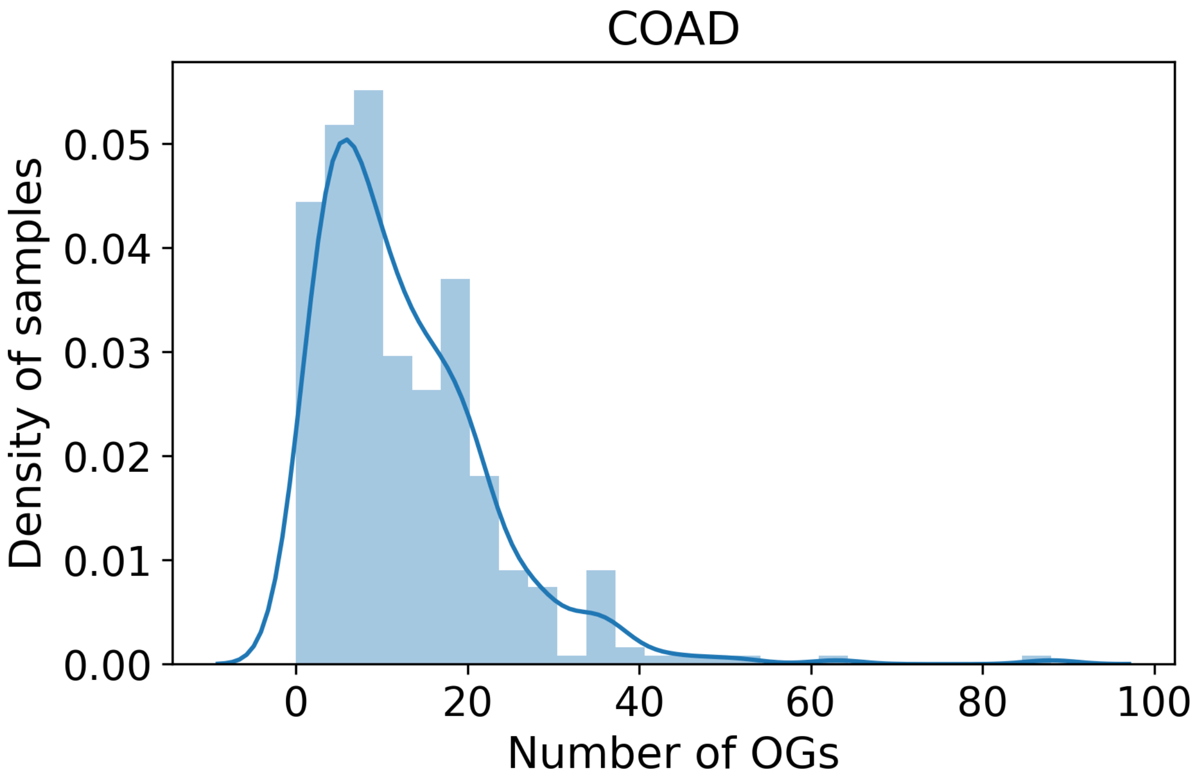


**Figure S53. Distribution of predicted oncogenes for samples in COAD.**

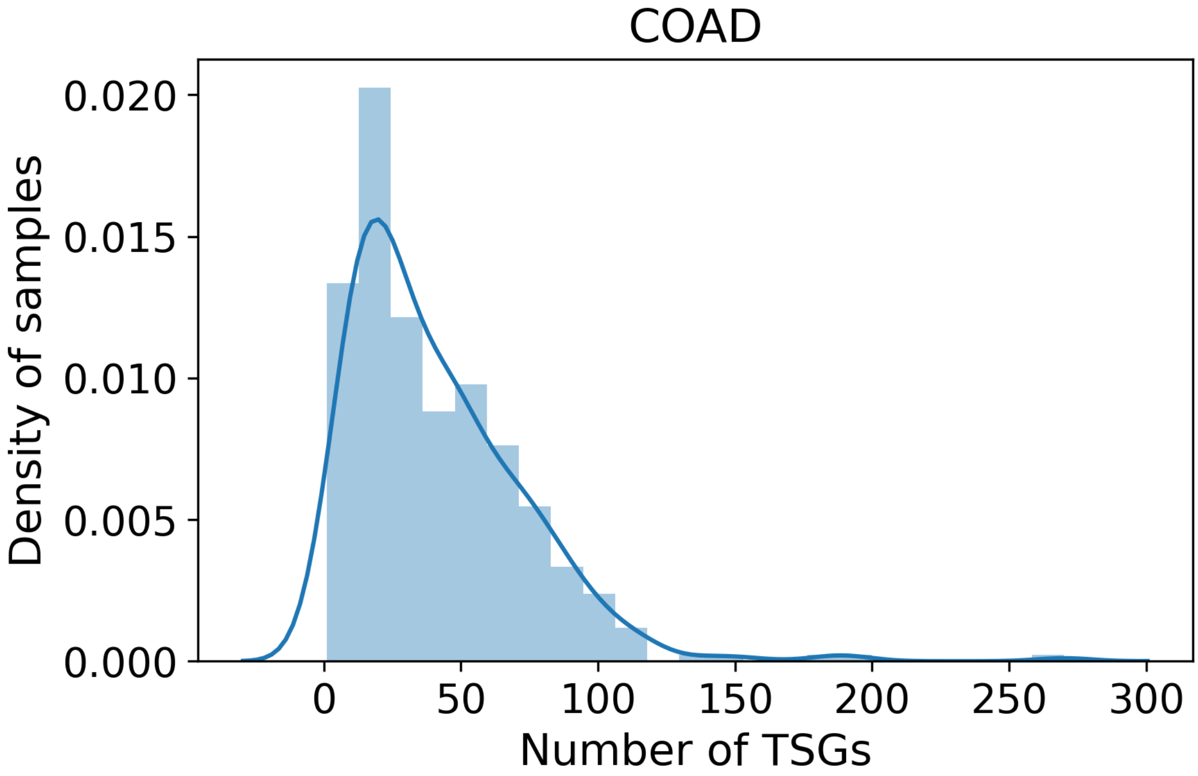


**Figure S54. Distribution of predicted tumour suppressor genes in COAD.**

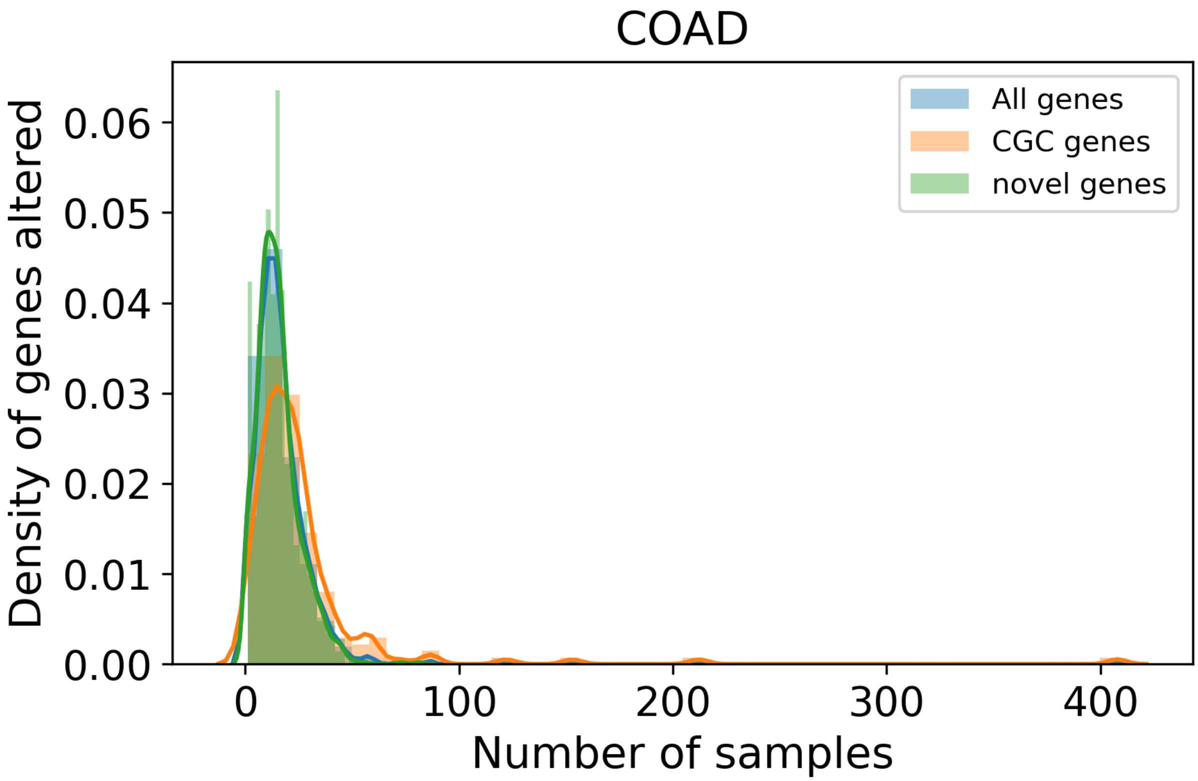


**Figure S55. Distribution of samples for genes predicted as driver in COAD.**

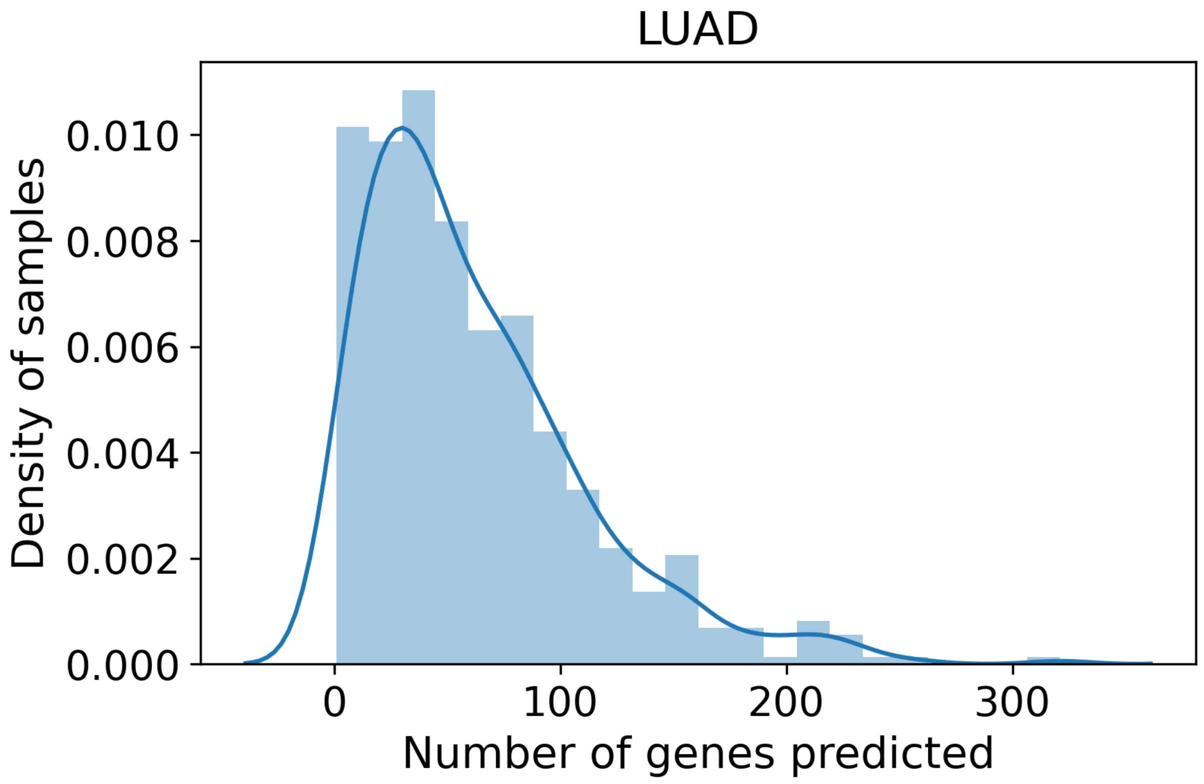


**Figure S56. Distribution of genes predicted for samples in LUAD.**

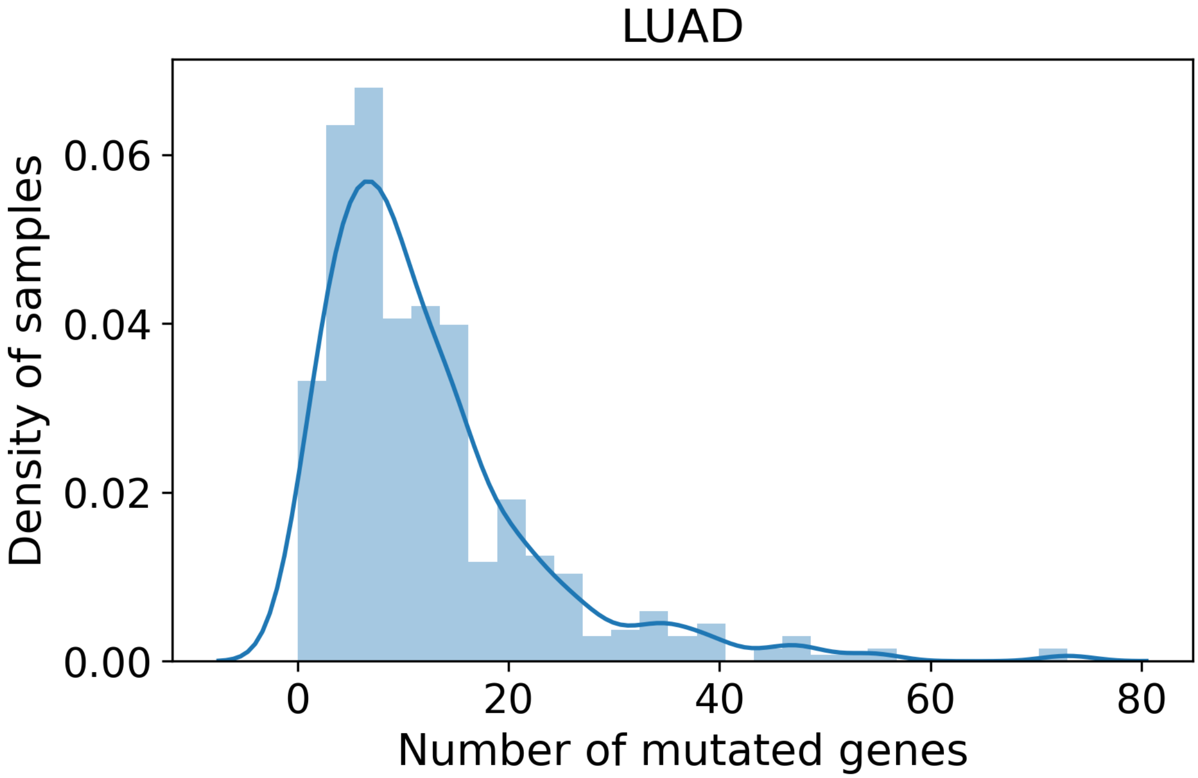


**Figure S57. Distribution of mutated genes predicted for samples in LUAD.**

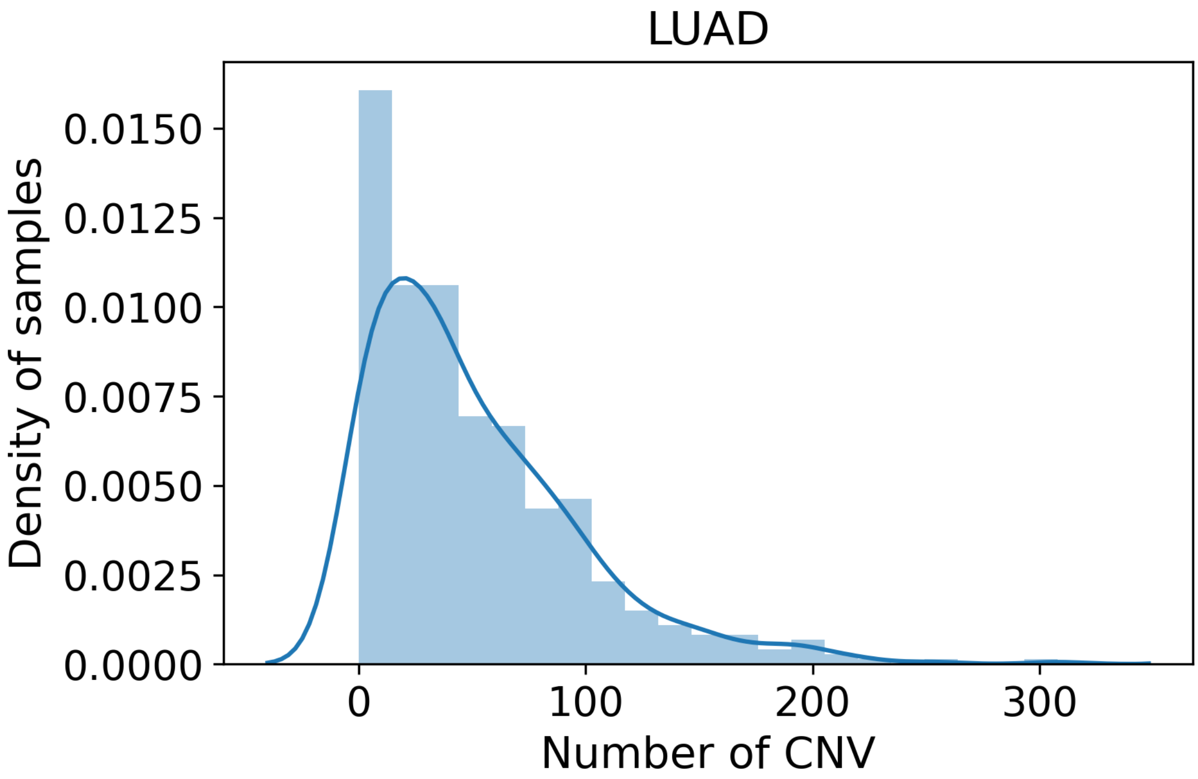


**Figure S58. Distribution of CNV altered genes predicted for samples in LUAD.**

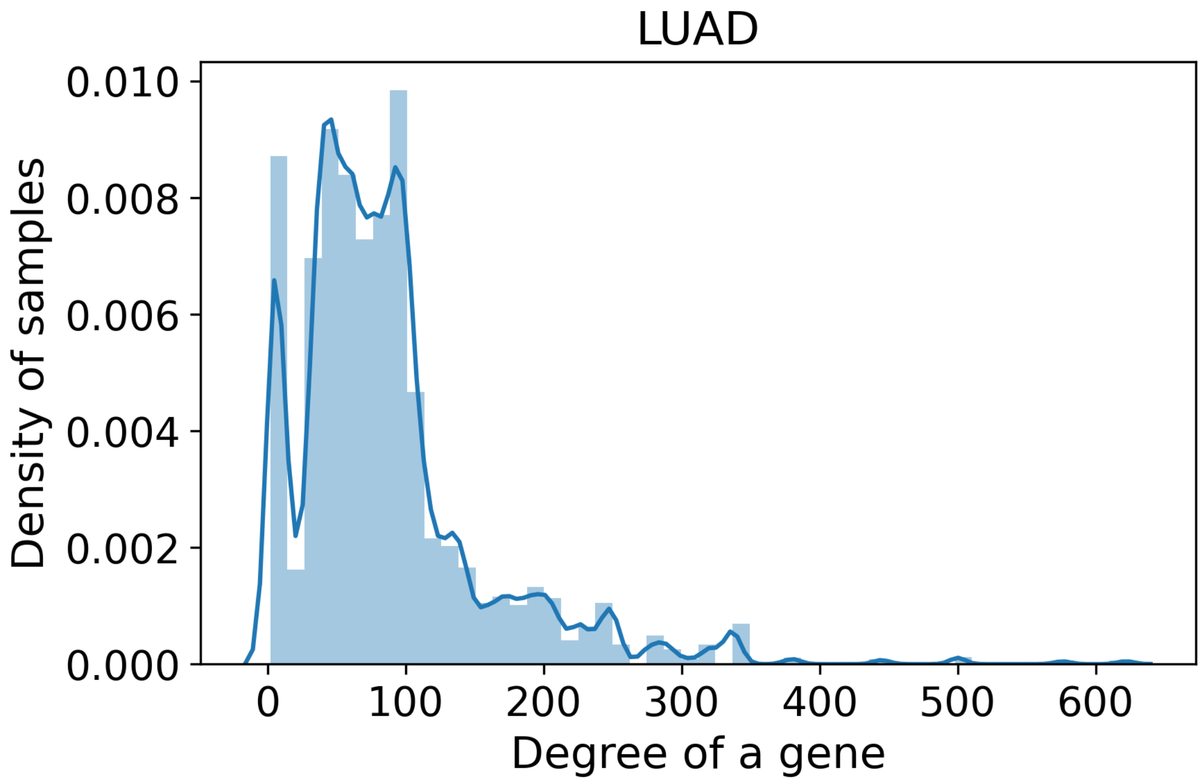


**Figure S59. Distribution of degree of genes predicted for samples in LUAD.**

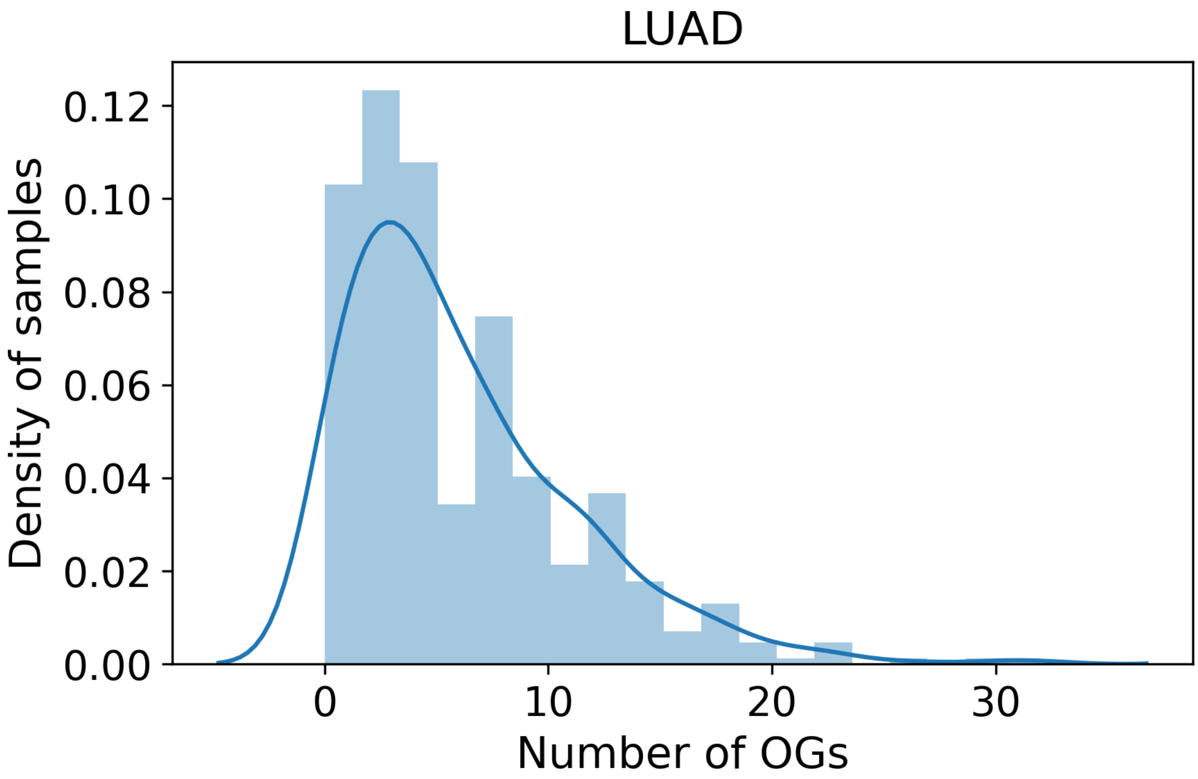


**Figure S60. Distribution of predicted oncogenes for samples in LUAD.**

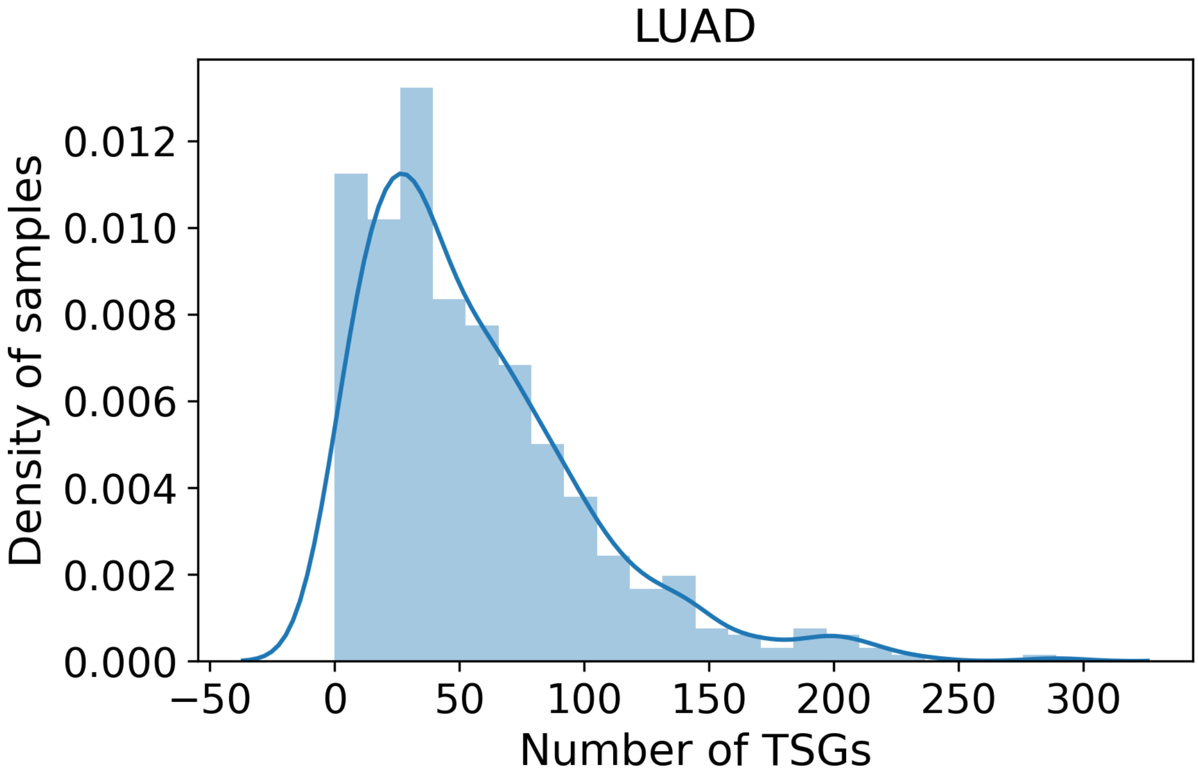


**Figure S61. Distribution of predicted tumour suppressor genes in LUAD.**

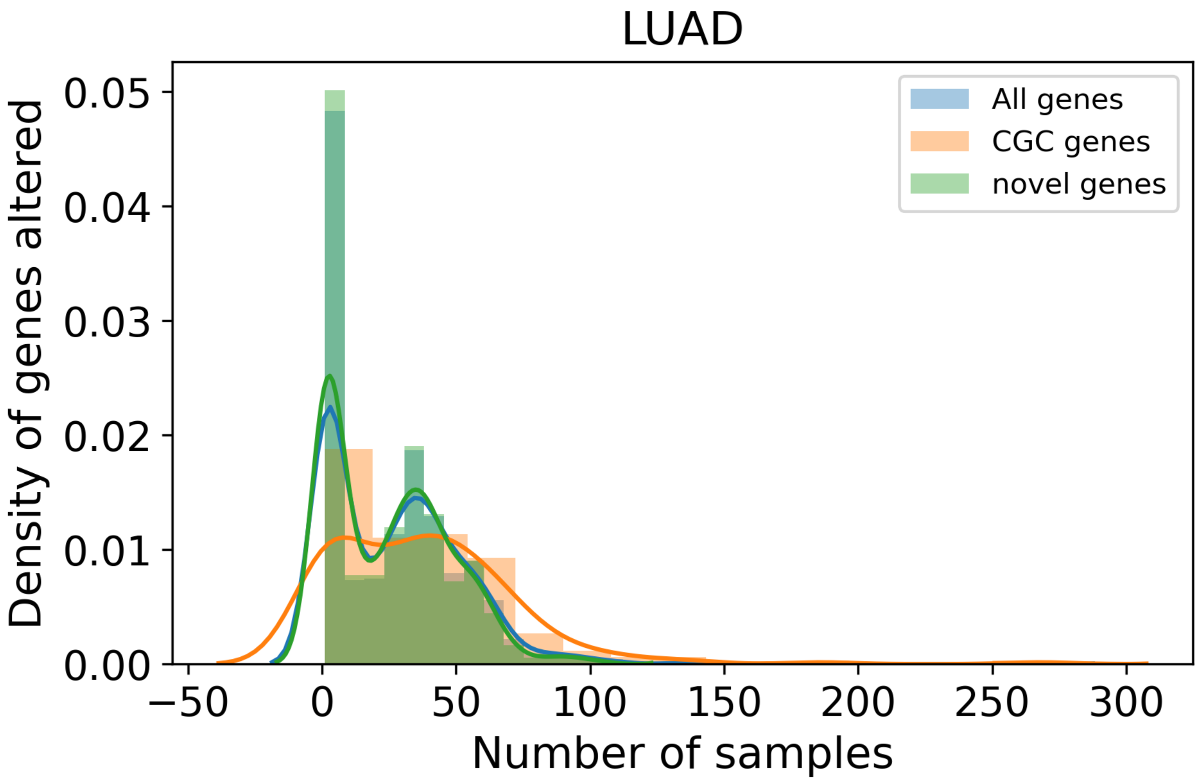


**Figure S62. Distribution of samples for genes predicted as driver in LUAD.**
